# Developmentally regulated activation of defense allows for rapid inhibition of infection in age-related resistance to *Phytophthora capsici* in cucumber fruit

**DOI:** 10.1101/2020.08.14.251603

**Authors:** Ben N. Mansfeld, Marivi Colle, Chunqiu Zhang, Ying-Chen Lin, Rebecca Grumet

**Affiliations:** Graduate Program in Plant Breeding, Genetics and Biotechnology and Department of Horticulture, Michigan State University, East Lansing, MI, USA; Key Laboratory of Biology and Genetic Improvement of Horticultural Crops (North China), Beijing Key Laboratory of Vegetable Germplasm Improvement, National Engineering Research Center for Vegetables, Beijing Academy of Agriculture and Forestry Sciences, Beijing, China

**Keywords:** Age-related resistance, ontogenic resistance, cucumber, *Phytophthora capsici*, plant defense, transcriptome, co-expression networks

## Abstract

**Background:** Age-related resistance (ARR) is a developmentally regulated phenomenon conferring resistance to pathogens or pests. Although ARR has been observed in several host-pathogen systems, the underlying mechanisms are largely uncharacterized. In cucumber, rapidly growing fruit are highly susceptible to *Phytophthora capsici* but become resistant as they complete exponential growth. We previously demonstrated that ARR is associated with the fruit peel and identified gene expression and metabolomic changes potentially functioning as preformed defenses.

**Results:** Here, we compare the response to infection in fruit at resistant and susceptible ages using microscopy, quantitative bioassays, and weighted gene co-expression analyses. We observed strong transcriptional changes unique to resistant aged fruit 2-4 hours post inoculation (hpi). Microscopy and bioassays confirmed this early response, with evidence of pathogen death and infection failure as early as 4 hpi and cessation of pathogen growth by 8-10 hpi. Expression analyses identified candidate genes involved in conferring the rapid response including those encoding transcription factors, hormone signaling pathways, and defenses such as reactive oxygen species metabolism and phenylpropanoid biosynthesis.

**Conclusion:** The early pathogen death and rapid defense response in resistant-aged fruit provide insight into potential mechanisms for ARR, implicating both pre-formed biochemical defenses and developmentally regulated capacity for pathogen recognition as key factors shaping age-related resistance.

## Background

Ontogenic, developmental, or age-related resistance (ARR), wherein plants or plant organs transition from susceptibility to resistance as a result of developmental changes [1–3], has been described in several different plant-pathogen systems and in crops such as pepper, grape, rice, wheat, and several cucurbit crops [4–8]. While providing protection in agricultural systems and potentially playing an important role in the evolution and optimization of host resistance [9], the molecular mechanisms controlling these resistances are not well understood.

Evidence from various systems suggests possible physical, chemical, or physiological changes that may result from age-dependent, non-pathogen specific investment in defense such as cell wall modifications, production of anti-microbial phytoanticipins, or altered hormone balance [2, 9, 10]. There are also some examples where ARR may result from developmentally regulated expression of a pathogen receptor. In rice, a developmental increase in expression of leucine-rich repeat (LRR)-kinase type genes, *Xa3/Xa26* and *Xa21*, that peaks at the maximum-tillering stage of growth, confers ARR to bacterial blight [7, 11]. Recently, in Arabidopsis, transcriptional control of the canonical immune receptor FLS2 was also shown to regulate ontogenic resistance to *Pseudomonas syringae* [12]. Thus, in these examples, a newly acquired ability to perceive the pathogen allows for an induced resistance response at the resistant age.

One model system for organ specific ARR is cucumber fruit rot by the oomycete *Phytophthora capsici* [13–15]. This soilborne hemibiotroph is a pathogen of many agriculturally important crops including numerous solanaceous and cucurbit species [16]. Infection is initiated by means of biflagellate zoospores which arrive via water from rain or irrigation [17]. Upon reaching the host target tissue, zoospores encyst, lose their flagella and germinate forming germination tubes [18]. The germination tubes penetrate the plant surface using appressoria and continue growing hyphae. During the early, biotrophic stages of infection, haustoria are formed mediating direct interaction with the host cells and nutrient acquisition. Studies in tomato show that the pathogen then transitions to necrotrophy at approximately 48 hours post-inoculation (hpi) and can produce sporangia for asexual reproduction as soon as 72 hpi [18, 19]. The transcriptome of *P. capsici* infection has been described in tomato leaves using microarray technology [19]. Two major transcriptomic responses were identified: 1) at initial infection (8 hpi) at which host primary metabolism was downregulated and specialized metabolism was upregulated, and 2) at the transition to necrotrophy (48 hpi) when large scale transcriptional reprograming occurs in the host [19].

In cucumber, the primary targets of infection of *P. capsici* are the fruit, which display symptoms of rot and tissue collapse followed by appearance of white mycelia and sporangia. Testing of the cucumber germplasm collection found that very young fruit [less than 8 days post pollination (dpp)] from the great majority of cucumber accessions are highly susceptible [20]. The genetics of inheritance of young fruit resistance have not yet been determined, and to date, no stable resistance to this pathogen is deployed in cultivated cucumber. However, an ARR was observed in a few cucumber cultivars and plant introduction lines wherein fruit become resistant as they near the end of their exponential growth phase, at ∼12-15 days post-pollination (dpp) [13]. As with young fruit resistance, the genetics of ARR have not been determined. However, transcriptional analyses indicate that the end of exponential fruit growth is accompanied by downregulation of transcription factors and other genes involved in growth and by increased expression of defense related genes [14, 15, 21]. This transcriptional transition away from growth and towards defense, corresponding with the end of rapid growth, is congruent with current understanding of optimal defense programing and utilization of resources [22]. With growth largely completed, fruit can invest in defense to protect the developing seeds during the period of seed maturation that precedes the final transition to fruit ripening at ∼35 dpp.

Further studies showed that the cucumber fruit peel is important for ARR [14]. Excised peels exhibit equivalent responses (susceptible or resistant) as do intact whole fruit, and if the peel is wounded or removed from resistant-aged fruit the underlying tissue is highly susceptible [14]. Furthermore, methanolic extracts from resistant-aged peels had inhibitory effects on pathogen growth and cucumber peels of resistant-aged fruit vs. susceptible age are enriched for genes associated with defense against biotic and abiotic stresses [14]. ARR to *P. capsici* also has been observed in whole pepper plants [4], chile pepper fruit [23], and several cucurbit fruits [6, 10]. Similar to cucumber, wounding overcame the resistance in chili pepper fruit and squash. The ARR in pepper and squash was suggested to result from structural changes in fruit surface properties such as cuticle thickness or cell wall lignification [10, 23].

Collectively, studies of ARR to date suggest that it may arise by means of preformed and/or induced defenses. In either case, a developmentally regulated difference in expression of the defense components is required. A comparison of uninoculated peel transcriptomes of ARR expressing and non-expressing cucumber fruit revealed the potential for either or both cases [15]. Of the 80 genes that were uniquely upregulated in ARR expressing fruit at the resistant age, four putative resistance genes (R-genes) as well as resistance related transcription factors were identified. Furthermore, this set of genes was highly enriched for specialized metabolism genes, including terpenoid synthesis and decoration genes, and untargeted metabolomic analyses of the same tissues revealed an increased accumulation of glycosylated terpenoids in the resistant tissue [15]. The accumulation of these preformed compounds may work in inhibiting infection, while at the same time developmentally regulated expression of *R*-genes may provide the ontogenic capacity to sense and respond to infection.

A highly resolved transcriptomic characterization of early of infection in ARR response could shed light on the mechanism by which this resistance is controlled, revealing if preformed or induced defenses are recruited. Furthermore, utilizing an ARR pathosystem allows a unique opportunity to examine both compatible and incompatible interactions within the same plant genotype. To our knowledge only one other transcriptomic study of ARR was performed, in that case apple leaves of different ages inoculated with *Venturia inaequalis* were sampled at 72 and 96 hpi [24]. In this study, our goal was to determine whether a developmentally regulated inducible mechanism contributes to ARR of cucumber fruit to *P. capsici*. To address this question, we performed microscopic, quantitative bioassay, and transcriptomic time courses characterizing the response of cucumber peel to inoculation with *P. capsici* at resistant (16 dpp) and susceptible (8 dpp) ages during the first 48 hpi. Differential expression analysis combined with weighted gene-co-expression network analyses and cubic spline regression analysis identified modules and defense-related genes which were uniquely induced in resistant fruit at early timepoints post-inoculation. The findings suggest that a developmentally acquired ability to initiate early defense responses may be crucial in conferring ARR to *P. capsici* in cucumber.

## Results

### The cultivar ‘Poinsett 76’ displays age-related resistance to P. capsici

Our previous ARR studies [6, 13–15] examined ‘Vlaspik’, an F_1_ hybrid commonly grown for processing cucumber production in the midwestern USA. To facilitate future analyses, we sought to identify a homozygous inbred cultivar expressing ARR. Testing of 22 inbred cucumber cultivars was performed using a detached fruit assay [13]. Hand-pollinated fruit were harvested at 8 and 16 dpp. The unwounded fruit surface was inoculated with droplets of *P. capsici* zoospore suspension and evaluated over a period of ten days using a 9-point disease score rating (Supplementary Table 1). Two of the inbred cultivars were identified to exhibit ARR, including the fresh market cultivar ‘Poinsett 76’ (Figure 1). As described for ‘Vlaspik’, very young ‘Poinsett 76’ fruit are extremely susceptible to infection, showing severe symptoms and extensive mycelial growth and sporulation (Figure 1 A). Fruits then become increasingly resistant as they complete their exponential growth phase. As fruits reached full size, at ∼16 dpp, they primarily exhibited localized necrosis at sites of inoculation, with occasional successful infection at inoculation sites (Figure 1 B).

**Table 1.**
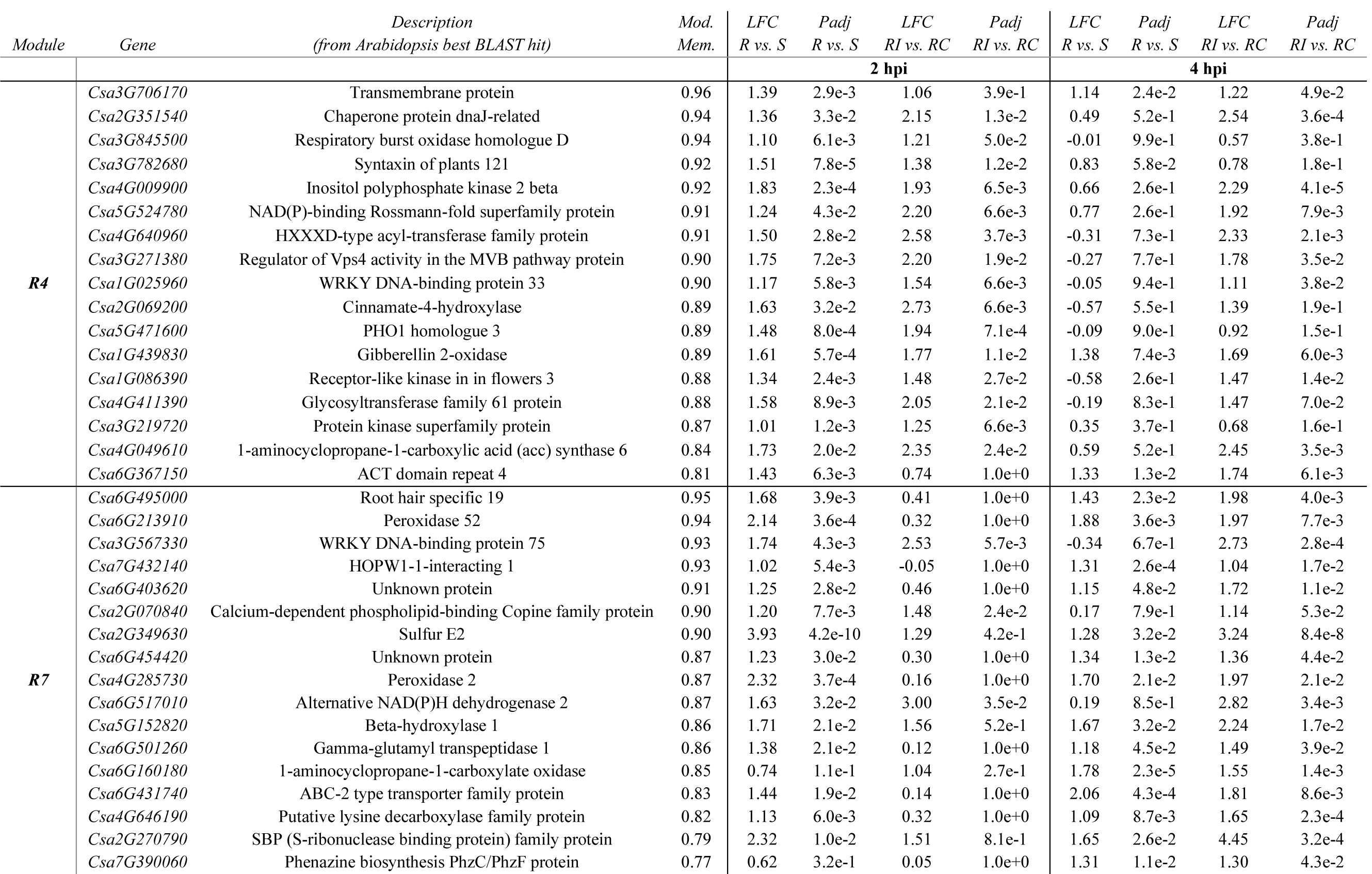
Genes with high module membership to Modules R4 and R7 that are differentially expressed in resistant-aged fruit compared to the uninoculated control and inoculated susceptible fruit. Mod. Mem. – Module membership; LFC – log_2_(Fold Change); Padj – Benjamini-Hochberg adjusted p-value; R – resistant 16 dpp; S – susceptible 8 dpp; RI – Resistant inoculated; RC – Resistant control.

**Figure 1.**
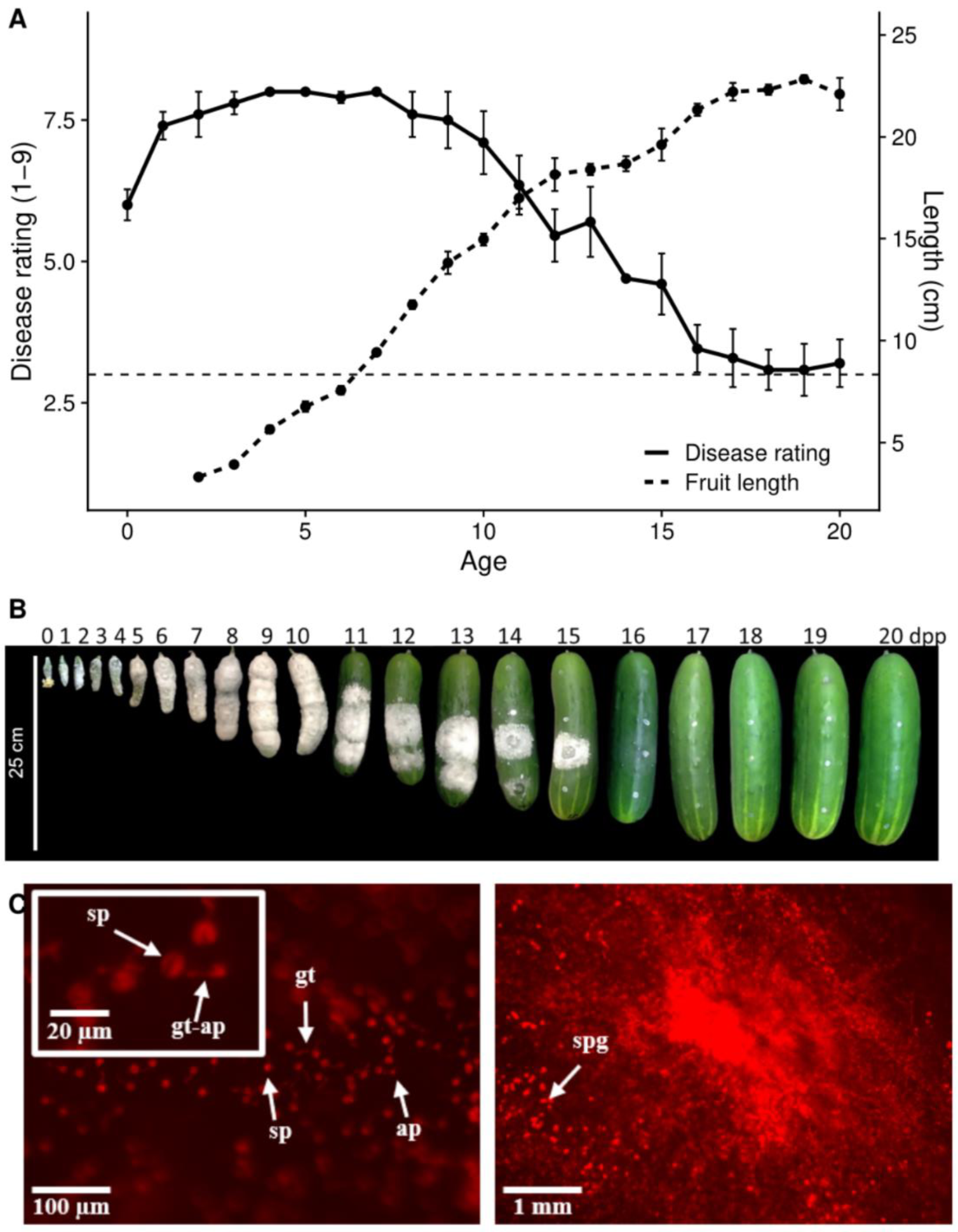
Cucumber cultivar ‘Poinsett 76’ exhibits age-related resistance to *P. capsici*. (**A**) Fruit length and disease rating (DR) as a function of fruit age. Fruit were ranked on a 1-9 DR scale (1=no symptom; 9=extensive mycelial growth and sporulation) at 5 days post-inoculation. The dotted line at DR = 3 represents localized necrosis. Points are means of 5-6 fruit. Error bars represent +/- standard error of the mean. (**B**) Representative fruit and disease progression at 5 dpi. (**C**) Fluorescently labeled *P. capsici* on <8 dpp cucumber fruit at 4 (left) and 72 (right) hpi. sp – spore; gt – germ tube; ap – appressoria; spg – sporangia.

### Age-dependent differential transcriptomic responses to infection

As a first step to explore the early stages of infection in susceptible age fruit (8 dpp) we observed germination and growth of a constitutively fluorescent isolate of *P. capsici* NY-0644-1 [25]. Peel sections from? Young fruit were inoculated with droplets of zoospore suspension, prepared for microscopy, and observed for 72 hours. Consistent with observations of *P. capsici* development on tomato leaves [19], appressoria formation was observed by 4 hpi, extensive growth by 24 hpi, and sporangia formation by 72 hpi (Figure 1 C).

Based on these results we compared transcriptomic responses of resistant (16 dpp) and susceptible (8 dpp) fruit peels at 0 (uninoculated), 4, 24, and 48 hpi. For each age and timepoint three fruit were inoculated with 6-12 droplets. All inoculation sites for a given fruit were harvested and pooled for sequencing providing ∼20M reads per sample. An average of ∼82% reads uniquely quasi-mapped to the cucumber transcriptome (Supplementary Figure 1). Pearson’s correlations of replicate samples were at least 96% (Supplementary Figure 2) showing high reproducibility among replicates.

**Figure 2.**
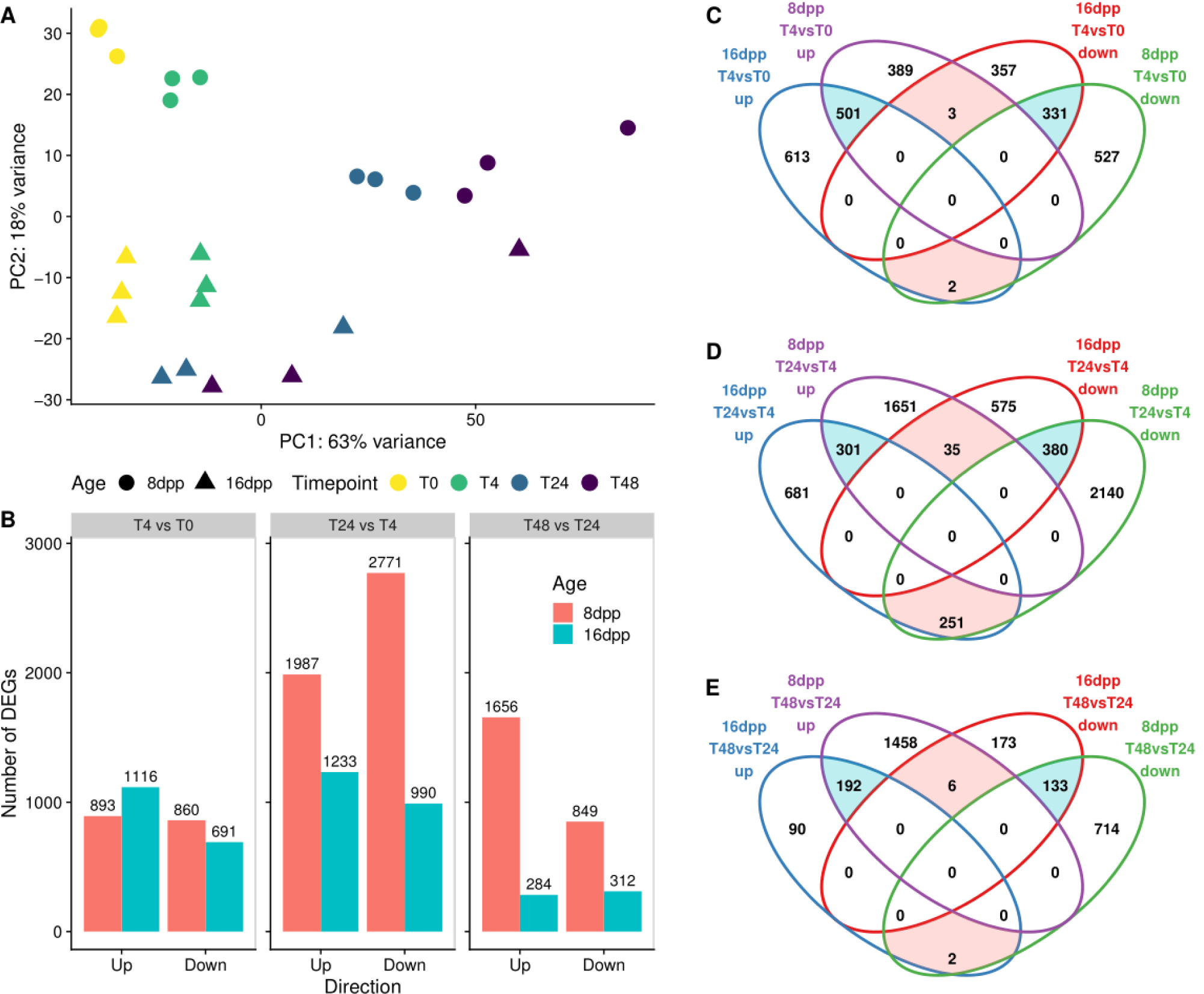
Age-dependent differential transcriptomic responses to infection. (**A**) Principal component analysis (PCA) of 8 and 16 dpp inoculated fruit at 0, 4, 24, and 48 hpi. (**B**) Number of differentially expressed genes in each of three within-age consecutive contrasts: 4 vs. 0 hpi, 24 vs 4 hpi, and 48 vs 24 hpi. (**C -E**) Venn diagrams comparing all significantly differentially expressed (adjusted P<0.05, fold change ≥ 2) genes in 8 and 16 dpp inoculated fruit at (**C**) 4 vs. 0 hpi, (**D**) 24 vs 4 hpi, and (**E**) 48 vs 24 hpi. Counts in red and blue denote up- and downregulated genes, respectively. Blue and red highlighted areas represent shared- and inverted-differential-expression, respectively.

To first identify any age-dependent differences between 8 and 16 dpp fruit, we performed transcriptional comparisons of the uninoculated control fruit peels. This analysis revealed that over 3400 and 4300 genes were up- and down-regulated genes with age, respectively. Gene Ontology (GO) term enrichment analyses (Supplementary Table 2) revealed that “specialized metabolic processes” (*p-value* = 7.50e-05) and “defense response to fungus” (*p-value* = 9.11e- 03) were among the upregulated terms. These results are consistent with our previous developmental analyses of other cucumber cultivars [14, 15] indicating potential differences in pre-formed factors contributing to resistance. Also consistent with prior studies and with the difference in developmental stages of the fruits, the most significantly enriched down-regulated term was “photosynthesis, light reaction” (*p-value* = 1.40e-26).

Principal component analysis (PCA) of the transcriptome data from inoculated and uninoculated samples showed that the first principal component largely reflected time post inoculation, while the second largely reflected fruit age (Figure 2). A similar transcriptional shift in direction and magnitude was observed along the positive direction of PC1 at 4 hpi regardless of age (8 dpp – circles, 16 dpp – triangles) of the tissue, suggesting a somewhat comparable initial response to infection (Figure 2 A). In contrast, subsequent timepoints (colors) exhibited differential transcriptional responses to infection as evidenced by the PCA. The susceptible 8 dpp samples progressively moved along the positive direction of PC1 with time, while resistant 16 dpp samples largely stayed in same position relative to PC1, suggesting little subsequent change in gene expression. One 16 dpp sample in each of 24 and 48 hpi timepoints exhibited transcription signatures that approached those of infected 8 dpp fruit at the same respective timepoints. As successful infection can occasionally occur on 16 dpp fruit, this was likely the case for those two samples. These samples had little effect on results (due to treatment of outlier genes in DESeq2), therefore analysis of differential gene expression was performed including the two samples.

### Resistant-age fruit mount a successful response by 24 hours post inoculation

To assess the changes in gene expression during infection we analyzed the transcriptional changes that occurred between consecutive timepoints at each fruit age (4 vs. 0 hpi, 24 vs. 4 hpi, and 48 vs. 24 hpi). Differential expression analysis showed that approximately 1800 genes were differentially expressed (up or down) at 4 hpi compared to uninoculated tissue, for both the 8dpp and 16dpp fruit, indicating a rapid and strong response to infection at both ages (Figure 2 B). However, as suggested by the PCA, progressive changes in gene expression were markedly different in the susceptible and resistant tissues at subsequent contrasts. While there was an increase in differentially expressed genes (DEG) in the susceptible 8 dpp fruit peels with time, (4758 and 2505 DEG at 24 and 48 hpi, respectively) the resistant 16 dpp samples had a smaller number of DEG at 24 hpi vs 4 hpi (2223) and by 48 hpi only about 500 genes were differentially expressed compared to 24 hpi. This approach allowed us to identify key points of transcriptional change during early infection.

To further understand the biological processes involved in the two responses, the most significantly enriched GO-terms (Fisher-*weight01* p-value < 0.01) were compared for each of the consecutive contrasts (Figure 3). At 4 hpi, upregulated genes in inoculated 8 and 16 dpp fruit were strongly enriched for defense related genes. “Response to wounding”, “defense response”,

**Figure 3.**
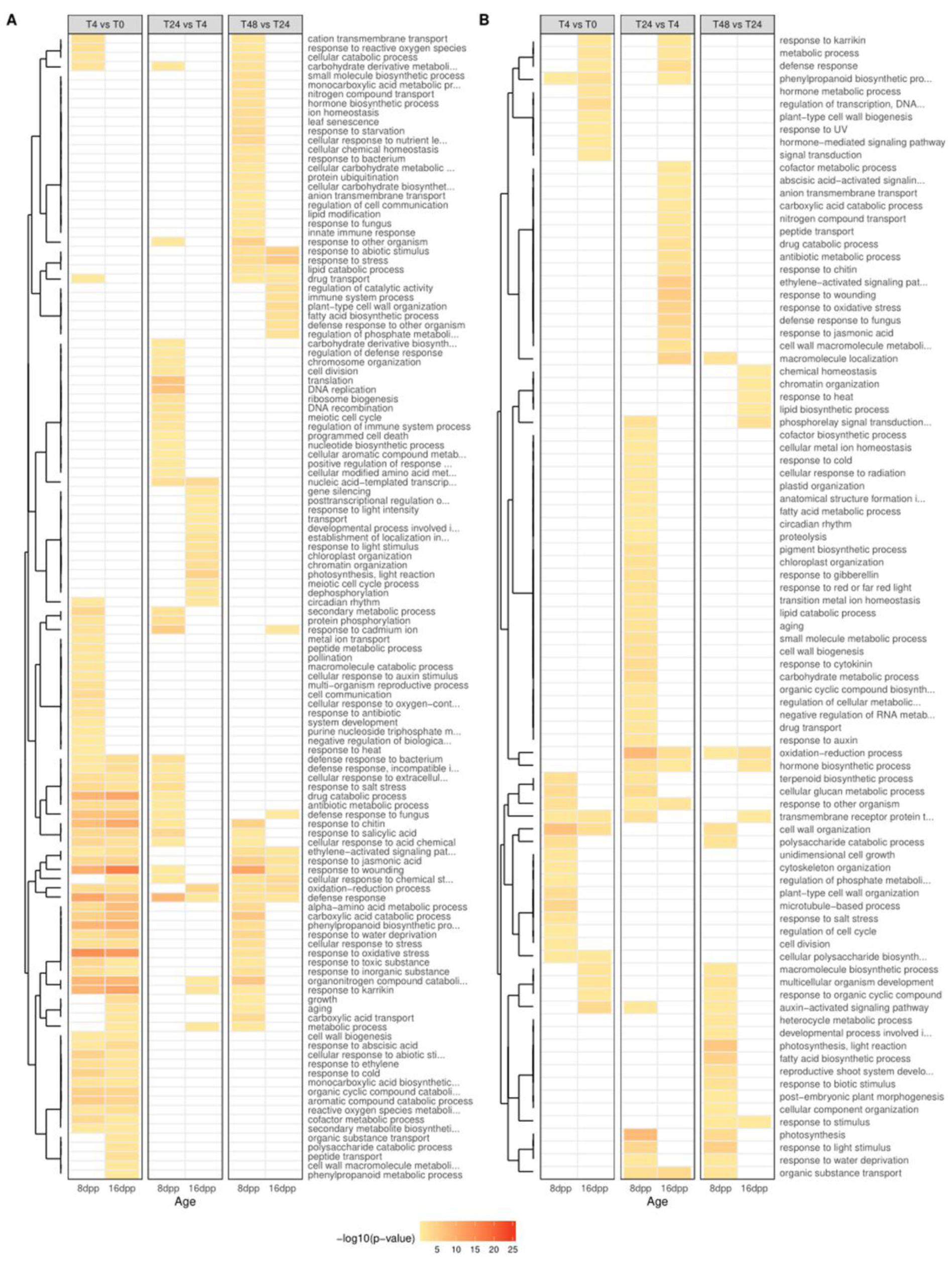
Resistant-aged fruit show transient upregulation of defense related genes. Gene Ontology enrichment of up- and down-regulated genes (**A** and **B**, respectively). Each row represents an enriched term at one of the three consecutive contrasts: 4 vs. 0 hpi, 24 vs 4 hpi, and 48 vs 24 hpi. Enrichment p-value threshold of 0.01. Terms clustered by Euclidean distances.

“response to chitin”, “phenypropanoid biosynthetic process, “response to oxidative stress”, “response to karrikin”, “organonitrogen compound catabolic process”, “defense response to fungus”, “aromatic compound catabolic process”, and “drug catabolic process” were the ten most enriched terms in both ages at this time point. Although the number and GO categories of genes differentially expressed at 4 hpi was comparable between the two ages, fewer than half of the DEG in the resistant samples was shared with those differentially expressed in the susceptible samples (Figure 2 C; blue shading). Analysis of the 613 genes uniquely upregulated in the resistant 16 dpp samples at 4 hpi revealed a potentially unique set of defense related genes involved in an early incompatible interaction (Supplementary File 2).

When comparing 24 hpi to 4 hpi, and 48 hpi to 24 hpi, less than 15% of the thousands of up- and downregulated genes were shared between the ages, respectively (Figure 2 D and E). Susceptible 8 dpp fruit continued to upregulate defense (top five enriched terms: “defense response”, “DNA replication”, “translation”, “response to cadmium ion”, and “response to salicylic acid”) while down-regulating photosynthetic processes and other homeostatic processes, such as carbohydrate metabolic processes. In contrast, by 24 hpi, resistant 16 dpp fruit were upregulating photosynthetic and growth-related genes and downregulating defense (top five downregulated terms: “response to wounding” “ethylene-activated signaling pathway”, “macromolecule localization”, “response to oxidative stress”, “defense response to fungus”) suggesting a return to normal state (Figure 3 B). This is especially evidenced by the large number of inversely regulated genes in the 8 dpp vs. 16 dpp samples at 24 hpi indicating an opposite response (red shading). The set of 251 genes upregulated in 16 dpp and downregulated in 8 dpp was strongly enriched for photosynthesis (*p-value* = 5.2e-13). Collectively these observations suggest that the resistant fruit have successfully mounted a defense by 24 hpi, indicating the importance of further investigation prior to that timepoint.

### Analysis of pathogen growth provides evidence for infection failure in the first 24 hours on resistant fruit

The transcriptomic suggestion of a rapid and potentially successful defense response within 24 hours prompted us to more closely investigate pathogen growth during the first 24 hours of infection using electron microscopy and a high throughput microplate assay. Samples were collected for Scanning Electron Microscopy (SEM) from 8 dpp and 16 dpp aged cucumber fruit inoculated with droplets of zoospore suspension (5 × 10^5^ spores/ml) at time intervals of 0, 2, 4, 8, 12, 18, and 24 hpi. At each timepoint, three samples were collected, each from a distinct inoculated fruit. Morphological differences between 8 dpp and 16 dpp fruit were readily observed; the younger susceptible fruits had smaller more densely packed cells and trichomes, as well as warts that produced valley regions that in some cases increased spore density due to the surface topography (Figure 4).

**Figure 4.**
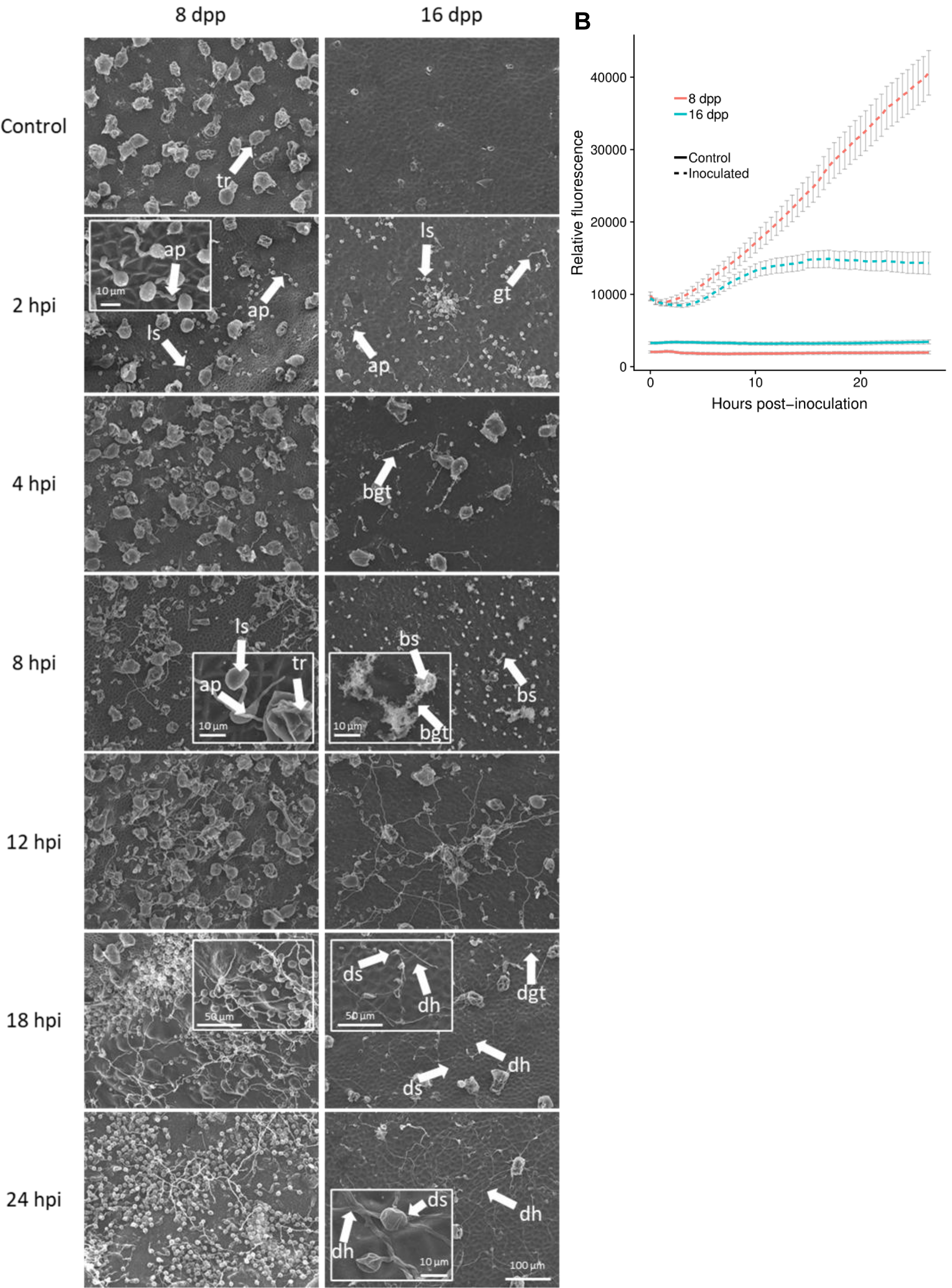
Evidence for infection failure in the first 24 hours post inoculation on resistant fruit. (**A**) Scanning electron micrographs of inoculated 8 and 16 dpp fruit. tr – trichrome; ls – live spore; ap – appressoria; gt – germ tube; bgt – burst germ tube; bs – burst spore; deflated spores (ds), deflated germ tubes (dgt), and deflated hyphae (dh). Bottom right scale bar for all frames, except for insets. (**B**) High-throughput *in vivo* analysis of pathogen growth on fruit plugs. For inoculated and control treatments, each point is a mean of 36 and 12 replicates, respectively. Error bars are +/- SEM.

At 2 hpi, encysted *P. capsici* spores were observed germinating on fruit of both ages. Formation of some appressoria was also observed as early as 2 hpi. By 4 and 8 hpi, some differences were observable between the resistant (16 dpp) and susceptible (8 dpp) fruit. While spores on susceptible fruit continued to germinate and form appressoria, in four of the six 16 dpp samples, lysed spores and germ tubes were observed, suggesting either preformed antimicrobial compounds or a rapidly induced defense response may inhibit successful infection as early as 4 hpi. As infection progressed, more evidence of failure to infect was observable in the majority of the resistant samples. By 18 and 24 hpi, deflated spores, germ tubes, and hyphae were observed on most of the resistant fruit samples, suggesting that spores that survived an initial defense response may be stopped at a later time, during the first 24 hours. No such histological signs of deflated or burst pathogen structures were observed at any timepoint in samples from susceptible 8 dpp fruit.

Fluorescence based *in vivo* infection assays provided further, quantitative evidence of inhibited infection in resistant aged fruit by 24 hpi. In these assays, fruit peel plugs (36 replicate fruit sections per fruit age, per experiment) were placed in a 96-well plate, inoculated with fluorescently labeled zoospores; fluorescence signal was measured hourly for 24 hours. After a short lag phase, the signal from the labeled *P. capsici* on inoculated susceptible 8 dpp samples grew linearly throughout the 24-hour period (Figure 4 B). However, on 16 dpp resistant-aged fruit samples, intensity of the fluorescent signal plateaued by 8 – 10 hpi, further suggesting early inhibition of pathogen growth. Together, the SEM and bioassay results bolster the transcriptional evidence suggesting that infection is thwarted by 24 hpi in 16 dpp cucumber fruit.

### Transcriptomic investigation of the first 24 hours post inoculation

Concurrent with the collection of samples for SEM (0, 2, 4, 8, 12, 18 and 24 hpi), inoculated and uninoculated tissue was harvested for a second transcriptome analysis using 3′ mRNA sequencing. In total, 78 3’-mRNA libraries were sequenced to an average depth of ∼5M reads/sample and an average of ∼53% reads quasi-mapping to the 3’-extended cucumber transcript sequences (Supplementary Figure 3). Two samples (8dpp_T12_Inoc_1 and 8dpp_T18_Cont_2) had aberrantly low read coverage (<0.5 M reads) and were excluded from analysis. A PCA comparison of timepoints shared between the two transcriptome experiments (0, 4 and 24 hpi) showed tight clustering of samples within their respective timepoints, indicating high reproducibility between the two experiments, performed in different years, and using different sequencing library technologies (Supplementary Figure 4).

The PCA of data from experiment 2 revealed modest changes in transcriptomic patterns for uninoculated samples of both ages (open symbols) relative to timepoint 0 (asterisks), likely reflecting a combination of diurnal changes and the effects of fruit detachment from the vine (Figure 5). In contrast, the inoculated samples (closed symbols) showed strong transcriptional changes, especially for the susceptible 8 dpp fruit (circles) (Figure 5 A). From 4 hpi and beyond, the inoculated 8 dpp samples exhibited a sequential transcriptomic transition. Conversely, samples collected from the resistant-aged 16 dpp fruit (triangles) all clustered together, and relatively closely to uninoculated fruit, from 4 hpi and beyond. Notably, at 2 hpi, samples from uninoculated 8 dpp fruit clustered with uninoculated control, while 2 hpi samples from 16 dpp fruit showed a clear difference from the uninoculated controls, suggesting an earlier response to infection in the resistant-aged fruit.

**Figure 5.**
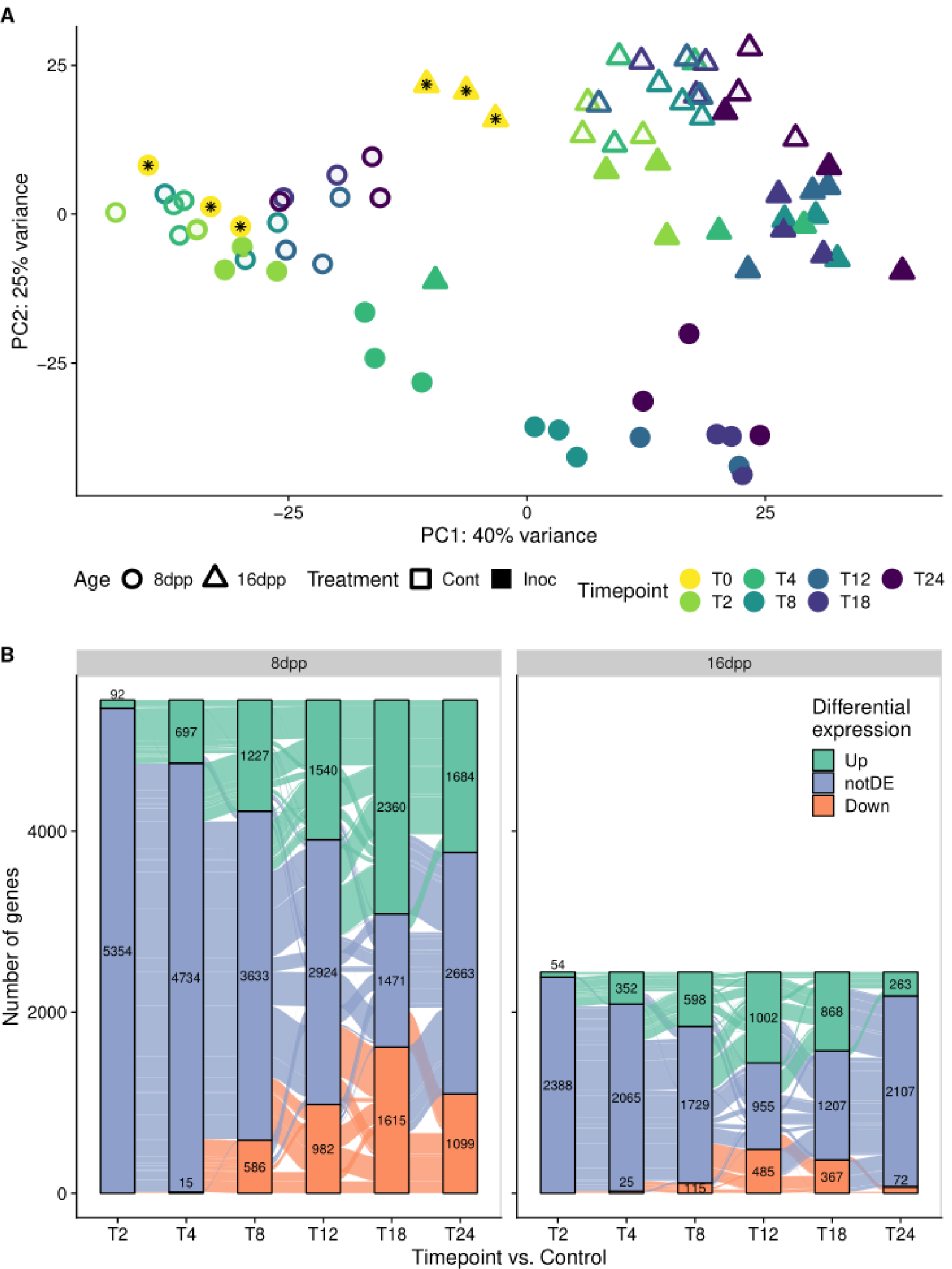
Differential transcriptional dynamics within the first 24 hours. (**A**) Principal component analysis of inoculated and control fruit of two ages, 8 and 16 dpp. Samples collected at time 0, which represent both Inoculated and Control treatments, are denoted by an asterisk. (**B**) Alluvial plots showing all genes with significantly changed expression at each timepoint compared to respective controls. Each stacked bar represents the numbers of genes either up-down- or not-differentially (notDE) regulated at sequential time points. Genes are grouped based on expression patterns throughout time.

To further understand the trends in gene expression changes in response to infection, all genes that were differentially expressed in at least one timepoint vs. the respective non-inoculated control were displayed as stacked bars (strata) in an alluvial plot (Figure 5 B). The alluvia (curves) illustrate progressive changes in gene expression over time post-inoculation (2h to 24h); each alluvium represents a group of genes that shares expression pattern over time. The large alluvia from the nonDE strata (blue) show a phased response to infection, in which certain genes are involved in early timepoints, while others in later stages of infection.

The figure reveals that response to infection is dramatically different in the two ages. Most evident is that more than double the number of unique genes is differentially expressed at least once in susceptible fruit compared to resistant fruit. Examination of the trends in 8 dpp susceptible fruit revealed a pattern of sequential accumulation of DEG, as previously suggested by the PCA. The number of DEG grew continuously until 18 hpi, and most genes which were differentially expressed at one timepoint continued to be differentially expressed at following timepoints. Of the 697 upregulated genes at 4 hpi, 424 are continuously upregulated in every subsequent timepoint. By 24 hpi, 1523 of the 1684 upregulated genes had been previously upregulated at one of the timepoints and then subsequently upregulated at all following timepoints. A similar trend is observable in downregulated genes.

Conversely, by following alluvia of genes initially upregulated at 4 hpi in the 16 dpp resistant-aged samples, it is evident that most are subsequently not differentially expressed at further timepoints. Of the 352 genes upregulated at 4 hpi, 140 genes are not differentially expressed at 8 hpi, and only 45 are continuously upregulated through 24 hpi. Though the number of DEG grows until 12 hpi, most genes are timepoint specific, being differentially expressed once or at most at two timepoints. Most striking is the fact that at 24 hpi, only 263 and 72 genes are up- and down-regulated, respectively, compared to uninoculated tissue, further confirming the culmination of the defense response in resistant-aged fruit.

### Gene co-expression is preserved but not expression patterns over time

To better identify transcriptional co-expression patterns in the data we employed Weighted Gene Co-expression Network Analysis (WGCNA) [26]. This analysis identifies genes with highly correlated transcriptional patterns and groups them into co-expression clusters or modules. These modules help understand biological processes and identify key genes associated with these processes [26]. Two independent response networks for the susceptible and resistant fruit (Figure 6) were assembled. After module merging (based on eigengene correlation), a total of 17 and 29 modules were defined for the susceptible and resistant inoculated networks, respectively (Supplementary Files 3 and 4).

**Figure 6.**
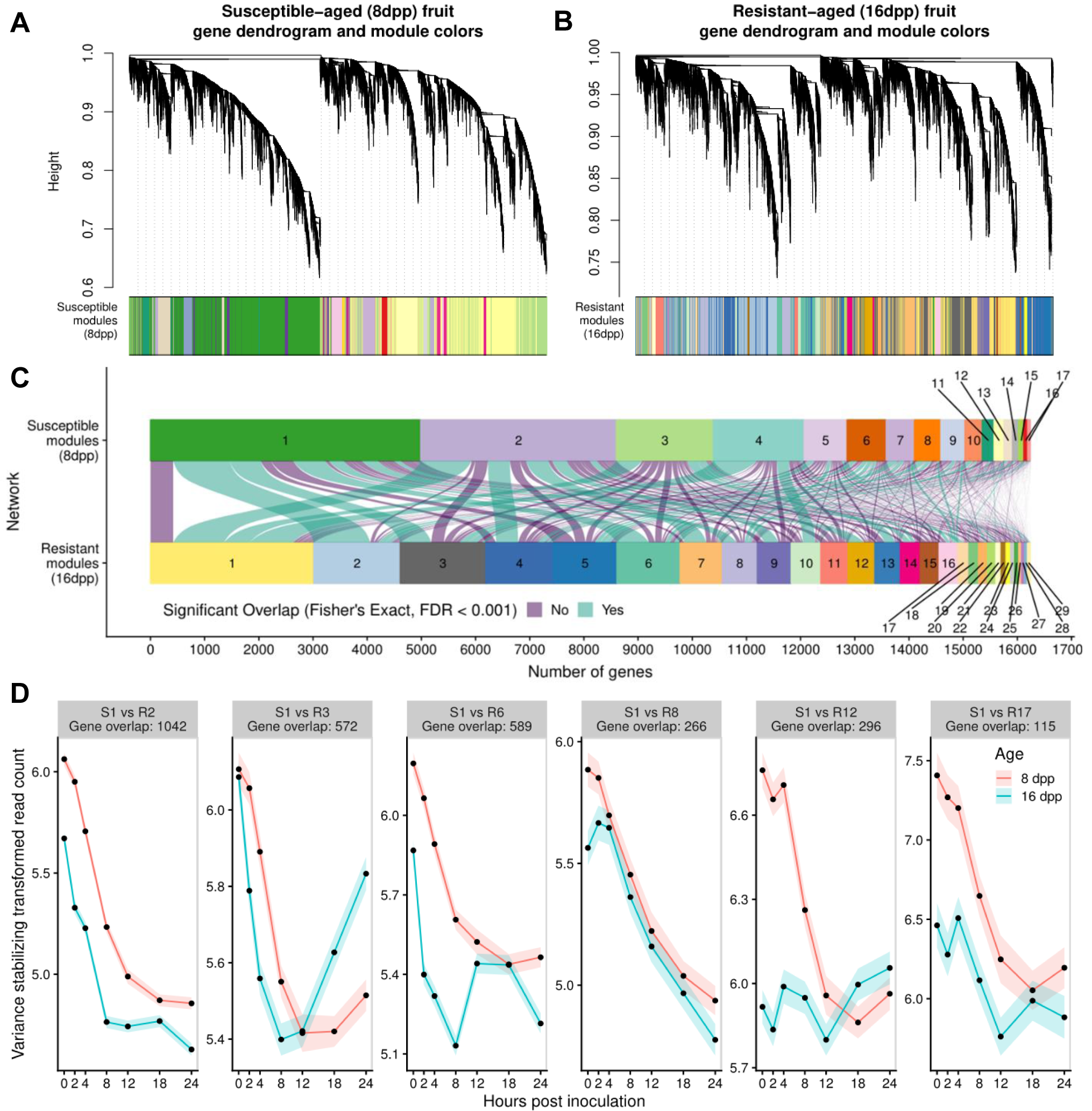
Defense response network reprogramming in the two fruit ages. Weighted gene co-expression analysis (WGCNA) was used to analyze all expressed genes in inoculated samples. Separate signed networks were analyzed for susceptible (**A**) and resistant (**B**) fruit. Dendrograms cluster the genes based on their topological dissimilarity. The colored bars underneath the dendrograms show co-expression module assignment. (**C**) Module overlap between the susceptible and resistant networks. Each curve connecting two modules in the different network represents a shared group of genes. Curve width is proportionate to the number of genes. Curve color shows if the overlap between modules is significant based on Fisher’s exact test and an FDR of < 0.001. (**D**) Variance stabilized expression patterns of genes in Susceptible module 1 (S1) that are also co-expressed in resistant modules (R2, R3, R6, R8, R12 and R17; Module overlap curves marked with an asterisk in panel **C**).

Module preservation statistics [27] showed that gene co-expression was preserved in many modules when comparing the susceptible and resistant networks (Supplementary Figure 5). Module gene overlap (Figure 6 C) also showed significant preservation (Fisher’s Exact test and FDR < 0.001) of gene co-expression between many modules of the two networks. For example, close to half of the expressed genes in the susceptible response are clustered in the Susceptible Module 1 (S1) which is consistently downregulated until 18 hpi (Figure 6 D). The genes in this module overlap with subsets of genes in six resistance modules (R2, R3, R6, R8, R12 and R17), indicating they are also co-expressed in the resistant network, but in a different manner. For instance, a subset of the co-expressed genes present in module S1 are co-expressed in resistant-age fruit in module R12; however, while the genes in S1 show a drop in expression, in resistant fruit they have minimal fluctuations in expression over time. Thus, while many groups of genes were expressed in concert over time, indicating coordinate regulation regardless of fruit age, the diverse module assignment and thus age-specific patterns of expression, suggests a reprogramming of the response network in resistant-aged fruit.

### Biological processes identified by weighted gene co-expression network analysis

Using gene module assignment defined by the resistant network, expression patterns of the genes within a given module were compared in control and inoculated fruit of both ages (Figure 7). Genes identified in many of the co-expression modules of infected fruit had distinct patterns from those in uninoculated control fruit, suggesting we were able to isolate specific defense responses in response to inoculation.

**Figure 7.**
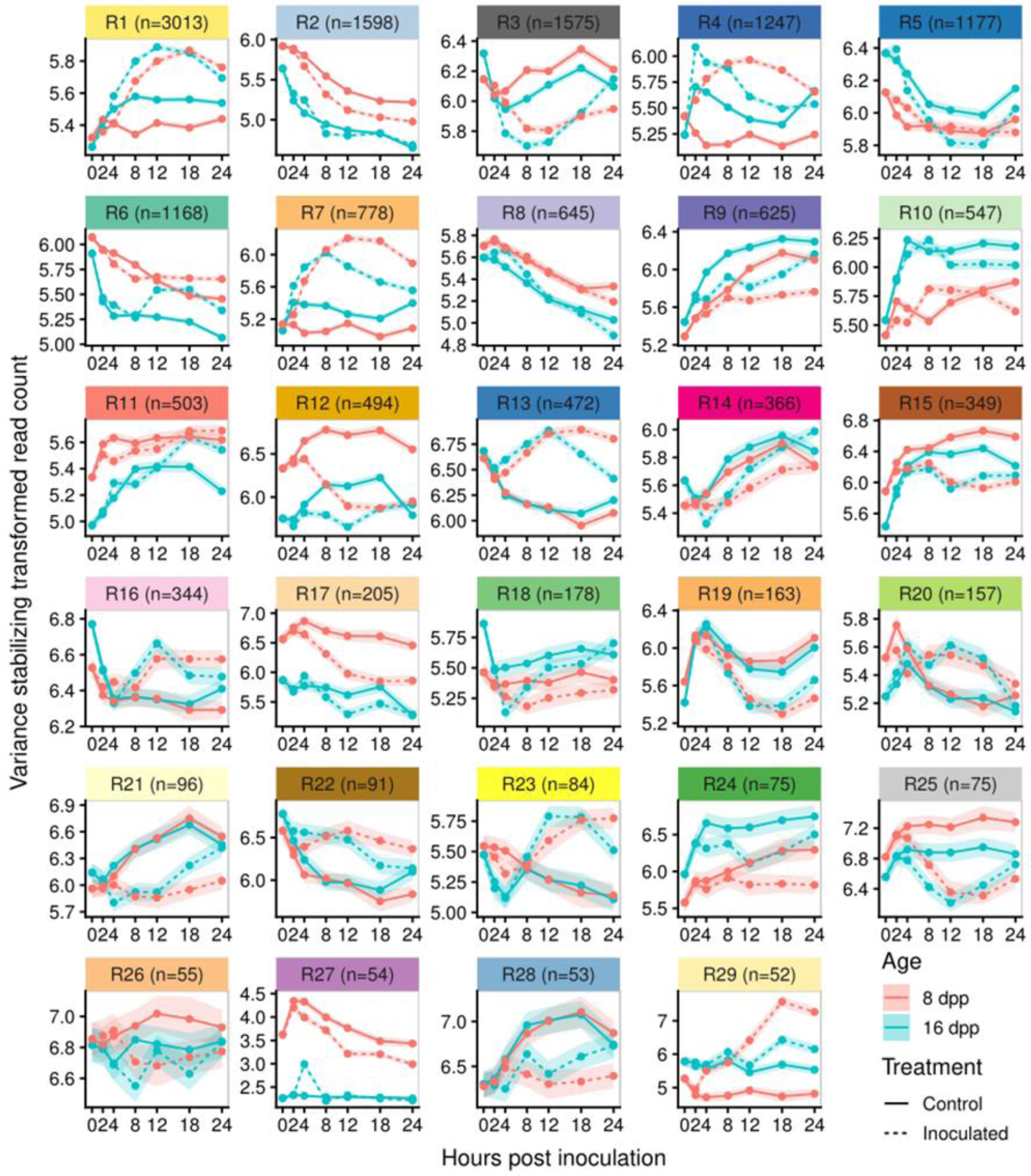
Resistant network module gene expression patterns. Each panel represents a module (number of genes indicated in label) from the infection co-expression network of resistant fruit (Figure 3.7A). Comparison of expression patterns of the module-defined genes in the different ages and treatments across time. Data are mean and SEM of variance stabilized normalized read counts (which is useful for pattern comparisons) for all genes in the module.

GO-term enrichment analysis of the different modules demonstrated the biological relevance and function of identified modules (Supplementary File 5). For example, Module R1 (Resistance Module 1) (n genes = 3013) showed patterns of increased expression in response to inoculation in both ages. Uninoculated tissue largely showed unchanging expression levels throughout the entire time course. GO term enrichment showed that this module was strongly associated with translation and ribosome biogenesis, suggesting induction of protein synthesis in response to infection.

In Modules R2 and R9 (n genes = 1598 and 625, respectively) all genes, regardless of age or treatment, showed a similar pattern of change over time. Genes in Module R2, which exhibited gradual decline, were enriched for carbohydrate metabolic process, lipid metabolic process, cell division, signal transduction and response to abiotic stimulus. Higher baseline expression of 8 dpp fruit is likely due to the different stage of development, as unharvested fruit at this age are still rapidly growing (Figure 1 B). Genes in Module R9 showed gradual increase and were enriched for “organonitrogen compound catabolic process”, “aromatic compound catabolic process” as well as” response to water deprivation”. The expression patterns of genes in these modules is probably a result of the fruit being detached during the analysis, i.e. deprived of carbohydrate source and subject to water loss.

Module R3 (n genes = 1575) showed a potentially circadian controlled expression pattern, with peak expression at 18 hpi (collected at 6:30 AM; sunrise at 6:15 AM) and almost identical patterns of expression in uninoculated fruit of both ages. In inoculated fruit of both ages, the circadian patterns of these genes were diminished, and they showed a prompt drop in expression levels after inoculation. By 24 hpi however, expression levels in resistant fruit returned to levels of uninoculated fruit, while those in susceptible fruit remained low. This module was enriched for processes involved in transcriptional regulation as well as response to light stimulus.

### Modules induced in early infection of resistant-aged fruit

Several modules indicated differential response to infection between the resistant and susceptible ages. Based on the PCA and microscopy, we were especially interested in modules that showed very early response differences. To identify such modules, expression patterns of genes, in the resistant network modules, were compared between 8 dpp and 16 dpp samples using regression analysis. Because gene expression patterns do not follow linear trends over time, a cubic spline basis function was applied to the time variable with three internal knots, splitting the time course into quartiles (0-3, 3-8, 8-15, and 15-24 hpi). Most of the modules (24/29) showed significant interaction effects between age and splined modeled time in at least one quartile, indicating that genes in each module behave differently across time in the two ages, further reflecting the reprogramming of module expression patterns (Supplementary Table 3). Of specific interest were early induced modules in 16 dpp peels vs. 8 dpp peels. Nine modules had significant interaction effects during the first two spline fractions, 0-3 hpi and 3-8 hpi (Figure 8).

**Figure 8.**
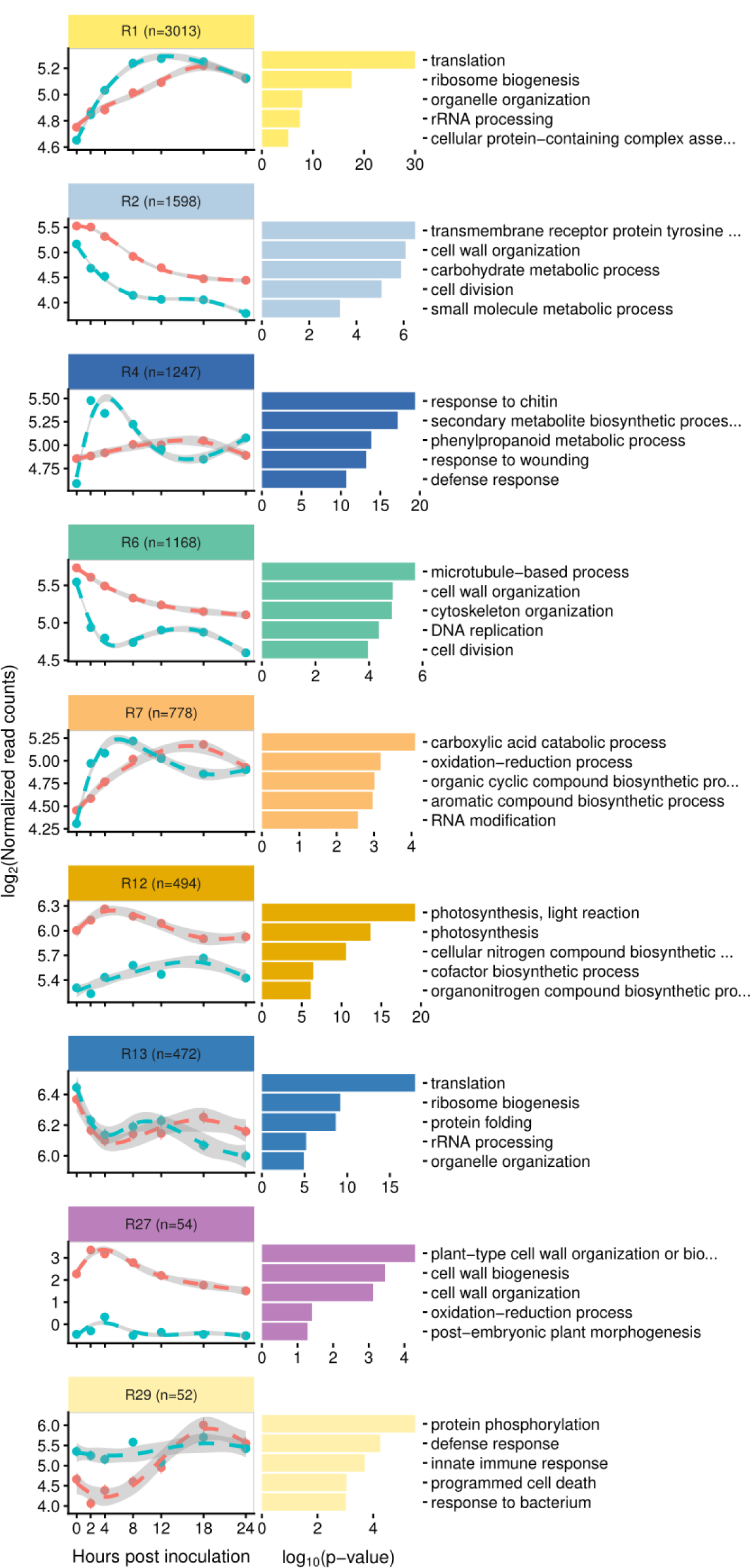
early response modules in the resistant network. Points represent the mean log_2_(normalized read count) of inoculated 8 dpp or 16 dpp fruit. Lines and grey ribbons are predicted values (and standard errors) based on regression of log_2_(normalized read count) by a natural cubic spline of time with three internal knots (0-3, 3-8, 8-15, and 15-24 hpi). Modules selected are those with a significant interaction effect in one or both first spline fractions (0-3 and 3-8 hpi), i.e. have age-dependent response to infection in early timepoints. The five most significantly enriched gene ontology terms are indicated next to each module.

Module R4 (n = 1247), is defined by a spike of increased expression in response to inoculation of resistant-aged fruit. Expression levels peak at 2 and 4 hpi followed by decrease in expression starting at 8 hpi. In the inoculated susceptible fruit, the genes identified in this module showed a minimal change in expression prior to 4 hpi. GO term enrichment showed that this module was strongly associated with defense and pathogen recognition, the five most enriched terms being “response to chitin”, “secondary metabolite biosynthetic process”, “phenylpropanoid metabolic process”, “response to wounding”, “defense response”.

Similarly, Module R7 (n = 778) also showed statistically different expression during early infection, with increased expression by 2 and 4 hpi in inoculated resistant-aged fruit. In susceptible fruit these genes show a more gradual increase in expression, which only matches that of the resistant-aged fruit at 8 hpi. This module was also strongly associated with defense as evidenced by enrichment for terms associated with canonical response mechanisms “carboxylic acid catabolic process”, “oxidation-reduction process”, “organic cyclic compound biosynthetic process”, “aromatic compound biosynthetic process”.

Other interaction effects and expression patterns of note are observed in other modules. Module R1, which as mentioned earlier is strongly enriched for translation-related processes and is activated earlier and more rapidly in inoculated resistant-aged fruit. A more rapid reduction in expression in genes of Module R6 (enriched for microtubule, cell wall and cell division related processes) is observed in resistant-aged fruit, suggesting a rapid response. Genes in Module R29 which are associated with defense and innate immune responses, are already activated at 0 hpi and show little change in expression in resistant-aged fruit. Interaction effects in Modules R2, R12 and R27 which are enriched for growth, cell wall and photosynthesis related terms, are most likely associated with baseline differences between the different ages.

### Identification of early response genes in resistant-aged fruit

Genes with high module membership are strongly correlated to the module eigengene. We used this measure to identify genes highly connected to Modules R4 and R7, as these modules showed patterns of increased expression at early infection timepoints in resistant fruit. The second criterion for selection was those genes that also showed significantly increased expression in resistant inoculated fruit at 2 and/or 4 hpi compared to both the uninoculated control and the inoculated susceptible fruit. We identified 34 genes with a greater than 2-fold expression at 2 and/or 4 hpi in both comparisons as well as a module membership greater than 0.75 (Table 1, Supplementary Figure 6). As expected, the expression patterns of these genes strongly conform to those of the Module eigengene, with a uniquely strong expression at early timepoints in resistant inoculated fruit. Many of these genes are annotated to have canonical functions in early defense response, for example associated with phenylpropanoid-derived compound metabolism, reactive oxygen species (ROS) metabolism, gibberellin and ethylene balance, vesicle transport, protein phosphorylation, as well as pathogen perception and response (Figure 9). The expression patterns observed for these genes (i.e., earlier increase in 16 dpp samples, but higher level at 24 hpi in 8 dpp samples) were confirmed by comparison to their expression in RNA-seq experiment 1 (Supplemental Figure 7 and 8). These data showed strong reproducibility (Pearson’s correlation; R = 0.85, *p-value* < 2.2e-16) in response between experiments performed in different years and with different sequencing library techniques.

**Figure 9.**
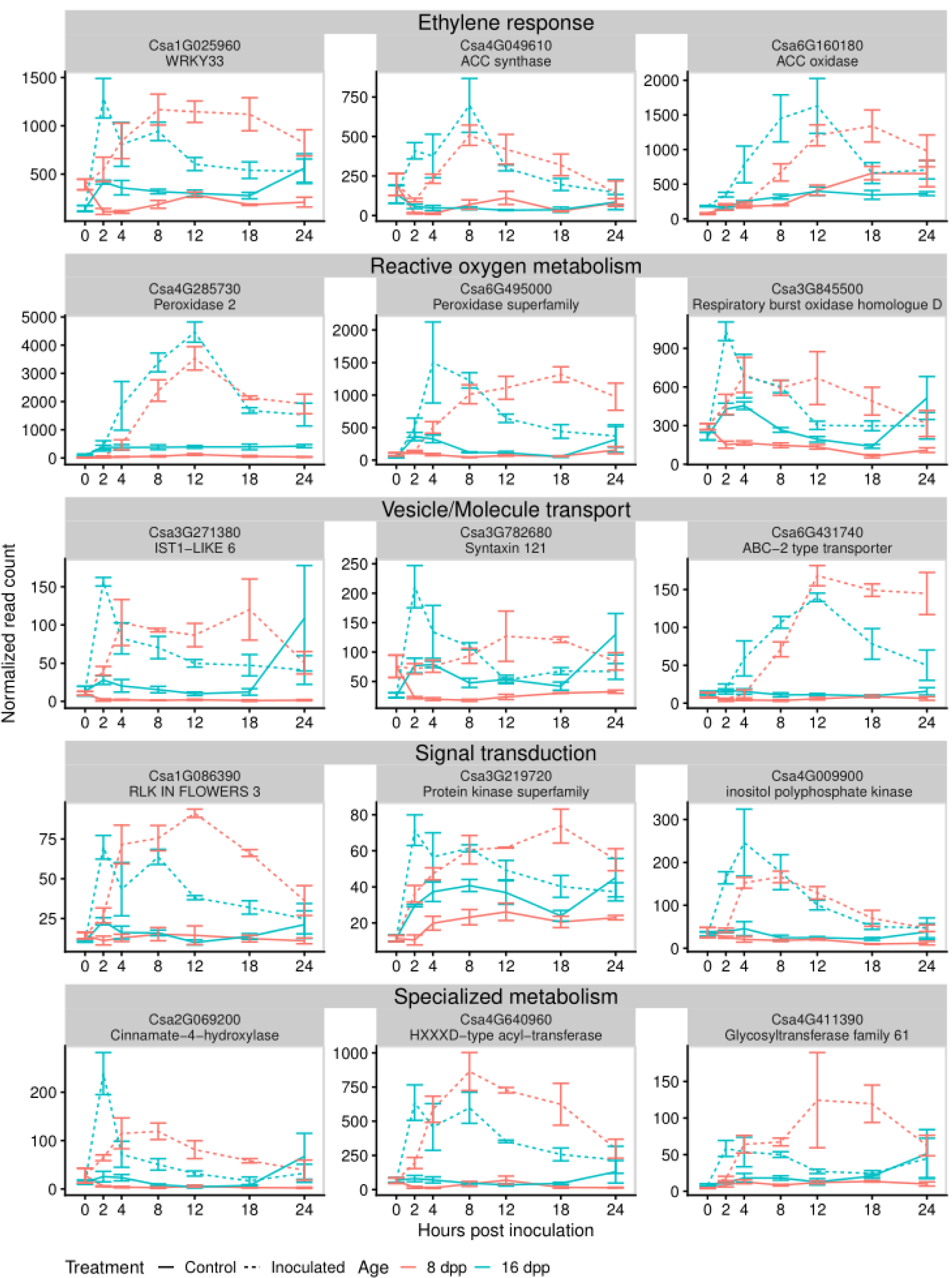
Genes induced in early responses to inoculation of resistant fruit. A subset of genes identified by a module membership > 0.75 and significant fold change of >2 in inoculated resistant fruit (16 dpp) compared to both control resistant fruit and inoculated susceptible fruit (8 dpp). Full list and expression patterns of identified genes available in Table 1 and Supplementary Figure 6. Mean expression for three biological replicates in each age and treatment. Error bars are +/- standard error of the mean.

## Discussion

### A rapid infection meets a rapid response

The infection process of *Phytophthora* spp. has been studied extensively on many plants and crops [28]. However, there are clear differences between the rates and severity of infection in different species and hosts. For example, *P. infestans* infections on potato and tomato are largely symptomless until 3 days post infection (dpi) [29]. As such, many transcriptome studies examine much later time points post inoculation [e.g., 29, 30]. In contrast, *P. capsici* has a rapid infection cycle with visible symptoms, often within 24 hpi, and can reach asexual sporulation by 2-3 dpi on a variety of species [6, 18].

Using both fluorescent and SEM, we observed that *P. capsici* infection of susceptible young cucumber fruit is also extremely rapid. On susceptible-aged fruit, hyphal growth was observed as early as 8 hpi and progressed rapidly throughout the first 24 hours, indicating a successful biotrophic infection. Using *in vivo* high-throughput bioassay, we quantitatively observed a detectable linear growth rate on cucumber fruit peel samples as early as 4-6 hpi, again suggesting quick establishment and rapid progression during this time. As described on other hosts [19], our fluorescent microscopy showed that by 72 hpi asexual reproductive sporangia have been formed in cucumber. It is this rapid infection and reproductive cycle that allows *P. capsici* to be such a devastating pathogen in the field.

Our prior studies have shown that certain cucumber cultivars and plant introgression lines [13–15] exhibit ARR at approximately 16 dpp, coinciding with the end of exponential fruit growth. Varieties that do not develop ARR remain highly susceptible to *P. capsici* and show severe symptoms of infection [13–15]. Evidently, inhibition of zoospore germination is not the mechanism of ARR; regardless of fruit age, within two hours of inoculation most *P. capsici* zoospores had already encysted and germinated (Figure 4), and some had formed appressoria. Strikingly however, the SEM of inoculated resistant fruit revealed histological signs, such as burst or lysed spores and germ tubes, as early as 4 hpi, and consistently by 8 hpi. Further confirmation of rapid inhibition of infection on the 16 dpp fruit was obtained from our quantitative *in vivo* bioassay showing cessation of pathogen growth by 8-10 hpi.

By 24 hpi, all three resistant-aged samples examined with SEM either showed no pathogen present, suggesting failure to remain attached to the fruit surface or deflated/unviable spores and hyphae. None of these histological signs were present in susceptible samples, on which infection proceeded normally. Similarly, a study of Port-Orford-cedar plants resistant to *P. lateralis* showed a reduction in pathogen presence as well as deflated hyphae and spores at 24 hpi [32]. Though there is limited histopathological evidence of such a severe defense response, similar signs of *Phytophthora* spp. (including *capsici)* inoculum death are observed after exogenous application of phytochemicals such as garlic root exudates or terpenoid-containing essential oils from oregano and other plants [33–36]. Evidence in other fungal pathosystems of spore- and hyphal membrane disruption by preformed or induced defensin proteins also exists [e.g., 36, 37]. This study and our prior work showed an upregulation of specialized metabolism associated genes in 16 dpp vs. 8 dpp uninoculated fruit peels. Thus, the rapid histological signs of pathogen death might result from preformed antimicrobial compounds, as was similarly implicated from peel extract assays and transcriptomic analysis of non-inoculated developing peels of other ARR cultivars [14, 15], or from a rapidly induced defense response.

The findings indicating rapid inhibition of infection were further supported by transcriptional evidence. Gene expression changes over time were markedly different between resistant and susceptible aged fruit; double the number of genes were involved in the susceptible response compared to the resistant one. Moreover, susceptible fruit exhibited progressive waves of gene expression changes peaking at 18 hpi, many associated with defense response (Figures 3 and 6 A). The susceptible response may be compared to a high-resolution transcriptional time series of the compatible response of Arabidopsis to infection by the fungal pathogen *Botrytis cinerea* [39]. In *Botrytis* infection of Arabidopsis, most differential expression also occurred at ∼18-30 hpi. As these timepoints are post-pathogen penetration, the increased gene expression at these times may represent a response which is too late to inhibit infection [39]. The increased expression at comparable timepoints in our data might suggest a similarly failed defense response in susceptible-aged fruit. Additionally, as was seen in successful infection of tomato by *P. capsici* [19], the susceptible cucumber fruit also exhibited down-regulation of genes associated with primary metabolism processes and photosynthesis in by 24 hpi.

Conversely, in resistant-aged fruit, an induced rapid activation of defense related genes at 2 and 4 hpi was followed by active downregulation of defense related genes by 24 hpi. Furthermore, at 24 hpi, relatively few genes were differentially expressed in inoculated resistant fruit compared to the control (Figure 5 B), providing an additional indication that the pathogen defense response is largely completed. This downregulation of defense makes intuitive sense if the pathogen has been eradicated or is no longer infecting the host, as was observed in the microscopic and *in vivo* assays. Apart from downregulation of defense, resistant-aged fruit showed upregulation of photosynthesis and other metabolic processes, suggesting a return to a “normal” or uninfected state after pathogen defeat. Rapid responses to *P. capsici* have also been observed in incompatible reaction on Arabidopsis leaves where failure to penetrate, ROS bursts, callose deposition, and hypersensitive cell death all occurred within 24 hpi [40]. Together the evidence from microscopy, bioassay and transcriptome studies suggested that factors, either preformed or induced, prior to 24 hpi are important in conferring ARR.

### A reprogramming of gene co-expression networks of infection at the resistant age

To better understand the effect of fruit age on gene expression patterns in response to *P. capsici* inoculation, the high-resolution transcriptomic time course was further analyzed using WGCNA [26]. This advanced analytical approach, which identifies gene co-expression modules, together with the unique ability afforded by an ARR pathosystem to examine both susceptible and resistant responses within the same plant genotype, helps us gain valuable insight into the mechanisms that confer plant disease resistance.

When comparing the gene co-expression networks of the resistant and susceptible interactions, we first observed a large difference in the number of modules identified in each network. Furthermore, module preservation patterns suggested that the co-expression response in resistant-age fruit was reprogrammed as compared to the network on susceptible fruit (Figure 6). While co-expressed sets of genes share similar regulatory mechanisms, these are employed at different times and with different patterns during the infection of resistant and susceptible-aged fruit. Network reprograming in responses to infection by *P. syringae* was similarly observed when comparing resistant wild type plants and effector-triggered-immunity compromised mutants of Arabidopsis [41]. Specifically, similar groups of genes were shown to be activated in compatible- and incompatible-interactions; however, their timing and expression patterns were altered.

While we identified several modules that exhibit similar expression patterns regardless of infection (i.e. likely a response of fruit detached from the plant), most modules were impacted by infection (Figure 7). The differences in expression patterns observed between inoculated and non-inoculated tissue reveal we were successful at identification of defense response co-expression modules. We focused our analysis on the differences in gene expression patterns of genes identified to be induced earlier by inoculation in the resistant network. In PCA, susceptible samples at 2 hpi clustered with non-inoculated controls (Figure 5 A). Conversely, resistant-aged samples at 2 hpi showed a distinct transition along both PC1 and 2, away from the uninoculated controls, suggesting a more rapid transcriptional defense response in these samples. We thus were interested in modules showing differential expression patterns at very early timepoints. As optimal timing of defense response can be crucial for successful resistance [42], the ability to mount a successful defense could be attributed to this early response.

By performing cubic spline regression on module gene expression curves, we identified gene co-expression modules associated with defense that are differentially activated in resistant-aged fruit as early as 2 and 4 hpi (Figure 8). Of specific interest were module R4 and R7; in both cases increases in gene expression were delayed in susceptible relative to resistant-aged fruit. Genes of interest were identified using a combination of module membership statistics and differential expression analysis (Figure 9). Among the 34 candidate genes of interest showing resistance-specific increase at 2 or 4 hpi, were two WRKY transcription factor homologs *Csa1G025960* (*AtWRKY33*, BLAST E = 2.8e-119) and *Csa3G567330* (*AtWRKY75*, BLAST E = 7.4e-44). While *AtWRKY75* is reported to be associated with phosphate deficiency [43], its cucumber homolog may function in defense to pathogen infection. *AtWRKY33*, however, has been shown to be important in resistance to *Alternaria brassicicola* and *Botrytis cinerea* in Arabidopsis [44, 45]. *AtWRKY33* is also rapidly induced by the flg22 epitope as part of microbe-associated molecular pattern immunity, with downstream targets involved in ethylene and camalexin synthesis as well as other transcription factors and pathogen receptors [46]. Furthermore, in a proteomic study of *P. capsici* infection, the tomato WRKY33 homolog protein was found to be induced by 8 hpi and localized to the nucleus [47].

Among the targets of WRKY33 in Arabidopsis are ethylene biosynthesis genes [46]. We further identified that the two ethylene synthesis genes *Csa4G049610* and *Csa6G160180*, encoding 1-aminocyclopropane-1-carboxylate synthase and 1-aminocyclopropane-1-carboxylate oxidase, respectively, both have high module membership with Modules R4 and R7 respectively. Although ethylene is generally thought to be important in defense response against fungi and necrotrophic pathogens [48], ethylene response, but not SA or JA, was shown to be crucial for inhibition of *P. capsici* growth in habanero pepper, [49]. Blocking ethylene perception by means of exogenous application of silver nitrate reduced this inhibition [49]. Moreover, silencing of ethylene signal transduction in *Nicotiana benthamiana* resulted in loss of ARR to *P. infestans* [50]. The uniquely increased expression of the cucumber *WRKY33* and downstream upregulation of ethylene synthesis observed in resistant fruit could thus be a central component in regulation of the successful defense response in cucumber ARR.

Consistent with a successful hypersensitive response in resistant-age fruit, this group of early induced genes also included three putative peroxidases (*Csa6G213910, Csa4G285730* and *Csa6G495000*), an NAD(P)H-dehydrogenase (*Csa6G517010*), as well as an NADPH/respiratory burst oxidase protein (*Csa3G845500*) that could all potentially serve in modulating ROS within the first few hpi [51]. Other genes identified to be potentially involved in defense are genes involved in vesicle transport (*Csa3G271380, Csa3G782680* and *Csa6G431740*), as well as a signal transduction (*Csa1G086390, Csa3G219720 and Csa4G009900*) and specialized metabolism (*Csa2G069200, Csa4G640960* and *Csa4G411390*). Upregulation of *Csa2G069200*, a putative cinnamate-4-hydroxylase, could be acting either in specialized metabolite synthesis, or perhaps upstream of other enzymes in a lignification response. Finally, *Csa2G070840*, which putatively encodes a calcium-dependent phospholipid-binding copine family protein, might also be important in fine-tuning the response to infection, as its homolog in Arabidopsis functions in stomatal closing during infection and regulation of several resistance receptor genes [52, 53]. All these genes are canonically involved with response to pathogens, and so their early activation in response to inoculation in resistant fruit could be crucial in conferring ARR by limiting pathogen establishment in early stages of infection.

### Cucumber ARR to P. capsici may be mediated by developmental regulation of basal defenses and receptor-like genes

While it is tempting to speculate that resistances in young cucumber fruit may have been lost as a trade off in breeding for rapid growth, screening of the highly diverse USDA cucumber germplasm collection suggests against this idea, as the great majority are highly susceptible when very young [20]. On the other hand, while not all lines show as dramatic an ARR to *P. capsici* as ‘Poinsett 76’ or ‘Vlaspik’, other cultivars and wild accessions do show increased expression of defense genes after exponential growth [15], as well as a tendency to become less susceptible as they approach full size [13]. This further suggests that ontogenic defenses in cucumber fruit may result from a combination of an increase in preformed basal, physical and biochemical defenses common to fruit development, and in some genotypes, pathogen-specific, inducible resistance.

The distinctive biology of the cucumber ARR pathosystem and high-resolution probe of the initial period of infection allowed us to tease out transcriptional processes that are unique in resistant tissue, with minimal introduction of artifacts derived from genetic difference between resistant and susceptible materials. The rapid transcriptional defense response as early as 2-4 hpi and subsequent pathogen death in resistant-aged fruit indicate that upstream components in defense signaling, such as pathogen receptors or transcription factors, are developmentally regulated and enable pathogen sensing at the resistant age. In our previous analyses of ARR expressing fruit, we observed developmentally-regulated accumulation of chemical compounds with inhibitory effects on the *P. capsici* growth *in vitro* [14, 15]. We further showed a developmental upregulation of four pathogen receptors that was unique to the ARR expressing genotype [15]. Here we observed similar upregulation of specialized metabolism and defense in uninoculated 16 dpp fruit. Together, these earlier and current results suggest a model (Figure 10) in which accumulated antimicrobial specialized metabolites *and* an early response to infection mediated by developmentally-regulated expression of receptor-like genes, contribute to ARR. Transcription factors such as WRKY33 are expressed, and their downstream targets including ethylene synthesis genes and other defense genes are activated. In resistant-aged fruit, metabolism of ROS is also rapidly activated, likely further mediating a strong defense.

**Figure 10.**
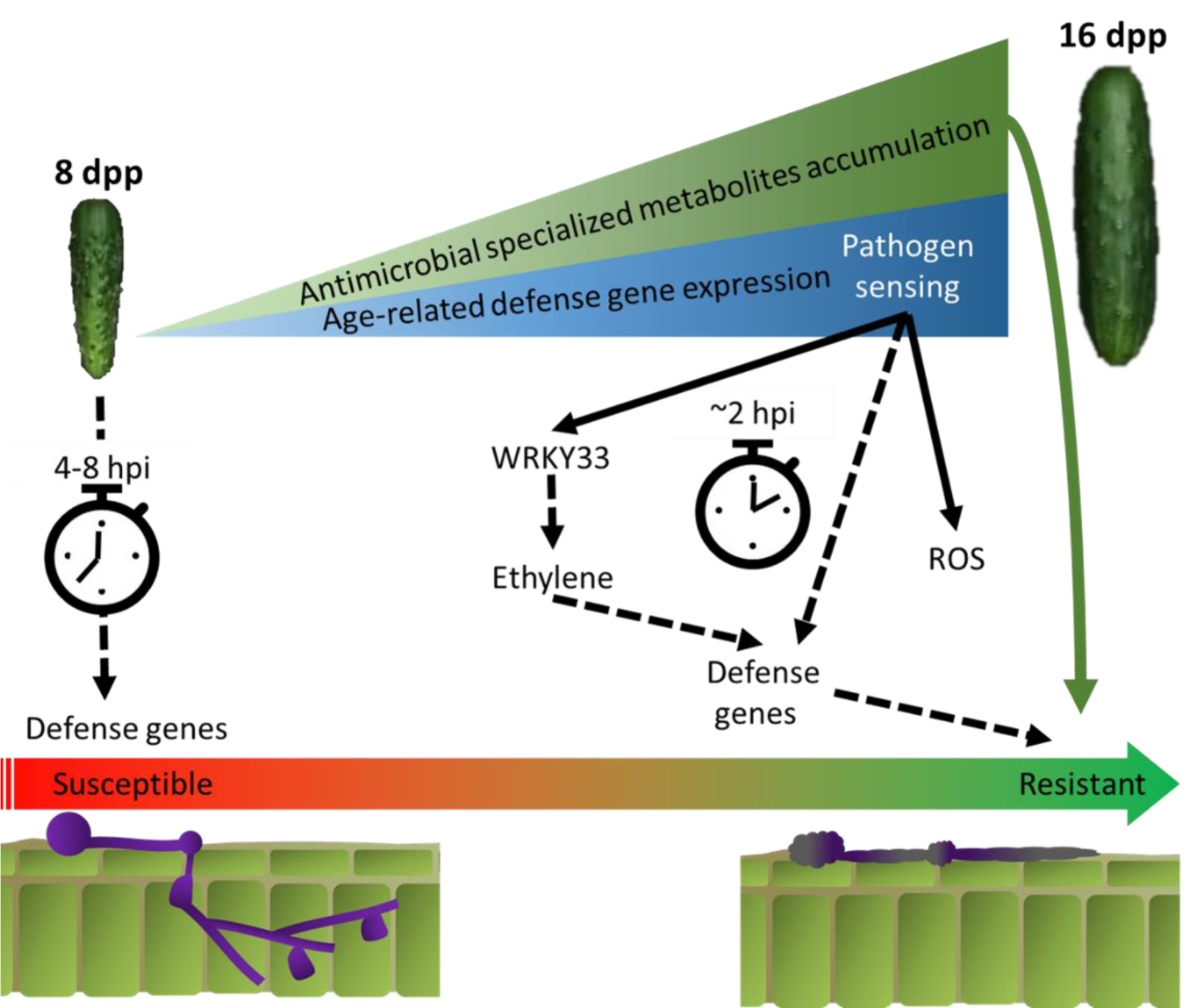
Hypothesized model for cucumber age-related resistance to *P. capsici*. In young susceptible fruit there is low accumulation of potentially antimicrobial specialized metabolites. Furthermore, a delayed response (> 8 hpi) to pathogen sensing may be too late to limit pathogen establishment. In resistant-aged fruit, the accumulation of metabolites could directly inhibit pathogen growth. Developmentally regulated expression of receptor-like gene(s) allows the sensing of pathogen-associated molecular patterns or effectors, and thus mediates an early response to infection. Transcription factors such as WRKY33 are expressed and their downstream targets including ethylene synthesis genes and other defense genes are activated. In resistant-aged fruit, metabolism of reactive oxygen species (ROS) is also activated in an early response to infection likely further mediating a strong defense response. By 24 hpi pathogen presence is limited on the fruit surface and mostly non-viable spores and hyphae remain.

## Conclusion

In conclusion, our studies indicate that ARR in the cucumber fruit - *P. capsici* system likely results from a combination of preformed and inducible defenses. Biochemical and other developmentally intrinsic defenses may act in concert with developmentally regulated expression of pathogen recognition-triggered, inducible responses. As production and maintenance of plant defenses is energy intensive and can be costly to fitness, an ideally timed and placed defense, is crucial for plant evolutionary success [9]. Plants expressing ARR optimize these defenses ontogenically, to express them at the opportune physiological age. For cucumber fruit, manifestation of these defenses coincides with the end of exponential fruit growth, protecting the developing seeds for the remaining weeks until the seeds and fruit reach maturity.

## Methods

### Plant Material

A set of 22 inbred cucumber cultivars was grown in the greenhouse as described in Ref. 21; three to ten fruits were collected from each cultivar at 16 dpp to test for ARR. Seeds of cucumber cv. ‘Poinsett 76’ were obtained from Seedway, LLC (Hall, NY). For all further experiments, greenhouse production of cucumber fruit (cv. ‘Poinsett 76’) were drip fertigated (1 L/day at 1-2% 20-20-20 fertilizer). In transcriptome experiment 1, two flowers were hand pollinated per plant, while in transcriptome experiment two flowers were tagged at anthesis and bee pollinated. In both experiments pollination was staggered, such that 8 and 16 dpp fruit were harvested on the same day. After fruit set only one fruit per plant was retained; any other fruits were removed, to prevent competition.

### Detached fruit inoculations and sample collection

Harvested fruit were briefly washed with distilled water and allowed to air-dry. Fruits were placed in incubation trays lined with wet paper towels, to maintain high humidity and covered with clear plastic tops. Zoospore suspensions were prepared from *P. capsici* isolate OP97 or NY-0644-1 expressing RFP [25] cultured on diluted V8 agar media (V8 juice 200 mL, CaCO_3_ 3 g, agar 15 g, distilled water 800 mL). After 7 days, the plates were flooded with 10 mL sterile distilled water to release zoospores. Two 10 µl aliquots of zoospore suspension were quantified using a Countess Cell Counter (Invitrogen) and the mean concentration was used for dilution. The suspension was diluted to a concentration of 5 × 10^5^ zoospores/mL. Fruits were then inoculated with ∼6 (8 dpp fruit) or ∼12 (16 dpp fruit), equally spaced, 30 µL droplets of the diluted zoospore suspension. Incubation was performed under constant light at 23-25 °C. For ARR screening development of disease symptoms such as water soaking and mycelial growth on each fruit was monitored daily for ten days. Fruits were evaluated using a disease rating in scale of 1-9 (1=no symptom; 9=extensive mycelial growth and sporulation).

Plant material was inoculated as described above for two transcriptome experiments; the first included fruit sampled at 0, 4, 24, and 48 hours post inoculation (hpi), and the second at 0, 2, 4, 8, 12, 18, and 24 hpi. In both experiments timepoint 0 was collected at 12:30 pm. At each subsequent timepoint, samples were collected from 6-12 inoculation sites per fruit. Samples from a given fruit were pooled to form a biological replicate. Three replicate fruits were sampled for each age at each timepoint. Thus, each timepoint had 3 biological replicates, each comprised of multiple inoculated tissue samples. In experiment 2, at timepoint 0, the three replicate samples were prepared from a single fruit. Fruits were removed from the incubation chamber and punches were made around each inoculation site using a No. 4 cork borer. Peel discs were subsequently collected by peeling the punched area using a vegetable peeler and immediately frozen in liquid nitrogen and stored at -80°C until RNA extraction. In experiment 2, samples taken from uninoculated parts of the fruit were used as the respective control for each time point.

### High-throughput RNA extraction

Samples were ground using a mortar and pestle in liquid nitrogen. RNA extraction was performed using the MagMAX Plant RNA Isolation Kit protocol (Thermo Fisher) with slight modifications: 100-150 mg of ground tissue were added to 1000 µL of lysis buffer. High-throughput RNA extraction was performed in 96-well format, on a KingFisher Flex Purification System (Thermo Fisher). Immediately after the run was complete, the 96-well plate was transferred to storage at -80°C. RNA concentration and quality were measured using Qubit 2.0 Fluorometer (Life Technologies, Inc.) and LabChip GX (Perkin Elmer) respectively. All samples had a minimum RNA quality score of 8.

### TruSeq Library preparation and sequencing

Libraries were prepared at Michigan State University’s Research Technology Support Facility, using the Illumina TruSeq Stranded mRNA Library Preparation Kit on a Sciclone G3 robot following manufacturer’s recommendations. An additional cleanup with 0.8X AmpureXP magnetic beads was performed after completion of library preparation. Quality control and quantification of completed libraries was performed using a combination of Qubit dsDNA HS and Advanced Analytical Fragment Analyzer High Sensitivity DNA assays. The libraries were divided into two pools of 15 libraries each. Pools were quantified using the Kapa Biosystems Illumina Library Quantification qPCR kit. Each pool was loaded onto one lane of an Illumina HiSeq 4000 flow cell and sequencing was performed in a 1×50 bp single read format using HiSeq 4000 SBS reagents. Base calling was accomplished by Illumina Real Time Analysis (RTA) v2.7.7 and output of RTA was demultiplexed and converted to FastQ format with Illumina Bcl2fastq v2.19.1.

### QuantSeq 3’-mRNA library preparation and sequencing

For the second experiment, QuantSeq 3’-mRNA FWD libraries (Lexogen) were prepared by the Cornell University, Institute of Biotechnology, Genomics Facility using the manufacturers guidelines. Quality control and quantification of completed libraries was performed using a combination of Qubit dsDNA HS and Advanced Analytical Fragment Analyzer High Sensitivity DNA assays. The libraries where then loaded on a single Illumina NextSeq500 lane and sequenced in a 1×86 bp single end format. Base calling was achieved by Illumina RTA v2.4.11 and output of RTA was demultiplexed and converted to FastQ format with Illumina Bcl2fastq v2.18.

### Sequencing read preprocessing and quasi-mapping

Experiment 1: Reads were cleaned, and adaptor sequences were removed using Trimmomatic v. 0.34 [54] with the following settings: LEADING:3 TRAILING:3 SLIDINGWINDOW:4:15 MINLEN:35. Quality control was performed using FastQC (http://www.bioinformatics.bbsrc.ac.uk/projects/fastqc). A cucumber transcriptome fasta file was made from the ‘Chinese Long’ (v2) [55, 56] genome using the *gffread* function from the cufflinks software package [57] and high-quality reads were then quasi-mapped to the transcriptome using Salmon v. 0.9.1 [58] with default settings.

Experiment 2: The quality of reads was assessed with FastQC and visualized using multiQC [59]. Subsequently, reads were processed using BBMap [60] with the following settings: ftl=12; k=13; ktrim=r; useshortkmers=t; mink=5; qtrim=r; trimq=10; minlength=20; int=f, and trimmed of any poly A sequences, adaptors and the first 12 nt (as recommended by the manufacturer of the library kit). To increase the mapping success rate, the ‘Chinese Long’ (v2) GFF3 file was amended to extend all transcript 3’UTRs by 1000 bases using a custom R script. If a gene model existed prior to 1000 bases, on the same strand, then the 3’UTR was extended only until 1 base before that next gene. The extended gene models were then used to extract a transcriptome fasta file as above. Reads were then quasi-mapped to this new transcriptome file using Salmon v 0.12.0 with the --noLengthCorrection option. Efficacy of Salmon for mapping 3’RNA reads was benchmarked by comparing to whole genome alignment using BWA [61] and htseq-count [62], with comparable results.

### Differential expression analysis

Read quantification data was imported into R using the tximport R package [63] and differential expression analysis was performed using DEseq2 [64] with log-fold-change-shrinkage. Contrasts were analyzed comparing sequential timepoints as well as each timepoint vs. uninoculated samples. DEG were called significant using an adjusted p-value (Benjamini-Hochberg adjustment [65]) and a false discovery rate of less than 5%. A cutoff expression change of above two-fold was used to define biological significance. Alluvial plots were drawn using the ggalluvial R package and venn diagrams were created using the overLapper script from Ref. 64.

### Weighted gene co-expression network analysis

Integer value transcript counts from experiment two, were imported into DESeq2 using the tximport R package [63]. Genes with less than 5 reads in greater than 75 of the total 78 samples were considered lowly expressed and excluded from the analysis, and 15202 genes remained. The normalized counts matrix was then transformed using the variance stabilizing transformation (VST) [67] using DESeq2 and imported into the WGCNA package pipeline [26].

Two separate signed networks were assigned for the inoculated susceptible- and resistant-aged fruit. For each network, the VST counts were used to calculate adjacency matrices using the biweight midcorrelation and a soft thresholding power of 12 (yielding a scale free topology fit of greater than 0.8). The adjacency matrices were used to calculate two topological overlap dissimilarity matrices which were subsequently used for forming gene clustering trees, using average distances. The gene trees were used for assigning co-expression modules using the dynamic tree cut algorithm with a minimum module size of 30 genes. Module eigengenes were correlated to each other and modules with similar expression patterns (dissimilarity < 0.25) were merged. Gene expression profiles of module genes from the infected resistant network (16 dpp) were plotted based on VST values and compared to control and 8 dpp expression patterns.

To identify modules with different expression patterns in inoculated tissue, read counts were first extracted and normalized by library size using DESeq2 *counts()* function. For each of the resistant network modules, the log_2_ (+ 0.5) of the normalized counts of all genes in that module was the dependent variable in a linear model where a natural cubic spline with 3 internal knots, at 3, 8, 15 hpi (as determined by the 0.25, 0.5 and 0.75 quartiles of time), was applied to the time variable using the *ns()* function from the splines R package:

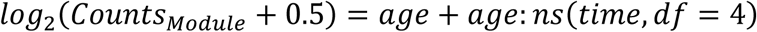

An analysis of variance was then performed to identify module with significant age × splined-time interactions. The summary of each of those linear models contains the interaction effects for each segment of the spline. Modules with interaction effects in segment 1 and/or 2 (0-3 and 3-8 hpi, respectively) were identified as early induced modules.

### Gene ontology term enrichment analysis

Gene Ontology (GO) term enrichment analysis was performed using the TopGO R package [68] with the entire set of fruit peel expressed genes set as background. Terms were considered enriched if they passed a *p*-value of 0.05 on the Fisher test with the “*weight01*” algorithm and a minimum node size of 100 genes. The previously updated GO term list for cucumber genes [15], was used for analysis. To visualize change of GO terms over consecutive contrasts heatmaps of -log_10_(Fisher *weight01 p*-values) were plotted using only terms with *P* < 0.01.

### Microscopy

Preliminary fluorescent microscopy of infection was performed using an EVOS FL Auto imaging system (ThermoFisher). Excised cucumber peels were affixed to the lid of a 100 mm petri dish using petroleum jelly and inoculated with 10 µL zoospore suspension (∼5 × 10^5^ spores/ml) of RFP-expressing isolate NY-0664-1 [25]. Petri dishes were then sealed with parafilm and carefully inverted and placed on the microscope table. Samples were observed at 2 and 4x magnification and images were captured every 30 minutes for 72 hours. Additional images were taken at 20x magnification at 4 hpi.

While collecting inoculated samples for transcriptome experiment 2, a ∼2 mm peel plug from the middle section of each fruit was also excised using a razor blade and fixed in 4% glutaraldehyde in 0.1M phosphate buffer for scanning electron microscopy. After overnight fixation in glutaraldehyde, samples were soaked in 0.1 M phosphate buffer for 40min. After consecutive dehydration in rising ethanol concentrations (25, 50, 75, 90, 100, 100, 100%; 1 hour each), samples were transferred to a Leica Microsystems EM CPD300 critical point dryer (Leica Microsystems) using liquefied carbon dioxide as the transitional fluid. Samples were then mounted on aluminum stubs using adhesive tabs (M.E. Taylor Engineering) and coated with osmium (∼10 nm thickness) in an NEOC-AT osmium coater (Meiwafosis Co., Ltd.). Samples were examined in a JEOL JSM-6610LV scanning electron microscope (JEOL Ltd.).

### High-throughput infection phenotyping

High-throughput *in vivo* disease phenotyping was performed as described in Ref. 67. Briefly, sixteen 6 mm diameter, 5 mm thick, peel tissue plugs were collected from each of three 8 and 16 dpp fruit using a biopsy punch. Plugs were placed in a 96-well black plate and subsequently inoculated with the constitutively fluorescing *P. capsici* isolate NY-0664-1 [25], or with distilled water (control - 4 plugs/fruit). Plates were read using a Tecan Spark Plate Reader (Tecan). Fluorescent measurements were taken in each well every hour, over the course of 24 hours at 28 °C. The excitation and emission settings were 536 and 586 nm, respectively. Gain was calculated from a well containing a mycelial mat, and the Z-position was set at 20000 µm.

## Supplemental Data

Supplementary Tables and Figures

Supplementary File 1: All code and analysis

Supplementary File 2: Genes involved in an early incompatible interaction

Supplementary File 3: Gene module assignment and normalized expression – resistant network

Supplementary File 4: Gene module assignment and normalized expression – susceptible network

Supplementary File 5: GO-term enrichment analysis of the resistant network modules

## List of abbreviations

ARR: Age-related resistance
DEG: Differentially expressed genes
dpp: Days post-pollination
GO: Gene Ontology
hpi: hours post-inoculation
ROS: Reactive oxygen species
SEM: Scanning electron microscopy
VST: Variance stabilizing transformation
PCA: Principal component analysis
WGCNA: Weighted Gene Co-expression Network Analysis

## Declarations

### Ethics approval and consent to participate

Not applicable

### Consent for publication

Not applicable

### Availability of data and materials

Raw reads for both experiments are deposited in the NCBI Sequence Read Archive (SRA) database under the accession number PRJNA575868.

**READ ONLY LINK FOR REVIEWERS:

https://dataview.ncbi.nlm.nih.gov/object/PRJNA575868?reviewer=a1174d4r0466lqonfcl6aa70od

All code and scripts for the entire analysis are available in Supplementary File 1 and at: https://github.com/bmansfeld/cucumberPcapInfection

## Author Contributions

BNM and RG conceived of the research and wrote the manuscript. BNM performed the RNA-seq experiments, transcriptome and network analyses. MC screened the cultivar collection for ARR and performed the developmental phenotyping for ARR in ‘Poinsett 76’. CZ and BNM developed and performed the *in vivo* bioassay. YL and BNM performed the fluorescent microscopy experiments. All authors have read and approved the manuscript.

## Acknowledgments

We thank the Sue Hammar for greenhouse and experiment assistance, and the Michigan State University (MSU) Research Technology Support Facility Genomics Core and Cornell University, Institute of Biotechnology, Genomics Facility, for library preparation and sequencing. We thank the MSU Center for Advanced Microscopy for scanning electron microscopy work. We also acknowledge the Michigan State University Research Center for Statistical Training and Consulting. Finally, we thank Drs. Cornelius Barry, Robin Buell, Brad Day, Sheng Yang He and Robert VanBuren for critical reading of the manuscript.

