## Supplementary Material for "Developmentally regulated activation of defense allows for rapid inhibition of infection in age-related resistance to *Phytophthora capsici* in cucumber fruit"

**Supplementary Table 1. Cucumber cultivars tested for age-related resistance to *P. capsici.*** Each line was tested using 3-10 fruit and was evaluated using a disease rating in scale of 1-9 (1=no symptom; 9=extensive mycelial growth and sporulation).

| **Cucumber Varieties** | **Disease rating**  **16 dpp** | **ARR (R/S)** | |
| --- | --- | --- | --- |
|  |  | 8 dpp | 16 dpp |
| Ashley | 7.0±0.0 | S | S |
| Boston Pickling | 5.0±0.7 | S | S |
| Boston Pickling Improved | 7.0±0.0 | S | S |
| Certified Organic Boothby's | 8.0±0.0 | S | S |
| Chinese Long | 6.5±0.5 | S | S |
| Delikatesse | 7.0±0.0 | S | S |
| Gy14 | 7.7±0.1 | S | S |
| Homemade Pickles | 5.1±0.1 | S | S |
| **Long Green Improved** | 2.5±0.3 | S | **R** |
| Miniature White | 7.6±0.4 | S | S |
| Muncher | 7.5±0.3 | S | S |
| National Pickling | 7.3±0.3 | S | S |
| Parisian Pickling | 7±0.0.0 | S | S |
| **Pointsett 76** | 3.0±0.2 | S | **R** |
| Rhinish Pickle | 4.5±1.0 | S | S |
| Russian Cucumber | 6.1±0.8 | S | S |
| Spacemaster 80 | 6.5±0.4 | S | S |
| Tanja | 8.0±0.0 | S | S |
| Tendergreen Burpless | 8.0±0.0 | S | S |
| **Vlaspik** | 3.0±0.9 | S | **R** |
| White Wonder | 7.3±0.2 | S | S |
| Zarnista | 4.4±0.5 | S | S |

**Supplementary Table 2. Top 15 most enriched Gene Ontology terms in 16 dpp vs. 8 dpp uninoculated fruit peels**

| Direction | GO.ID | Term | Annotated | Significant | Expected | Fisher.weight01 |
| --- | --- | --- | --- | --- | --- | --- |
| Up | GO:0019748 | secondary metabolic process | 251 | 26 | 11.79 | 7.50E-05 |
| Up | GO:0006518 | peptide metabolic process | 643 | 16 | 30.21 | 0.00016 |
| Up | GO:0044703 | multi-organism reproductive process | 136 | 17 | 6.39 | 0.00044 |
| Up | GO:0006790 | sulfur compound metabolic process | 256 | 24 | 12.03 | 0.00057 |
| Up | GO:0051186 | cofactor metabolic process | 486 | 30 | 22.83 | 0.00072 |
| Up | GO:0008152 | metabolic process | 9213 | 451 | 432.83 | 0.00109 |
| Up | GO:0014070 | response to organic cyclic compound | 294 | 28 | 13.81 | 0.00189 |
| Up | GO:0040008 | regulation of growth | 181 | 17 | 8.5 | 0.00515 |
| Up | GO:0048364 | root development | 376 | 24 | 17.66 | 0.00518 |
| Up | GO:0055114 | oxidation-reduction process | 1249 | 78 | 58.68 | 0.00569 |
| Up | GO:0015893 | drug transport | 112 | 12 | 5.26 | 0.00633 |
| Up | GO:0009723 | response to ethylene | 254 | 19 | 11.93 | 0.00806 |
| Up | GO:0006575 | cellular modified amino acid metabolic p... | 102 | 11 | 4.79 | 0.00835 |
| Up | GO:0046777 | protein autophosphorylation | 131 | 13 | 6.15 | 0.00868 |
| Up | GO:0050832 | defense response to fungus | 239 | 20 | 11.23 | 0.00911 |
| Down | GO:0019684 | photosynthesis, light reaction | 115 | 66 | 16.73 | 1.40E-26 |
| Down | GO:0015979 | photosynthesis | 201 | 113 | 29.25 | 1.60E-18 |
| Down | GO:0055114 | oxidation-reduction process | 1249 | 275 | 181.75 | 9.80E-14 |
| Down | GO:0000272 | polysaccharide catabolic process | 130 | 44 | 18.92 | 2.50E-08 |
| Down | GO:0044271 | cellular nitrogen compound biosynthetic ... | 2861 | 382 | 416.32 | 7.10E-07 |
| Down | GO:0071555 | cell wall organization | 323 | 81 | 47 | 7.60E-07 |
| Down | GO:0051188 | cofactor biosynthetic process | 272 | 63 | 39.58 | 3.00E-06 |
| Down | GO:0006073 | cellular glucan metabolic process | 171 | 48 | 24.88 | 3.30E-06 |
| Down | GO:0016114 | terpenoid biosynthetic process | 110 | 35 | 16.01 | 3.30E-06 |
| Down | GO:0005975 | carbohydrate metabolic process | 804 | 189 | 116.99 | 3.30E-06 |
| Down | GO:0048646 | anatomical structure formation involved ... | 167 | 47 | 24.3 | 3.80E-06 |
| Down | GO:0006633 | fatty acid biosynthetic process | 167 | 47 | 24.3 | 3.80E-06 |
| Down | GO:0051276 | chromosome organization | 343 | 69 | 49.91 | 6.40E-06 |
| Down | GO:0009658 | chloroplast organization | 148 | 42 | 21.54 | 9.70E-06 |
| Down | GO:0042446 | hormone biosynthetic process | 185 | 49 | 26.92 | 1.50E-05 |

**Supplementary Table 3. Modules with significant age X splined-time interactions.** Interaction effects from ANOVA results with p.value < 0.05.

| Module Label | Sum squares | Mean squares | F statistic | p.value |
| --- | --- | --- | --- | --- |
| 1 | 329.9997 | 82.49991 | 15.93437 | <1E-12 |
| 2 | 314.3932 | 78.5983 | 14.95523 | <1E-12 |
| 3 | 556.1415 | 139.0354 | 27.25719 | <1E-12 |
| 4 | 1372.471 | 343.1178 | 64.26949 | <1E-12 |
| 5 | 330.5371 | 82.63427 | 14.55474 | <1E-12 |
| 6 | 405.3507 | 101.3377 | 23.74939 | <1E-12 |
| 7 | 940.1047 | 235.0262 | 42.18563 | <1E-12 |
| 8 | 126.2244 | 31.5561 | 5.047284 | 0.00046 |
| 9 | 73.84148 | 18.46037 | 3.210159 | 0.012096 |
| 10 | 172.7907 | 43.19768 | 8.52806 | 7E-07 |
| 11 | 135.2212 | 33.80529 | 7.716836 | 3.3E-06 |
| 12 | 503.9448 | 125.9862 | 25.39578 | <1E-12 |
| 13 | 223.3057 | 55.82643 | 13.36232 | <1E-12 |
| 14 | 106.5474 | 26.63684 | 6.508226 | 3.15E-05 |
| 15 | 207.5917 | 51.89793 | 10.68372 | <1E-12 |
| 16 | 72.2291 | 18.05727 | 4.95054 | 0.00055 |
| 18 | 98.17228 | 24.54307 | 5.643348 | 0.000156 |
| 19 | 68.86208 | 17.21552 | 3.174732 | 0.012901 |
| 20 | 55.10566 | 13.77642 | 3.567641 | 0.006519 |
| 21 | 41.93763 | 10.48441 | 2.463965 | 0.043103 |
| 22 | 52.10105 | 13.02526 | 3.387247 | 0.008964 |
| 23 | 41.79785 | 10.44946 | 3.288753 | 0.010641 |
| 27 | 104.5677 | 26.14192 | 11.52098 | <1E-12 |
| 29 | 410.3042 | 102.5761 | 19.21226 | <1E-12 |

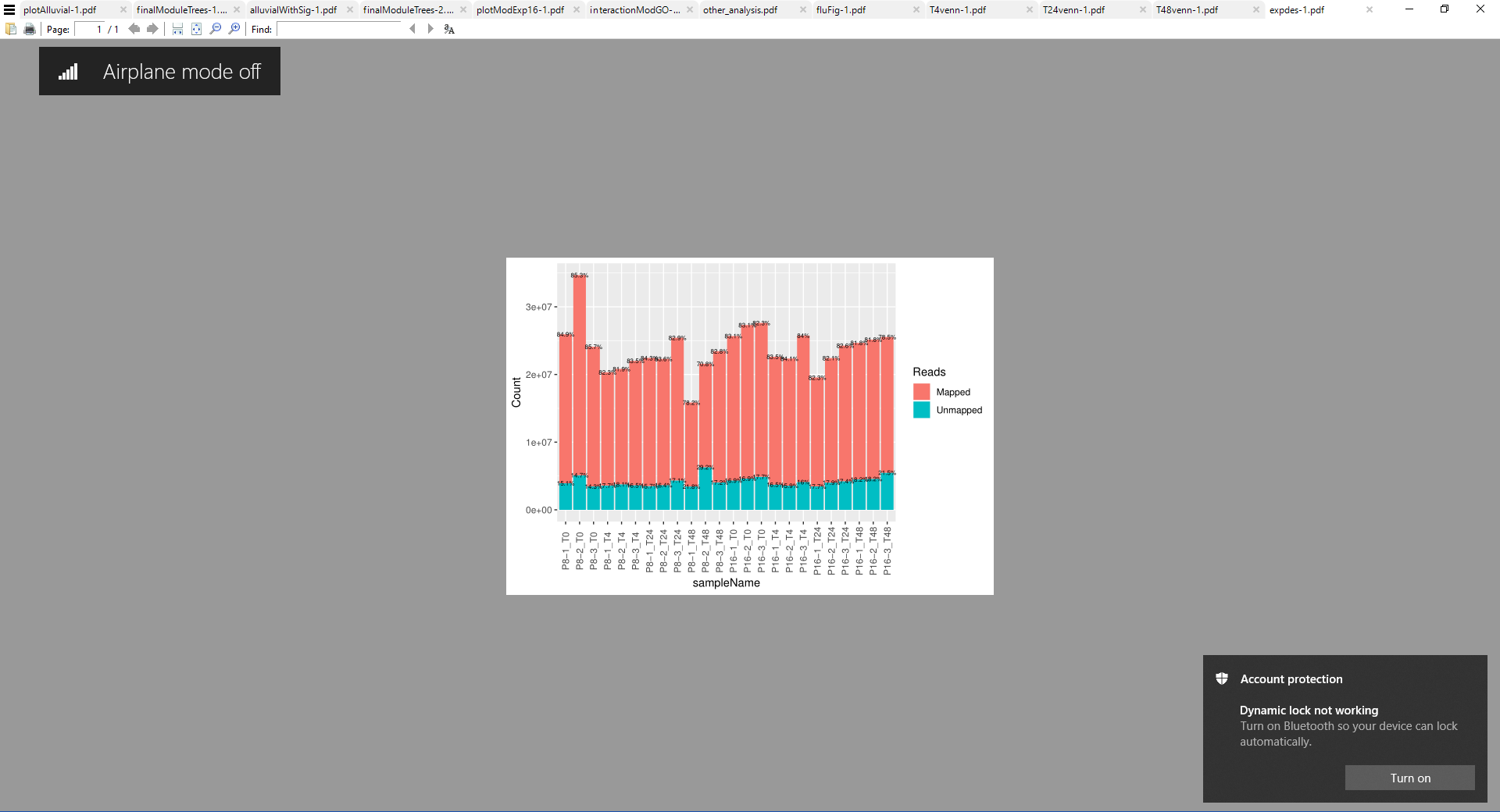
**Supplementary Figure 1. Transcriptome experiment 1: RNAseq Reads quasi-mapping to the cucumber transcriptome**

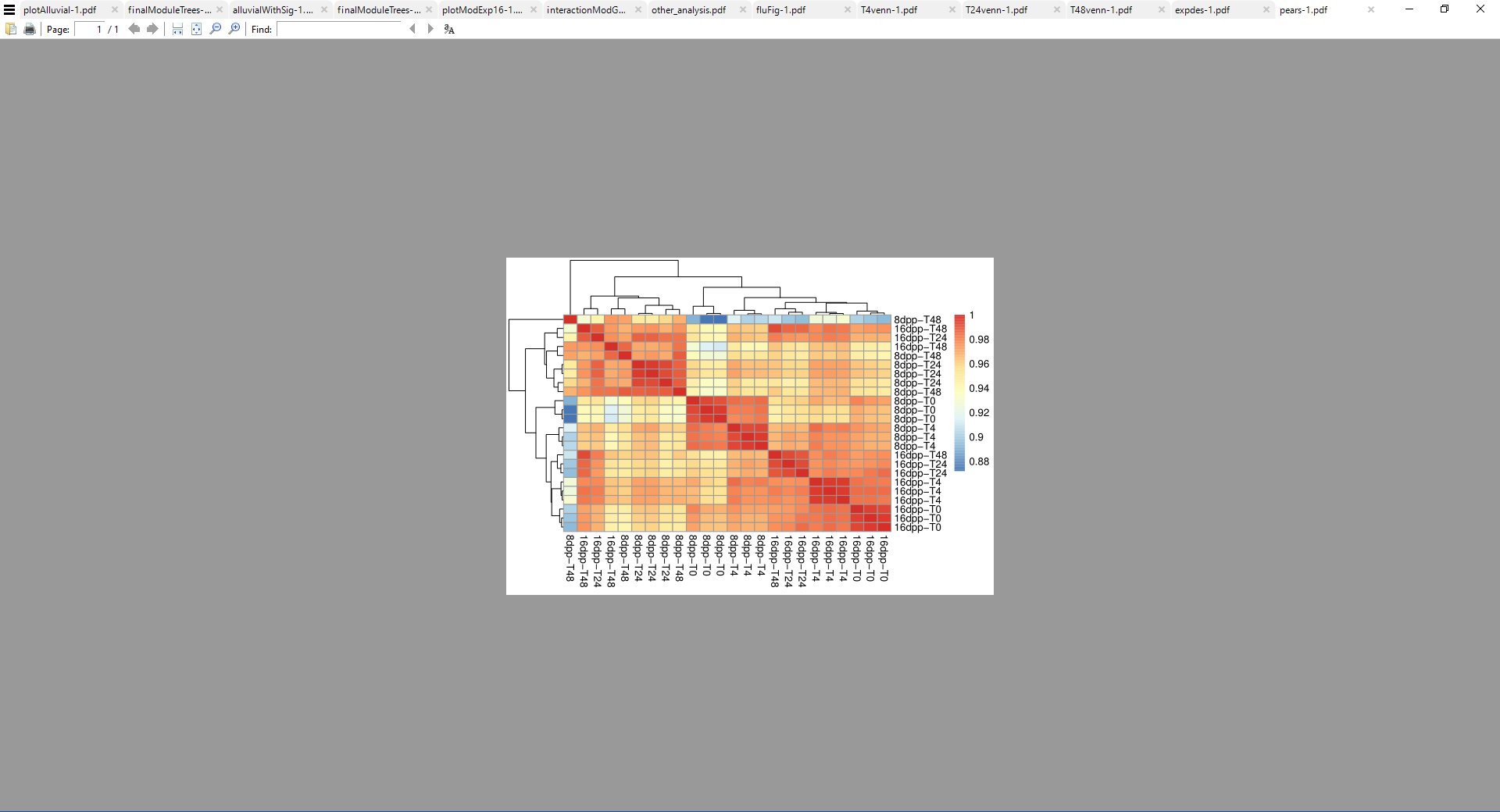

**Supplementary Figure 2. Between sample Pearson’s correlations**

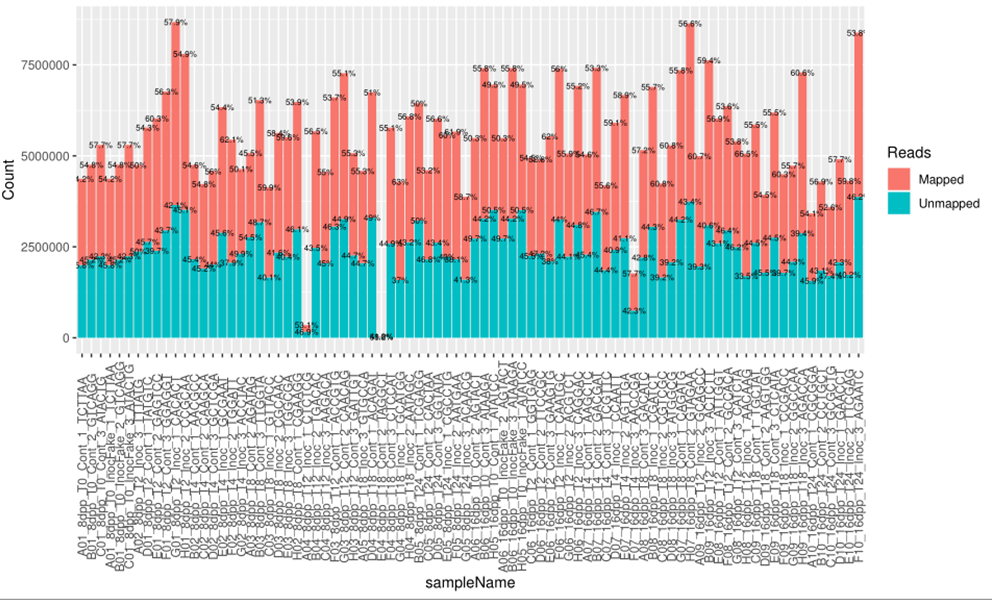
**Supplementary Figure 3. Transcriptome experiment 2: 3’mRNA-seq reads quasi-mapping to a 3’-extended cucumber transcriptome.**

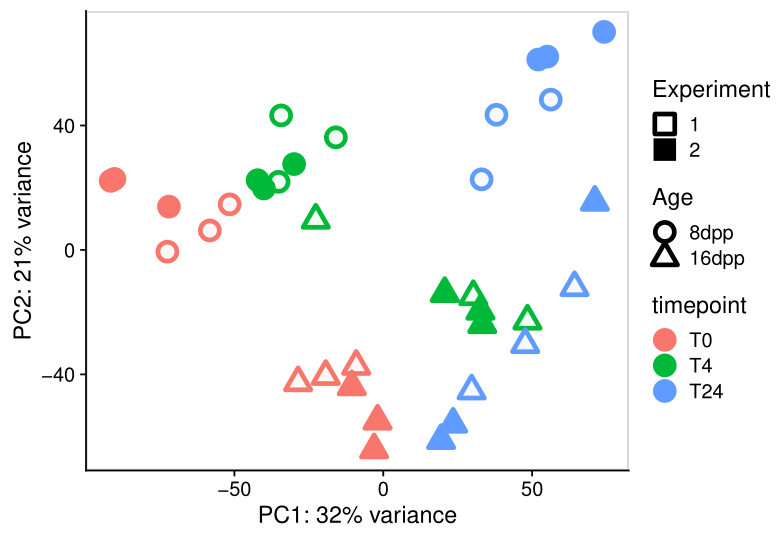
**Supplementary Figure 4. Principal component analysis of shared timepoints in both transcriptome experiments performed in different years with different sequencing library technologies.** Samples at timepoint 0 and inoculated samples at 4 and 24 hpi of 8 dpp and 16 dpp fruit. Variance stabilizing transformed counts from both experiments (Exp1 – Illumina TruSeq mRNA; Exp2 – Lexogen 3’-mRNA QuantSeq) were adjusted to account for experimental batch effects using the function *removeBatchEffects()* from the ‘limma’ R package.

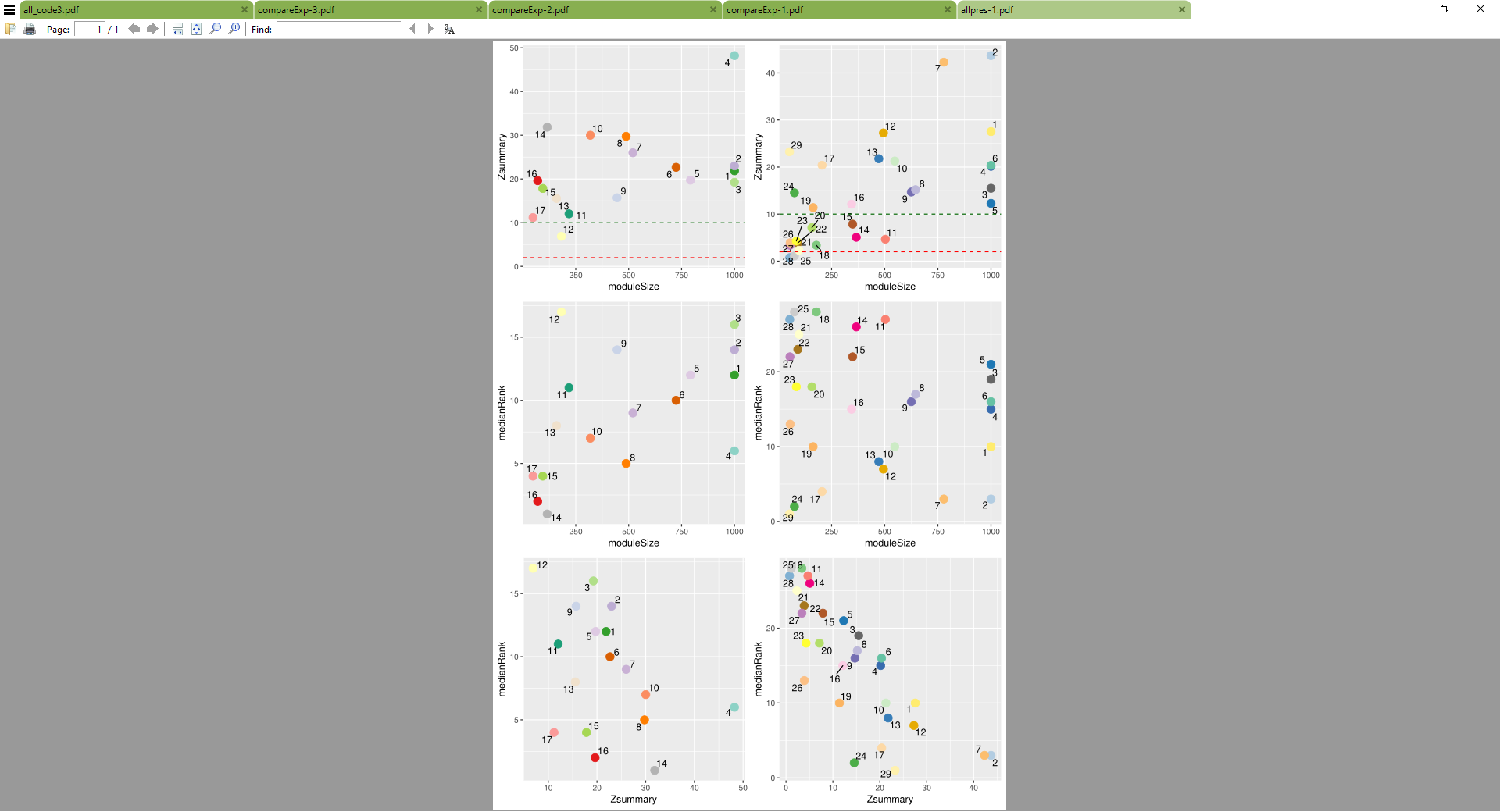
**Supplementary Figure 5. Module preservation statistics for susceptible and resistant weighted gene co-expression networks.** Zsummary scores by module size for 8 dpp (A) and 16 dpp (B). Median Rank scores by module size for 8 dpp (C) and 16 dpp (D). Median rank scores by Zsummary scores for 8 dpp (E) and 16 dpp (F).

**A**

**C**

**E**

**B**

**D**

**F**

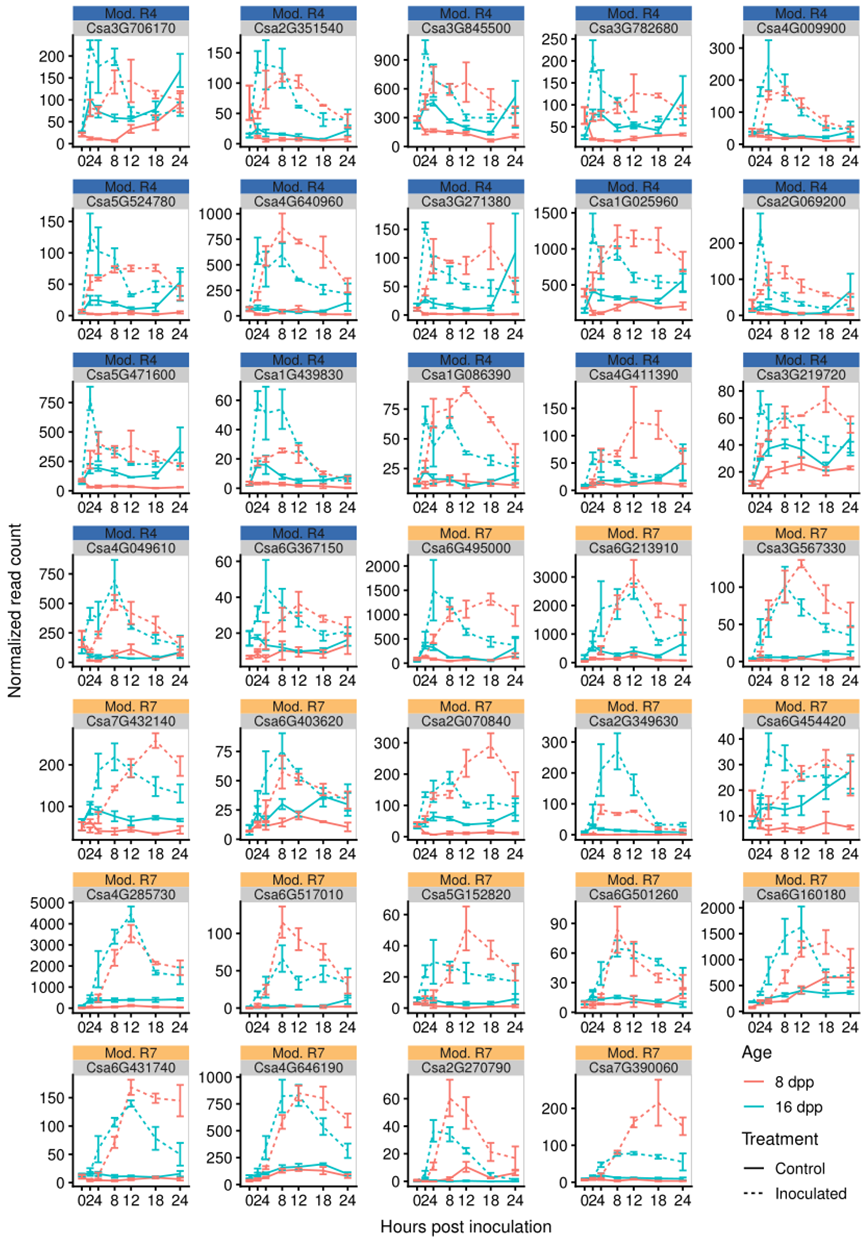
**Supplementary Figure 6. Genes induced in early responses to inoculation of resistant fruit.** All Genes identified by a module membership > 0.75 and significant fold change of >2 in inoculated resistant fruit (16 dpp) compared to both control resistant fruit and inoculated susceptible fruit (8 dpp). Mean expression for three biological replicates in each age and treatment. error bars are +/- standard error of the mean.

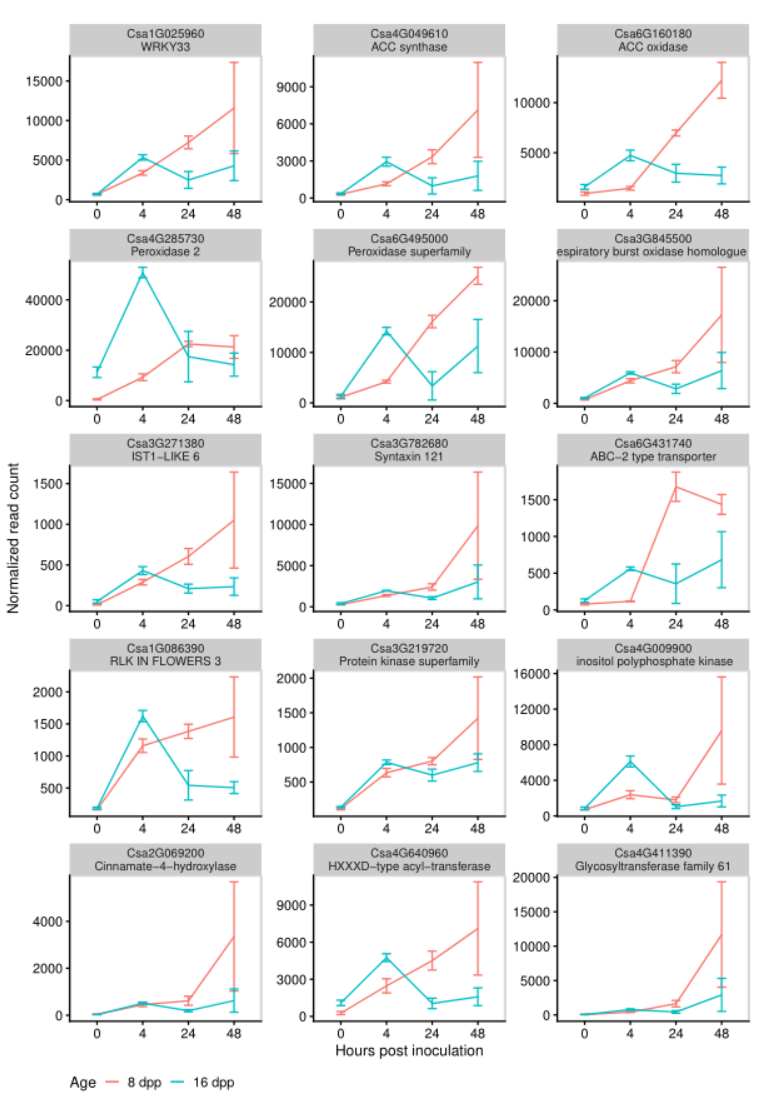

**Supplementary Figure 7. Validation of expression patterns of genes induced in early responses to inoculation of resistant fruit.** Expression patterns in RNA-seq experiment 1 of a subset of genes identified in RNA-seq experiment 2. The experiments were performed in a different year and with a different sequencing library technique. Normalized gene expression of inoculated resistant (16 dpp) and susceptible fruit (8 dpp). Mean expression for three biological replicates in each age and treatment. Error bars are +/- standard error of the mean.

**Supplementary Figure 8. Between experiment correlation of expression levels of genes induced in early responses to inoculation of resistant fruit.** Log10 of average read counts RNA-seq experiment 1 of a subset of genes identified in RNA-seq experiment 2. At each age and timepoint the average read count of three replicates was compared between experiments. The experiments were performed in a different year and with a different sequencing library technique. Pearson correlation R and p-value reported in the figure.

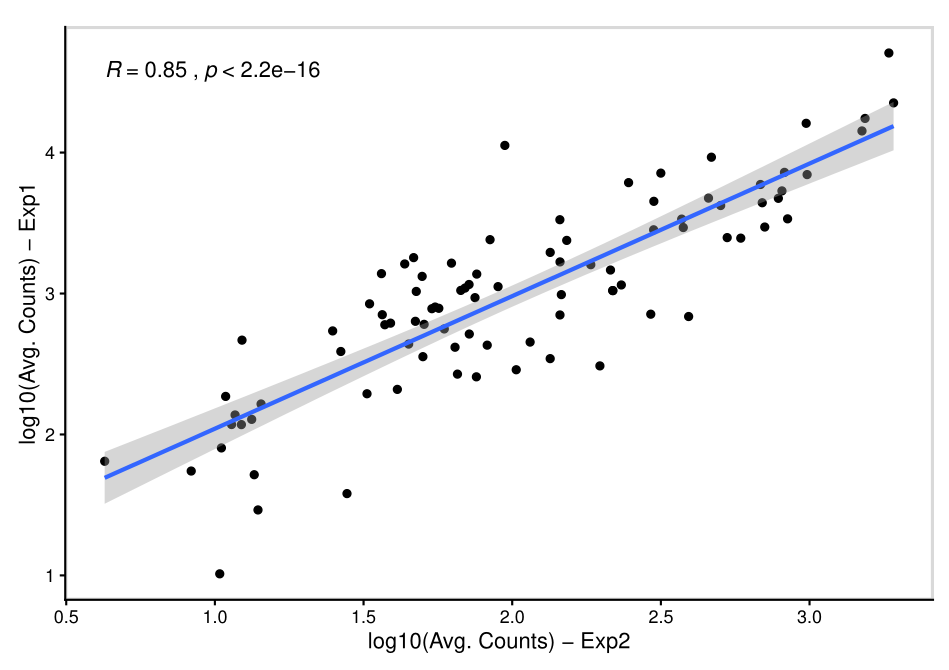
