## Supplementary File 1 for "Developmentally regulated activation of defense allows for rapid inhibition of infection in age-related resistance to *Phytophthora capsici* in cucumber fruit"

Supplementary File 1: Analysis workflow

Ben N. Mansfeld, Marivi Colle, Chunqiu Zhang, Ying-Chen Lin, Rebecca Grumet

04/07/2019

```
# source("https://bioconductor.org/biocLite.R")
# biocLite("tximport")
# bioCLite("impute")
library(tidyverse)
library(UpSetR)
library(grid)
library(tximport)
library(stringr)
library(readr)
library(DESeq2)
library(kableExtra)
library(rjson)
library(ggalluvial)
library(WGCNA)
library(pheatmap)
library(RColorBrewer)
library(splines)
library(rvest)
library(readxl)
library(magick)
library(topGO)
```

#### Physiology and other experiments

Disease rating vs. growth

```
loaded <- read.csv(file = "diseaserating_length_marivi.csv", header = TRUE)

drPlot <- ggplot(loaded) +
    stat_summary(
        aes(x = Age, y = `Disease.rating`, linetype = "Disease rating"),
        fun.y = mean,
        geom = "line",
        na.rm = TRUE,
        size = 1
    ) +
```

```
    stat_summary(
        aes(x = Age, y = `Disease.rating`),
        fun.y = mean,
        geom = "point",
        na.rm = TRUE,
        size = 2
    ) +
    stat_summary(
        aes(x = Age, y = `Disease.rating`),
        fun.data = mean_se,
        geom = "errorbar",
        na.rm = TRUE,
        width = 0.2
    ) +
    stat_summary(
        aes(
            x = Age,
            y = `Length` / 2.777,
            linetype = "Fruit length"
        ),
        fun.y = mean,
        geom = "line",
        na.rm = TRUE,
        size = 1
    ) +
    stat_summary(
        aes(x = Age, y = `Length` / 2.777),
        fun.y = mean,
        geom = "point",
        na.rm = TRUE,
        size = 2
    ) +
    stat_summary(
        aes(x = Age, y = `Length` / 2.777),
        fun.data = mean_se,
        geom = "errorbar",
        na.rm = TRUE,
        width = 0.2
    ) +
    scale_y_continuous(
        name = "Disease rating (1-9)",
        sec.axis = sec_axis( ~ . * 2.777, name = "Length (cm)"),
        limits = c(1, 9)
    ) +
    scale_linetype() +
    geom_hline(yintercept = 3, linetype = 2) +
    cowplot::theme_cowplot(font_size = 14) +
    theme(
        legend.title = element_blank(),
        legend.position = c(0.7, 0.1),
        legend.direction = "vertical"
    )
```

Fluorecent plate reader data:

```r
rawData <- readxl::read_xlsx(path = "flu_2018-10-6_raw_data.xlsx",
                             sheet = 1,
                             skip = 61,
                             n_max = 1822,
                             col_names = FALSE)

wells <- rawData[seq(from = 1, to = nrow(rawData), by = 19), 1, drop = TRUE]
time <- as.numeric(rawData[2, 2:ncol(rawData)])

df <- rawData[rep(seq(from = 0, nrow(rawData) - 1, by = 19), each = 12) + 6:17, ]
df <- cbind(rep(wells, each = 12), df)
names(df) <- c("well", "subPos", time)

#summarize the data
df_summary <- df %>%
  gather(key = "time", value = "RFU", -well, -subPos) %>%
  mutate(RFU = as.numeric(RFU),
         time = as.numeric(time),
         col = str_sub(well, start = 2),
         row = str_sub(well, start = 1, end = 1)) %>%
  group_by(well, time, col, row) %>%
  summarise(avgRFU = mean(RFU), maxRFU = max(RFU), minRFU = min(RFU)) %>%
  ungroup() %>%
  mutate(col = as.numeric(col))


# assign treatments
df_summary <- df_summary %>%
  mutate(Age = ifelse(col <= 6, "8 dpp", "16 dpp"),
         Treatment = ifelse(row %in% c("G", "H"), "Control", "Inoculated")) %>%
  mutate(Age = fct_relevel(Age, "8 dpp"))

# filter out H1
df_summary <- df_summary %>%
  filter(!well %in% c("H1")) %>%
  na.omit()
```

Plot fluorecense summarized over time:

```r
#plot summarised data
df_summary %>%
  ggplot(aes(x = time / 3600, y = maxRFU)) +
  stat_summary(
    data = filter(df_summary, Treatment == "Inoculated"),
    fun.data = mean_se,
    geom = "errorbar",
    aes(x = time / 3600, y = maxRFU, group = Age),
    alpha = 0.2
  ) +
  stat_summary(
    data = filter(df_summary, Treatment == "Control"),
    fun.data = mean_se,
    geom = "errorbar",
    aes(x = time / 3600, y = maxRFU, group = Age),
```

```
    alpha = 0.2
  ) +

  stat_summary(aes(color = Age, linetype = Treatment), fun.y = "mean", geom = "line", size = 1) +
  labs(x = "Hours post-inoculation", y = "Relative fluorescence") +
  cowplot::theme_cowplot(font_size = 14) +
    theme(legend.title = element_blank(),
          legend.position = c(0.1, 0.7),
          legend.direction = "vertical")
```

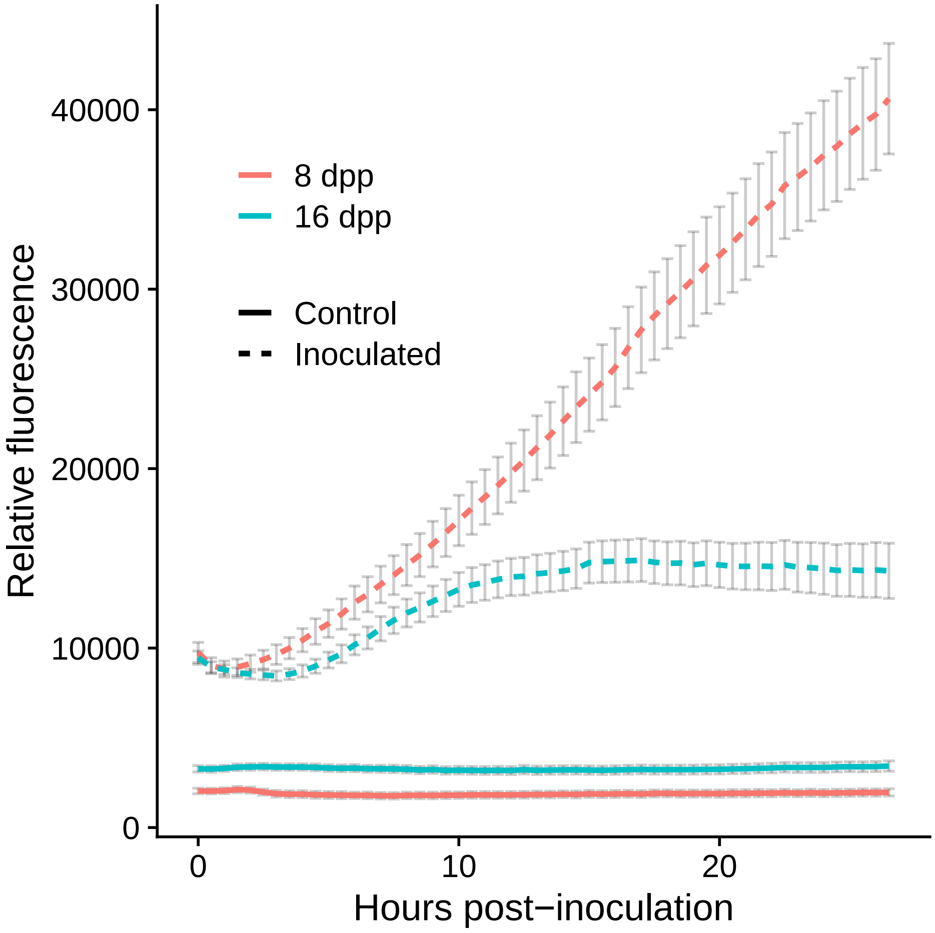


Figure 1:

```
mic <- cowplot::ggdraw() + cowplot::draw_image(image = "fluMic2.png")
poin <- cowplot::ggdraw() + cowplot::draw_image(image = "poinsett.png")

cowplot::plot_grid(drPlot, poin, mic,
                   ncol = 1,
                   labels = "AUTO",
                   rel_heights = c(3, 1.5, 1.8))
```

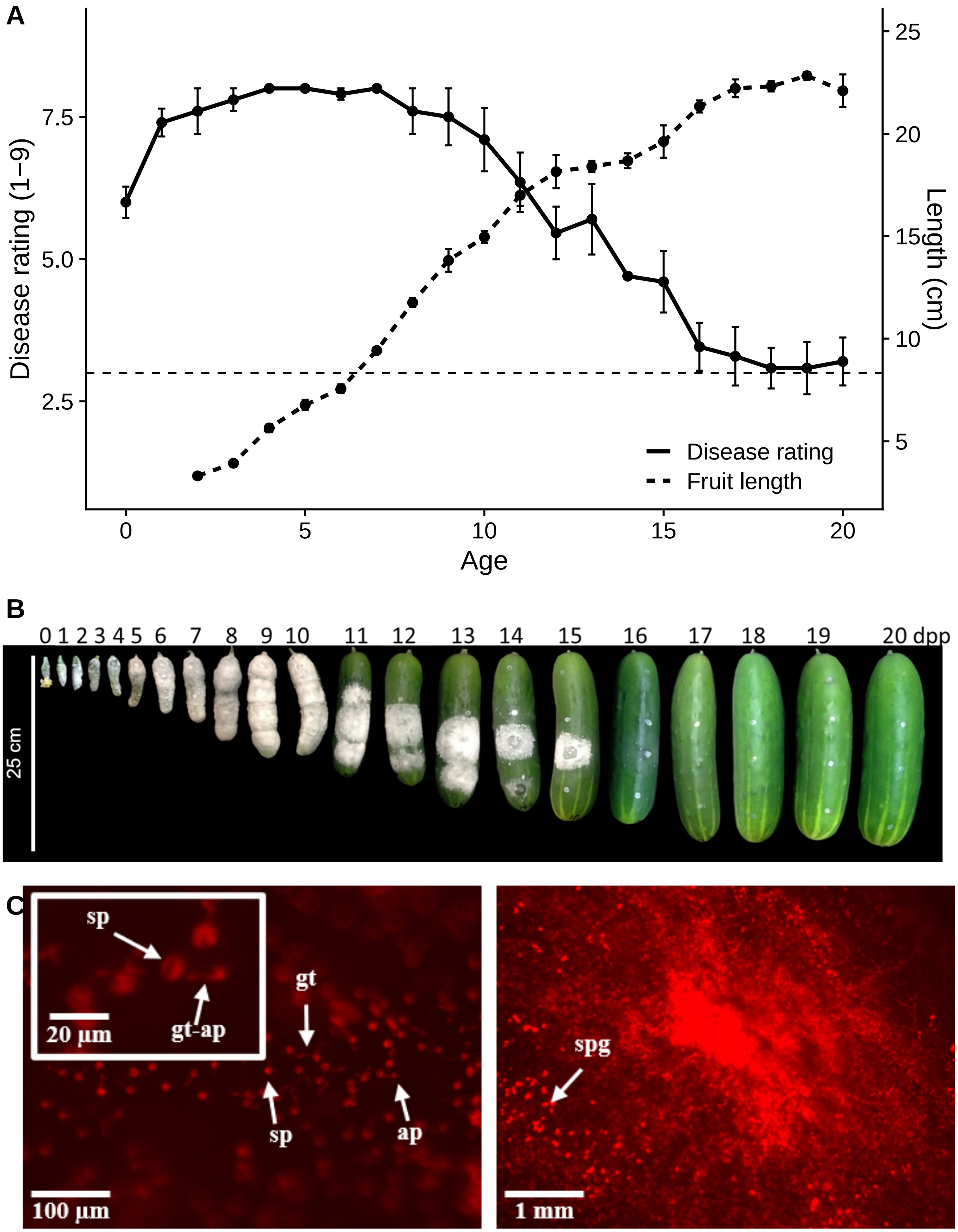


```r
pdf(file = "fig1.pdf", width = 7, 9)
cowplot::plot_grid(drPlot, poin, mic,
                   ncol = 1,
```

```
                    labels = "AUTO",
                    rel_heights = c(3, 1.5, 1.8))
dev.off()
```

```
## pdf
##   2
```

### Transcriptome Experiment 1

#### Salmon mapping

```
mkdir infectionRNAseq
mkdir genome
mkdir cl9930-v2

# Download the genome files
wget ftp://cucurbitgenomics.org/pub/cucurbit/genome/cucumber/Chinese_long/v2/*
```

Make a cdna fasta file for the cuke transcriptome

```
module load cufflinks/2.2.1

gzip -dc "~/infectionRNAseq/genome/cl9930-v2/cucumber_ChineseLong_v2.gff3.gz" > \
cucumber_ChineseLong_v2.gff3

gzip -dc "~/infectionRNAseq/genome/cl9930-v2/cucumber_ChineseLong_v2_genome.fa.gz" > \
cucumber_ChineseLong_v2_genome.fa

gffread -g cucumber_ChineseLong_v2_genome.fa \
-w cucumber_ChineseLong_v2_cdna.fa \
cucumber_ChineseLong_v2.gff3
```

```
#!/bin/bash -login
#PBS -l walltime=02:00:00
#PBS -l nodes=1:ppn=17,feature=gbe,mem=8gb
#PBS -t 1-30

#set wd
cd ${PBS_O_WORKDIR}
mkdir FastQC/
mkdir FastQC/Cleaned

# get the filename in the files.txt file at position PBS_ARRAY_INDEX
SAMPLENAME=`ls *001.fastq.gz | head -n +${PBS_ARRAYID} | tail -n 1`

module load Trimmomatic

java -jar $TRIM/trimmomatic SE \
-threads 16 \
-phred33 ${SAMPLENAME} ${SAMPLENAME%.fast*}_trimmed.fastq.gz \
ILLUMINACLIP:$ADAPTERS/TruSeq-SE.fa:2:30:10 \
LEADING:3 \
TRAILING:3 \
SLIDINGWINDOW:4:15 \
```

```
MINLEN:35

module load FastQC/0.11.3
fastqc -f fastq -noextract ${SAMPLENAME%.fast*}_trimmed.fastq.gz -o FastQC/Cleaned
```

Building a salmon index file

```
salmon index -t cucumber_ChineseLong_v2_cdna.fa -i cucumber_ChineseLong_v2_salmon_index
```

Run Salmon on the samples:

```
for fn in `ls rawReads/*trimmed.fastq.gz`;
do
nodir=${fn#rawReads/}
echo "Processing sample ${fn%_S*}"
salmon quant -i genome/cl9930-v2/cucumber_ChineseLong_v2_salmon_index -l A \
         -r ${fn} \
         -p 8 -o quants/${nodir%_S*}_quant
done
```

#### Differntial expression with DESeq2

load GO term functions and files.

```
source("topGO functions.R")

CukeGO <- read.csv(file = "CukeGO.csv", header = TRUE, stringsAsFactors = FALSE)

#prepare as named list for use with topGO
GOList<-setNames(nm = CukeGO$X, strsplit(CukeGO$GOTerm, "; "))
```

#### Differential expression analysis

Import using tximport

```
samples <- read.csv(file = "samples_exp1.csv", row.names = 1)
files <- file.path("quants", samples$directory, "quant.sf")
names(files) <- row.names(samples)

samples$timepoint <- as.factor(paste0("T", samples$timepoint))
samples$age <- relevel(samples$age, "8dpp")
samples$timepoint <- fct_relevel(samples$timepoint, "T0", "T4", "T24", "T48")

samples$condition <-
    fct_relevel(
    paste0(samples$age, "_", samples$timepoint),
    "8dpp_T0",
    "8dpp_T4",
    "8dpp_T24",
    "8dpp_T48",
    "16dpp_T0",
    "16dpp_T4",
    "16dpp_T24",
    "16dpp_T48"
    )
```

Table 1: The experimental design

| age | timepoint | Reps |
|---|---|---|
| 8dpp | T0 | 3 |
| 8dpp | T4 | 3 |
| 8dpp | T24 | 3 |
| 8dpp | T48 | 3 |
| 16dpp | T0 | 3 |
| 16dpp | T4 | 3 |
| 16dpp | T24 | 3 |
| 16dpp | T48 | 3 |

```
samples$name <- row.names(samples)

# make tx2gene file
tx2gene <- read.table(file = files[1], sep = "\t", header = TRUE)
tx2gene <- tx2gene[1]
tx2gene$geneid <- str_replace(tx2gene$Name, pattern = "\\..*", replacement = "")

files <- file.path("quants", samples$directory, "quant.sf")
names(files) <- row.names(samples)

txiAll <- tximport(files, type = "salmon", tx2gene = tx2gene, dropInfReps = TRUE)

ddsTxi <- DESeqDataSetFromTximport(txiAll, colData = samples, design = ~ condition)
ddsTxi_filt <- ddsTxi[rowSums(counts(ddsTxi)) >= 25, ]

dds_exp1 <- DESeq(ddsTxi_filt)
```

Reads mapping summary

```
# Import JSON files for sample read mapping results

jsons <- file.path("quants", samples$directory, "aux_info", "meta_info.json")
samples$totalReads <- sapply(jsons, FUN = function(x) {
    fromJSON(file = x)$num_processed
})

samples$Mapped <- sapply(jsons, FUN = function(x) {
    fromJSON(file = x)$num_mapped
})
```

```
expDesign <- samples %>%
    group_by(age, timepoint) %>% summarise(Reps = n())
knitr::kable(as.data.frame(expDesign), caption = "The experimental design") %>%
    kable_styling(bootstrap_options = "striped", full_width = FALSE)
```

```
samples %>%
    mutate(sampleName = rownames(.),
    Unmapped = totalReads - Mapped) %>%
    arrange(age, timepoint) %>%
    mutate(sampleName = fct_inorder(sampleName)) %>%
    gather(
```

```r
    key = "Reads",
    value = "Count",
    -directory,
    -age,
    -name,
    -timepoint,
    -condition,
    -sampleName,
    -totalReads
    ) %>%
    mutate(Percent = round(Count / totalReads * 100, digits = 1)) %>%
    group_by(Reads) %>%
    summarize(mean(Count), mean(Percent))
```

```
## # A tibble: 2 x 3
##   Reads    `mean(Count)` `mean(Percent)`
##   <chr>            <dbl>           <dbl>
## 1 Mapped       19649607.            82.3
## 2 Unmapped      4179976.            17.7
```

```r
samples %>%
    mutate(sampleName = rownames(.),
    Unmapped = totalReads - Mapped) %>%
    arrange(age, timepoint) %>%
    mutate(sampleName = fct_inorder(sampleName)) %>%
    gather(
    key = "Reads",
    value = "Count",
    -directory,
    -age,
    -name,
    -timepoint,
    -condition,
    -sampleName,
    -totalReads
    ) %>%
    ggplot(aes(
        x = sampleName,
        y = Count,
        fill = Reads,
        label = paste0(round(Count / totalReads * 100, digits = 1), "%")
        )) +
        geom_bar(stat = "identity", position = "stack") +
        geom_text(
        size = 2,
        stat = "identity",
        position = "stack",
        hjust = 0.5
        ) +
        theme(axis.text.x = element_text(angle = 90, vjust = 0.5))
```

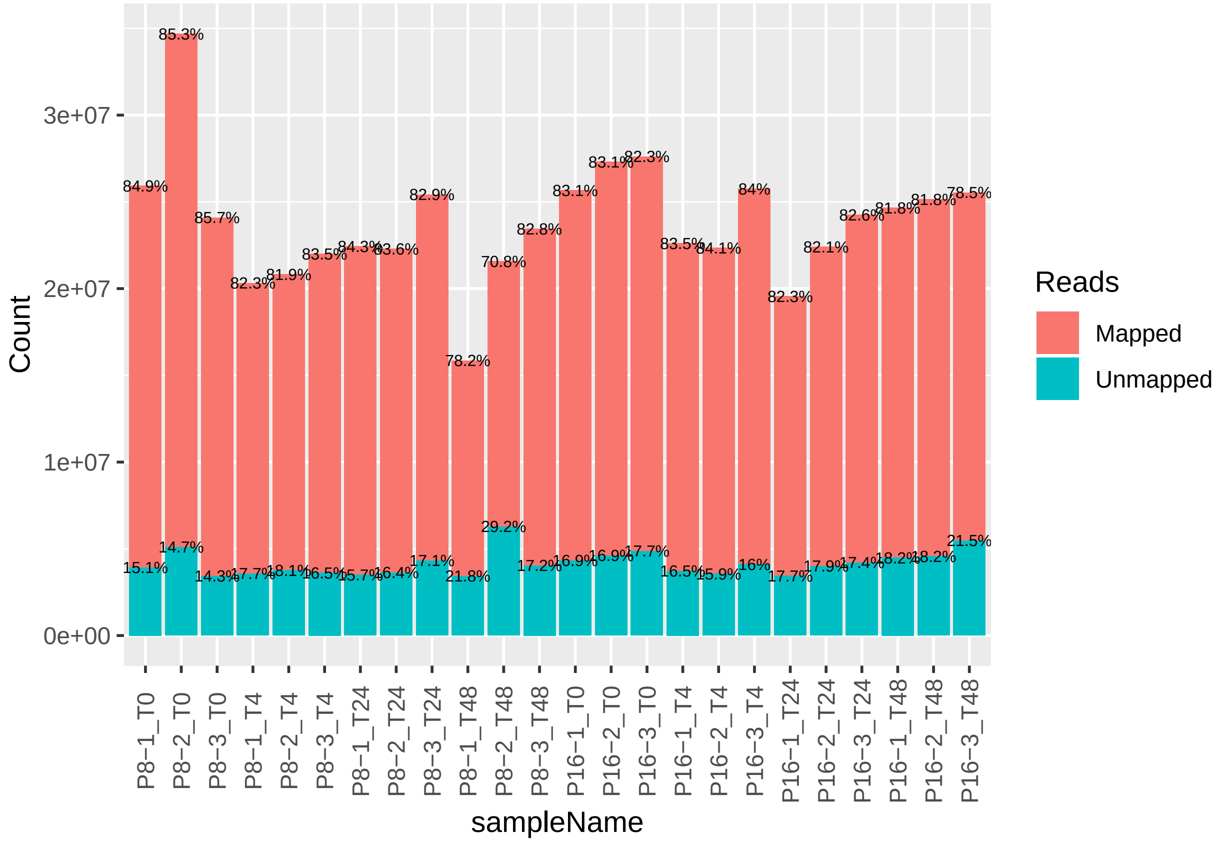


Principal component analysis

```
rld <- rlog(dds_exp1, blind = TRUE)

pcaData <- plotPCA(rld, returnData=TRUE, ntop = 500)

percentVar <- round(100 * attr(pcaData, "percentVar"))

pcaData <- pcaData %>%
    mutate(Age = dds_exp1$age,
           Timepoint = dds_exp1$timepoint)

pcaPlot1 <- ggplot(pcaData, aes(PC1, PC2, color = Timepoint, shape = Age)) +
    geom_point(size=5) +
    xlab(paste0("PC1: ",percentVar[1],"% variance")) +
    ylab(paste0("PC2: ",percentVar[2],"% variance")) +
    cowplot::theme_cowplot() +
    theme(strip.text =          element_text(
        colour = "grey10",
        size = rel(0.8),
        margin = margin(0.8 * 7, 0.8 * 7, 0.8 * 7, 0.8 * 7)
        ),
        legend.position="bottom") +
    scale_color_viridis_d(direction = -1)
   #ggrepel::geom_text_repel(aes(label = name)) # for samples names if needed

pcaPlot1
```

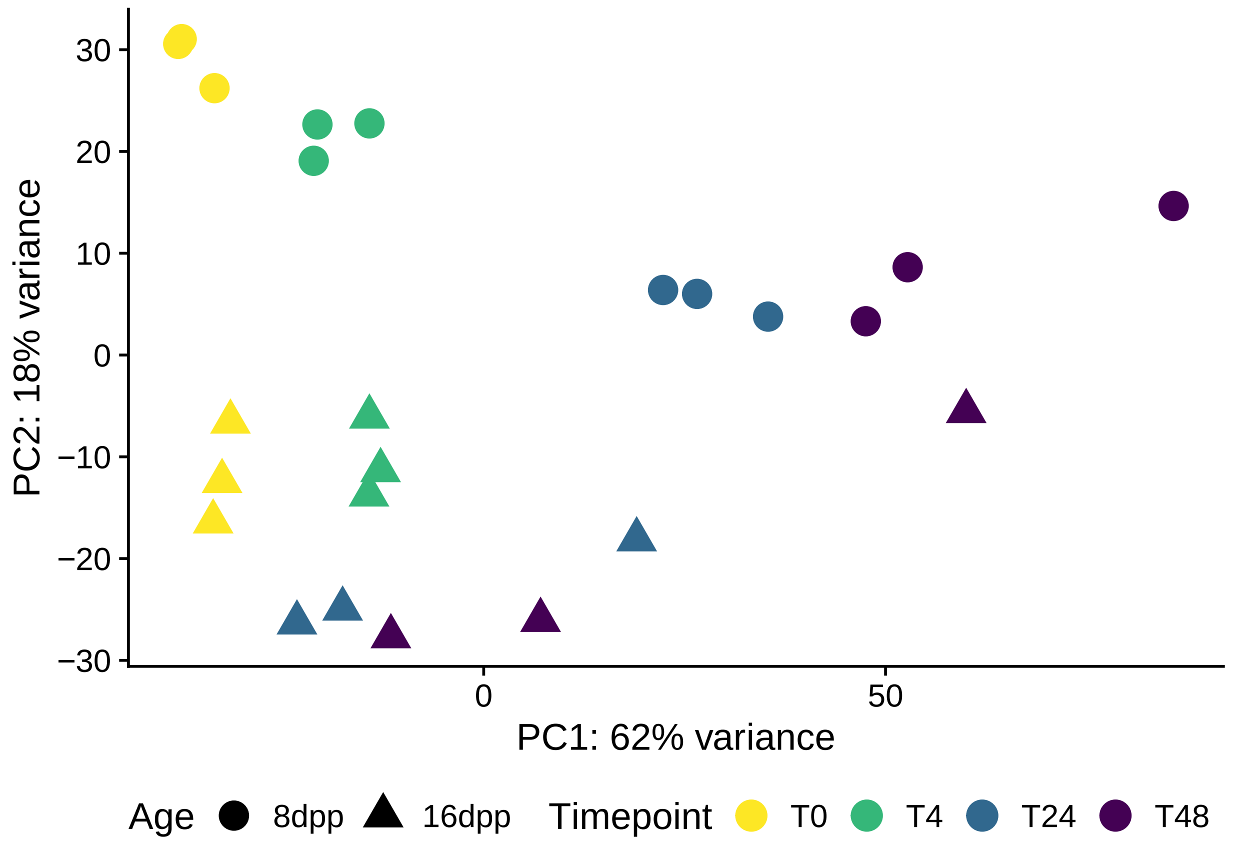


Get between sample within treatment Pearson's correlations and plot

```
rld_df <- as.data.frame(assay(rld))
cor(rld_df[, dds_exp1$age == "16dpp" & dds_exp1$timepoint == "T0"], method="pearson")
```

```
##            P16-1_T0  P16-2_T0  P16-3_T0
## P16-1_T0 1.0000000 0.9962891 0.9958321
## P16-2_T0 0.9962891 1.0000000 0.9938945
## P16-3_T0 0.9958321 0.9938945 1.0000000
```

```
cor(rld_df[, dds_exp1$age == "16dpp" & dds_exp1$timepoint == "T4"], method="pearson")
```

```
##            P16-1_T4  P16-2_T4  P16-3_T4
## P16-1_T4 1.0000000 0.9970227 0.9963668
## P16-2_T4 0.9970227 1.0000000 0.9968560
## P16-3_T4 0.9963668 0.9968560 1.0000000
```

```
cor(rld_df[, dds_exp1$age == "16dpp" & dds_exp1$timepoint == "T24"], method="pearson")
```

```
##             P16-1_T24 P16-2_T24 P16-3_T24
## P16-1_T24 1.0000000 0.9935373 0.9768389
## P16-2_T24 0.9935373 1.0000000 0.9753957
## P16-3_T24 0.9768389 0.9753957 1.0000000
```

```
cor(rld_df[, dds_exp1$age == "16dpp" & dds_exp1$timepoint == "T48"], method="pearson")
```

```
##             P16-1_T48 P16-2_T48 P16-3_T48
## P16-1_T48 1.0000000 0.9933915 0.9750264
```

```
## P16-2_T48 0.9933915 1.0000000 0.9633022
## P16-3_T48 0.9750264 0.9633022 1.0000000
```

```r
cor(rld_df[, dds_exp1$age == "8dpp" & dds_exp1$timepoint == "T0"], method="pearson")
```

```
##           P8-1_T0   P8-2_T0   P8-3_T0
## P8-1_T0 1.0000000 0.9943148 0.9922322
## P8-2_T0 0.9943148 1.0000000 0.9967608
## P8-3_T0 0.9922322 0.9967608 1.0000000
```

```r
cor(rld_df[, dds_exp1$age == "8dpp" & dds_exp1$timepoint == "T4"], method="pearson")
```

```
##           P8-1_T4   P8-2_T4   P8-3_T4
## P8-1_T4 1.0000000 0.9933884 0.9943525
## P8-2_T4 0.9933884 1.0000000 0.9964352
## P8-3_T4 0.9943525 0.9964352 1.0000000
```

```r
cor(rld_df[, dds_exp1$age == "8dpp" & dds_exp1$timepoint == "T24"], method="pearson")
```

```
##            P8-1_T24  P8-2_T24  P8-3_T24
## P8-1_T24 1.0000000 0.9947419 0.9934408
## P8-2_T24 0.9947419 1.0000000 0.9965143
## P8-3_T24 0.9934408 0.9965143 1.0000000
```

```r
cor(rld_df[, dds_exp1$age == "8dpp" & dds_exp1$timepoint == "T48"], method="pearson")
```

```
##            P8-1_T48  P8-2_T48  P8-3_T48
## P8-1_T48 1.0000000 0.9724116 0.9888864
## P8-2_T48 0.9724116 1.0000000 0.9690032
## P8-3_T48 0.9888864 0.9690032 1.0000000
```

```r
cor<-as.matrix(cor(rld_df, method="pearson"))
rownames(cor) <- colnames(cor) <- with(colData(dds_exp1), paste(age, timepoint, sep="-"))
pheatmap::pheatmap(cor)
```

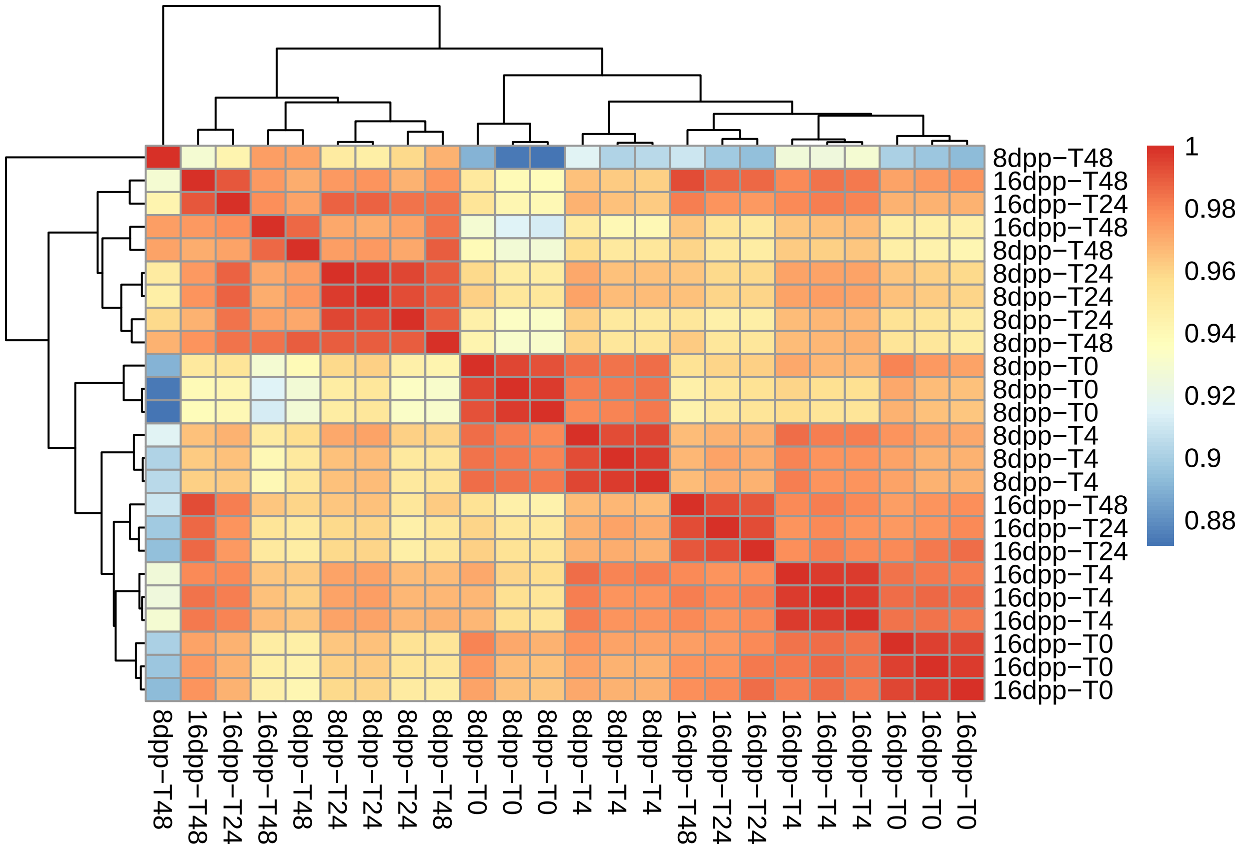


Plot Euclidean distances between samples as well

```
distsRL <- dist(t(assay(rld)))
mat <- as.matrix(distsRL)
rownames(mat) <-  colData(rld)$condition
colnames(mat) <-  colData(rld)$sampleNO

#hmcol <- colorRampPalette(brewer.pal(9, "Blues"))(255)
pheatmap(mat)
```

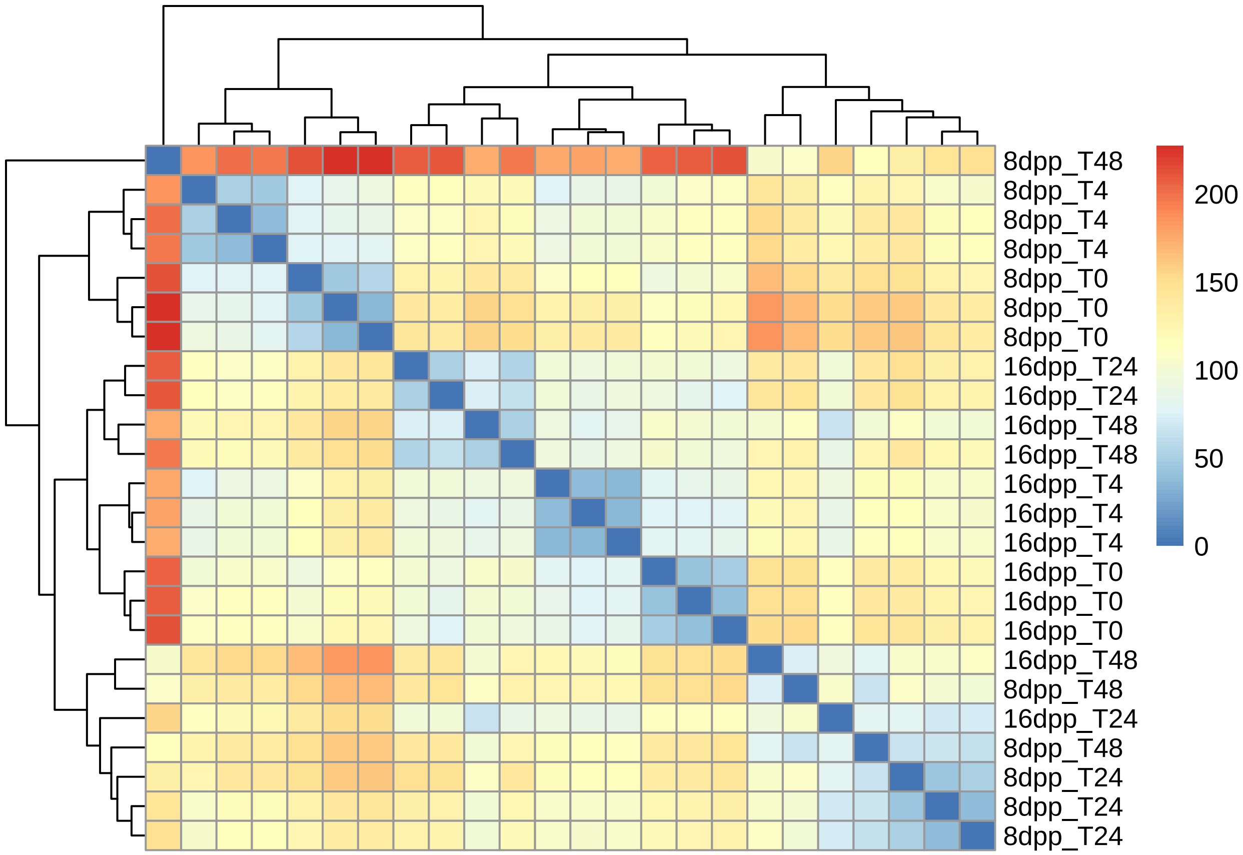


Comparison of 8 dpp and 16 dpp T0 - uninoculated fruit:

```
DE_AgeT0 <- lfcShrink(dds_exp1, "condition_16dpp_T0_vs_8dpp_T0")

summary(DE_AgeT0, 0.05)
```

```
##
## out of 19667 with nonzero total read count
## adjusted p-value < 0.05
## LFC > 0 (up)       : 3413, 17%
## LFC < 0 (down)     : 4391, 22%
## outliers [1]       : 18, 0.092%
## low counts [2]     : 0, 0%
## (mean count < 1)
## [1] see 'cooksCutoff' argument of ?results
## [2] see 'independentFiltering' argument of ?results
```

```
DE_AgeT0 %>%
    as.data.frame() %>%
    mutate(geneid = rownames(.)) %>%
    dplyr::filter(padj < 0.05 & abs(log2FoldChange) >= 1) %>%
    mutate(Direction = fct_relevel(ifelse(log2FoldChange > 0, "Up", "Down"), "Up")) %>%
    group_by(Direction) %>%
    do({tgd <- runTopGoAnalysis(DEgeneSet = .$geneid,
                                dds = dds_exp1,
                                GOdb = GOList,
                                onts = "BP",
```

```
                              nodeSize = 100)
     exportGOtable(tgd, "Fisher.weight01", n = 15) %>%
         mutate(Fisher.weight01 = as.numeric(Fisher.weight01))
         }
     )
```

```
## # A tibble: 30 x 7
## # Groups:   Direction [2]
##    Direction GO.ID   Term         Annotated Significant Expected Fisher.weight01
##    <fct>     <chr>   <chr>            <int>       <int>    <dbl>           <dbl>
##  1 Up        GO:001... secondary m...      251          26    11.8         0.000075
##  2 Up        GO:000... peptide met...      643          16    30.2         0.00016
##  3 Up        GO:004... multi-organ...      136          17     6.39        0.00044
##  4 Up        GO:000... sulfur comp...      256          24    12.0         0.00057
##  5 Up        GO:005... cofactor me...      486          30    22.8         0.00072
##  6 Up        GO:000... metabolic p...     9213         451   433.          0.00109
##  7 Up        GO:001... response to...      294          28    13.8         0.00189
##  8 Up        GO:004... regulation ...      181          17     8.5         0.00515
##  9 Up        GO:004... root develo...      376          24    17.7         0.00518
## 10 Up        GO:005... oxidation-r...     1249          78    58.7         0.00569
## # ... with 20 more rows
```

Extract results from dds. These are comparisons (contrasts) of consequtive timepoints within each age. Log2 Fold Changes are shrunken using the “normal” method.

```
comparsion <- 1:3
timepoints <- c("T0", "T4", "T24", "T48")


conditions <- paste0("8dpp_", timepoints)
timeContrasts8 <- bind_rows(lapply(comparsion, function(x) {
    lfc <- lfcShrink(dds_exp1, contrast = c("condition", conditions[x + 1], conditions[x]))
    as.data.frame(lfc) %>%
    mutate(
    geneid = rownames(.),
    Contrast = paste0(timepoints[x + 1]," vs ", timepoints[x])
    )
}))

conditions <- paste0("16dpp_", timepoints)
timeContrasts16 <- bind_rows(lapply(comparsion, function(x) {
    lfc <- lfcShrink(dds_exp1, contrast = c("condition", conditions[x + 1], conditions[x]))
    as.data.frame(lfc) %>%
    mutate(
    geneid = rownames(.),
    Contrast = paste0(timepoints[x + 1]," vs ", timepoints[x])
    )
}))

timeContrasts <- bind_rows("8dpp" = timeContrasts8, "16dpp" = timeContrasts16, .id = "Age") %>%
    mutate(Contrast = fct_inorder(Contrast),
           Age = fct_relevel(Age, "8dpp")
           )
```

Plot a summary of the differentially expressed genes in each contrast.

```r
sumPlot1 <- timeContrasts %>%
    filter(padj < 0.05 & abs(log2FoldChange) >= 1) %>%
    mutate(Direction = fct_relevel(ifelse(log2FoldChange > 0, "Up", "Down"), "Up")) %>%
    group_by(Contrast, Age, Direction) %>%
    ggplot(aes(x = Direction, fill = Age)) +
    geom_bar(position = "dodge") +
    geom_text(stat = 'count', aes(label = ..count..), vjust = -0.5, position = position_dodge(0.90)) +
    # facet_wrap(~ Contrast) +
    scale_y_continuous(expand = expand_scale(mult = c(0.05, 0.1)), name = "Number of DEGs") +
    cowplot::theme_cowplot(font_size = 14) +
    theme(strip.text = element_text(
        colour = "grey10",
        size = rel(0.8),
        margin = margin(0.5 * 7, 0.5 * 7, 0.5 * 7, 0.5 * 7)
        ),
        legend.position = c(0.825, 0.825),
        axis.line=element_line()) +
    lemon::facet_rep_wrap(~ Contrast) +
     cowplot::panel_border()

sumPlot1
```

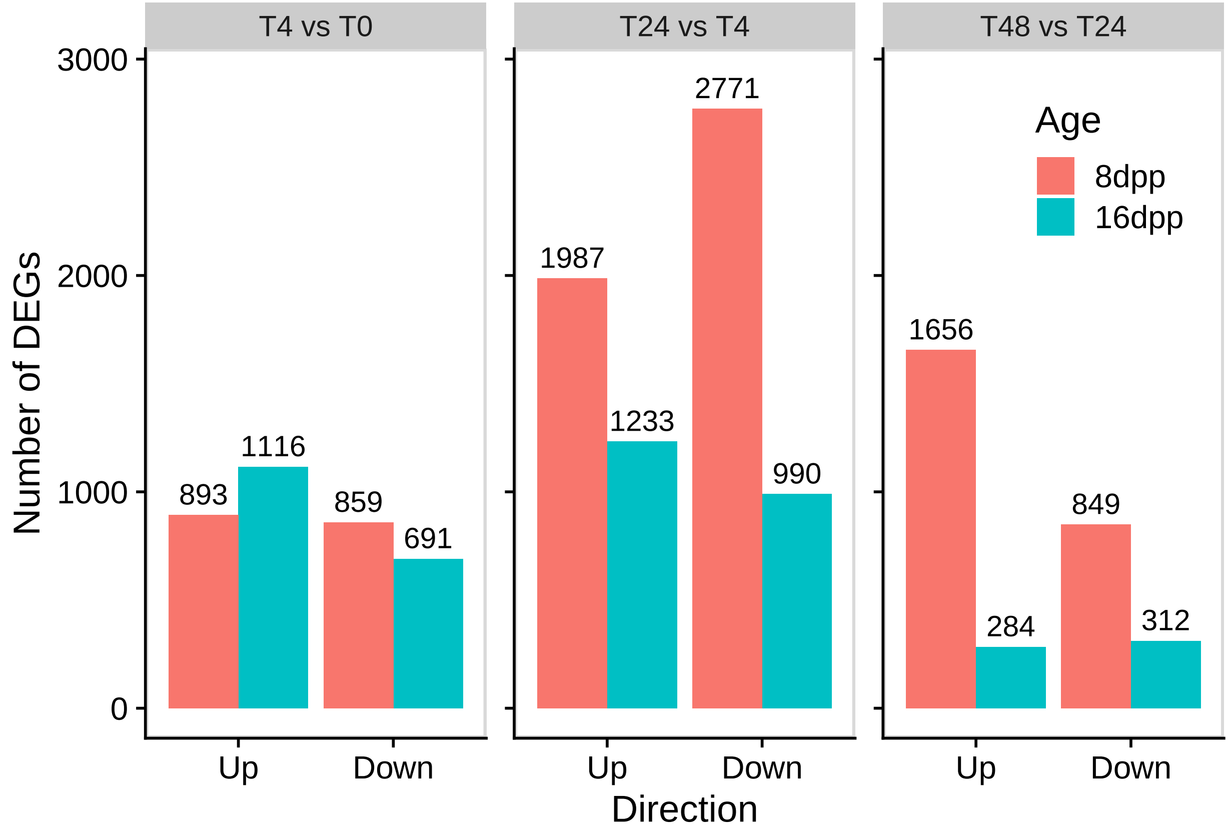


Extract Gene Ontology terms for each contrast:

```r
allGOres <- timeContrasts %>%
    filter(padj < 0.05 & abs(log2FoldChange) >= 1) %>%
    mutate(Direction = fct_relevel(ifelse(log2FoldChange > 0, "Up", "Down"), "Up")) %>%
```

```r
    group_by(Contrast, Age, Direction) %>%
    # filter(Age == "16dpp", Contrast == "T4 vs T0", Direction == "Up") %>%
    do({tgd <- runTopGoAnalysis(DEgeneSet = .$geneid,
                                dds = dds_exp1,
                                GOdb = GOList,
                                onts = "BP",
                                nodeSize = 100)
       exportGOtable(tgd, "Fisher.weight01", n = 500) %>%
           mutate(Fisher.weight01 = as.numeric(Fisher.weight01))
           }
       )
```

Tables of results

```r
for (i in unique(allGOres$Contrast)) {
print(
kable(allGOres %>%
          filter(Contrast == i ) %>%
          top_n(n = 10, wt = dplyr::desc(Fisher.weight01)), digits = 20) %>%
    kable_styling(bootstrap_options = "striped", full_width = FALSE, latex_options="scale_down") %>%
    kableExtra::kable_styling(latex_options = c("HOLD_position"))
)
}
```

| Contrast | Age | Direction | GO.ID | Term | Annotated | Significant | Expected | Fisher.weight01 |
|---|---|---|---|---|---|---|---|---|
| T4 vs T0 | 8dpp | Up | GO:0006979 | response to oxidative stress | 380 | 66 | 17.47 | 2.10e-19 |
| T4 vs T0 | 8dpp | Up | GO:0009611 | response to wounding | 259 | 43 | 11.91 | 1.90e-13 |
| T4 vs T0 | 8dpp | Up | GO:0006952 | defense response | 1018 | 118 | 46.81 | 2.50e-13 |
| T4 vs T0 | 8dpp | Up | GO:0009699 | phenylpropanoid biosynthetic process | 133 | 29 | 6.12 | 1.70e-12 |
| T4 vs T0 | 8dpp | Up | GO:0080167 | response to karrikin | 141 | 29 | 6.48 | 7.80e-12 |
| T4 vs T0 | 8dpp | Up | GO:0042737 | drug catabolic process | 151 | 30 | 6.94 | 8.60e-12 |
| T4 vs T0 | 8dpp | Up | GO:1901565 | organonitrogen compound catabolic proces... | 489 | 41 | 22.48 | 2.50e-11 |
| T4 vs T0 | 8dpp | Up | GO:0050832 | defense response to fungus | 239 | 36 | 10.99 | 3.10e-10 |
| T4 vs T0 | 8dpp | Up | GO:0010200 | response to chitin | 134 | 26 | 6.16 | 3.70e-10 |
| T4 vs T0 | 8dpp | Up | GO:0019439 | aromatic compound catabolic process | 215 | 23 | 9.89 | 1.20e-07 |
| T4 vs T0 | 8dpp | Down | GO:0071555 | cell wall organization | 323 | 48 | 14.75 | 1.40e-09 |
| T4 vs T0 | 8dpp | Down | GO:0007017 | microtubule-based process | 137 | 20 | 6.26 | 4.00e-06 |
| T4 vs T0 | 8dpp | Down | GO:0006073 | cellular glucan metabolic process | 171 | 21 | 7.81 | 3.50e-05 |
| T4 vs T0 | 8dpp | Down | GO:0016114 | terpenoid biosynthetic process | 110 | 16 | 5.02 | 3.80e-05 |
| T4 vs T0 | 8dpp | Down | GO:0009664 | plant-type cell wall organization | 101 | 15 | 4.61 | 5.20e-05 |
| T4 vs T0 | 8dpp | Down | GO:0000272 | polysaccharide catabolic process | 130 | 17 | 5.94 | 8.80e-05 |
| T4 vs T0 | 8dpp | Down | GO:0051707 | response to other organism | 839 | 40 | 38.31 | 3.80e-04 |
| T4 vs T0 | 8dpp | Down | GO:0009651 | response to salt stress | 524 | 40 | 23.93 | 1.04e-03 |
| T4 vs T0 | 8dpp | Down | GO:0009826 | unidimensional cell growth | 234 | 23 | 10.69 | 3.11e-03 |
| T4 vs T0 | 8dpp | Down | GO:0007169 | transmembrane receptor protein tyrosine ... | 135 | 14 | 6.16 | 3.49e-03 |
| T4 vs T0 | 16dpp | Up | GO:0009611 | response to wounding | 259 | 61 | 15.32 | 0.00e+00 |
| T4 vs T0 | 16dpp | Up | GO:0006979 | response to oxidative stress | 380 | 67 | 22.47 | 7.80e-17 |
| T4 vs T0 | 16dpp | Up | GO:0080167 | response to karrikin | 141 | 37 | 8.34 | 6.50e-15 |
| T4 vs T0 | 16dpp | Up | GO:0042737 | drug catabolic process | 151 | 38 | 8.93 | 1.20e-14 |
| T4 vs T0 | 16dpp | Up | GO:0010200 | response to chitin | 134 | 35 | 7.92 | 4.10e-14 |
| T4 vs T0 | 16dpp | Up | GO:0009699 | phenylpropanoid biosynthetic process | 133 | 33 | 7.86 | 1.10e-12 |
| T4 vs T0 | 16dpp | Up | GO:1901565 | organonitrogen compound catabolic proces... | 489 | 53 | 28.92 | 7.60e-12 |
| T4 vs T0 | 16dpp | Up | GO:1901605 | alpha-amino acid metabolic process | 276 | 40 | 16.32 | 1.40e-10 |
| T4 vs T0 | 16dpp | Up | GO:0046395 | carboxylic acid catabolic process | 131 | 29 | 7.75 | 4.70e-10 |
| T4 vs T0 | 16dpp | Up | GO:0050832 | defense response to fungus | 239 | 40 | 14.13 | 2.10e-09 |
| T4 vs T0 | 16dpp | Down | GO:0009734 | auxin-activated signaling pathway | 150 | 18 | 5.53 | 1.10e-05 |
| T4 vs T0 | 16dpp | Down | GO:0006355 | regulation of transcription, DNA-templat... | 1745 | 97 | 64.38 | 3.90e-05 |
| T4 vs T0 | 16dpp | Down | GO:0071555 | cell wall organization | 323 | 26 | 11.92 | 7.40e-05 |
| T4 vs T0 | 16dpp | Down | GO:0009699 | phenylpropanoid biosynthetic process | 133 | 15 | 4.91 | 1.20e-04 |
| T4 vs T0 | 16dpp | Down | GO:0009059 | macromolecule biosynthetic process | 2827 | 126 | 104.30 | 1.18e-03 |
| T4 vs T0 | 16dpp | Down | GO:0008152 | metabolic process | 9213 | 350 | 339.92 | 1.39e-03 |
| T4 vs T0 | 16dpp | Down | GO:0007169 | transmembrane receptor protein tyrosine ... | 135 | 13 | 4.98 | 1.47e-03 |
| T4 vs T0 | 16dpp | Down | GO:0042445 | hormone metabolic process | 261 | 14 | 9.63 | 1.92e-03 |
| T4 vs T0 | 16dpp | Down | GO:0007275 | multicellular organism development | 2197 | 101 | 81.06 | 2.11e-03 |
| T4 vs T0 | 16dpp | Down | GO:0080167 | response to karrikin | 141 | 13 | 5.20 | 2.17e-03 |

| Contrast | Age | Direction | GO.ID | Term | Annotated | Significant | Expected | Fisher.weight01 |
|---|---|---|---|---|---|---|---|---|
| T24 vs T4 | 8dpp | Up | GO:0006952 | defense response | 1018 | 172 | 103.56 | 2.80e-11 |
| T24 vs T4 | 8dpp | Up | GO:0006260 | DNA replication | 119 | 36 | 12.11 | 1.10e-09 |
| T24 vs T4 | 8dpp | Up | GO:0006412 | translation | 564 | 94 | 57.38 | 1.30e-09 |
| T24 vs T4 | 8dpp | Up | GO:0046686 | response to cadmium ion | 335 | 66 | 34.08 | 9.90e-08 |
| T24 vs T4 | 8dpp | Up | GO:0009751 | response to salicylic acid | 145 | 33 | 14.75 | 7.00e-06 |
| T24 vs T4 | 8dpp | Up | GO:0006468 | protein phosphorylation | 935 | 135 | 95.12 | 7.70e-05 |
| T24 vs T4 | 8dpp | Up | GO:0009651 | response to salt stress | 524 | 81 | 53.31 | 8.10e-05 |
| T24 vs T4 | 8dpp | Up | GO:0042254 | ribosome biogenesis | 273 | 47 | 27.77 | 1.60e-04 |
| T24 vs T4 | 8dpp | Up | GO:0097659 | nucleic acid-templated transcription | 1867 | 178 | 189.93 | 2.00e-04 |
| T24 vs T4 | 8dpp | Up | GO:0042742 | defense response to bacterium | 306 | 51 | 31.13 | 2.80e-04 |
| T24 vs T4 | 8dpp | Down | GO:0055114 | oxidation-reduction process | 1249 | 267 | 184.64 | 4.40e-11 |
| T24 vs T4 | 8dpp | Down | GO:0015979 | photosynthesis | 201 | 114 | 29.71 | 1.10e-10 |
| T24 vs T4 | 8dpp | Down | GO:0009416 | response to light stimulus | 640 | 140 | 94.61 | 5.40e-06 |
| T24 vs T4 | 8dpp | Down | GO:0006073 | cellular glucan metabolic process | 171 | 46 | 25.28 | 2.60e-05 |
| T24 vs T4 | 8dpp | Down | GO:0005975 | carbohydrate metabolic process | 804 | 158 | 118.86 | 3.50e-05 |
| T24 vs T4 | 8dpp | Down | GO:0009735 | response to cytokinin | 238 | 58 | 35.18 | 6.00e-05 |
| T24 vs T4 | 8dpp | Down | GO:0071702 | organic substance transport | 923 | 119 | 136.45 | 1.20e-04 |
| T24 vs T4 | 8dpp | Down | GO:0042446 | hormone biosynthetic process | 185 | 46 | 27.35 | 2.00e-04 |
| T24 vs T4 | 8dpp | Down | GO:0042546 | cell wall biogenesis | 143 | 29 | 21.14 | 3.50e-04 |
| T24 vs T4 | 8dpp | Down | GO:0007169 | transmembrane receptor protein tyrosine ... | 135 | 35 | 19.96 | 4.90e-04 |
| T24 vs T4 | 16dpp | Up | GO:0019684 | photosynthesis, light reaction | 115 | 23 | 7.30 | 7.50e-07 |
| T24 vs T4 | 16dpp | Up | GO:0055114 | oxidation-reduction process | 1249 | 119 | 79.33 | 3.90e-06 |
| T24 vs T4 | 16dpp | Up | GO:0097659 | nucleic acid-templated transcription | 1867 | 129 | 118.59 | 1.90e-05 |
| T24 vs T4 | 16dpp | Up | GO:0006325 | chromatin organization | 230 | 26 | 14.61 | 1.30e-04 |
| T24 vs T4 | 16dpp | Up | GO:0009658 | chloroplast organization | 148 | 22 | 9.40 | 1.60e-04 |
| T24 vs T4 | 16dpp | Up | GO:0051649 | establishment of localization in cell | 439 | 17 | 27.88 | 4.50e-04 |
| T24 vs T4 | 16dpp | Up | GO:0009416 | response to light stimulus | 640 | 72 | 40.65 | 7.40e-04 |
| T24 vs T4 | 16dpp | Up | GO:0080167 | response to karrikin | 141 | 19 | 8.96 | 1.50e-03 |
| T24 vs T4 | 16dpp | Up | GO:1903046 | meiotic cell cycle process | 102 | 15 | 6.48 | 1.91e-03 |
| T24 vs T4 | 16dpp | Up | GO:0016311 | dephosphorylation | 149 | 14 | 9.46 | 1.92e-03 |
| T24 vs T4 | 16dpp | Down | GO:0009611 | response to wounding | 259 | 37 | 13.81 | 4.10e-08 |
| T24 vs T4 | 16dpp | Down | GO:0009873 | ethylene-activated signaling pathway | 158 | 27 | 8.42 | 7.10e-08 |
| T24 vs T4 | 16dpp | Down | GO:0033036 | macromolecule localization | 710 | 31 | 37.85 | 1.30e-06 |
| T24 vs T4 | 16dpp | Down | GO:0006979 | response to oxidative stress | 380 | 37 | 20.26 | 1.40e-06 |
| T24 vs T4 | 16dpp | Down | GO:0050832 | defense response to fungus | 239 | 31 | 12.74 | 4.10e-06 |
| T24 vs T4 | 16dpp | Down | GO:0071702 | organic substance transport | 923 | 56 | 49.20 | 9.30e-06 |
| T24 vs T4 | 16dpp | Down | GO:0009753 | response to jasmonic acid | 205 | 26 | 10.93 | 3.60e-05 |
| T24 vs T4 | 16dpp | Down | GO:0010200 | response to chitin | 134 | 18 | 7.14 | 2.70e-04 |
| T24 vs T4 | 16dpp | Down | GO:0042737 | drug catabolic process | 151 | 19 | 8.05 | 4.30e-04 |
| T24 vs T4 | 16dpp | Down | GO:0016999 | antibiotic metabolic process | 128 | 17 | 6.82 | 4.50e-04 |

| Contrast | Age | Direction | GO.ID | Term | Annotated | Significant | Expected | Fisher.weight01 |
|---|---|---|---|---|---|---|---|---|
| T48 vs T24 | 8dpp | Up | GO:0009611 | response to wounding | 259 | 63 | 21.87 | 7.400e-15 |
| T48 vs T24 | 8dpp | Up | GO:0046395 | carboxylic acid catabolic process | 131 | 33 | 11.06 | 7.700e-09 |
| T48 vs T24 | 8dpp | Up | GO:1901565 | organonitrogen compound catabolic proces... | 489 | 65 | 41.29 | 1.000e-08 |
| T48 vs T24 | 8dpp | Up | GO:0009753 | response to jasmonic acid | 205 | 40 | 17.31 | 4.300e-07 |
| T48 vs T24 | 8dpp | Up | GO:0051707 | response to other organism | 839 | 119 | 70.85 | 5.400e-07 |
| T48 vs T24 | 8dpp | Up | GO:0010200 | response to chitin | 134 | 30 | 11.31 | 5.900e-07 |
| T48 vs T24 | 8dpp | Up | GO:0009414 | response to water deprivation | 318 | 51 | 26.85 | 6.100e-06 |
| T48 vs T24 | 8dpp | Up | GO:0031669 | cellular response to nutrient levels | 113 | 25 | 9.54 | 6.300e-06 |
| T48 vs T24 | 8dpp | Up | GO:0042594 | response to starvation | 108 | 24 | 9.12 | 8.900e-06 |
| T48 vs T24 | 8dpp | Up | GO:1901605 | alpha-amino acid metabolic process | 276 | 40 | 23.31 | 1.300e-05 |
| T48 vs T24 | 8dpp | Down | GO:0019684 | photosynthesis, light reaction | 115 | 22 | 5.32 | 1.200e-08 |
| T48 vs T24 | 8dpp | Down | GO:0009416 | response to light stimulus | 640 | 59 | 29.59 | 2.700e-07 |
| T48 vs T24 | 8dpp | Down | GO:0006633 | fatty acid biosynthetic process | 167 | 24 | 7.72 | 7.500e-07 |
| T48 vs T24 | 8dpp | Down | GO:0015979 | photosynthesis | 201 | 36 | 9.29 | 2.900e-05 |
| T48 vs T24 | 8dpp | Down | GO:0090567 | reproductive shoot system development | 408 | 24 | 18.86 | 2.000e-04 |
| T48 vs T24 | 8dpp | Down | GO:0071555 | cell wall organization | 323 | 30 | 14.93 | 6.000e-04 |
| T48 vs T24 | 8dpp | Down | GO:0007275 | multicellular organism development | 2197 | 113 | 101.57 | 6.200e-04 |
| T48 vs T24 | 8dpp | Down | GO:0009607 | response to biotic stimulus | 856 | 48 | 39.57 | 7.900e-04 |
| T48 vs T24 | 8dpp | Down | GO:0000272 | polysaccharide catabolic process | 130 | 15 | 6.01 | 9.800e-04 |
| T48 vs T24 | 8dpp | Down | GO:0033036 | macromolecule localization | 710 | 22 | 32.82 | 1.300e-03 |
| T48 vs T24 | 16dpp | Up | GO:0006950 | response to stress | 2648 | 72 | 39.15 | 1.400e-08 |
| T48 vs T24 | 16dpp | Up | GO:0009628 | response to abiotic stimulus | 1764 | 35 | 26.08 | 2.900e-07 |
| T48 vs T24 | 16dpp | Up | GO:0009664 | plant-type cell wall organization | 101 | 9 | 1.49 | 1.900e-05 |
| T48 vs T24 | 16dpp | Up | GO:0055114 | oxidation-reduction process | 1249 | 37 | 18.46 | 3.700e-05 |
| T48 vs T24 | 16dpp | Up | GO:0070887 | cellular response to chemical stimulus | 889 | 27 | 13.14 | 3.800e-05 |
| T48 vs T24 | 16dpp | Up | GO:0006633 | fatty acid biosynthetic process | 167 | 10 | 2.47 | 1.900e-04 |
| T48 vs T24 | 16dpp | Up | GO:0050790 | regulation of catalytic activity | 180 | 10 | 2.66 | 3.500e-04 |
| T48 vs T24 | 16dpp | Up | GO:0002376 | immune system process | 327 | 11 | 4.83 | 6.900e-04 |
| T48 vs T24 | 16dpp | Up | GO:0009753 | response to jasmonic acid | 205 | 10 | 3.03 | 9.700e-04 |
| T48 vs T24 | 16dpp | Up | GO:0019220 | regulation of phosphate metabolic proces... | 109 | 7 | 1.61 | 1.170e-03 |
| T48 vs T24 | 16dpp | Down | GO:0055114 | oxidation-reduction process | 1249 | 40 | 21.75 | 1.300e-04 |
| T48 vs T24 | 16dpp | Down | GO:0000160 | phosphorelay signal transduction system | 192 | 10 | 3.34 | 2.700e-04 |
| T48 vs T24 | 16dpp | Down | GO:0042446 | hormone biosynthetic process | 185 | 10 | 3.22 | 1.530e-03 |
| T48 vs T24 | 16dpp | Down | GO:0008610 | lipid biosynthetic process | 473 | 19 | 8.24 | 3.870e-03 |
| T48 vs T24 | 16dpp | Down | GO:0009408 | response to heat | 178 | 9 | 3.10 | 4.040e-03 |
| T48 vs T24 | 16dpp | Down | GO:0006325 | chromatin organization | 230 | 8 | 4.01 | 7.350e-03 |
| T48 vs T24 | 16dpp | Down | GO:0050896 | response to stimulus | 4815 | 114 | 83.85 | 8.160e-03 |
| T48 vs T24 | 16dpp | Down | GO:0048878 | chemical homeostasis | 318 | 8 | 5.54 | 8.530e-03 |
| T48 vs T24 | 16dpp | Down | GO:0007169 | transmembrane receptor protein tyrosine ... | 135 | 7 | 2.35 | 9.420e-03 |
| T48 vs T24 | 16dpp | Down | GO:0009414 | response to water deprivation | 318 | 12 | 5.54 | 1.008e-02 |

Extract the top terms in T4 vs T0 regardless of age:

```
allGOres %>%
    filter(Contrast == "T4 vs T0") %>%
    filter(Direction == "Up") %>%
    group_by(Age) %>%
    top_n(n = 10, wt = dplyr::desc(Fisher.weight01)) %>%
    pull(Term) %>% unique()
```

```
##  [1] "response to oxidative stress"
##  [2] "response to wounding"
##  [3] "defense response"
##  [4] "phenylpropanoid biosynthetic process"
##  [5] "response to karrikin"
##  [6] "drug catabolic process"
##  [7] "organonitrogen compound catabolic proces..."
##  [8] "defense response to fungus"
##  [9] "response to chitin"
## [10] "aromatic compound catabolic process"
## [11] "alpha-amino acid metabolic process"
## [12] "carboxylic acid catabolic process"
```

Plot heatmaps of GO terms over consequtive contrasts (Figure 3):

```
library(ggdendro)
source("myheatmap.R")

legend <- cowplot::get_legend(allGOres %>%
                         ungroup() %>%
                         filter(Fisher.weight01 < 0.01) %>%
                         #separate(col = Contrast, into = c("Age", "Contrast", "Direction"), sep = "_",
                         mutate(Term = factor(Term, unique(Term)),
                                Contrast = paste(Age, Contrast, sep = "_")) %>%
                         #mutate(Term = factor(Term,) %>%
                         #mutate(Age = fct_relevel(Age, "P8")) %>%
                         ggplot() +
                         geom_tile(aes(
                             x = Age,
                             y = factor(str_trunc(as.character(Term), 40, "right")),
                             fill = -(Fisher.weight01)
                         )) +
                         geom_vline(xintercept = 1.5, color = "grey92") +
                         scale_y_discrete(position = "right") +
                         scale_fill_gradient(name = "-log10(p-value)",
                                             low = "#ffeda0",
                                             high = "#f03b20",
                                             limits = c(1.30103, 25)) +
                         theme(legend.position = "bottom",
                               text = element_text(size = 10),
                               plot.margin = unit(c(0, 0, 0, 0), "cm"),
                               legend.margin = unit(c(0, 0, 0, 0), "cm")
                               )
)

### Plot all

p1 <- myheatmap2(allGOres ,"Up", trim = 35, pval = 0.01)
p2 <- myheatmap2(allGOres, "Down", trim = 35, pval = 0.01)

pall <- cowplot::ggdraw(cowplot::plot_grid(
    cowplot::plot_grid(
        p1,
        p2,
        ncol = 2,
        align = 'h',
        rel_widths = c(1, 1),
        labels = "AUTO"
    ),
    cowplot::plot_grid(
        NULL,
        legend,
        NULL,
        ncol = 3,
        rel_widths = c(0.8, 0.2, 1)
    ),
    nrow = 2,
```

```
    rel_heights = c(1, 0.05)
))

pall
```

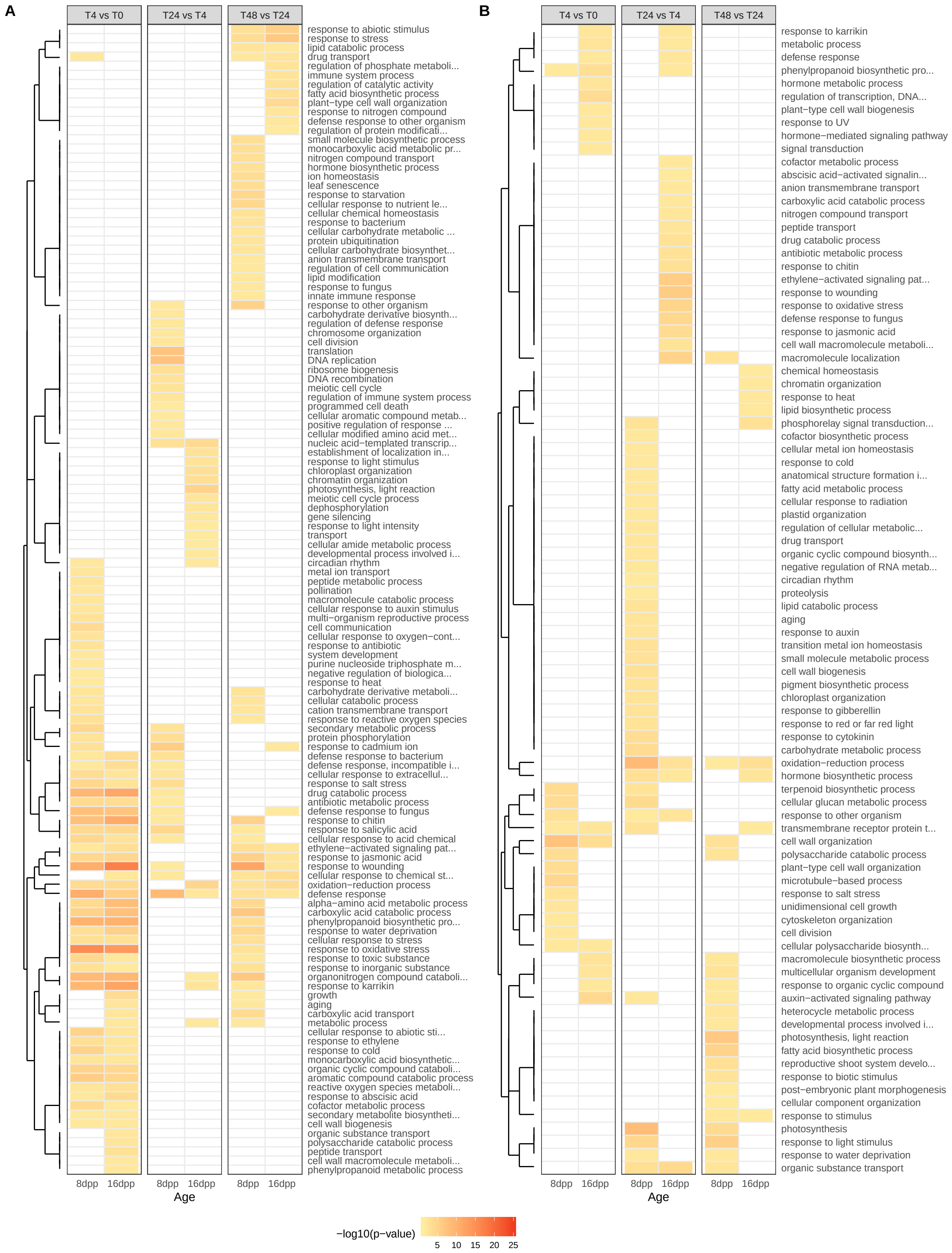
A
T4 vs T0
T24 vs T4
T48 vs T24
response to abiotic stimulus
response to stress
lipid catabolic process
drug transport
regulation of phosphate metaboli...
immune system process
regulation of catalytic activity
fatty acid biosynthetic process
plant−type cell wall organization
response to nitrogen compound
defense response to other organism
regulation of protein modificati...
small molecule biosynthetic process
monocarboxylic acid metabolic pr...
nitrogen compound transport
hormone biosynthetic process
ion homeostasis
leaf senescence
response to starvation
cellular response to nutrient le...
cellular chemical homeostasis
response to bacterium
cellular carbohydrate metabolic ...
protein ubiquitination
cellular carbohydrate biosynthet...
anion transmembrane transport
regulation of cell communication
lipid modification
response to fungus
innate immune response
response to other organism
carbohydrate derivative biosynth...
regulation of defense response
chromosome organization
cell division
translation
DNA replication
ribosome biogenesis
DNA recombination
meiotic cell cycle
regulation of immune system process
programmed cell death
cellular aromatic compound metab...
positive regulation of response ...
cellular modified amino acid met...
nucleic acid−templated transcrip...
establishment of localization in...
response to light stimulus
chloroplast organization
chromatin organization
photosynthesis, light reaction
meiotic cell cycle process
dephosphorylation
gene silencing
response to light intensity
transport
cellular amide metabolic process
developmental process involved i...
circadian rhythm
metal ion transport
peptide metabolic process
pollination
macromolecule catabolic process
cellular response to auxin stimulus
multi−organism reproductive process
cell communication
cellular response to oxygen−cont...
response to antibiotic
system development
purine nucleoside triphosphate m...
negative regulation of biologica...
response to heat
carbohydrate derivative metaboli...
cellular catabolic process
cation transmembrane transport
response to reactive oxygen species
secondary metabolic process
protein phosphorylation
response to cadmium ion
defense response to bacterium
defense response, incompatible i...
cellular response to extracellul...
response to salt stress
drug catabolic process
antibiotic metabolic process
defense response to fungus
response to chitin
response to salicylic acid
cellular response to acid chemical
ethylene−activated signaling pat...
response to jasmonic acid
response to wounding
cellular response to chemical st...
oxidation−reduction process
defense response
alpha−amino acid metabolic process
carboxylic acid catabolic process
phenylpropanoid biosynthetic pro...
response to water deprivation
cellular response to stress
response to oxidative stress
response to toxic substance
response to inorganic substance
organonitrogen compound cataboli...
response to karrikin
growth
aging
carboxylic acid transport
metabolic process
cellular response to abiotic sti...
response to ethylene
response to cold
monocarboxylic acid biosynthetic...
organic cyclic compound cataboli...
aromatic compound catabolic process
reactive oxygen species metaboli...
response to abscisic acid
cofactor metabolic process
secondary metabolite biosyntheti...
cell wall biogenesis
organic substance transport
polysaccharide catabolic process
peptide transport
cell wall macromolecule metaboli...
phenylpropanoid metabolic process
8dpp 16dpp
8dpp 16dpp
8dpp 16dpp
Age
B
T4 vs T0
T24 vs T4
T48 vs T24
response to karrikin
metabolic process
defense response
phenylpropanoid biosynthetic pro...
hormone metabolic process
regulation of transcription, DNA...
plant−type cell wall biogenesis
response to UV
hormone−mediated signaling pathway
signal transduction
cofactor metabolic process
abscisic acid−activated signalin...
anion transmembrane transport
carboxylic acid catabolic process
nitrogen compound transport
peptide transport
drug catabolic process
antibiotic metabolic process
response to chitin
ethylene−activated signaling pat...
response to wounding
response to oxidative stress
defense response to fungus
response to jasmonic acid
cell wall macromolecule metaboli...
macromolecule localization
chemical homeostasis
chromatin organization
response to heat
lipid biosynthetic process
phosphorelay signal transduction...
cofactor biosynthetic process
cellular metal ion homeostasis
response to cold
anatomical structure formation i...
fatty acid metabolic process
cellular response to radiation
plastid organization
regulation of cellular metabolic...
drug transport
organic cyclic compound biosynth...
negative regulation of RNA metab...
circadian rhythm
proteolysis
lipid catabolic process
aging
response to auxin
transition metal ion homeostasis
small molecule metabolic process
cell wall biogenesis
pigment biosynthetic process
chloroplast organization
response to gibberellin
response to red or far red light
response to cytokinin
carbohydrate metabolic process
oxidation−reduction process
hormone biosynthetic process
terpenoid biosynthetic process
cellular glucan metabolic process
response to other organism
transmembrane receptor protein t...
cell wall organization
polysaccharide catabolic process
plant−type cell wall organization
microtubule−based process
response to salt stress
unidimensional cell growth
cytoskeleton organization
cell division
cellular polysaccharide biosynth...
macromolecule biosynthetic process
multicellular organism development
response to organic cyclic compound
auxin−activated signaling pathway
heterocycle metabolic process
developmental process involved i...
photosynthesis, light reaction
fatty acid biosynthetic process
reproductive shoot system develo...
response to biotic stimulus
post−embryonic plant morphogenesis
cellular component organization
response to stimulus
photosynthesis
response to light stimulus
response to water deprivation
organic substance transport
8dpp 16dpp
8dpp 16dpp
8dpp 16dpp
Age
−log10(p−value)
5 10 15 20 25

```
pdf("fig3.pdf", width = 12, height = 16)
pall
dev.off()
```

```
## pdf
##   2
```

Venn diagrams of differntially expressed genes at each consecutive contrast:

```
# Imports required functions. Originally from http://faculty.ucr.edu/~tgirke/Documents/R_BioCond/My_R_S
source("overLapper.R.txt")

# ellipse plotting functions:
source(file = "plotellipse.R")
```

```
venninput <- timeContrasts %>%
    filter(padj < 0.05 & abs(log2FoldChange) >= 1) %>%
    filter(Contrast == "T4 vs T0") %>%
    mutate(Direction = fct_relevel(ifelse(log2FoldChange > 0, "Up", "Down"), "Up")) %>%
    group_by(Contrast, Age, Direction) %>%
    do(data = (.$geneid)) %>%
    arrange(Direction, desc(Age)) %>%
    with(set_names(data, paste(Contrast, Age, Direction)))


OLlist <-
    overLapper(setlist = venninput,
               sep = "_",
               type = "vennsets")

counts <- list(sapply(OLlist$Venn_List, length))

venndf <- data.frame(
    count = as.vector(lengths(OLlist$Venn_List)),
    y = c(4, 4, 3, 1, 4, 3, 1, 3, 1, 2, 3, 1, 2, 2, 2),
    x = c(1, 3, 4, 4, 2, 1, 1, 3, 3, 4, 2, 2, 1, 3, 2)
)

#create venn ellipses
ellipse1 <-
    as.data.frame(
        plotellipse(
            center = c(2.6, 1.4),
            radius = c(1.15, 2.25),
            rotate = 90,
            segments = 360,
            xlab = "",
            ylab = "",
            col = lines[1],
            axes = FALSE,
            main = mymain,
            sub = mysub,
            lwd = mylwd
        )
    )
```

```r
ellipse2 <-
    as.data.frame(
        plotellipse(
            center = c(2.5, 2.5),
            radius = c(1.15, 2.25),
            rotate = 90,
            segments = 360,
            xlab = "",
            ylab = "",
            col = lines[1],
            axes = FALSE,
            main = mymain,
            sub = mysub,
            lwd = mylwd
        )
    )
ellipse3 <-
    as.data.frame(
        plotellipse(
            center = c(2.5, 2.5),
            radius = c(1.15, 2.25),
            rotate = 0,
            segments = 360,
            xlab = "",
            ylab = "",
            col = lines[1],
            axes = FALSE,
            main = mymain,
            sub = mysub,
            lwd = mylwd
        )
    )
ellipse4 <-
    as.data.frame(
        plotellipse(
            center = c(1.4, 2.6),
            radius = c(1.15, 2.25),
            rotate = 0,
            segments = 360,
            xlab = "",
            ylab = "",
            col = lines[1],
            axes = FALSE,
            main = mymain,
            sub = mysub,
            lwd = mylwd
        )
    )

library("sp")
poly1 <- Polygon(ellipse1)
poly2 <- Polygon(ellipse2)
poly3 <- Polygon(ellipse3)
```

```
poly4 <- Polygon(ellipse4)
# create SpatialPolygons objects
p1 <- SpatialPolygons(list(Polygons(list(poly1), "p1")))
p2 <- SpatialPolygons(list(Polygons(list(poly2), "p2")))
p3 <- SpatialPolygons(list(Polygons(list(poly3), "p3")))
p4 <- SpatialPolygons(list(Polygons(list(poly4), "p4")))

# highlight intersections
library(rgeos)
int1 <- gDifference(gDifference(gIntersection(p1, p4), p2), p3)
highlight1<-as.data.frame(slot(int1@polygons[[1]]@Polygons[[1]], "coords"))
highlight1$group <- "1"

int2 <- gDifference(gDifference(gIntersection(p2, p3), p1), p4)
highlight2<-as.data.frame(slot(int2@polygons[[1]]@Polygons[[1]], "coords"))
highlight2$group <- "1"

int3 <- gDifference(gDifference(gIntersection(p1, p2), p3), p4)
highlight3<-as.data.frame(slot(int3@polygons[[1]]@Polygons[[1]], "coords"))
highlight3$group <- "2"

int4 <- gDifference(gDifference(gIntersection(p3, p4), p1), p2)
highlight4<-as.data.frame(slot(int4@polygons[[1]]@Polygons[[1]], "coords"))
highlight4$group <- "2"

highlights <- rbind(highlight1, highlight2, highlight3, highlight4)


ellipses <- rbind(ellipse1, ellipse2, ellipse3, ellipse4)
ellipses$contrast <-
    as.factor(rep(
        c(
            "16dppT4vsT0_up",
            "8dppT4vsT0_up",
            "16dppT4vsT0_down",
            "8dppT4vsT0_down"
        ),
        each = 361
    ))
vennlabels <-
    data.frame(
        contrast = factor(
            c(
                "16dpp_T4vsT0_up",
                "8dpp_T4vsT0_up",
                "16dpp_T4vsT0_down",
                "8dpp_T4vsT0_down"
            )
        ),
        x = c(-2.85, -2, 2, 2.85),
        y = c(4.75, 5.5, 5.5, 4.75)
    )
```

```r
T4venn <- ggplot(data = venndf,
       aes(x = (x - y) / sqrt(2),
           y = (y + x) / sqrt(2))) +
    geom_path(data = ellipses,
              aes(
                  x = (V2 - V1) / sqrt(2),
                  y = (V1 + V2) / sqrt(2),
                  color = contrast
              ),
              size = 1) +
    geom_polygon(data = highlights,
                 aes(x = (y - x) / sqrt(2),
                     y = (y + x) / sqrt(2),
                     fill = group),
                 alpha = 0.25
    ) +
    scale_color_manual(values = c("#e41a1c", "#377eb8", "#4daf4a", "#984ea3")) +
    geom_text(
        data = vennlabels,
        aes(
            x = x,
            y = y,
            label = gsub('_', '\n', contrast)
            ),
        color = c("#377eb8", "#984ea3", "#e41a1c", "#4daf4a"),
        fontface = "bold"
    ) +
    geom_text(
        label = venndf$count,
        nudge_y = -0.10,
        fontface = "bold"
    ) +
    cowplot::theme_cowplot() +
    theme(
        legend.position = "none",
        axis.line = element_blank(),
        axis.text.x = element_blank(),
        axis.text.y = element_blank(),
        axis.ticks = element_blank(),
        axis.title.x = element_blank(),
        axis.title.y = element_blank(),
        panel.background = element_blank(),
        panel.border = element_blank(),
        panel.grid.major = element_blank(),
        panel.grid.minor = element_blank(),
        plot.background = element_blank(),
        plot.margin = unit(c(0.1, 0, 0, 0), "cm")
    ) +
    scale_x_continuous(expand = expand_scale(mult = 0.1)) +
    scale_y_continuous(expand = expand_scale(mult = 0.1)) +
    coord_cartesian(clip = "off", expand = TRUE)
```

GO terms of uniquely upregulated genes in 16dpp at T4vsT0

```r
tgd <- runTopGoAnalysis(DEgeneSet = OLlist$Venn_List$`T4 vs T0 16dpp Up`, dds = dds_exp1, GOdb = GOList
kable(exportGOtable(tgd, "Fisher.weight01", n = 10) %>%
          mutate(Fisher.weight01 = as.numeric(Fisher.weight01)), digits = 20) %>%
    kableExtra::kable_styling(latex_options = c("HOLD_position", "scale_down"),
                              full_width = FALSE)
```

| GO.ID | Term | Annotated | Significant | Expected | Fisher.weight01 |
|---|---|---|---|---|---|
| GO:0009611 | response to wounding | 259 | 29 | 8.49 | 7.90e-09 |
| GO:0015833 | peptide transport | 601 | 13 | 19.69 | 6.30e-06 |
| GO:1901605 | alpha-amino acid metabolic process | 276 | 20 | 9.04 | 5.90e-05 |
| GO:0009414 | response to water deprivation | 318 | 23 | 10.42 | 3.40e-04 |
| GO:1901565 | organonitrogen compound catabolic proces... | 489 | 25 | 16.02 | 3.80e-04 |
| GO:0010200 | response to chitin | 134 | 13 | 4.39 | 4.60e-04 |
| GO:1901698 | response to nitrogen compound | 218 | 22 | 7.14 | 1.23e-03 |
| GO:0046395 | carboxylic acid catabolic process | 131 | 12 | 4.29 | 1.25e-03 |
| GO:0040007 | growth | 534 | 24 | 17.49 | 1.27e-03 |
| GO:0009620 | response to fungus | 308 | 22 | 10.09 | 1.68e-03 |

Write these genes and their Arabidopsis best hits (from cucurbitgenomics.org) to a csv (Supp File 2):

```r
timeContrasts %>%
    filter(
        padj < 0.05 & abs(log2FoldChange) >= 1,
        geneid %in% OLlist$Venn_List$`T4 vs T0 16dpp Up`,
        Age == "16dpp",
        Contrast == "T4 vs T0"
    ) %>%
    rowwise() %>%
    mutate(Annotation = paste(html_text(
        html_nodes(
            read_html(paste0(
                "http://cucurbitgenomics.org/feature/gene/", str_replace(geneid, " ", "")
            )),
            "#11 .odd:nth-child(1) td:nth-child(1) , #11 .odd:nth-child(1) td:nth-child(2) , #11 .odd:n
        )
    ), collapse = "__")) %>%
    separate(
        Annotation,
        into = c("Match Name", "E-value", "Identity", "Description"),
        sep = "__"
    ) %>%
    write.csv(file = "UniqUp16T4.csv")
```

GO terms of uniquely upregulated genes in 8dpp at T4vsT0

```r
tgd <- runTopGoAnalysis(DEgeneSet = OLlist$Venn_List$`T4 vs T0 8dpp Up`,
                        dds = dds_exp1,
                        GOdb = GOList,
                        onts = "BP",
                        nodeSize = 100)
kable(exportGOtable(tgd, "Fisher.weight01", n = 10) %>%
          mutate(Fisher.weight01 = as.numeric(Fisher.weight01)), digits = 20) %>%
    kableExtra::kable_styling(latex_options = c("HOLD_position", "scale_down"),
                              full_width = FALSE)
```

| GO.ID | Term | Annotated | Significant | Expected | Fisher.weight01 |
|---|---|---|---|---|---|
| GO:0044703 | multi-organism reproductive process | 136 | 8 | 2.67 | 7.10e-06 |
| GO:0006518 | peptide metabolic process | 643 | 10 | 12.61 | 9.90e-05 |
| GO:0019748 | secondary metabolic process | 251 | 12 | 4.92 | 1.40e-04 |
| GO:0007154 | cell communication | 1524 | 41 | 29.88 | 1.80e-04 |
| GO:0009636 | response to toxic substance | 152 | 11 | 2.98 | 2.10e-04 |
| GO:0006979 | response to oxidative stress | 380 | 19 | 7.45 | 7.90e-04 |
| GO:0006575 | cellular modified amino acid metabolic p... | 102 | 8 | 2.00 | 8.90e-04 |
| GO:0046686 | response to cadmium ion | 335 | 16 | 6.57 | 9.80e-04 |
| GO:0009651 | response to salt stress | 524 | 21 | 10.27 | 1.61e-03 |
| GO:1901565 | organonitrogen compound catabolic proces... | 489 | 13 | 9.59 | 1.77e-03 |

```r
venninput <- timeContrasts %>%
    filter(padj < 0.05 & abs(log2FoldChange) >= 1) %>%
    filter(Contrast == "T24 vs T4") %>%
    mutate(Direction = fct_relevel(ifelse(log2FoldChange > 0, "Up", "Down"), "Up")) %>%
    group_by(Contrast, Age, Direction) %>%
    do(data = (.$geneid)) %>%
    arrange(Direction, desc(Age)) %>%
    with(set_names(data, paste(Contrast, Age, Direction)))

OLlist <-
    overLapper(setlist = venninput,
               sep = "_",
               type = "vennsets")

counts <- list(sapply(OLlist$Venn_List, length))

venndf <- data.frame(
    count = as.vector(lengths(OLlist$Venn_List)),
    y = c(4, 4, 3, 1, 4, 3, 1, 3, 1, 2, 3, 1, 2, 2, 2),
    x = c(1, 3, 4, 4, 2, 1, 1, 3, 3, 4, 2, 2, 1, 3, 2)
)

ellipses <- rbind(ellipse1, ellipse2, ellipse3, ellipse4)
ellipses$contrast <-
    as.factor(rep(
        c(
            "16dppT24vsT4_up",
            "8dppT24vsT4_up",
            "16dppT24vsT4_down",
            "8dppT24vsT4_down"
        ),
        each = 361
    ))
vennlabels <-
    data.frame(
        contrast = factor(
            c(
                "16dpp_T24vsT4_up",
                "8dpp_T24vsT4_up",
                "16dpp_T24vsT4_down",
                "8dpp_T24vsT4_down"
            )
```

```r
        ),
        x = c(-2.85, -2, 2, 2.85),
        y = c(4.75, 5.5, 5.5, 4.75)
    )

T24venn <- ggplot(data = venndf,
       aes(x = (x - y) / sqrt(2),
           y = (y + x) / sqrt(2))) +
    geom_path(data = ellipses,
              aes(
                  x = (V2 - V1) / sqrt(2),
                  y = (V1 + V2) / sqrt(2),
                  color = contrast
              ),
              size = 1) +
    geom_polygon(data = highlights,
                 aes(x = (y - x) / sqrt(2),
                     y = (y + x) / sqrt(2),
                     fill = group),
                 alpha = 0.25
    ) +
    scale_color_manual(values = c("#e41a1c", "#377eb8", "#4daf4a", "#984ea3")) +
    geom_text(
        data = vennlabels,
        aes(
            x = x,
            y = y,
            label = gsub('_', '\n', contrast)
            ),
        color = c("#377eb8", "#984ea3", "#e41a1c", "#4daf4a"),
        fontface = "bold"
    ) +
    geom_text(
        label = venndf$count,
        nudge_y = -0.10,
        fontface = "bold"
    ) +
    cowplot::theme_cowplot() +
    theme(
        legend.position = "none",
        axis.line = element_blank(),
        axis.text.x = element_blank(),
        axis.text.y = element_blank(),
        axis.ticks = element_blank(),
        axis.title.x = element_blank(),
        axis.title.y = element_blank(),
        panel.background = element_blank(),
        panel.border = element_blank(),
        panel.grid.major = element_blank(),
        panel.grid.minor = element_blank(),
        plot.background = element_blank(),
        plot.margin = unit(c(0.1, 0, 0, 0), "cm")
    ) +
```

```r
    scale_x_continuous(expand = expand_scale(mult = 0.1)) +
    scale_y_continuous(expand = expand_scale(mult = 0.1)) +
    coord_cartesian(clip = "off", expand = TRUE)
```

GO terms of inversely regulated genes at 24 hpi (Up 16 down 8dpp)

```r
tgd <- runTopGoAnalysis(DEgeneSet = OLlist$Venn_List$`T24 vs T4 16dpp Up_T24 vs T4 8dpp Down`,
                        dds = dds_exp1,
                        GOdb = GOList,
                        onts = "BP",
                        nodeSize = 100)

kable(exportGOtable(tgd, "Fisher.weight01", n = 10) %>%
        mutate(Fisher.weight01 = as.numeric(Fisher.weight01)), digits = 20) %>%
    kableExtra::kable_styling(latex_options = c("HOLD_position", "scale_down"),
                              full_width = FALSE)
```

| GO.ID | Term | Annotated | Significant | Expected | Fisher.weight01 |
|---|---|---|---|---|---|
| GO:0019684 | photosynthesis, light reaction | 115 | 17 | 1.63 | 5.20e-13 |
| GO:0055114 | oxidation-reduction process | 1249 | 35 | 17.68 | 7.60e-05 |
| GO:0048869 | cellular developmental process | 662 | 12 | 9.37 | 9.30e-04 |
| GO:0006732 | coenzyme metabolic process | 262 | 11 | 3.71 | 2.05e-03 |
| GO:0007169 | transmembrane receptor protein tyrosine ... | 135 | 7 | 1.91 | 3.11e-03 |
| GO:0009658 | chloroplast organization | 148 | 7 | 2.10 | 5.15e-03 |
| GO:0006790 | sulfur compound metabolic process | 256 | 8 | 3.62 | 5.22e-03 |
| GO:0015979 | photosynthesis | 201 | 22 | 2.85 | 5.58e-03 |
| GO:0048229 | gametophyte development | 312 | 12 | 4.42 | 6.40e-03 |
| GO:0043603 | cellular amide metabolic process | 751 | 14 | 10.63 | 8.06e-03 |

```r
venninput <- timeContrasts %>%
    filter(padj < 0.05 & abs(log2FoldChange) >= 1) %>%
    filter(Contrast == "T48 vs T24") %>%
    mutate(Direction = fct_relevel(ifelse(log2FoldChange > 0, "Up", "Down"), "Up")) %>%
    group_by(Contrast, Age, Direction) %>%
    do(data = (.$geneid)) %>%
    arrange(Direction, desc(Age)) %>%
    with(set_names(data, paste(Contrast, Age, Direction)))


OLlist <-
    overLapper(setlist = venninput,
               sep = "_",
               type = "vennsets")

counts <- list(sapply(OLlist$Venn_List, length))

venndf <- data.frame(
    count = as.vector(lengths(OLlist$Venn_List)),
    y = c(4, 4, 3, 1, 4, 3, 1, 3, 1, 2, 3, 1, 2, 2, 2),
    x = c(1, 3, 4, 4, 2, 1, 1, 3, 3, 4, 2, 2, 1, 3, 2)
)

ellipses <- rbind(ellipse1, ellipse2, ellipse3, ellipse4)
ellipses$contrast <-
```

```r
    as.factor(rep(
        c(
            "16dppT48vsT24_up",
            "8dppT48vsT24_up",
            "16dppT48vsT24_down",
            "8dppT48vsT24_down"
        ),
        each = 361
    ))

vennlabels <-
    data.frame(
        contrast = factor(
            c(
                "16dpp_T48vsT24_up",
                "8dpp_T48vsT24_up",
                "16dpp_T48vsT24_down",
                "8dpp_T48vsT24_down"
            )
        ),
        x = c(-2.85, -2, 2, 2.85),
        y = c(4.75, 5.5, 5.5, 4.75)
    )

T48venn <- ggplot(data = venndf,
       aes(x = (x - y) / sqrt(2),
           y = (y + x) / sqrt(2))) +
    geom_path(data = ellipses,
              aes(
                  x = (V2 - V1) / sqrt(2),
                  y = (V1 + V2) / sqrt(2),
                  color = contrast
              ),
              size = 1) +
    geom_polygon(data = highlights,
                 aes(x = (y - x) / sqrt(2),
                     y = (y + x) / sqrt(2),
                     fill = group),
                 alpha = 0.25
    ) +
    scale_color_manual(values = c("#e41a1c", "#377eb8", "#4daf4a", "#984ea3")) +
    geom_text(
        data = vennlabels,
        aes(
            x = x,
            y = y,
            label = gsub('_', '\n', contrast)
            ),
        color = c("#377eb8", "#984ea3", "#e41a1c", "#4daf4a"),
        fontface = "bold"
    ) +
    geom_text(
        label = venndf$count,
```

```
        nudge_y = -0.10,
        fontface = "bold"
    ) +
    cowplot::theme_cowplot() +
    theme(
        legend.position = "none",
        axis.line = element_blank(),
        axis.text.x = element_blank(),
        axis.text.y = element_blank(),
        axis.ticks = element_blank(),
        axis.title.x = element_blank(),
        axis.title.y = element_blank(),
        panel.background = element_blank(),
        panel.border = element_blank(),
        panel.grid.major = element_blank(),
        panel.grid.minor = element_blank(),
        plot.background = element_blank(),
        plot.margin = unit(c(0.1, 0, 0, 0), "cm")
    ) +
    scale_x_continuous(expand = expand_scale(mult = 0.1)) +
    scale_y_continuous(expand = expand_scale(mult = 0.1)) +
    coord_cartesian(clip = "on", expand = TRUE)
```

```
vennPlots <- cowplot::plot_grid(T4venn, T24venn, T48venn,
                                label_size = 16,
                                labels = c("C", "D", "E"),
                                align = "v",
                                axis = "l",
                                ncol = 1,
                                rel_widths = c(0.1, 1, 1, 1))
```

Make Figure 2 PCA, Summary of DEGs and Venn diagrams:

```
expPlots <- cowplot::plot_grid(pcaPlot1, sumPlot1,
                               align = "v",
                               labels = c("A", "B"),
                               label_size = 16,
                               axis = "l",
                               ncol = 1
                               )

cowplot::plot_grid(
    expPlots,
    NULL,
    vennPlots,
    ncol = 3,
    align = "v",
    axis = "b",
    rel_widths = c(2, 0.1, 1.5)
)
```

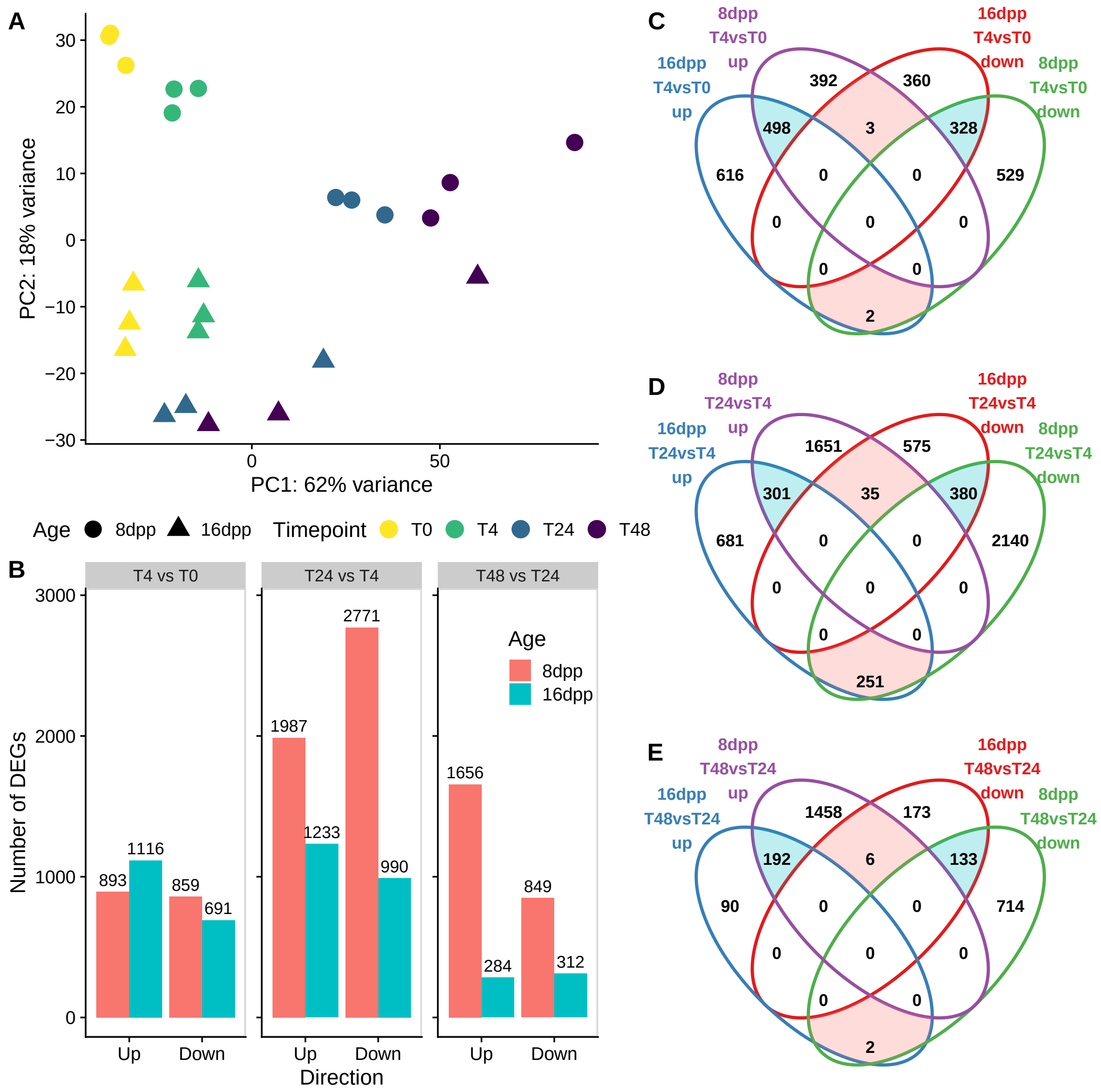


```r
pdf("fig2.pdf", width = 12, height = 10)
cowplot::plot_grid(expPlots, NULL, vennPlots, ncol = 3, rel_widths = c(2, 0.1, 1.5))
dev.off()
```

```
## pdf
##   2
```

### Transciptome Experiment 2

#### QuantSeq 3' mRNA mapping pipeline

##### Read cleanup and Quality control

```
mkdir infection24_3prime
cd infection24_3prime
mkdir qualitycheck

# To install multiQC we need to work with Anaconda python environment
module load Anaconda2/4.2.0
export PATH=$PATH:$HOME/anaconda2/bin
conda create --name multiQC
source activate multiQC

# Install the multiQC package
conda install -c bioconda multiqc
source deactivate

# QC before cleaning
module load fastQC
fastqc --outdir qualitycheck --format fastq --threads 8 *fastq.gz

export PATH=$PATH:$HOME/anaconda2/bin
source activate multiQC
multiqc -o ~/infection24_3prime/ -n multiqc_raw ~/infection24_3prime/qualitycheck/
source deactivate

# Make a polyA fasta file for trimming
printf '>\nAAAAAAAAAAAAAAAAAAA' > polyA.fa
gzip polyA.fa

# Trim and clean reads with BBMap
module load BBMap
for sample in *fastq.gz;
do cat $sample | bbduk.sh in=stdin.fq.gz out=${sample}_trimmed_clean \
ref=polyA.fa.gz,$ADAPTERS/TruSeq3-SE.fa \
ftl=12 k=13 ktrim=r useshortkmers=t mink=5 qtrim=r trimq=10 minlength=20 int=f ;
done

# QC after trimming
mkdir qc_trimmed_clean

fastqc --outdir qc_trimmed_clean --format fastq --threads 8 *fastq.gz_trimmed_clean

export PATH=$PATH:$HOME/anaconda2/bin
source activate multiQC
multiqc -o ~/infection24_3prime/ \
-n multiqc_clean ~/infection24_3prime/qc_trimmed_clean/
source deactivate
```

##### Make extended transcriptome file

Extract chromosome sizes:

```
wget https://github.com/bedops/bedops/releases/download/v2.4.35/bedops_linux_x86_64-v2.4.35.tar.bz2
tar jxvf bedops_linux_x86_64-v2.4.35.tar.bz2
export PATH=$PATH:$HOME/bin

cd ~/infectionRNAseq/genome/cl9930-v2
gff2bed < cucumber_ChineseLong_v2.gff3 > cucumber_ChineseLong_v2.bed

module load SAMtools
cut -f 1,2 cucumber_ChineseLong_v2_genome.fa.fai > chrom.sizes
```

Transcript 3'UTR extension script in R:

```
gff <- read.delim('cucumber_ChineseLong_v2.gff3', header = FALSE, sep = '\t')
chromSizes <- read.table(file = "chrom.sizes") %>%
  dplyr::rename(chrom = V1,
         maxLength = V2)

names(gff) <- c("chrom", "source", "feature",
                "start", "end", "score",
                "strand", "phase", "attr")

gff <- gff %>%
  mutate(parent = ifelse(str_detect(attr, "CsaUN"),
                         str_remove(str_extract(string = attr,
                                               pattern = "ID=CsaUNG[0-9]*"),
                                    pattern = "ID="),
                         str_remove(str_extract(string = attr,
                                               pattern = "ID=Csa[0-9]G[0-9]*"),
                                    pattern = "ID=")
                         )
         )

extend <- 1000
gff_name <- paste0("cucumber_ChineseLong_v2_extended3UTR_", extend,".gff3")

gff_new <- left_join(gff, chromSizes) %>%
  group_by(parent, feature) %>%
  mutate(id = row_number()) %>% # add id index for each feature
  mutate(
    sug_start = case_when(
      feature %in% c("gene", "mRNA") &
        strand == "-" ~ start - extend,
      feature == "three_prime_utr" &
        id == max(id) & strand == "-" ~ start - extend, #use max id to get the last 3'UTR
      feature == "exon" &
        id == max(id) & strand == "-" ~ start - extend, #use max id to get the last exon,
      TRUE ~ as.numeric(start)
    )
  ) %>%
  mutate(
    sug_end = case_when(
      feature %in% c("gene", "mRNA") &
        strand == "+" ~ end + extend,
      feature == "three_prime_utr" &
```

```
        id == max(id) & strand == "+" ~ end + extend, #use max id to get the last 3'UTR
      feature == "exon" &
        id == max(id) & strand == "+" ~ end + extend, #use max id to get the last exon
      TRUE ~ as.numeric(end)
    )
  )  %>%
  ungroup() %>%
  mutate(sug_start = as.integer(ifelse(sug_start <= 0, 1, sug_start)),  # limit on the left
         sug_end = as.integer(ifelse(sug_end > maxLength, maxLength, sug_end)) # limit on the right
  )

gff_new2 <- gff_new %>% # Find the distances between genes and the next start
  left_join(gff_new %>% # site on the same strand
              group_by(chrom, strand) %>%
              arrange(chrom, strand, start) %>%
              filter(feature %in% c("gene")) %>%
              mutate(next_gene = ifelse(strand == "-",
                                        lag(end, 1),
                                        lead(start, 1))
              ) %>% mutate(next_gene = ifelse(is.na(next_gene),
                                         ifelse(strand == "-", 0, maxLength + 1),
                                         next_gene))
            ) %>%
  group_by(chrom) %>%
    fill(next_gene) %>% # Identify suggested start sites that overlap the next gene
    mutate(
    start_ext = case_when(
      feature %in% c("gene", "mRNA") &
        sug_start <= next_gene & strand == "-" ~ next_gene + 1, # if there is an overlap stop one base
      feature == "three_prime_utr" &
        id == max(id) & sug_start <= next_gene & strand == "-" ~ next_gene + 1, #use max id to get the
      feature == "exon" &
        id == max(id) & sug_start <= next_gene & strand == "-" ~ next_gene + 1, #use max id to get the
      TRUE ~ as.numeric(sug_start)
    )
  ) %>%
  mutate(
    end_ext = case_when(
      feature %in% c("gene", "mRNA") &
        sug_end >= next_gene & strand == "+" ~ next_gene - 1,
      feature == "three_prime_utr" &
        id == max(id) & sug_end >= next_gene & strand == "+" ~ next_gene - 1, #use max id to get the la
      feature == "exon" &
        id == max(id) & sug_end >= next_gene & strand == "+" ~ next_gene - 1, #use max id to get the la
      TRUE ~ as.numeric(sug_end)
    )
  ) %>%
  ungroup() %>%
  mutate(start_ext = as.integer(ifelse(start_ext <= 0, 1, start_ext)),  # limit on the left
         end_ext = as.integer(ifelse(end_ext > maxLength, maxLength, end_ext)) # limit on the right
  ) %>%
  mutate(delta = (end_ext - start_ext) - (end - start)) %>% # check the difference before and after ext
  mutate(final_start = ifelse(delta < 0, start, start_ext),
```

```
         final_end = ifelse(delta < 0, end, end_ext),
         final_delta = (final_end - final_start) - (end - start))

gff_new2 %>%
  filter(feature == "gene") %>%
  ggplot() +
  geom_density(aes(x = final_delta))

gff_out <- gff_new2 %>%
  mutate(start = final_start,
         end = final_end
         ) %>%
  dplyr::select(-parent, -id, -maxLength,
         -sug_start, -sug_end, -start_ext,
         -end_ext, -final_start, -final_end,
         -delta, -final_delta, -next_gene
         )

write_delim(gff_out,
            path = gff_name,
            delim = "\t",
            col_names = FALSE)
```

Use cufflinks gffread to output a fasta file based on the new gff3 file:

```
module load Cufflinks/2.2.1
gffread -g ../genome/cl9930-v2/cucumber_ChineseLong_v2_genome.fa -w cucumber_ChineseLong_v2_cdna_extend
```

##### Salmon mapping

```
wget https://github.com/COMBINE-lab/salmon/releases/download/v0.12.0/salmon-0.12.0_linux_x86_64.tar.gz
tar xvfz salmon-0.12.0_linux_x86_64.tar.gz

SALMON=salmon-0.12.0_linux_x86_64/bin

# Index transcriptome
$SALMON/salmon index -t cucumber_ChineseLong_v2_cdna_extended3UTR_1000.fa -i cucumber_ChineseLong_v2_cd

# Perform the mapping using Salmon
mkdir quants_1000

for fn in `ls clean_reads/*fastq`;
do
nodir=${fn#clean_reads/}
echo "Processing sample ${nodir}"
$SALMON/salmon quant -i cucumber_ChineseLong_v2_cdna_extended3UTR_1000_salmon_index \
    -l SF \
    -r ${fn} \
    -p 16 \
    -o quants_1000/${nodir%_S*}_quant \
    --noLengthCorrection \
    --validateMappings;
done
```

#### Differential expression pipeline

##### Import data using tximport

```
dirs <- list.files("./quants_1000")
samples <- as_tibble(stringr::str_split(dirs, pattern = "\\.", n = 2, simplify = T)[, 1]) %>%
  separate(col = value, into = c("well", "age", "timepoint", "treatment", "rep"), sep = "_", remove = F[illegible]
  dplyr::rename(name = value) %>%
  mutate(directory = dirs)

# duplicate the controll T0 samples to have Inoc T0 for statistical purposes
fake_inocT0 <- samples %>%
  filter(timepoint == "T0") %>%
  mutate(treatment = "Inoc",
         name = str_replace(name, "Cont", "InocFake"))

samples <- bind_rows(samples, fake_inocT0)

files <- file.path("quants_1000", samples$directory, "quant.sf")
names(files) <- samples$name

samples <- samples %>%
  mutate(
  timepoint = fct_relevel(timepoint, "T0", "T2", "T4", "T8", "T12", "T18", "T24"),
  age = fct_relevel(age, "8dpp"),
  treatment = fct_relevel(samples$treatment, "Cont"),
  condition = as_factor(
  paste0(samples$age, "_", samples$timepoint, "_", samples$treatment)
  ),
  condition = fct_relevel(
  condition,
  "8dpp_T0_Cont",
  "8dpp_T2_Cont",
  "8dpp_T4_Cont",
  "8dpp_T8_Cont",
  "8dpp_T12_Cont",
  "8dpp_T18_Cont",
  "8dpp_T24_Cont",
  "16dpp_T0_Cont",
  "16dpp_T2_Cont",
  "16dpp_T4_Cont",
  "16dpp_T8_Cont",
  "16dpp_T12_Cont",
  "16dpp_T18_Cont",
  "16dpp_T24_Cont",
  "8dpp_T0_Inoc",
  "8dpp_T2_Inoc",
  "8dpp_T4_Inoc",
  "8dpp_T8_Inoc",
  "8dpp_T12_Inoc",
  "8dpp_T18_Inoc",
  "8dpp_T24_Inoc",
  "16dpp_T0_Inoc",
  "16dpp_T2_Inoc",
```

```
  "16dpp_T4_Inoc",
  "16dpp_T8_Inoc",
  "16dpp_T12_Inoc",
  "16dpp_T18_Inoc",
  "16dpp_T24_Inoc"
  )
  )

# make tx2gene file
tx2gene <- read.table(file = files[1], sep = "\t", header = TRUE)
tx2gene <- tx2gene[1]
tx2gene$geneid <- str_replace(tx2gene$Name, pattern = "\\..*", replacement = "")

txiAll <- tximport(files, type = "salmon", tx2gene = tx2gene, dropInfReps = TRUE, countsFromAbundance =

ddsTxiAll <- DESeqDataSetFromMatrix(round(txiAll$counts), colData = samples, design = ~ condition)

# filter low expression genes: genes were 75 samples have less than 5 reads
ddsTxiAll_filt <- ddsTxiAll[rowSums(counts(ddsTxiAll) <= 5) < 75, ]
```

Reads mapping stats

```
# Import JSON files for sample read mapping results

jsons <- file.path("quants_1000", samples$directory, "aux_info", "meta_info.json")
samples$totalReads <- sapply(jsons, FUN = function(x) {
    fromJSON(file = x)$num_processed
})

samples$Mapped <- sapply(jsons, FUN = function(x) {
    fromJSON(file = x)$num_mapped
})
```

```
expDesign <- samples %>%
    group_by(age, timepoint, treatment) %>% summarise(Reps = n())

knitr::kable(as.data.frame(expDesign), caption = "The experimental design") %>%
    kable_styling(bootstrap_options = "striped", full_width = FALSE)
```

```
samples %>%
    mutate(sampleName = name,
           Unmapped = totalReads - Mapped) %>%
    arrange(age, timepoint) %>%
    mutate(sampleName = fct_inorder(sampleName)) %>%
    gather(key = "Reads", value = "Count", -directory, -age, -timepoint, -condition, -sampleName, -total
    ggplot(aes(x = sampleName, y = Count, fill = Reads,label = paste0(round(Count / totalReads * 100, di
    geom_bar(stat = "identity", position = "stack") +
    geom_text(size = 2, stat = "identity", position = "stack", hjust = 0.5) +
    theme(axis.text.x = element_text(angle = 90, vjust = 0.5))
```

Table 2: The experimental design

| age | timepoint | treatment | Reps |
|---|---|---|---|
| 8dpp | T0 | Cont | 3 |
| 8dpp | T0 | Inoc | 3 |
| 8dpp | T2 | Cont | 3 |
| 8dpp | T2 | Inoc | 3 |
| 8dpp | T4 | Cont | 3 |
| 8dpp | T4 | Inoc | 3 |
| 8dpp | T8 | Cont | 3 |
| 8dpp | T8 | Inoc | 3 |
| 8dpp | T12 | Cont | 3 |
| 8dpp | T12 | Inoc | 3 |
| 8dpp | T18 | Cont | 3 |
| 8dpp | T18 | Inoc | 3 |
| 8dpp | T24 | Cont | 3 |
| 8dpp | T24 | Inoc | 3 |
| 16dpp | T0 | Cont | 3 |
| 16dpp | T0 | Inoc | 3 |
| 16dpp | T2 | Cont | 3 |
| 16dpp | T2 | Inoc | 3 |
| 16dpp | T4 | Cont | 3 |
| 16dpp | T4 | Inoc | 3 |
| 16dpp | T8 | Cont | 3 |
| 16dpp | T8 | Inoc | 3 |
| 16dpp | T12 | Cont | 3 |
| 16dpp | T12 | Inoc | 3 |
| 16dpp | T18 | Cont | 3 |
| 16dpp | T18 | Inoc | 3 |
| 16dpp | T24 | Cont | 3 |
| 16dpp | T24 | Inoc | 3 |

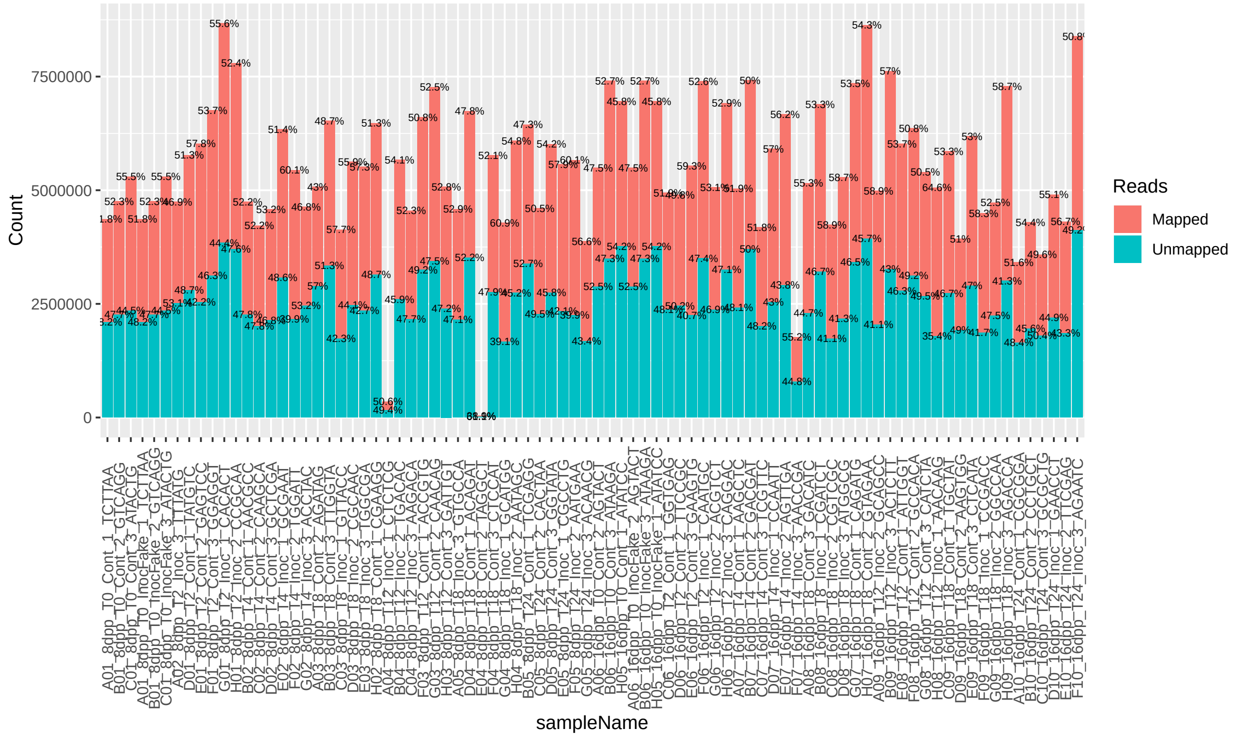


```r
# remove samples with less than 1M reads

ddsTxiAll_filt <- ddsTxiAll_filt[, samples$totalReads > 1e6]

# average reads mapping

samples %>%
  filter(totalReads > 1e6) %>%
  summarize(avgReadMapping = mean(Mapped / totalReads))
```

```
## # A tibble: 1 x 1
##   avgReadMapping
##            <dbl>
## 1          0.533
```

##### DESeq2 analysis

```r
dds <- DESeq(ddsTxiAll_filt)

save(dds, file = "infectionTC_8vs16_ContvsInoc_dds.RData")
```

Principal component analysis

```r
vst <- vst(dds, blind = TRUE)

pcaData <- plotPCA(vst, returnData=TRUE, ntop = 500)

percentVar <- round(100 * attr(pcaData, "percentVar"))

pcaData <- pcaData %>%
    mutate(Age = vst$age,
```

```
           Timepoint = vst$timepoint,
           Treatment = vst$treatment)

pcaExp2 <- ggplot(pcaData, aes(PC1, PC2, color = Timepoint, shape = Age)) +
    geom_point(size = 4, stroke = 2) +
    geom_point(aes(alpha = Treatment, fill = Timepoint), size = 5, stroke = 0) +
    scale_shape_manual(values = c(21, 24, 21, 24)) +
    scale_alpha_manual(values=c(0.05, 1, 0.05, 1)) +
    xlab(paste0("PC1: ",percentVar[1],"% variance")) +
    ylab(paste0("PC2: ",percentVar[2],"% variance")) +
    geom_point(data = filter(pcaData, Timepoint == "T0"),
               aes(x = PC1, y = PC2, shape = Timepoint),
               color = "black",
               shape = 8,
               size = 2) +
    cowplot::theme_cowplot() +
    guides(
    shape = guide_legend(order = 1),
    fill = guide_legend(order = 0, nrow=2, byrow=TRUE),
    alpha = guide_legend(title = "Treatment",
                                override.aes = list(shape = c(22, 15), stroke = 2, alpha = 1))
  ) +
  scale_color_viridis_d(direction = -1) +
  scale_fill_viridis_d(direction = -1) +
    theme(legend.position = "bottom",
          legend.direction = "horizontal") +
    coord_fixed(ratio = percentVar[2] / percentVar[1])

pcaExp2
```

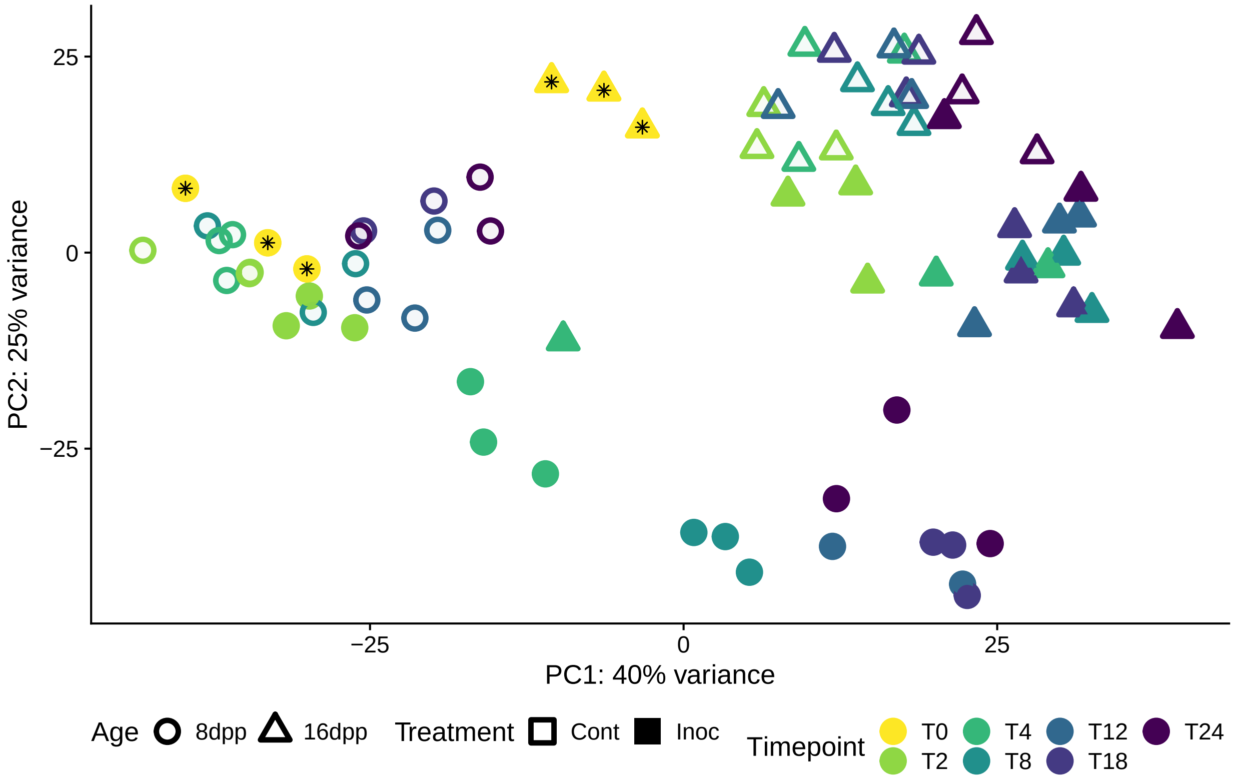


Differential expression contrast results

```
IvsC_T2_8dpp <- lfcShrink(dds, contrast = c("condition", "8dpp_T2_Inoc", "8dpp_T2_Cont"))
IvsC_T4_8dpp <- lfcShrink(dds, contrast = c("condition", "8dpp_T4_Inoc", "8dpp_T4_Cont"))
IvsC_T8_8dpp <- lfcShrink(dds, contrast = c("condition", "8dpp_T8_Inoc", "8dpp_T8_Cont"))
IvsC_T12_8dpp <- lfcShrink(dds, contrast = c("condition", "8dpp_T12_Inoc", "8dpp_T12_Cont"))
IvsC_T18_8dpp <- lfcShrink(dds, contrast = c("condition", "8dpp_T18_Inoc", "8dpp_T18_Cont"))
IvsC_T24_8dpp <- lfcShrink(dds, contrast = c("condition", "8dpp_T24_Inoc", "8dpp_T24_Cont"))

IvsC_T2_16dpp <- lfcShrink(dds, contrast = c("condition", "16dpp_T2_Inoc", "16dpp_T2_Cont"))
IvsC_T4_16dpp <- lfcShrink(dds, contrast = c("condition", "16dpp_T4_Inoc", "16dpp_T4_Cont"))
IvsC_T8_16dpp <- lfcShrink(dds, contrast = c("condition", "16dpp_T8_Inoc", "16dpp_T8_Cont"))
IvsC_T12_16dpp <- lfcShrink(dds, contrast = c("condition", "16dpp_T12_Inoc", "16dpp_T12_Cont"))
IvsC_T18_16dpp <- lfcShrink(dds, contrast = c("condition", "16dpp_T18_Inoc", "16dpp_T18_Cont"))
IvsC_T24_16dpp <- lfcShrink(dds, contrast = c("condition", "16dpp_T24_Inoc", "16dpp_T24_Cont"))

timeContrasts <-
  bind_rows(
  "IvsC_T2_8dpp" = as.data.frame(IvsC_T2_8dpp),
  "IvsC_T4_8dpp" = as.data.frame(IvsC_T4_8dpp),
  "IvsC_T8_8dpp" = as.data.frame(IvsC_T8_8dpp),
  "IvsC_T12_8dpp" = as.data.frame(IvsC_T12_8dpp),
  "IvsC_T18_8dpp" = as.data.frame(IvsC_T18_8dpp),
  "IvsC_T24_8dpp" = as.data.frame(IvsC_T24_8dpp),
  "IvsC_T2_16dpp" = as.data.frame(IvsC_T2_16dpp),
  "IvsC_T4_16dpp" = as.data.frame(IvsC_T4_16dpp),
  "IvsC_T8_16dpp" = as.data.frame(IvsC_T8_16dpp),
  "IvsC_T12_16dpp" = as.data.frame(IvsC_T12_16dpp),
```

```
  "IvsC_T18_16dpp" = as.data.frame(IvsC_T18_16dpp),
  "IvsC_T24_16dpp" = as.data.frame(IvsC_T24_16dpp)
  , .id = "Contrast") %>%
  mutate(geneid = rep(rownames(dds), 12))
```

Alluvial plots for all DE genes

```
lodesTC <- timeContrasts %>%
    mutate(Direction = case_when(
        padj < 0.05 & log2FoldChange >= 1 ~ "Up",
        padj < 0.05 & log2FoldChange <= -1 ~ "Down",
        TRUE ~ "notDE"
    )) %>%
    mutate(Direction = fct_relevel(Direction, "Down", "Up", "notDE"),
           Age = str_extract(Contrast, "[0-9]*dpp"),
           Contrast = str_remove(Contrast, "_[0-9]*dpp")) %>%
    mutate(Contrast = fct_inorder(Contrast),
           Age = fct_relevel(Age, "8dpp")
    ) %>%
    dplyr::select(geneid, Direction, Contrast, Age) %>%
    group_by(geneid, Age) %>%
    filter(sum(as.numeric(Direction)) < 18) %>% # because as.numeric('notDE') = 3
    summarise(flow = paste0(Direction, collapse = "_")) %>%
    group_by(Age, flow) %>%
    summarise(freq = n()) %>%
    separate(col = flow, into = c("T2", "T4", "T8", "T12", "T18", "T24")) %>%
    mutate(first = T4,
           second = T8,
           third = T12,
           forth = T18,
           fifth = T24,
           sixth = T24) %>%
    to_lodes_form(key = "Contrast", axes = 2:7) %>%
    mutate(diffExp = fct_relevel(stratum, "Up", "notDE")) %>%
    mutate(flowColor = case_when(
      Contrast == "T2" ~ first,
      Contrast == "T4" ~ second,
      Contrast == "T8" ~ third,
      Contrast == "T12" ~ forth,
      Contrast == "T18" ~ fifth,
      Contrast == "T24" ~ sixth
    )) %>%
  group_by(Age, Contrast, diffExp) %>%
  mutate(stratumSize = sum(freq),
         number = stratumSize/ n())

alluvialExp2 <- lodesTC%>%
    ggplot(
        aes(
            x = Contrast,
            stratum = diffExp,
            alluvium = alluvium,
            y = freq,
            fill = diffExp,
```

```
            label = freq
        )
    ) +
    geom_flow(
        aes(fill = as.factor(flowColor)),
        stat = "alluvium",
        alpha = 0.75,
        size = 0.5
    ) +
    geom_stratum(width = 0.5) +
    geom_text(aes(label = ifelse(stratumSize > 100, number, NA)),
              stat = "stratum",
              size = 3,
              position = position_nudge(y = 0)) +
    labs(x = "Timepoint vs. Control", y = "Number of genes") +
    scale_fill_manual(values = c("#fc8d62", "#8da0cb", "#66c2a5"),
                      breaks = c("Up", "notDE", "Down"),
                      name = "Differential\nexpression") +
    ggrepel::geom_text_repel(stat = "stratum",
                             size = 3,
                             nudge_y = 100,
                             aes(label = ifelse(stratumSize <= 100,
                                                number,
                                                NA
                                                )
                                 )
                             ) +
    cowplot::theme_cowplot(font_size = 14) +
    theme(strip.text = element_text(colour = "grey10",
                                    size = rel(0.8),
                                    margin = margin(0.5 * 7,
                                                    0.5 * 7,
                                                    0.5 * 7,
                                                    0.5 * 7)
                                    ),
          legend.position = c(0.85, 0.85)
          ) +
    lemon::facet_rep_wrap(~ Age) +
    cowplot::panel_border()

alluvialExp2
```

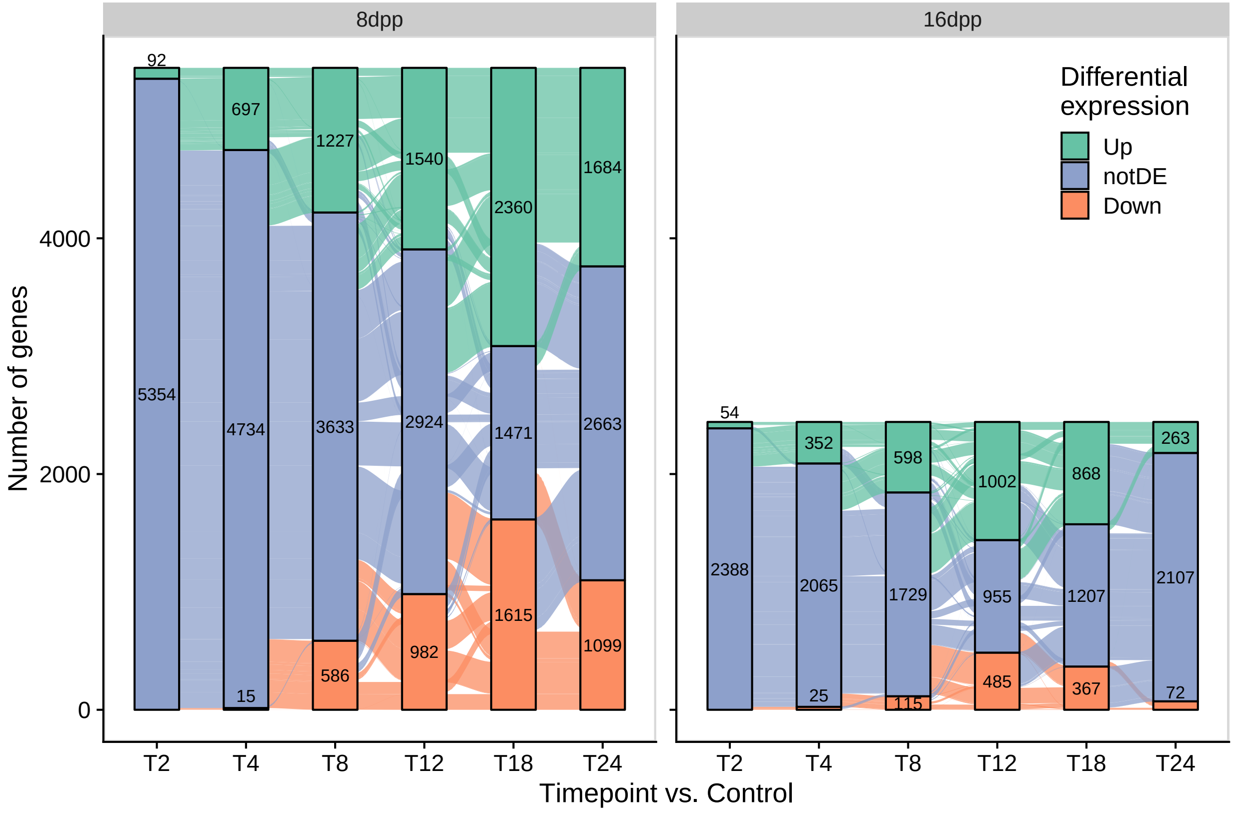


Get some summary numbers for alluvial results

```
# upregulated genes at 4 hpi and are continuously upregulated in every subsequent timepoint
lodesTC %>%
    filter(Age == "8dpp", Contrast == "T4", diffExp == "Up") %>%
    filter_at(.vars = c("second","third", "forth", "fifth", "sixth"),
              all_vars(. == "Up")) %>%
    summarize(sum(freq))
```

```
## # A tibble: 36 x 4
## # Groups:   Age, Contrast [12]
##    Age   Contrast diffExp `sum(freq)`
##    <fct> <fct>    <fct>         <int>
##  1 8dpp  T2       Up                0
##  2 8dpp  T2       notDE             0
##  3 8dpp  T2       Down              0
##  4 8dpp  T4       Up              424
##  5 8dpp  T4       notDE             0
##  6 8dpp  T4       Down              0
##  7 8dpp  T8       Up                0
##  8 8dpp  T8       notDE             0
##  9 8dpp  T8       Down              0
## 10 8dpp  T12      Up                0
## # ... with 26 more rows
```

```
# how many of the upregulated 8dpp genes are newly upregulated at T24
lodesTC %>%
    filter(Age == "8dpp", Contrast == "T24", diffExp == "Up") %>%
```

```r
    filter_at(.vars = c("second","third", "forth"), all_vars(. != "Up")) %>%
    summarise(sum(freq))
```

```
## # A tibble: 36 x 4
## # Groups:   Age, Contrast [12]
##    Age   Contrast diffExp `sum(freq)`
##    <fct> <fct>    <fct>         <int>
##  1 8dpp  T2       Up                0
##  2 8dpp  T2       notDE             0
##  3 8dpp  T2       Down              0
##  4 8dpp  T4       Up                0
##  5 8dpp  T4       notDE             0
##  6 8dpp  T4       Down              0
##  7 8dpp  T8       Up                0
##  8 8dpp  T8       notDE             0
##  9 8dpp  T8       Down              0
## 10 8dpp  T12      Up                0
## # ... with 26 more rows
```

```r
# how many 16 dpp genes upregulated at 4 hpi, are not differentially expressed at 8 hpi
lodesTC %>%
    filter(Age == "16dpp", Contrast == "T4", diffExp == "Up", second != "Up") %>%
    summarise(sum(freq))
```

```
## # A tibble: 36 x 4
## # Groups:   Age, Contrast [12]
##    Age   Contrast diffExp `sum(freq)`
##    <fct> <fct>    <fct>         <int>
##  1 8dpp  T2       Up                0
##  2 8dpp  T2       notDE             0
##  3 8dpp  T2       Down              0
##  4 8dpp  T4       Up                0
##  5 8dpp  T4       notDE             0
##  6 8dpp  T4       Down              0
##  7 8dpp  T8       Up                0
##  8 8dpp  T8       notDE             0
##  9 8dpp  T8       Down              0
## 10 8dpp  T12      Up                0
## # ... with 26 more rows
```

```r
# how many continuously upregulated through 24 hpi
lodesTC %>%
    filter(Age == "16dpp", Contrast == "T4", diffExp == "Up") %>%
    filter_at(.vars = c("second","third", "forth", "fifth", "sixth"),
              all_vars(. == "Up")) %>%
    summarize(sum(freq))
```

```
## # A tibble: 36 x 4
## # Groups:   Age, Contrast [12]
##    Age   Contrast diffExp `sum(freq)`
##    <fct> <fct>    <fct>         <int>
##  1 8dpp  T2       Up                0
##  2 8dpp  T2       notDE             0
##  3 8dpp  T2       Down              0
##  4 8dpp  T4       Up                0
```

```
##   5 8dpp  T4       notDE             0
##   6 8dpp  T4       Down              0
##   7 8dpp  T8       Up                0
##   8 8dpp  T8       notDE             0
##   9 8dpp  T8       Down              0
##  10 8dpp  T12      Up                0
## # ... with 26 more rows
```

```
cowplot::plot_grid(pcaExp2, alluvialExp2, align = "v", axis = "l", ncol = 1, labels = "AUTO")
```

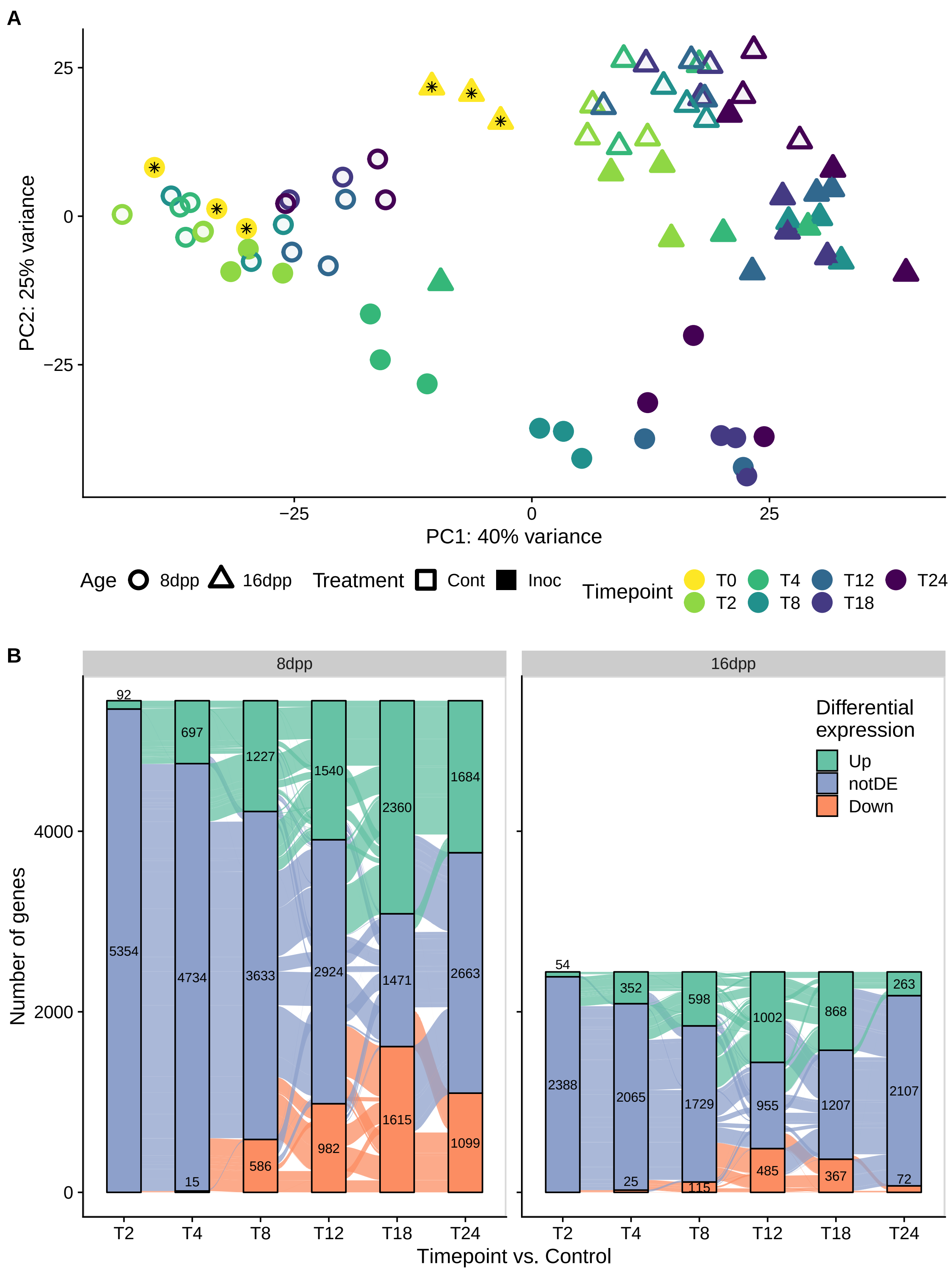
A
PC2: 25% variance
PC1: 40% variance
25
0
−25
−25
0
25
Age 8dpp 16dpp
Treatment Cont Inoc
Timepoint T0 T2 T4 T8 T12 T18 T24
B
8dpp
16dpp
Number of genes
4000
2000
0
92
5354
697
4734
15
1227
3633
586
1540
2924
982
2360
1471
1615
1684
2663
1099
54
2388
352
2065
25
598
1729
115
1002
955
485
868
1207
367
263
2107
72
Differential expression
Up
notDE
Down
T2
T4
T8
T12
T18
T24
Timepoint vs. Control

```
pdf(file = "fig5.pdf", width = 9, height =  12)
cowplot::plot_grid(pcaExp2, alluvialExp2, align = "v", axis = "l", ncol = 1, labels = "AUTO")
dev.off()
```

```
## pdf
##   2
```

Just as a check we will compare the experiments using pca Supplemental Figure 4.

```
vst_48hrs <- vst(dds_exp1, blind = TRUE)
vst_48hrs <- vst_48hrs[, vst_48hrs$timepoint != "T48"]

vst_filt <- vst[, vst$timepoint %in% c("T0", "T4", "T24")]

vst_48hrs <- vst_48hrs[rownames(vst_48hrs) %in% rownames(vst_filt), ]
vst_filt <- vst_filt[rownames(vst_filt) %in% rownames(vst_48hrs), ]

vst_48hrs_8dpp <- vst_48hrs[, vst_48hrs$age == "8dpp"]
vst_48hrs_16dpp <- vst_48hrs[, vst_48hrs$age == "16dpp"]


vst_filt_8dpp <- vst_filt[, vst_filt$age == "8dpp" & vst_filt$treatment == "Inoc"]
vst_filt_16dpp <- vst_filt[, vst_filt$age == "16dpp" & vst_filt$treatment == "Inoc"]

both_8dpp <- cbind(assay(vst_filt_8dpp), assay(vst_48hrs_8dpp))
both_16dpp <- cbind(assay(vst_filt_16dpp), assay(vst_48hrs_16dpp))
bothAges <- cbind(both_8dpp, both_16dpp)

pcaBoth <- prcomp(t(bothAges))

percentVar <- pcaBoth$sdev^2 / sum( pcaBoth$sdev^2 )

as.data.frame(prcomp(t(bothAges))$x) %>%
  mutate(sample = rownames(.),
         timepoint = fct_relevel(str_extract(sample, pattern = "T[0-9]*"), "T24", after = Inf),
         exp = c(rep(c("1", "2"), each = 9), rep(c("1", "2"), each = 9)),
         age = rep(c("8dpp", "16dpp"), each = 18)
           ) %>%
    mutate(Age = fct_relevel(age, "8dpp")) %>%
  ggplot(aes(x = PC1, y = PC2, color = timepoint, shape = Age)) +
    geom_point(size = 4, stroke = 2) +
    geom_point(aes(fill = timepoint, shape = Age, alpha = exp), size = 5, stroke = 0) +
    cowplot::theme_cowplot(font_size = 14) +
    scale_shape_manual(values = c(21, 24, 21, 24)) +
    scale_alpha_manual(values = c(0.05, 1, 0.05, 1)) +
    guides(alpha = guide_legend(title = "Experiment",
                                override.aes = list(shape = c(22, 15), stroke = 2, alpha = 1))) +
    xlab(paste0("PC1: ",round(percentVar[1] * 100),"% variance")) +
    ylab(paste0("PC2: ",round(percentVar[2] * 100),"% variance")) +
    theme(
        strip.text = element_text(
        colour = "grey10",
        size = rel(0.8),
        margin = margin(0.5 * 7, 0.5 * 7, 0.5 * 7, 0.5 * 7)
        )) +
```

```r
  # lemon::facet_rep_wrap(~ Age, scales = "free", ncol = 1) +
  cowplot::panel_border()
```

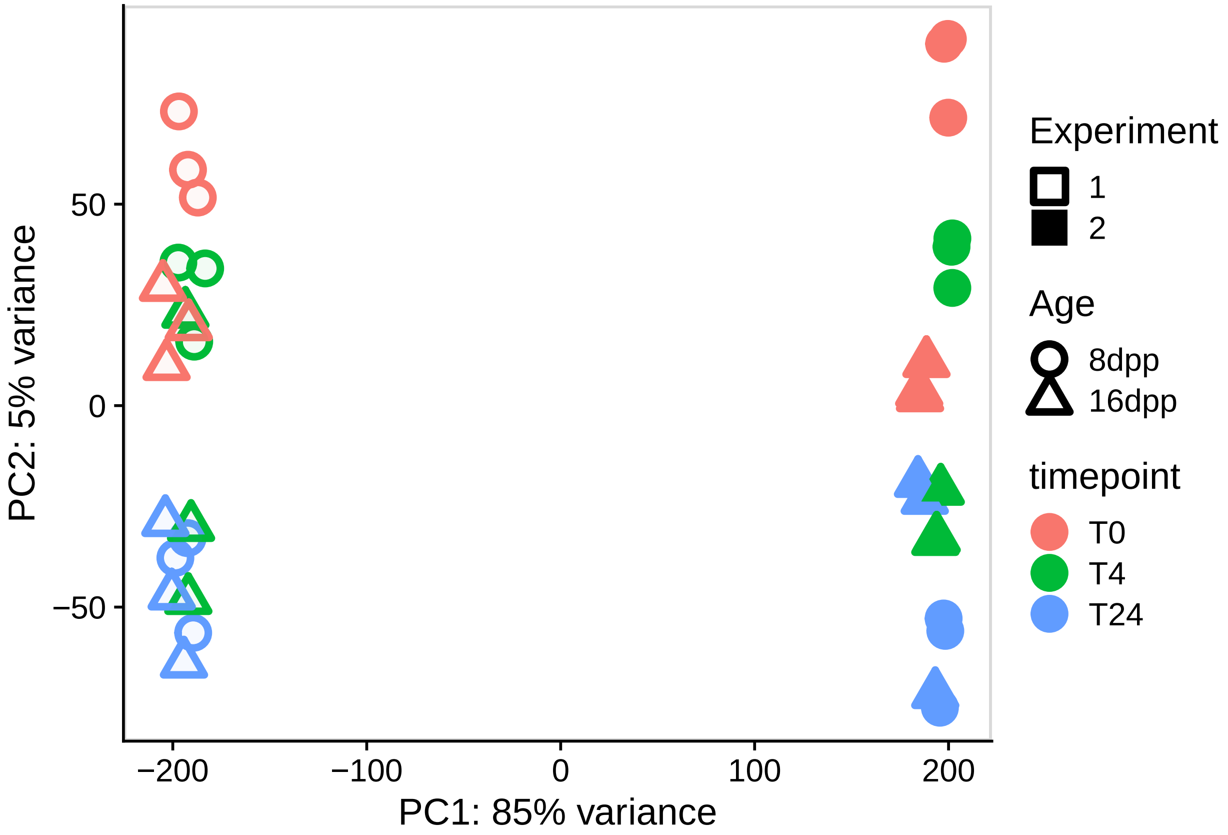


Correct for batch effects:

```r
bothAges_BatchEff <- limma::removeBatchEffect(x = bothAges,
                                              batch = c(rep(c("1", "2"), each = 9),
                                                        rep(c("1", "2"), each = 9))
                                              )

pcaBoth_BatchEff <- prcomp(t(bothAges_BatchEff))

percentVar <- pcaBoth_BatchEff$sdev^2 / sum( pcaBoth_BatchEff$sdev^2 )

as.data.frame(prcomp(t(bothAges_BatchEff))$x) %>%
  mutate(sample = rownames(.),
         timepoint = fct_relevel(str_extract(sample, pattern = "T[0-9]*"), "T24", after = Inf),
         exp = c(rep(c("1", "2"), each = 9), rep(c("1", "2"), each = 9)),
         age = rep(c("8dpp", "16dpp"), each = 18)
           ) %>%
    mutate(Age = fct_relevel(age, "8dpp")) %>%
  ggplot(aes(x = PC1, y = PC2, color = timepoint, shape = Age)) +
    geom_point(size = 4, stroke = 2) +
    geom_point(aes(fill = timepoint, shape = Age, alpha = exp), size = 5, stroke = 0) +
    cowplot::theme_cowplot(font_size = 14) +
    scale_shape_manual(values = c(21, 24, 21, 24)) +
```

```
    scale_alpha_manual(values = c(0.05, 1, 0.05, 1)) +
    guides(alpha = guide_legend(title = "Experiment",
                                override.aes = list(shape = c(22, 15), stroke = 2, alpha = 1))) +
    xlab(paste0("PC1: ",round(percentVar[1] * 100),"% variance")) +
    ylab(paste0("PC2: ",round(percentVar[2] * 100),"% variance")) +
    theme(
        strip.text = element_text(
        colour = "grey10",
        size = rel(0.8),
        margin = margin(0.5 * 7, 0.5 * 7, 0.5 * 7, 0.5 * 7)
        )) +
    # lemon::facet_rep_wrap(~ Age, scales = "free", ncol = 1) +
    cowplot::panel_border()
```

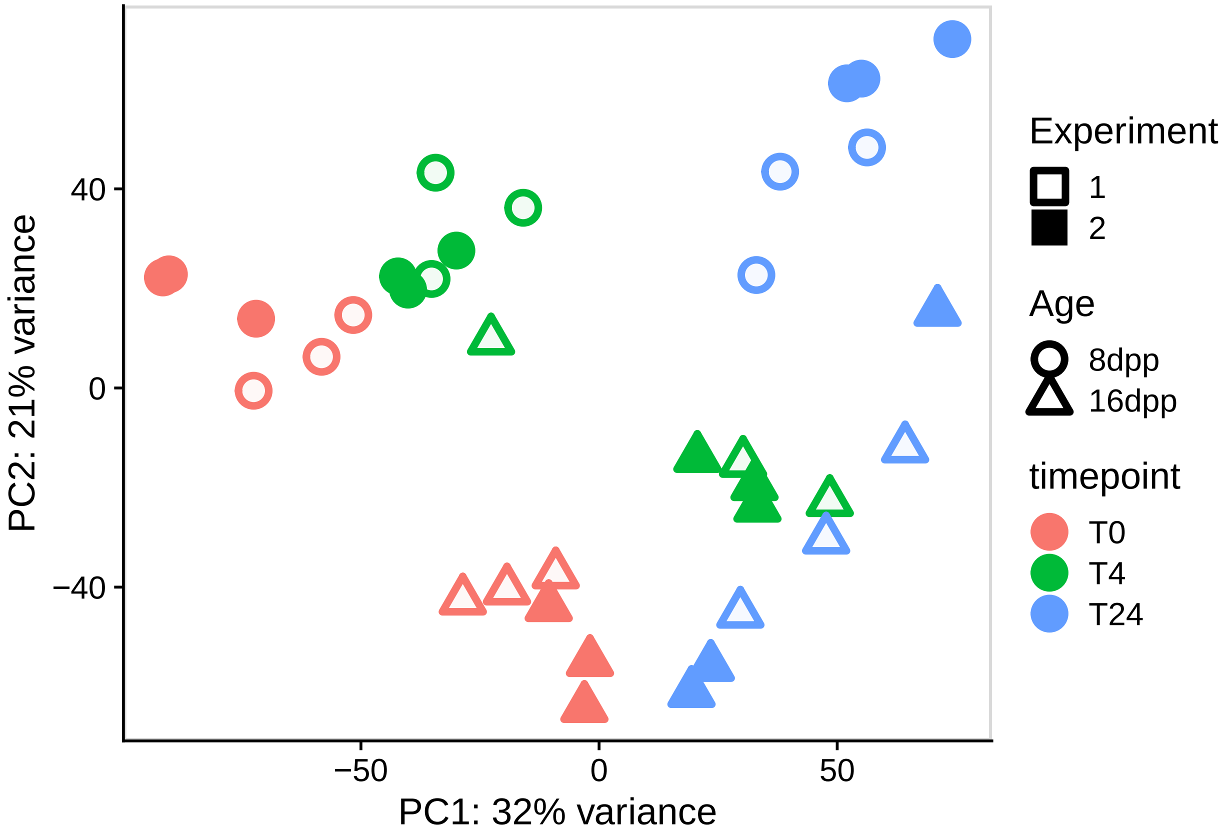


##### Weighted gene co-expression analysis

We then separate the samples in to the relevant networks

```
select_union <- rownames(vst) ### ALL GENES ###

topVar_16dpp <- assay(vst)[select_union, vst$treatment == "Inoc" & vst$age == "16dpp"]

topVar_8dpp <- assay(vst)[select_union, vst$treatment == "Inoc" & vst$age == "8dpp"]


infection8dpp <- t(topVar_8dpp)
```

```r
infection16dpp <- t(topVar_16dpp)

control16dpp <- t(assay(vst)[, vst$treatment == "Cont" & vst$age == "16dpp"])
control8dpp <- t(assay(vst)[, vst$treatment == "Cont" & vst$age == "8dpp"])
```

Checking and filtering genes and samples based on WGCNA guidlines

```r
gsg <- goodSamplesGenes(infection16dpp, verbose = 3)
```

```
##  Flagging genes and samples with too many missing values...
##   ..step 1
##   ..Excluding 1 genes from the calculation due to too many missing samples or zero variance.
##   ..step 2
```

```r
gsg$allOK
```

```
## [1] FALSE
```

```r
if (!gsg$allOK)
{
# Optionally, print the gene and sample names that were removed:
if (sum(!gsg$goodGenes) > 0)
printFlush(paste("Removing genes:",
                 paste(colnames(infection16dpp)[!gsg$goodGenes], collapse = ", ")))

if (sum(!gsg$goodSamples) > 0)
printFlush(paste("Removing samples:",
                 paste(rownames(infection16dpp)[!gsg$goodSamples], collapse = ", ")))

# Remove the offending genes and samples from the data:
infection16dpp <- infection16dpp[gsg$goodSamples, gsg$goodGenes]
}
```

```
## Removing genes: Csa3G141830
```

```r
gsg <- goodSamplesGenes(infection8dpp, verbose = 3)
```

```
##  Flagging genes and samples with too many missing values...
##   ..step 1
```

```r
gsg$allOK
```

```
## [1] TRUE
```

```r
if (!gsg$allOK)
{
# Optionally, print the gene and sample names that were removed:
if (sum(!gsg$goodGenes) > 0)
printFlush(paste("Removing genes:",
                 paste(colnames(infection8dpp)[!gsg$goodGenes], collapse = ", ")))

if (sum(!gsg$goodSamples) > 0)
printFlush(paste("Removing samples:",
                 paste(rownames(infection8dpp)[!gsg$goodSamples], collapse = ", ")))

# Remove the offending genes and samples from the data:
infection8dpp <- infection8dpp[gsg$goodSamples, gsg$goodGenes]
}
```

```
if (!identical(colnames(infection16dpp), colnames(infection8dpp))) {
  message("Genes not identical")
  message(paste("Removing genes from 16dpp:",
                paste(colnames(infection16dpp)[!colnames(infection16dpp) %in% colnames(infection8dpp)],
                      collapse = ", ")))
  infection16dpp <- infection16dpp[, colnames(infection16dpp) %in% colnames(infection8dpp)]
  message(paste("Removing genes from 8dpp:",
                paste(colnames(infection16dpp)[!colnames(infection8dpp) %in% colnames(infection16dpp)],
                      collapse = ", ")))
  infection8dpp <- infection8dpp[, colnames(infection8dpp) %in% colnames(infection16dpp)]
} else {
message("Treatments have same genes")
}
```

Plotting sample dendrograms to identify if there are outlier samples that act as outgroups to the whole set.

```
## Plot the sample dendros
sampleTree16 <- hclust(dist(infection16dpp), method = "average")
sampleTree8 <- hclust(dist(infection8dpp), method = "average")
```

Looks like all are good to include in the analysis

```
par(cex = 0.6);
par(mar = c(0,4,2,0))
plot(sampleTree16, main = "Sample clustering to detect outliers 16dpp", sub="", xlab="", cex.lab = 1.5,
     cex.axis = 1.5, cex.main = 2)
```

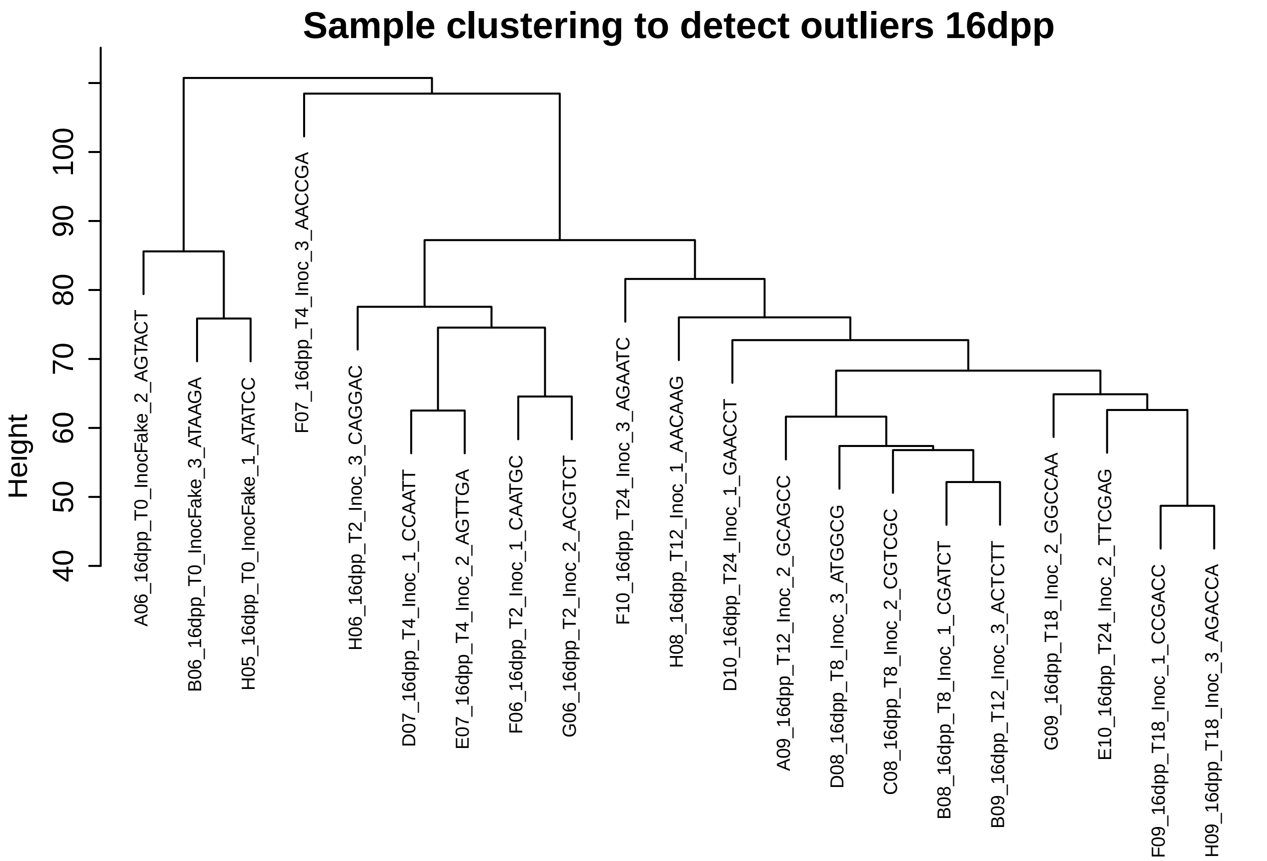

```r
plot(sampleTree8, main = "Sample clustering to detect outliers 8dpp", sub="", xlab="", cex.lab = 1.5,
     cex.axis = 1.5, cex.main = 2)
```

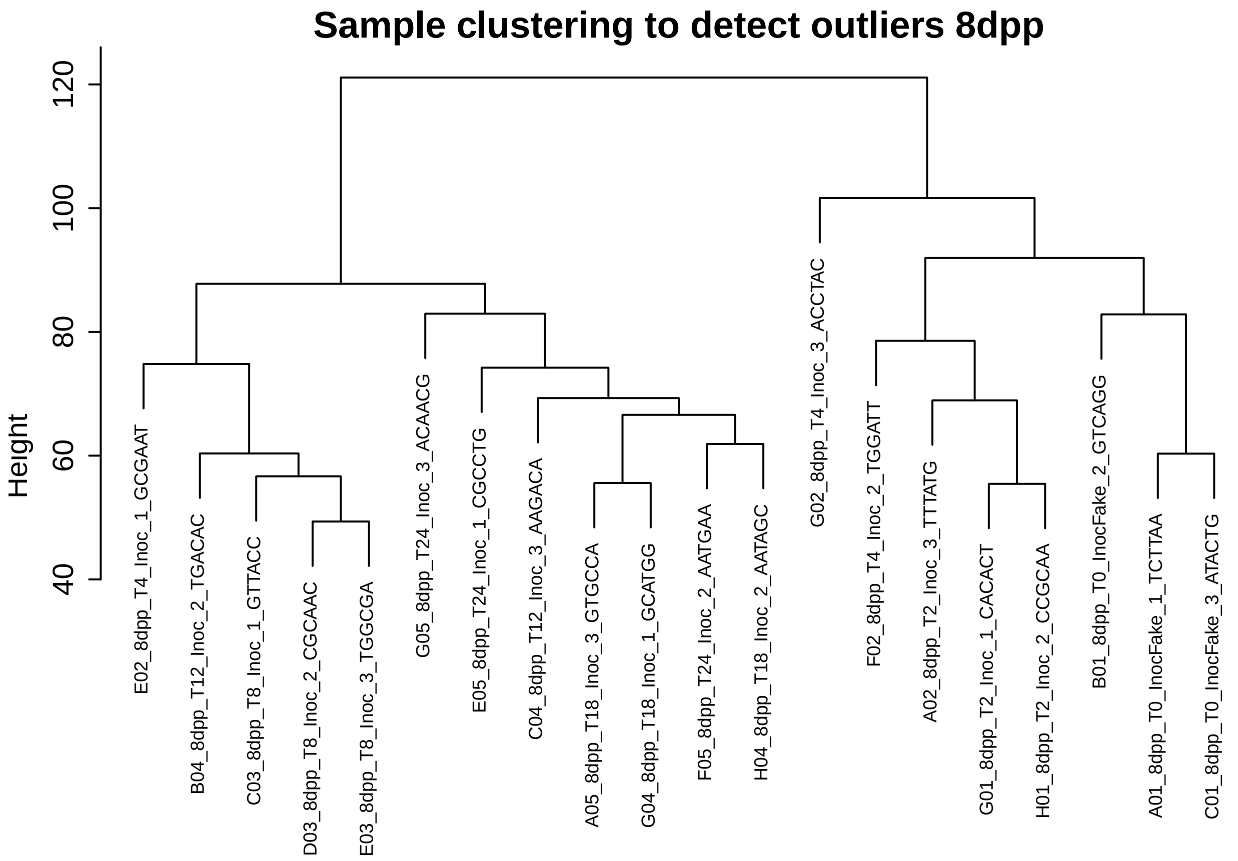


Calculate scale free topology to try to choose the best power for adjacency calculation We are using signed networks and the bicor correlation function.

```r
# Choose a set of soft-thresholding powers
powers <- c(c(1:10), seq(from = 12, to=30, by=2))
# Call the network topology analysis function
sft16 <-
  pickSoftThreshold(
    infection16dpp,
    powerVector = powers,
    verbose = 0,
    networkType = "signed",
    corFnc = "bicor",
    corOptions = "maxPOutliers = 0.1"
  )

sft8 <-
  pickSoftThreshold(
  infection8dpp,
  powerVector = powers,
  verbose = 0,
  networkType = "signed",
  corFnc = "bicor",
```

```
  corOptions = "maxPOutliers = 0.1"
  )

bind_rows("16dpp" = sft16$fitIndices, "8dpp" = sft8$fitIndices, .id = "network") %>%
ggplot() +
  geom_text(aes(x = Power, y = -sign(slope) * SFT.R.sq, label = Power, color = network)) +
  geom_abline(intercept = 0.8, slope = 0, color = "red") +
  ylab("Scale Free Topology Model Fit,signed R^2")
```

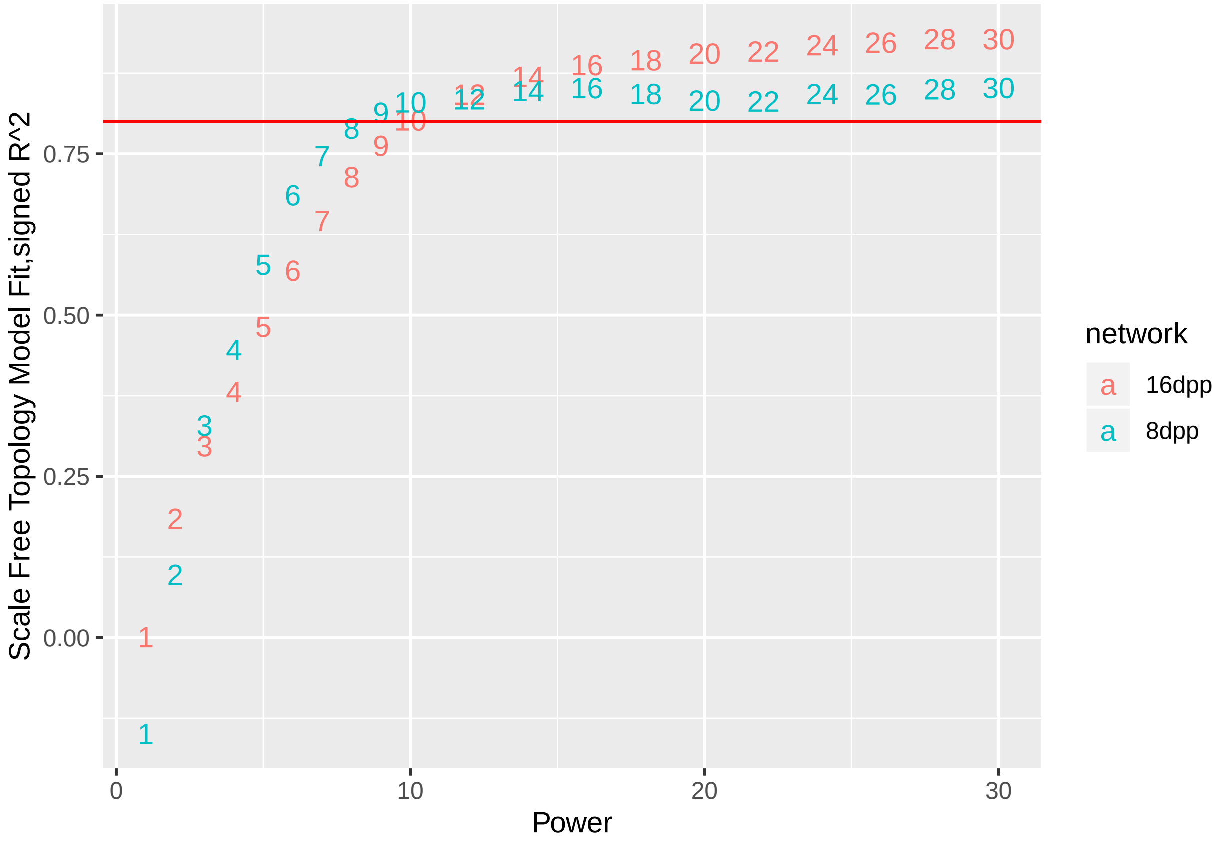


We choose 12 as a power because the both networks reach a scale free topology R^2 fit of greater than 0.8

```
# calculate adjacency matrix using bicor
softPower <- 12;
adjacency16 <-
  adjacency(
  infection16dpp,
  power = softPower,
  type = "signed",
  corFnc = "bicor",
  corOptions = "maxPOutliers = 0.1"
  )
```

```
## alpha: 1.000000
```

```
adjacency8 <-
  adjacency(
  infection8dpp,
```

```
  power = softPower,
  type = "signed",
  corFnc = "bicor",
  corOptions = "maxPOutliers = 0.1"
  )
```

```
## alpha: 1.000000
```

We then take the adjacency matrix and calculate the Topological Overlap Matrix

```
# TOMType doesn't matter in signed networks
dissTOM16 <- 1 - TOMsimilarity(adjacency16, TOMType = "unsigned")
```

```
## ..connectivity..
## ..matrix multiplication (system BLAS)..
## ..normalization..
## ..done.
```

```
dissTOM8 <- 1 - TOMsimilarity(adjacency8, TOMType = "unsigned")
```

```
## ..connectivity..
## ..matrix multiplication (system BLAS)..
## ..normalization..
## ..done.
```

Plot the gene dendrograms based on their dissimilarty

```
geneTree16 <- hclust(as.dist(dissTOM16), method = "average");
geneTree8 <- hclust(as.dist(dissTOM8), method = "average");

plot(geneTree16, xlab = "", sub = "", main = "Gene clustering on TOM-based dissimilarity 16dpp",
     labels = FALSE, hang = 0.04);
```

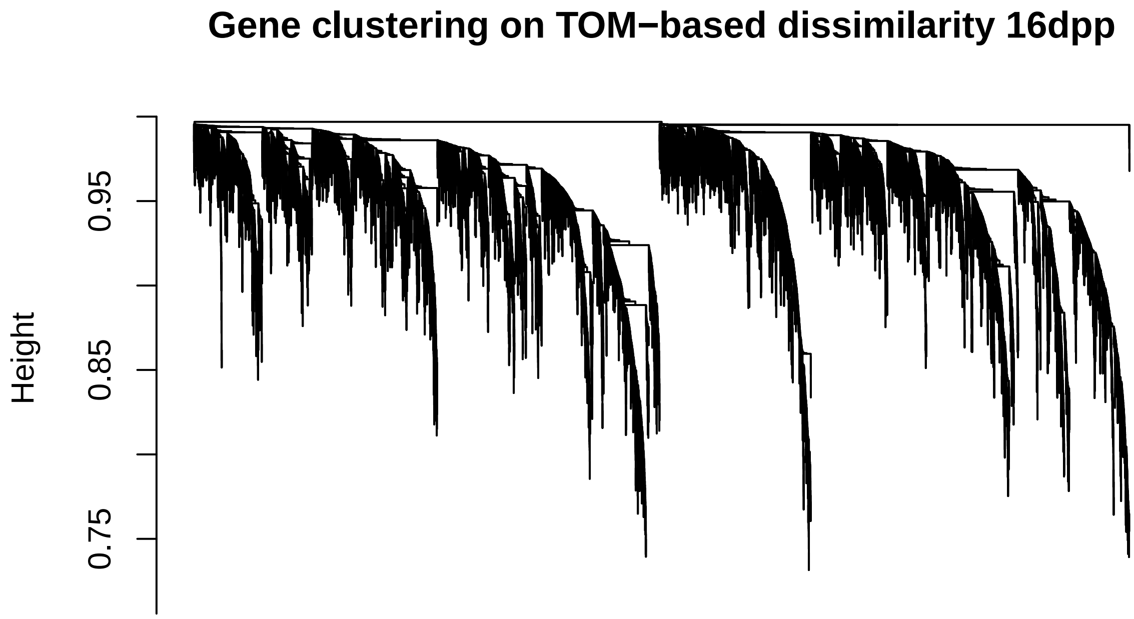


```r
plot(geneTree8, xlab = "", sub = "", main = "Gene clustering on TOM-based dissimilarity 8dpp",
     labels = FALSE, hang = 0.04);
```

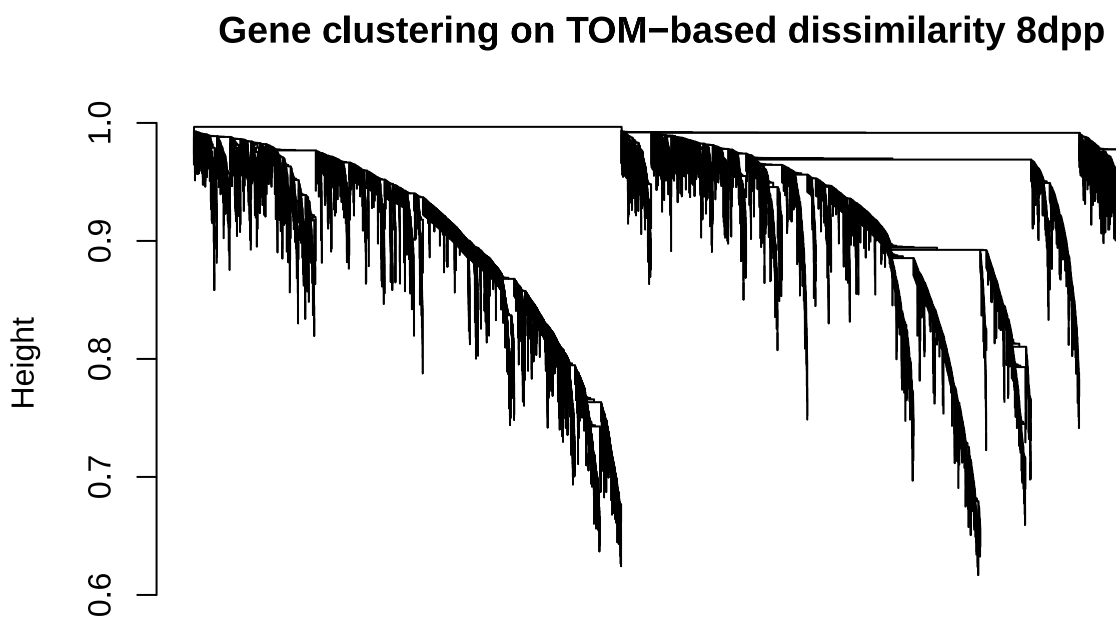


We then use the dynamic tree cut algorithm to identify first pass modules. These will then be merged.

```
minModuleSize <- 30;
# Module identification using dynamic tree cut:
dynamicMods16 <- cutreeDynamic(dendro = geneTree16,
                              distM = dissTOM16,
                              deepSplit = 2,
                              pamRespectsDendro = FALSE,
                              minClusterSize = minModuleSize);
```

```
##  ..cutHeight not given, setting it to 0.995  ===>  99% of the (truncated) height range in dendro.
##  ..done.
```

```
dynamicMods8 <- cutreeDynamic(dendro = geneTree8,
                              distM = dissTOM8,
                              deepSplit = 2,
                              pamRespectsDendro = FALSE,
                              minClusterSize = minModuleSize);
```

```
##  ..cutHeight not given, setting it to 0.994  ===>  99% of the (truncated) height range in dendro.
##  ..done.
```

```
# Convert numeric lables into colors
dynamicColors16 <- labels2colors(dynamicMods16)
table(dynamicColors16)
```

```
## dynamicColors16
##   antiquewhite4         bisque4           black            blue           brown
##              57              94             614            1444             797
```

```
##           brown4           coral1           coral2             cyan        darkgreen
##               94               59               55              334              208
##         darkgrey      darkmagenta   darkolivegreen       darkorange      darkorange2
##              181              133              134              170               96
##          darkred    darkseagreen4    darkslateblue    darkturquoise      floralwhite
##              208               63               93              182               98
##            green      greenyellow           grey60        honeydew1            ivory
##              671              404              262               72              102
##   lavenderblush3        lightcyan       lightcyan1       lightgreen       lightpink4
##               73              270              103              241               75
## lightsteelblue1      lightyellow          magenta           maroon     mediumorchid
##              104              227              527               75               54
##    mediumpurple3     midnightblue      navajowhite2           orange       orangered4
##              104              276               75              178              111
##    paleturquoise  palevioletred3             pink            plum1            plum2
##              152               78              570              117               92
##           purple              red        royalblue      saddlebrown           salmon
##              451              653              227              157              351
##          salmon4          sienna3          skyblue         skyblue1         skyblue2
##               84              128              162               46               53
##         skyblue3        steelblue              tan         thistle1         thistle2
##              119              154              370               91               91
##        turquoise           violet            white           yellow          yellow4
##             1746              136              163              752               52
##      yellowgreen
##              126
```

```
dynamicColors8 <- labels2colors(dynamicMods8)
table(dynamicColors8)
```

```
## dynamicColors8
##          black           blue          brown           cyan      darkgreen
##            633           2381           2152            219            137
##       darkgrey     darkorange        darkred  darkturquoise          green
##            119             96            150            131           1147
##    greenyellow         grey60      lightcyan     lightgreen    lightyellow
##            383            185            210            182            159
##        magenta   midnightblue         orange  paleturquoise           pink
##            451            218            115             70            542
##         purple            red      royalblue    saddlebrown         salmon
##            445            890            151             86            233
##        skyblue      steelblue            tan      turquoise         violet
##             94             86            364           2608             48
##          white         yellow
##             94           1460
```

ID first module eigengenes and then cluster based on dissimilarity

```
## calculate eigengenes

MEList16 <- moduleEigengenes(infection16dpp, colors = dynamicColors16)
MEs16 <- MEList16$eigengenes

MEList8 <- moduleEigengenes(infection8dpp, colors = dynamicColors8)
MEs8 <- MEList8$eigengenes
```

```
# Calculate dissimilarity of module eigengenes
MEDiss16 <- 1-cor(MEs16)
MEDiss8 <- 1-cor(MEs8)
```

Let's merge at three different thresholds

```
# Cluster module eigengenes
METree16 <- hclust(as.dist(MEDiss16), method = "average");
# Plot the result

par(cex = 0.6);
par(mar = c(0,4,2,0))
plot(METree16, main = "Clustering of module eigengenes",
     xlab = "", sub = "")
abline(h=0.15, col = "red")
abline(h=0.20, col = "blue")
abline(h=0.25, col = "green")
abline(h=0.30, col = "purple")
```

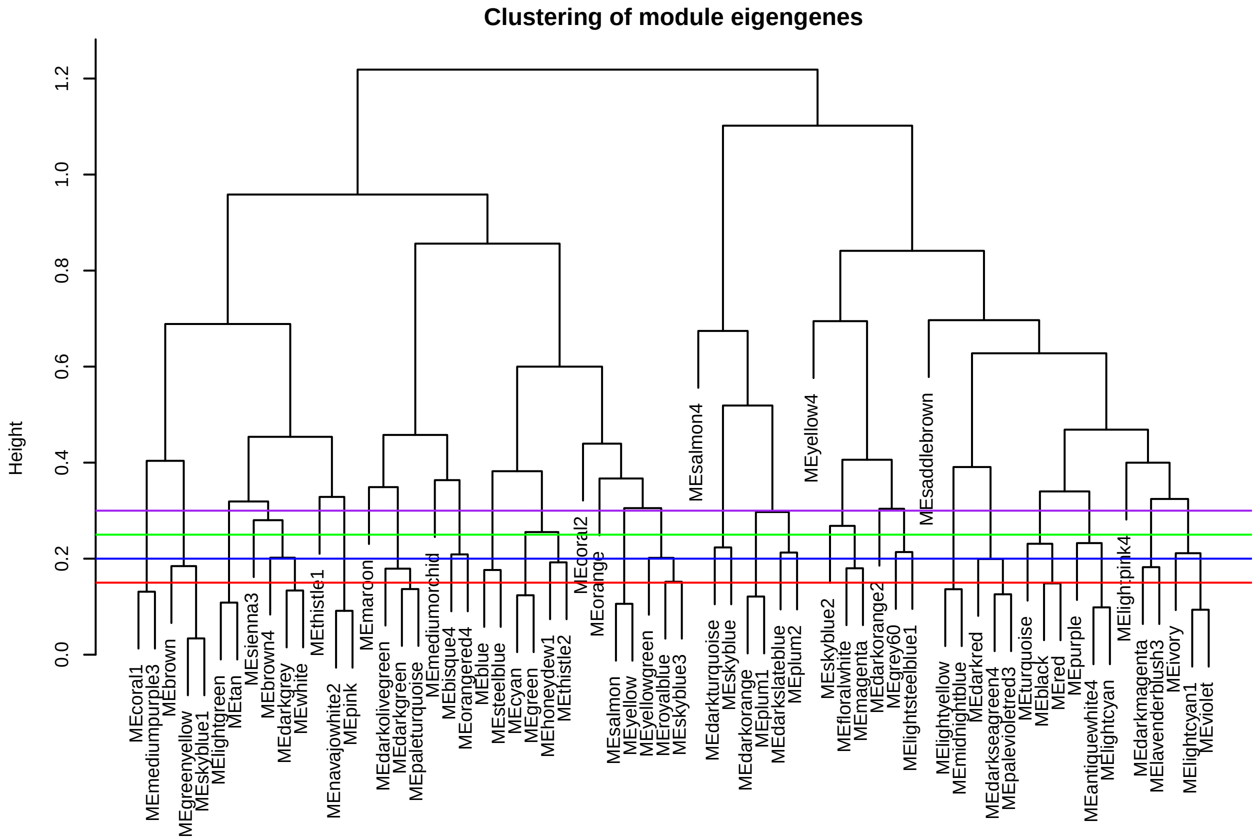


```
## find a good merge

MEDissThres <- 0.15
# Call an automatic merging function
merge16.15 <-
mergeCloseModules(infection16dpp,
dynamicColors16,
cutHeight = MEDissThres,
```

```r
verbose = 0)
# The merged module colors
merged16Colors.15 <- merge16.15$colors

# Eigengenes of the new merged modules:
merged16MEs.15 <- merge16.15$newMEs


MEDissThres <- 0.20
# Call an automatic merging function
merge16.20 <-
mergeCloseModules(infection16dpp,
dynamicColors16,
cutHeight = MEDissThres,
verbose = 0)
# The merged module colors
merged16Colors.20 <- merge16.20$colors

# Eigengenes of the new merged modules:
merged16MEs.20 <- merge16.20$newMEs

MEDissThres <- 0.25
# Call an automatic merging function
merge16.25 <-
mergeCloseModules(infection16dpp,
dynamicColors16,
cutHeight = MEDissThres,
verbose = 0)
# The merged module colors
merged16Colors.25 <- merge16.25$colors

# Eigengenes of the new merged modules:
merged16MEs.25 <- merge16.25$newMEs

MEDissThres <- 0.30
# Call an automatic merging function
merge16.30 <-
mergeCloseModules(infection16dpp,
dynamicColors16,
cutHeight = MEDissThres,
verbose = 0)
# The merged module colors
merged16Colors.30 <- merge16.30$colors

# Eigengenes of the new merged modules:
merged16MEs.30 <- merge16.30$newMEs
```

And the same for the 8dpp network

```r
# Cluster module eigengenes
METree8 <- hclust(as.dist(MEDiss8), method = "average");
# Plot the result

par(cex = 0.6);
```

```r
par(mar = c(0,4,2,0))
plot(METree8, main = "Clustering of module eigengenes",
     xlab = "", sub = "")
abline(h=0.15, col = "red")
abline(h=0.20, col = "blue")
abline(h=0.25, col = "green")
abline(h=0.30, col = "purple")
```

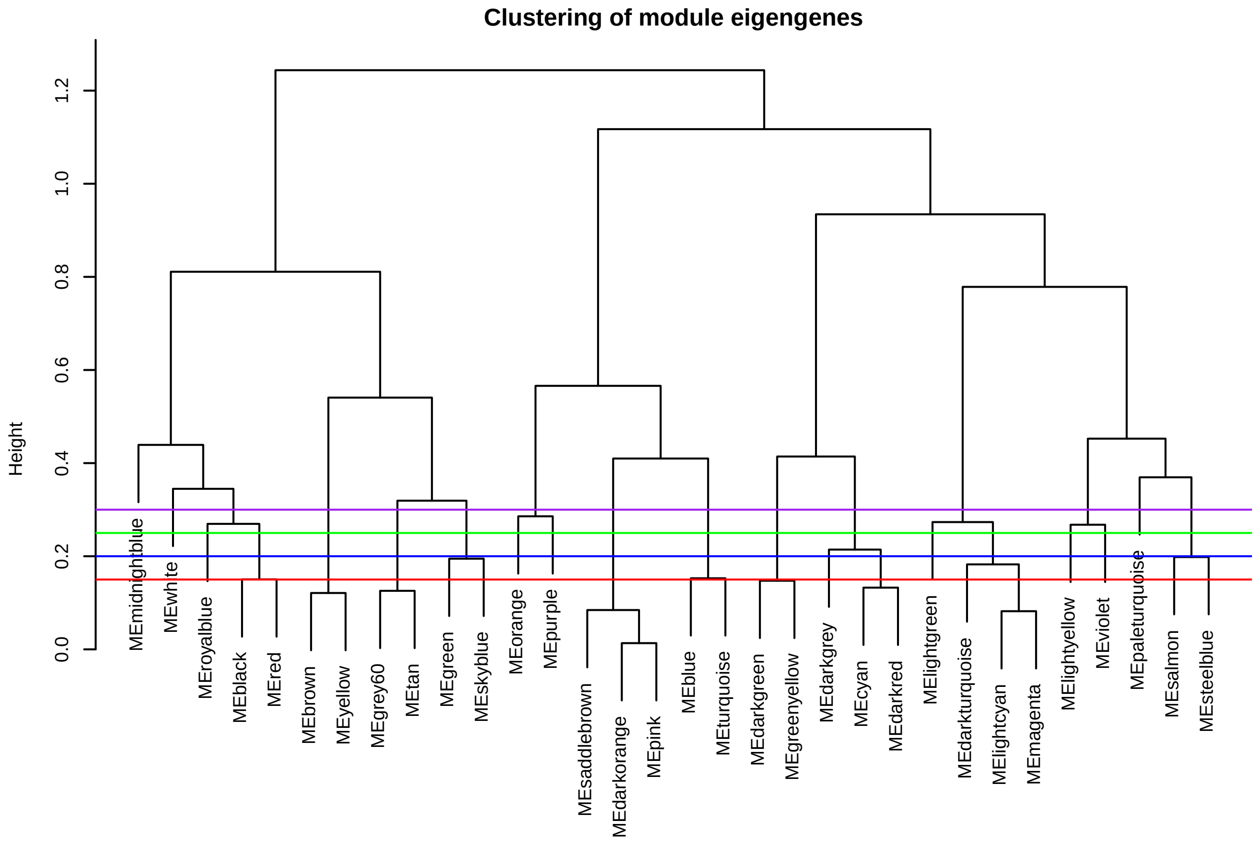


```r
## find a good merge

MEDissThres <- 0.15
# Call an automatic merging function
merge8.15 <-
mergeCloseModules(infection8dpp,
dynamicColors8,
cutHeight = MEDissThres,
verbose = 0)
# The merged module colors
merged8Colors.15 <- merge8.15$colors

# Eigengenes of the new merged modules:
merged8MEs.15 <- merge8.15$newMEs


MEDissThres <- 0.20
# Call an automatic merging function
```

```r
merge8.20 <-
mergeCloseModules(infection8dpp,
dynamicColors8,
cutHeight = MEDissThres,
verbose = 0)
# The merged module colors
merged8Colors.20 <- merge8.20$colors

# Eigengenes of the new merged modules:
merged8MEs.20 <- merge8.20$newMEs

MEDissThres <- 0.25
# Call an automatic merging function
merge8.25 <-
mergeCloseModules(infection8dpp,
dynamicColors8,
cutHeight = MEDissThres,
verbose = 0)
# The merged module colors
merged8Colors.25 <- merge8.25$colors

# Eigengenes of the new merged modules:
merged8MEs.25 <- merge8.25$newMEs

MEDissThres <- 0.30
# Call an automatic merging function
merge8.30 <-
mergeCloseModules(infection8dpp,
dynamicColors8,
cutHeight = MEDissThres,
verbose = 0)
# The merged module colors
merged8Colors.30 <- merge8.30$colors

# Eigengenes of the new merged modules:
merged8MEs.30 <- merge8.30$newMEs
```

Ok lets compare the merged modules to the orginal dynamic tree cut and the gene dendrogram

```r
plotDendroAndColors(geneTree16, cbind(dynamicColors16, merged16Colors.15, merged16Colors.20, merged16Col
                    c("Dynamic Tree Cut", "Merged 0.15", "Merged 0.20", "Merged 0.25", "Merged 0.30"),
                    dendroLabels = FALSE, hang = 0.03,
                    addGuide = TRUE, guideHang = 0.05,
                    main = "Cluster dendrogram - 16dpp")
```

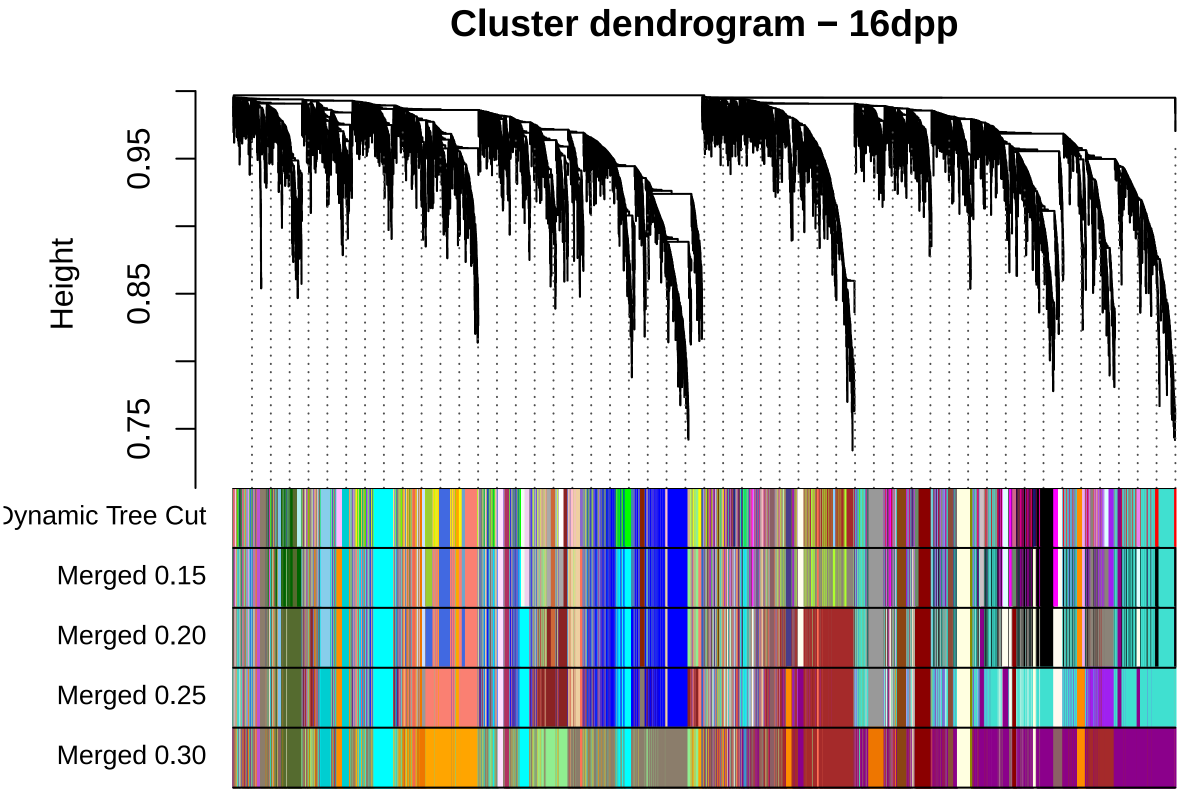


```
plotDendroAndColors(geneTree8, cbind(dynamicColors8, merged8Colors.15, merged8Colors.20, merged8Colors.2
                    c("Dynamic Tree Cut", "Merged 0.15", "Merged 0.20", "Merged 0.25", "Merged 0.30"),
                    dendroLabels = FALSE, hang = 0.03,
                    addGuide = TRUE, guideHang = 0.05,
                    main = "Cluster dendrogram - 8dpp")
```

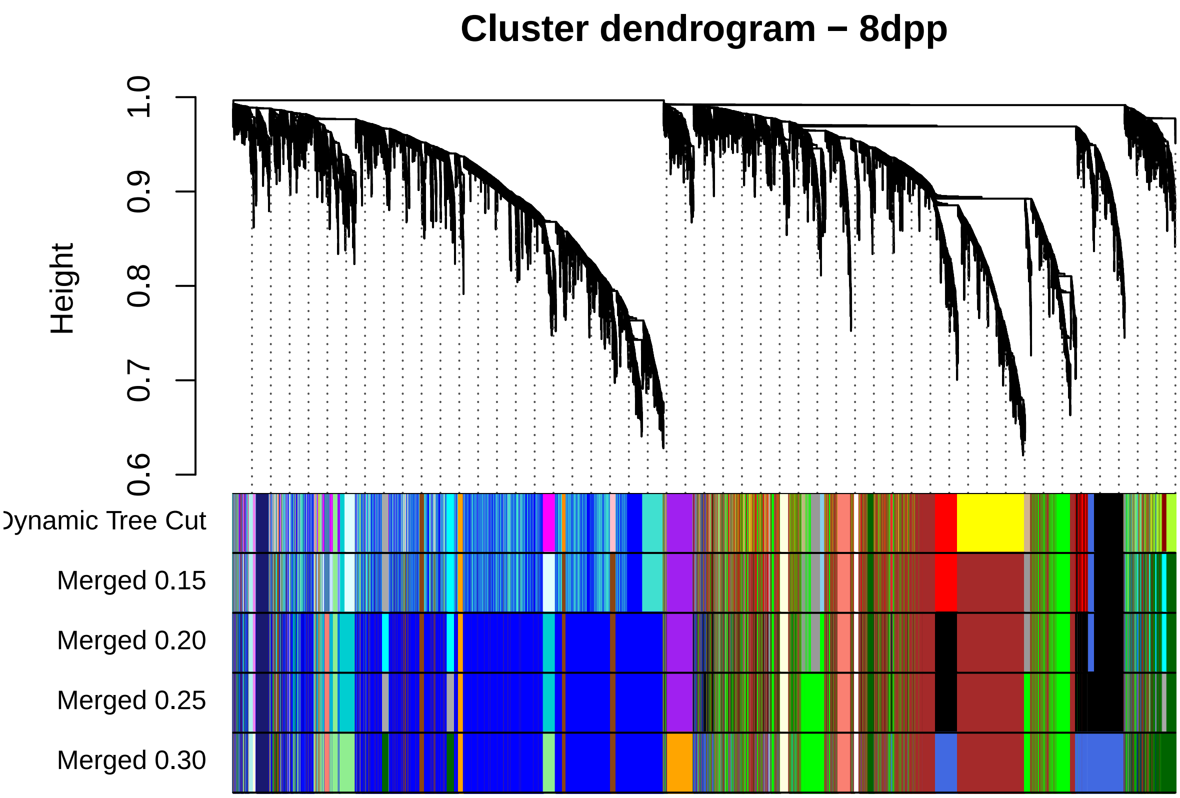


We choose the 0.25 merge level

```r
# Rename to selected merged moduleColors
moduleColors16 <- merged16Colors.25
moduleColors8 <- merged8Colors.25

# Construct numerical labels corresponding to the colors
finalColors16 <- names(sort(table(moduleColors16), decreasing = TRUE))
finalColors16 <- finalColors16[!finalColors16 %in% "grey"]

finalColors8 <- names(sort(table(moduleColors8), decreasing = TRUE))
finalColors8 <- finalColors8[!finalColors8 %in% "grey"]

colorOrder <- c("grey", finalColors16)
moduleLabels16 <- match(moduleColors16, colorOrder) - 1

colorOrder <- c("grey", finalColors8)
moduleLabels8 <- match(moduleColors8, colorOrder) - 1

table(moduleLabels16)
```

```
## moduleLabels16
##    1    2    3    4    5    6    7    8    9   10   11   12   13   14   15   16
## 3013 1598 1575 1247 1177 1168  778  645  625  547  503  494  472  366  349  344
##   17   18   19   20   21   22   23   24   25   26   27   28   29
##  205  178  163  157   96   91   84   75   75   55   54   53   52
```

```r
table(moduleLabels8)
```

```
## moduleLabels8
##    1    2    3    4    5    6    7    8    9   10   11   12   13   14   15   16
## 4989 3612 1790 1674  792  724  520  488  445  319  218  182  159  115   94   70
##   17
##   48
```

```r
#overlapTable(moduleLabels16, moduleLabels8)
```

Recolor with prettier colors

```r
qual_col_pals <- brewer.pal.info[brewer.pal.info$category == 'qual',]

col_vector <- unique(unlist(mapply(brewer.pal, qual_col_pals$maxcolors, rownames(qual_col_pals))))

# recolor modules so there are no overlaps between the network colors
RNGkind(sample.kind = "Rounding")
set.seed(7)
customColorOrder16 <-
c(
"grey",
sample(col_vector,
       size = length(unique(moduleLabels16[!moduleLabels16 %in% 0])),
       replace = FALSE
       )
)
moduleColors16.custom <- customColorOrder16[moduleLabels16 + 1]

set.seed(9)
customColorOrder8 <-
c(
"grey",
sample(col_vector[!col_vector %in% unique(moduleColors16.custom)],
       size = length(unique(moduleLabels8[!moduleLabels8 %in% 0])),
       replace = FALSE)
)
moduleColors8.custom <- customColorOrder8[moduleLabels8 + 1]
```

Selected the new MEs after merging

```r
MEs16Lables <- moduleEigengenes(infection16dpp, colors = moduleLabels16)$eigengenes
MEs8Lables <- moduleEigengenes(infection8dpp, colors = moduleLabels8)$eigengenes

MEs16Colors <- moduleEigengenes(infection16dpp, colors = moduleColors16.custom)$eigengenes
MEs8Colors <- moduleEigengenes(infection8dpp, colors = moduleColors8.custom)$eigengenes
```

Plot the final gene trees and modules

```r
plotDendroAndColors(geneTree16,
                    cbind(moduleColors16.custom),
                    "Resistant\nmodules\n(16dpp)",
                    dendroLabels = FALSE, hang = 0.03,
                    addGuide = TRUE, guideHang = 0.05,
                    main = "Resistant-aged (16dpp) fruit\ngene dendrogram and module colors")
```

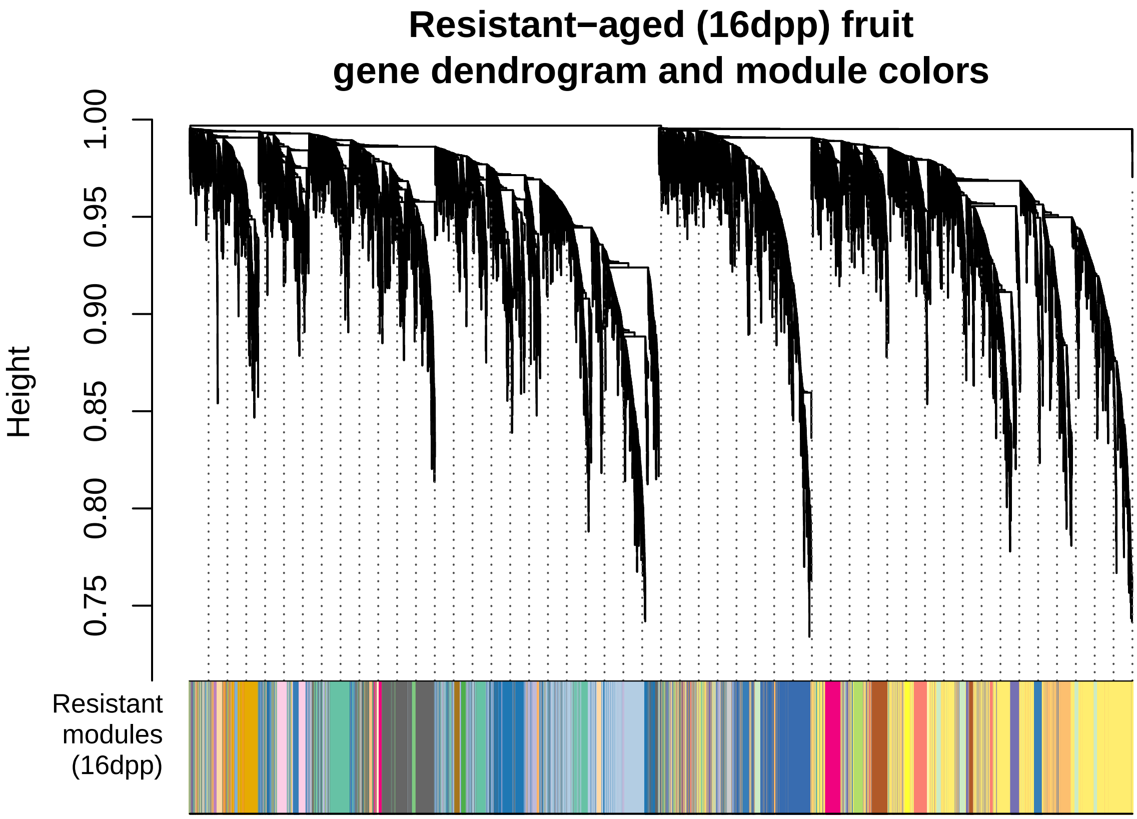


```r
plotDendroAndColors(geneTree8,
                    cbind(moduleColors8.custom),
                    "Susceptible\nmodules\n(8dpp)",
                    dendroLabels = FALSE, hang = 0.03,
                    addGuide = TRUE, guideHang = 0.05,
                    main = "Susceptible-aged (8dpp) fruit\ngene dendrogram and module colors")
```

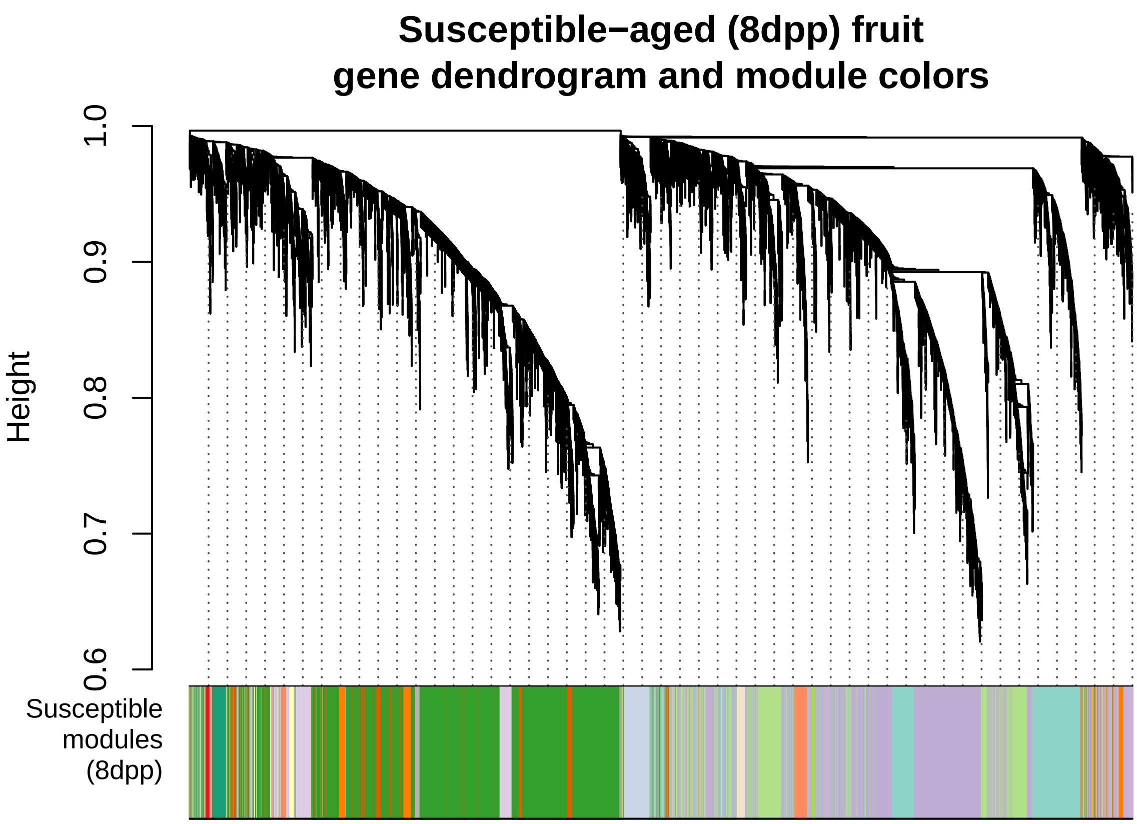


```
#
# plotDendroAndColors(geneTree8,
#                     cbind(moduleColors8.custom),
#                     "Susceptible\nmodules\n(8dpp)",
#                     dendroLabels = FALSE, hang = 0.03,
#                     addGuide = TRUE, guideHang = 0.05,
#                     main = "Susceptible-aged (8dpp) fruit\ngene dendrogram and module colors")

pdf(file = "fig6A.pdf", width = 6.5, height = 4.5)
    plotDendroAndColors(geneTree8,
                    cbind(moduleColors8.custom),
                    "Susceptible\nmodules\n(8dpp)",
                    dendroLabels = FALSE, hang = 0.03,
                    addGuide = TRUE, guideHang = 0.05,
                    main = "Susceptible-aged (8dpp) fruit\ngene dendrogram and module colors")
dev.off()
```

```
## pdf
##   2
```

```
pdf(file = "fig6B.pdf", width = 6.5, height = 4.5)
    plotDendroAndColors(geneTree16,
                    cbind(moduleColors16.custom),
                    "Resistant\nmodules\n(16dpp)",
                    dendroLabels = FALSE, hang = 0.03,
                    addGuide = TRUE, guideHang = 0.05,
                    main = "Resistant-aged (16dpp) fruit\ngene dendrogram and module colors")
```

```
dev.off()
```

```
## pdf
##   2
```

Let's examine the module preservation. This step is really long computationally and extremely verbose so this is not run here to save time and space in the document.

```
# Number of data sets that we work with
nSets <- 2;
# Object that will contain the expression data
multiExpr <- list();
multiExpr[[1]] = list(data = infection16dpp)
multiExpr[[2]] = list(data = infection8dpp)

# Names for the two sets
setLabels <- c("16dpp", "8dpp");
# Important: components of multiExpr must carry identificating names
names(multiExpr) <- setLabels
# Display the dimensions of the expression data (if you are confused by this construct, ignore it):
lapply(multiExpr, lapply, dim)


# Create an object (list) holding the module labels for each set:
colorList <- list(moduleColors16.custom, moduleColors8.custom);
# Components of the list must be named so that the names can be matched to the names of multiExpr
names(colorList) <- setLabels;

mp_infected <-
  modulePreservation(
    multiExpr,
    colorList,
    networkType = "signed",
    corFnc = "bicor",
    corOptions = "maxPOutliers = 0.1",
    referenceNetworks = c(1:2),
    loadPermutedStatistics = FALSE,
    nPermutations = 200,
    verbose = 3
  )
```

Preservation stats for 16dpp network:

```
presData16dppRef <- tibble(moduleColor = rownames(mp_infected$preservation$Z[[1]][[2]]),
                           Zsummary = mp_infected$preservation$Z[[1]][[2]]$Zsummary.pres,
                           medianRank = mp_infected$preservation$observed[[1]][[2]]$medianRank.pres,
                           moduleSize = mp_infected$preservation$Z[[1]][[2]]$moduleSize) %>%
  filter(!moduleColor == "gold") %>%
  mutate(moduleLabel = match(moduleColor, names(sort(table(moduleColors16.custom), decreasing = TRUE))))

pr1_16 <- presData16dppRef %>%
  ggplot(aes(x = moduleSize, y = Zsummary)) +
  geom_point(aes(color = moduleColor), size = 4) +
  ggrepel::geom_text_repel(aes(label = moduleLabel), force = 5) +
  geom_hline(yintercept = 2, linetype = 2, color = "red") +
  geom_hline(yintercept = 10, linetype = 2, color = "darkgreen") +
```

```r
  scale_color_identity()

pr2_16 <- presData16dppRef %>%
  ggplot(aes(x = moduleSize, y = medianRank)) +
  geom_point(aes(color = moduleColor), size = 4) +
  ggrepel::geom_text_repel(aes(label = moduleLabel), force = 5) +
  scale_color_identity()

pr3_16 <- presData16dppRef %>%
  ggplot(aes(x = Zsummary, y = medianRank)) +
  geom_point(aes(color = moduleColor), size = 4) +
  ggrepel::geom_text_repel(aes(label = moduleLabel), force = 5) +
  scale_color_identity()
```

Now presrevation of the 8dpp network:

```r
presData8dppRef <- tibble(moduleColor = rownames(mp_infected$preservation$Z[[2]][[1]]),
                          Zsummary = mp_infected$preservation$Z[[2]][[1]]$Zsummary.pres,
                          medianRank = mp_infected$preservation$observed[[2]][[1]]$medianRank.pres,
                          moduleSize = mp_infected$preservation$Z[[2]][[1]]$moduleSize) %>%
  filter(!moduleColor == "gold") %>%
  mutate(moduleLabel = match(moduleColor, names(sort(table(moduleColors8.custom), decreasing = TRUE))))

pr1_8 <- presData8dppRef %>%
  ggplot(aes(x = moduleSize, y = Zsummary)) +
  geom_point(aes(color = moduleColor), size = 4) +
  ggrepel::geom_text_repel(aes(label = moduleLabel), force = 5) +
  geom_hline(yintercept = 2, linetype = 2, color = "red") +
  geom_hline(yintercept = 10, linetype = 2, color = "darkgreen") +
  scale_color_identity()

pr2_8 <- presData8dppRef %>%
  ggplot(aes(x = moduleSize, y = medianRank)) +
  geom_point(aes(color = moduleColor), size = 4) +
  ggrepel::geom_text_repel(aes(label = moduleLabel), force = 5) +
  scale_color_identity()

pr3_8 <- presData8dppRef %>%
  ggplot(aes(x = Zsummary, y = medianRank)) +
  geom_point(aes(color = moduleColor), size = 4) +
  ggrepel::geom_text_repel(aes(label = moduleLabel), force = 5) +
  scale_color_identity()
```

```r
cowplot::plot_grid(pr1_8, pr1_16, pr2_8, pr2_16, pr3_8, pr3_16, ncol = 2)
```

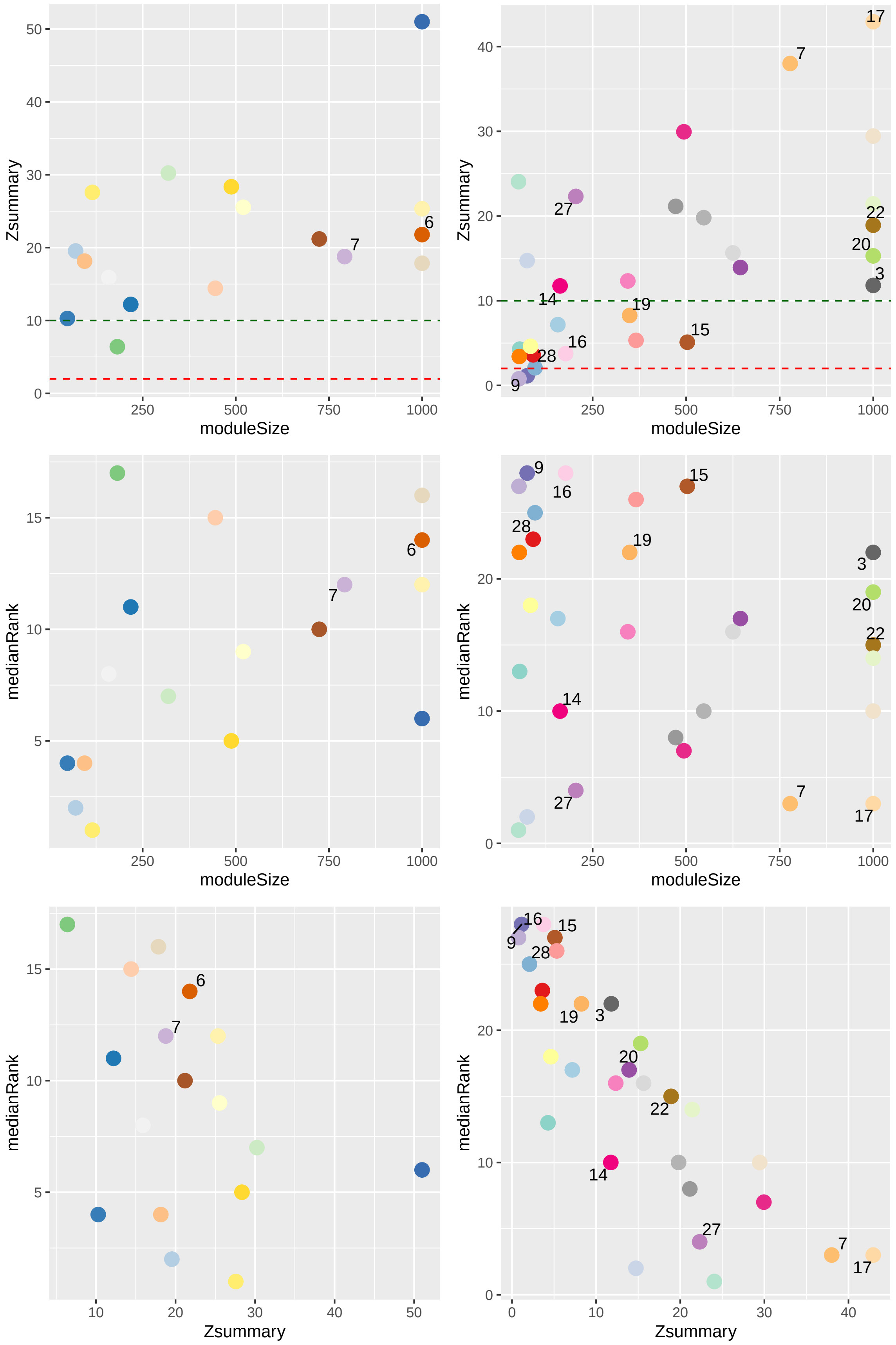
Zsummary
moduleSize
7
6
Zsummary
moduleSize
17
7
27
22
20
3
14
19
15
16
28
9
medianRank
moduleSize
6
7
medianRank
moduleSize
9
16
15
28
19
3
20
22
14
27
7
17
medianRank
Zsummary
6
7
medianRank
Zsummary
16
15
9
28
19
3
20
22
14
27
7
17

Plot alluvial comparison of modules

```
networkLodes <- as.data.frame(overlapTable(moduleLabels16, moduleLabels8)$countTable) %>%
  mutate(module16dpp = rownames(.)) %>%
  gather(key = "module8dpp", value = "overlap", -module16dpp) %>%
  left_join(as.data.frame(overlapTable(moduleLabels16, moduleLabels8)$pTable) %>%
              mutate(module16dpp = rownames(.)) %>%
              gather(key = "module8dpp", value = "pvalue", -module16dpp)
  ) %>%
  mutate(
    # module16dpp = fct_relevel(module16dpp, as.character(c(1:23, 0))),
    # module8dpp = fct_relevel(module8dpp, as.character(1:14)),
    `16dpp` = fct_rev(fct_inorder(rep(customColorOrder16[-1], length(customColorOrder8[-1])))),
    `8dpp` = fct_rev(fct_inorder(rep(customColorOrder8[-1], each = length(customColorOrder16[-1]))))
  ) %>%
  to_lodes_form(key = "Network", value = "module", id = "Cohort", -pvalue, -overlap, -module16dpp, -modu
  mutate(padj = p.adjust(pvalue),
         isSig = ifelse(padj < 0.001, "Yes", "No"),
         moduleSize = c(
           rep(sort(table(moduleColors16.custom), decreasing = TRUE), length(customColorOrder8[-1])),
           rep(sort(table(moduleColors8.custom), decreasing = TRUE), each = length(customColorOrder16[-
         ),
         moduleLabel = ifelse(Network == "16dpp", as.character(module16dpp), as.character(module8dpp))
         )


sixC <- ggplot(data = networkLodes,
         aes(
           x = Network,
           y = overlap,
           stratum = module,
           alluvium = Cohort
         )) +
    geom_stratum(aes(fill = module, color = module)) +
    scale_fill_identity() +
    scale_color_identity() +
    scale_alpha_manual(values=c(0.25, 1)) +
    ggnewscale::new_scale_fill() +
    ggnewscale::new_scale_color() +
    geom_flow(aes(fill = isSig)) +
    scale_fill_viridis_d(direction = 1, begin = 0, end = 0.55,
                       name = "Significant Overlap (Fisher's Exact, FDR < 0.001)"
                      ) +
    geom_text(stat = "stratum", aes(label = ifelse(
    moduleSize > 250, as.character(moduleLabel), NA
    ))) +
    ggrepel::geom_text_repel(data = filter(networkLodes, Network == "8dpp"),
                           stat = "stratum", aes(label = ifelse(moduleSize <= 250,
                                                                moduleLabel,
                                                                NA)),
                           nudge_x = .5) +
    ggrepel::geom_text_repel(data = filter(networkLodes, Network == "16dpp"),
                           stat = "stratum", aes(label = ifelse(moduleSize <= 250,
                                                                moduleLabel,
```

```
                                                                    NA)),
                          nudge_x = -.5) +
    coord_flip() +
    scale_y_continuous(breaks = seq(0, length(infection16dpp) + 1000, 1000),
                       name = "Number of genes") +
    scale_x_discrete(labels = c("8dpp" = "Susceptible\nmodules\n(8dpp)",
                                 "16dpp" = "Resistant\nmodules\n(16dpp)")) +
     cowplot::theme_cowplot(font_size = 14) +
    theme(legend.direction = "horizontal", legend.position = c(0.05, 0.1))

sixC
```

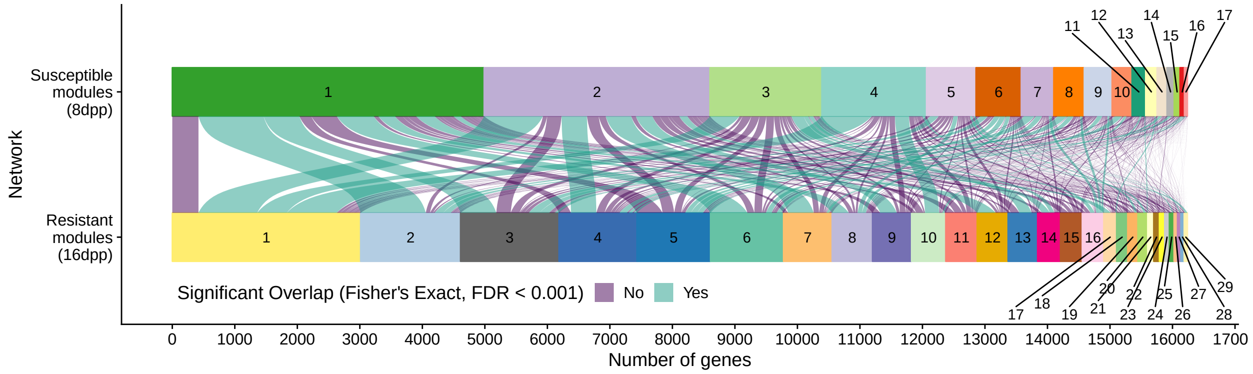


Plot VST expression data for each module

```
moduleExp16 <-
  as.data.frame(assay(vst[colnames(infection16dpp), ])) %>%
  mutate(gene = rownames(.),
         module = moduleColors16.custom,
         moduleLabel = moduleLabels16) %>%
  gather("sample", "counts",-gene, -module, -moduleLabel) %>%
  mutate(age = rep(vst$age, each = ncol(infection16dpp)),
         timepoint = rep(str_remove(vst$timepoint, "T"), each =  ncol(infection16dpp)),
         treatment = rep(vst$treatment, each =  ncol(infection16dpp))
  ) %>%
  left_join(., as_tibble(table(moduleColors16.custom)), by = c("module" = "moduleColors16.custom")) %>%
  mutate(timepoint = as.numeric(timepoint),
         module = fct_infreq(module),
         facetLabel = fct_infreq(paste0("R", moduleLabel, " (n=", n,")"))
         )

modSubset <- unique(moduleExp16$moduleLabel)
modSubset <- modSubset[!modSubset %in% 0]
```

Plot variance stabilized transformed read counts for resistant modules and overlay corrisponding values for other treament conditions. Figure 7.

```
p1 <- moduleExp16 %>%
  filter(moduleLabel %in% modSubset) %>%
  ggplot() +
  stat_summary(
    data = filter(moduleExp16, treatment == "Inoc" &
                    moduleLabel %in% modSubset),
    fun.data = mean_se,
```

```
    geom = "ribbon",
    aes(x = timepoint, y = counts, fill = age),
    alpha = 0.2
  ) +
  stat_summary(
    data = filter(moduleExp16, treatment == "Cont" &
                    moduleLabel %in% modSubset),
    fun.data = mean_se,
    geom = "ribbon",
    aes(x = timepoint, y = counts, fill = age),
    alpha = 0.2
  ) +
  stat_summary(
    fun.y = mean,
    geom = "line",
    aes(
      x = timepoint,
      y = counts,
      color = age,
      linetype = treatment
    )
  ) +
  stat_summary(
    fun.y = mean,
    geom = "point",
    aes(
      x = timepoint,
      y = counts,
      group = paste0(age, treatment)
    ), shape = 16,
    color = rep(rep(c("#00BFC4", "#00BFC4", "#F8766D", "#F8766D"), each = 7), length(modSubset))
  ) +
  facet_wrap(~ facetLabel, scales = "free", ncol = 5) +
  scale_x_continuous(breaks = c(0, 2, 4, 8, 12, 18, 24)) +
  labs(x = "Hours post inoculation", y = "Variance stabilizing transformed read count") +
  guides(linetype = guide_legend(title="Treatment"),
         color = guide_legend(title = "Age"),
         fill = guide_legend(title = "Age")) +
  scale_linetype_discrete(labels = c("Control", "Inoculated")) +
  scale_color_discrete(labels = c("8 dpp", "16 dpp")) +
  scale_fill_discrete(labels = c("8 dpp", "16 dpp")) +
  cowplot::theme_cowplot(font_size = 11) +
    theme(strip.text = element_text(
        colour = "grey10",
        size = rel(0.8),
        margin = margin(0.5 * 7, 0.5 * 7, 0.5 * 7, 0.5 * 7)
        )
    ) +
    theme(legend.position=c(0.85, 0.05),
        legend.box = 'vertical',
        legend.margin = margin(t = -5)) +
    cowplot::panel_border()
```

```
g1 <- ggplot_gtable(ggplot_build(p1))
stripr <- which(grepl('strip-t', g1$layout$name) & grepl("gTree", g1$grobs))

fills <- names(sort(table(moduleColors16.custom), decreasing = T))
fills <- names(sort(table(moduleColors16.custom), decreasing = T))[unlist(rev(split(1:length(fills), ce[illegible]


k <- 1

for (i in stripr) {
    j <- which(grepl('rect', g1$grobs[[i]]$grobs[[1]]$childrenOrder))
    g1$grobs[[i]]$grobs[[1]]$children[[j]]$gp$fill <- fills[k]
    k <- k+1
}
grid::grid.draw(g1)
```

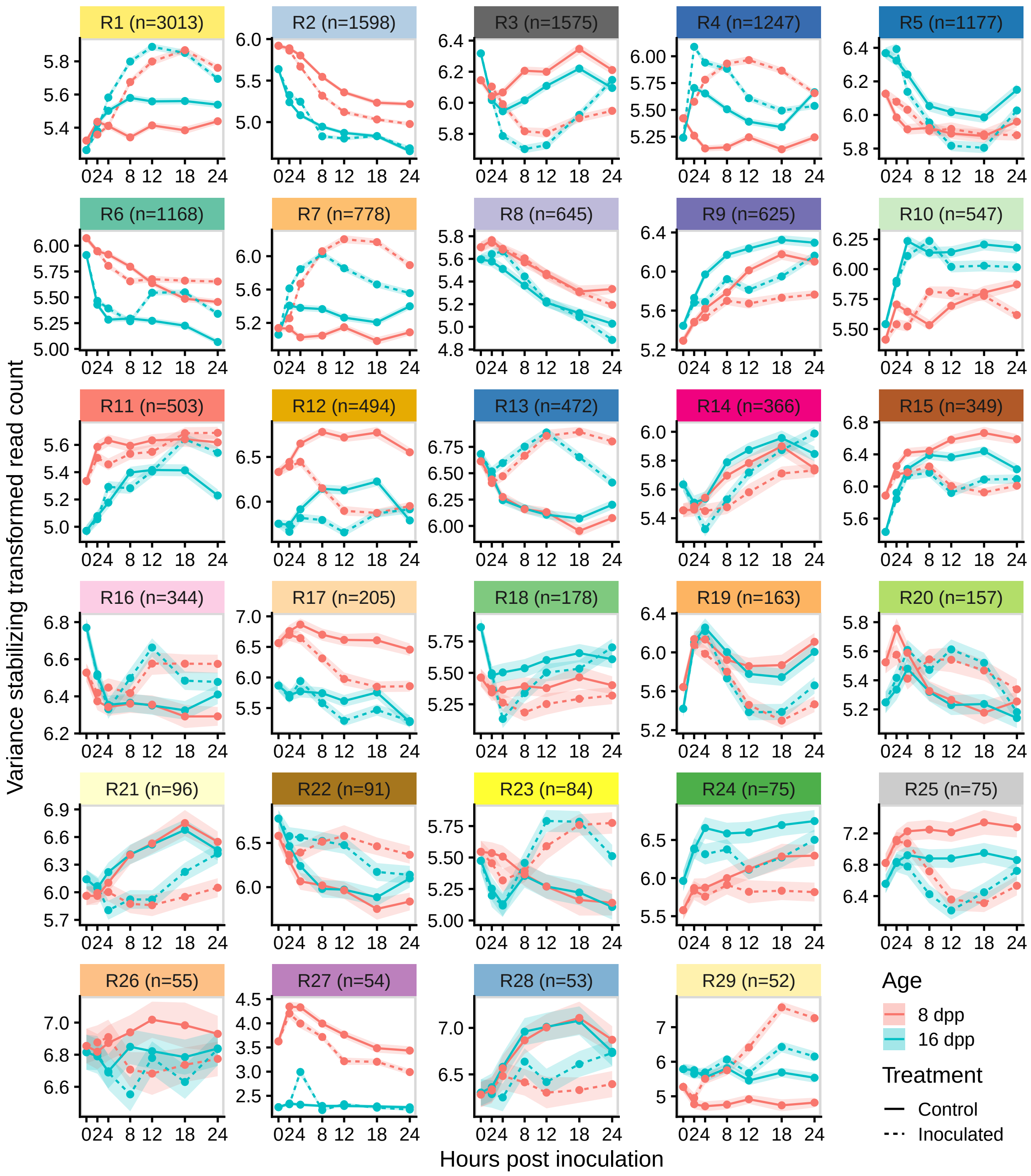


Plot VST expression data for each module in susceptible network:

```
moduleExp8 <-
    as.data.frame(assay(vst[colnames(infection8dpp), ])) %>%
    mutate(gene = rownames(.),
           module = moduleColors8.custom,
           moduleLabel = moduleLabels8) %>%
    gather("sample", "counts",-gene, -module, -moduleLabel) %>%
    mutate(age = rep(vst$age, each = ncol(infection8dpp)),
```

```r
          timepoint = rep(str_remove(vst$timepoint, "T"), each =  ncol(infection8dpp)),
          treatment = rep(vst$treatment, each =  ncol(infection8dpp))
    ) %>%
    left_join(., as_tibble(table(moduleColors8.custom)), by = c("module" = "moduleColors8.custom")) %>%
    mutate(timepoint = as.numeric(timepoint),
          module = fct_infreq(module),
          facetLabel = fct_infreq(paste0("S", moduleLabel, " (n=", n,")"))
    )

modSubset <- unique(moduleExp8$moduleLabel)
modSubset <- modSubset[!modSubset %in% 0]
```

Plot variance stabilized transformed read counts for susceptible modules and overlay corrisponding values for other treament conditions.

```r
moduleExp8  %>%
  filter(moduleLabel %in% modSubset) %>%
  ggplot() +
  stat_summary(
    data = filter(moduleExp8, treatment == "Inoc" &
                    moduleLabel %in% modSubset),
    fun.data = mean_se,
    geom = "ribbon",
    aes(x = timepoint, y = counts, fill = age),
    alpha = 0.2
  ) +
  stat_summary(
    data = filter(moduleExp8, treatment == "Cont" &
                    moduleLabel %in% modSubset),
    fun.data = mean_se,
    geom = "ribbon",
    aes(x = timepoint, y = counts, fill = age),
    alpha = 0.2
  ) +
  stat_summary(
    fun.y = mean,
    geom = "line",
    aes(
      x = timepoint,
      y = counts,
      color = age,
      linetype = treatment
    )
  ) +
  stat_summary(
    fun.y = mean,
    geom = "point",
    aes(
      x = timepoint,
      y = counts,
      group = paste0(age, treatment)
    ), shape = 16,
    color = rep(rep(c("#00BFC4", "#00BFC4", "#F8766D", "#F8766D"), each = 7), length(modSubset))
  ) +
```

```
facet_wrap(~ facetLabel, scales = "free", ncol = 5) +
scale_x_continuous(breaks = c(0, 2, 4, 8, 12, 18, 24)) +
labs(x = "Hours post inoculation", y = "Variance stabilizing transformed read count") +
guides(linetype = guide_legend(title="Treatment"),
       color = guide_legend(title = "Age"),
       fill = guide_legend(title = "Age")) +
scale_linetype_discrete(labels = c("Control", "Inoculated")) +
scale_color_discrete(labels = c("8 dpp", "16 dpp")) +
scale_fill_discrete(labels = c("8 dpp", "16 dpp")) +
cowplot::theme_cowplot(font_size = 11) +
  theme(strip.text = element_text(
      colour = "grey10",
      size = rel(0.8),
      margin = margin(0.5 * 7, 0.5 * 7, 0.5 * 7, 0.5 * 7)
      )
  ) +
  theme(legend.position=c(0.85, 0.05),
      legend.box = 'vertical',
      legend.margin = margin(t = -5)) +
  cowplot::panel_border()
```

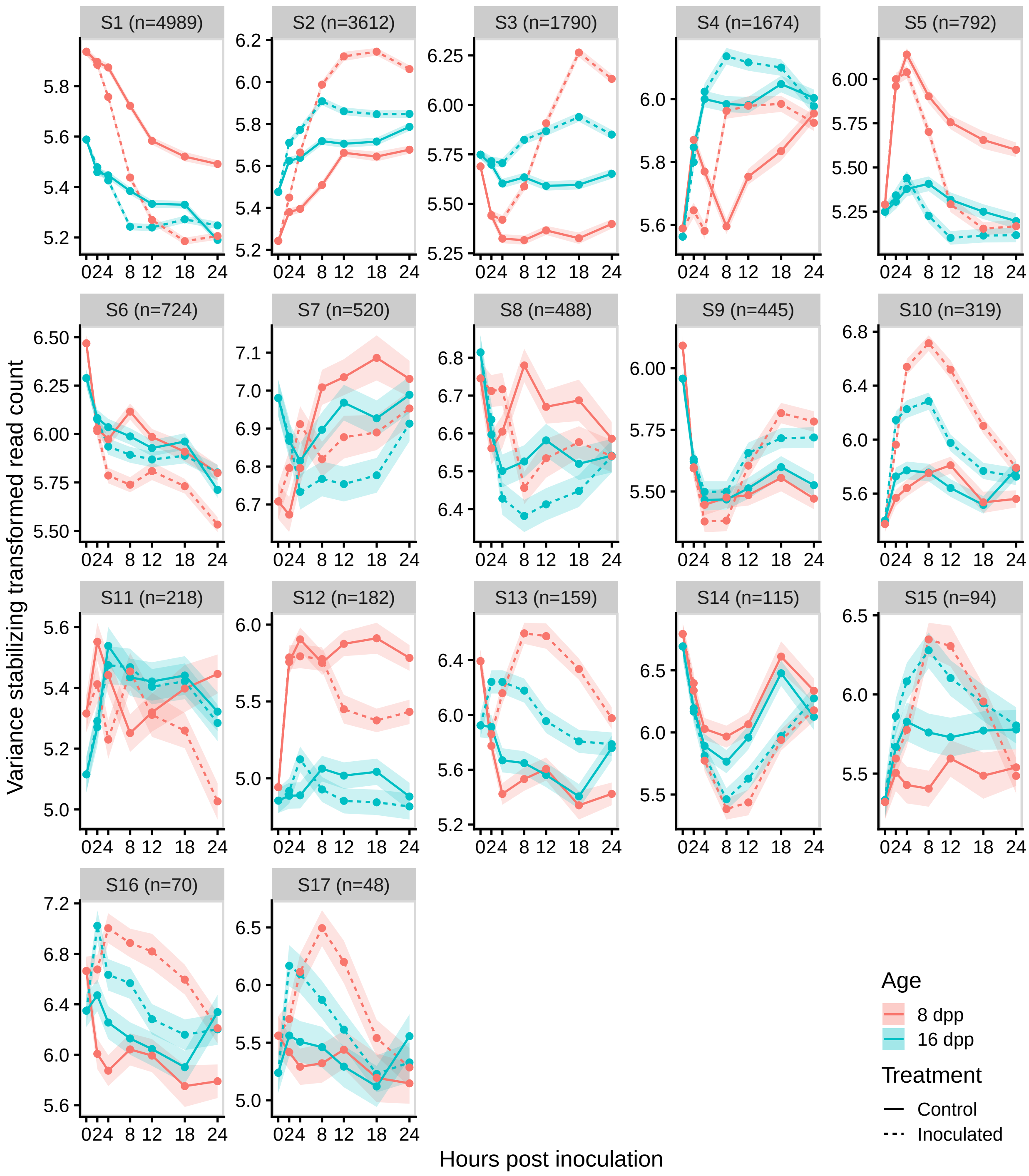


```
# g1 <- ggplot_gtable(ggplot_build(p1))
# stripr <- which(grepl('strip-t', g1$layout$name) & grepl("gTree", g1$grobs))
#
# fills <- names(sort(table(moduleColors8.custom), decreasing = T))
# fills <- names(sort(table(moduleColors8.custom), decreasing = T))[unlist(rev(split(1:length(fills), c
#
#
# k <- 1
```

```
#
# for (i in stripr) {
#     j <- which(grepl('rect', g1$grobs[[i]]$grobs[[1]]$childrenOrder))
#     g1$grobs[[i]]$grobs[[1]]$children[[j]]$gp$fill <- fills[k]
#     k <- k+1
# }
# grid::grid.draw(g1)
```

Arrange the read count data in tidy format and add module numbers and log counts

```
modulecounts16 <-
  as.data.frame(counts(dds[colnames(infection16dpp), ], normalized = T)) %>%
  mutate(gene = rownames(.),
         module = moduleColors16.custom,
         moduleLabel = moduleLabels16) %>%
  gather("sample", "counts",-gene, -module, -moduleLabel) %>%
  mutate(age = rep(vst$age, each = ncol(infection16dpp)),
         timepoint = rep(str_remove(vst$timepoint, "T"), each =  ncol(infection16dpp)),
         treatment = rep(vst$treatment, each =  ncol(infection16dpp))
  ) %>%
  left_join(., as_tibble(table(moduleColors16.custom)), by = c("module" = "moduleColors16.custom")) %>%
  mutate(timepoint = as.numeric(timepoint),
         module = fct_infreq(module),
         facetLabel = fct_infreq(paste0("R", moduleLabel, " (n=", n,")")),
         logCounts = log2(counts + 0.5)
  )

modulecounts8 <-
  as.data.frame(counts(dds[colnames(infection8dpp), ], normalized = T)) %>%
  mutate(gene = rownames(.),
         module = moduleColors8.custom,
         moduleLabel = moduleLabels8) %>%
  gather("sample", "counts",-gene, -module, -moduleLabel) %>%
  mutate(age = rep(vst$age, each = ncol(infection8dpp)),
         timepoint = rep(str_remove(vst$timepoint, "T"), each =  ncol(infection8dpp)),
         treatment = rep(vst$treatment, each =  ncol(infection8dpp))
  ) %>%
  left_join(., as_tibble(table(moduleColors8.custom)), by = c("module" = "moduleColors8.custom")) %>%
  mutate(timepoint = as.numeric(timepoint),
         module = fct_infreq(module),
         facetLabel = fct_infreq(paste0("S", moduleLabel, " (n=", n,")")),
         logCounts = log2(counts + 0.5)
  )
```

Genes in ModS1 and their expression patterns in other modules. Figure 6D

```
networkLodes %>%
    filter(module8dpp == 1, isSig == "Yes")
```

```
##    module16dpp module8dpp overlap        pvalue Cohort Network  module
## 1            2          1    1042 3.194379e-198      2   16dpp #B3CDE3
## 2            3          1     572  3.320226e-07      3   16dpp #666666
## 3            6          1     589  2.472749e-48      6   16dpp #66C2A5
## 4            8          1     266  5.160602e-09      8   16dpp #BEBADA
## 5           12          1     296  3.509969e-42     12   16dpp #E6AB02
```

```
## 6           17          1     115  3.280177e-14     17   16dpp #FED9A6
## 7            2          1    1042 3.194379e-198      2    8dpp #33A02C
## 8            3          1     572  3.320226e-07      3    8dpp #33A02C
## 9            6          1     589  2.472749e-48      6    8dpp #33A02C
## 10           8          1     266  5.160602e-09      8    8dpp #33A02C
## 11          12          1     296  3.509969e-42     12    8dpp #33A02C
## 12          17          1     115  3.280177e-14     17    8dpp #33A02C
##              padj isSig moduleSize moduleLabel
## 1  3.149658e-195   Yes       1598           2
## 2   2.994843e-04   Yes       1575           3
## 3   2.403512e-45   Yes       1168           6
## 4   4.737433e-06   Yes        645           8
## 5   3.404670e-39   Yes        494          12
## 6   3.089926e-11   Yes        205          17
## 7  3.149658e-195   Yes       4989           1
## 8   2.994843e-04   Yes       4989           1
## 9   2.403512e-45   Yes       4989           1
## 10  4.737433e-06   Yes       4989           1
## 11  3.404670e-39   Yes       4989           1
## 12  3.089926e-11   Yes       4989           1
```

```r
S1_R2_genes <- moduleExp8 %>%
    dplyr::select(gene, moduleLabel) %>%
    left_join(dplyr::select(moduleExp16, gene, moduleLabel), by = "gene", suffix = c(".8dpp", ".16dpp"))
    filter(moduleLabel.8dpp == 1) %>%
    filter(moduleLabel.16dpp == 2) %>%
    pull(gene) %>%
    unique()

S1_R3_genes <- moduleExp8 %>%
    dplyr::select(gene, moduleLabel) %>%
    left_join(dplyr::select(moduleExp16, gene, moduleLabel), by = "gene", suffix = c(".8dpp", ".16dpp"))
    filter(moduleLabel.8dpp == 1) %>%
    filter(moduleLabel.16dpp == 3) %>%
    pull(gene) %>%
    unique()

S1_R6_genes <- moduleExp8 %>%
    dplyr::select(gene, moduleLabel) %>%
    left_join(dplyr::select(moduleExp16, gene, moduleLabel), by = "gene", suffix = c(".8dpp", ".16dpp"))
    filter(moduleLabel.8dpp == 1) %>%
    filter(moduleLabel.16dpp == 6) %>%
    pull(gene) %>%
    unique()

S1_R8_genes <- moduleExp8 %>%
    dplyr::select(gene, moduleLabel) %>%
    left_join(dplyr::select(moduleExp16, gene, moduleLabel), by = "gene", suffix = c(".8dpp", ".16dpp"))
    filter(moduleLabel.8dpp == 1) %>%
    filter(moduleLabel.16dpp == 8) %>%
    pull(gene) %>%
    unique()

S1_R12_genes <- moduleExp8 %>%
```

```
    dplyr::select(gene, moduleLabel) %>%
    left_join(dplyr::select(moduleExp16, gene, moduleLabel), by = "gene", suffix = c(".8dpp", ".16dpp")
    filter(moduleLabel.8dpp == 1) %>%
    filter(moduleLabel.16dpp == 12) %>%
    pull(gene) %>%
    unique()

S1_R17_genes <- moduleExp8 %>%
    dplyr::select(gene, moduleLabel) %>%
    left_join(dplyr::select(moduleExp16, gene, moduleLabel), by = "gene", suffix = c(".8dpp", ".16dpp")
    filter(moduleLabel.8dpp == 1) %>%
    filter(moduleLabel.16dpp == 17) %>%
    pull(gene) %>%
    unique()

consModExp <- bind_rows("S1 vs R2\nGene overlap: 1042" = filter(moduleExp8, gene %in% S1_R2_genes),
                        "S1 vs R3\nGene overlap: 572" = filter(moduleExp8, gene %in% S1_R3_genes),
                        "S1 vs R6\nGene overlap: 589" = filter(moduleExp8, gene %in% S1_R6_genes),
                        "S1 vs R8\nGene overlap: 266" = filter(moduleExp8, gene %in% S1_R8_genes),
                        "S1 vs R12\nGene overlap: 296" = filter(moduleExp8, gene %in% S1_R12_genes),
                        "S1 vs R17\nGene overlap: 115" = filter(moduleExp8, gene %in% S1_R17_genes),
                        .id = "modComp") %>%
    mutate(modComp = fct_inorder(modComp))

sixD <- filter(consModExp, treatment == "Inoc")  %>%
  ggplot() +
  stat_summary(
    data = filter(consModExp, treatment == "Inoc"),
    fun.data = mean_se,
    geom = "ribbon",
    aes(x = timepoint, y = counts, fill = age),
    alpha = 0.2
  ) +
  # stat_summary(
  #   data = filter(consModExp, treatment == "Cont"),
  #   fun.data = mean_se,
  #   geom = "ribbon",
  #   aes(x = timepoint, y = counts, fill = age),
  #   alpha = 0.2
  # ) +
  stat_summary(
    fun.y = mean,
    geom = "line",
    aes(
      x = timepoint,
      y = counts,
      color = age,
      # linetype = treatment
    )
  ) +
  stat_summary(
    fun.y = mean,
    geom = "point",
```

```
    aes(
      x = timepoint,
      y = counts,
      group = paste0(age, treatment)
    ), shape = 16,
    # color = rep(rep(c("#00BFC4", "#00BFC4", "#F8766D", "#F8766D"), each = 7), 6)
  ) +
  facet_wrap(~ modComp, scales = "free", ncol = 6) +
  scale_x_continuous(breaks = c(0, 2, 4, 8, 12, 18, 24)) +
  labs(x = "Hours post inoculation", y = "Variance stabilizing transformed read count") +
  guides(#linetype = guide_legend(title="Treatment"),
         color = guide_legend(title = "Age"),
         fill = guide_legend(title = "Age")) +
  scale_color_discrete(labels = c("8 dpp", "16 dpp")) +
  scale_fill_discrete(labels = c("8 dpp", "16 dpp")) +
  cowplot::theme_cowplot(font_size = 11) +
    theme(strip.text = element_text(
        colour = "grey10",
        size = rel(0.8),
        margin = margin(0.5 * 7, 0.5 * 7, 0.5 * 7, 0.5 * 7)
        )
    ) +
    theme(legend.position = c(0.925, 0.875),
        # legend.position="bottom",
        # legend.box = 'vertical',
        legend.margin = margin(t = -5)) +
    cowplot::panel_border()

sixD
```

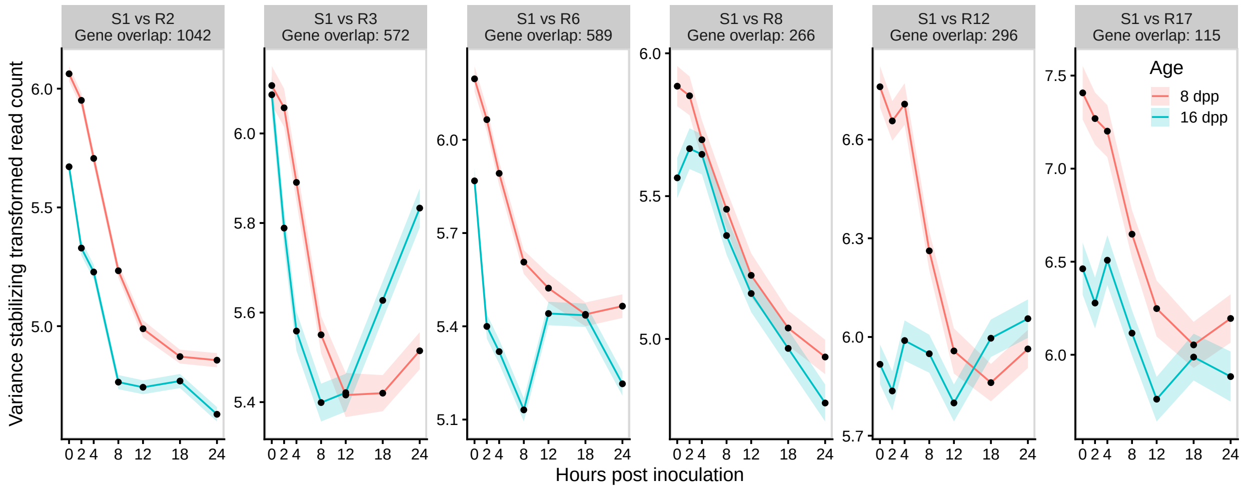


Figure 6:

```
sixA <- cowplot::ggdraw() + cowplot::draw_image(image = "fig6A.png")
sixB <- cowplot::ggdraw() + cowplot::draw_image(image = "fig6B.png")

sixAB <- cowplot::plot_grid(sixA, sixB,
                            ncol = 2,
```

```r
                       labels = "AUTO")

cowplot::plot_grid(sixAB,
                       sixC,
                       sixD,
                       ncol = 1,
                       labels = c("", "C", "D"),
                       rel_heights = c(3, 2, 2.1))
```

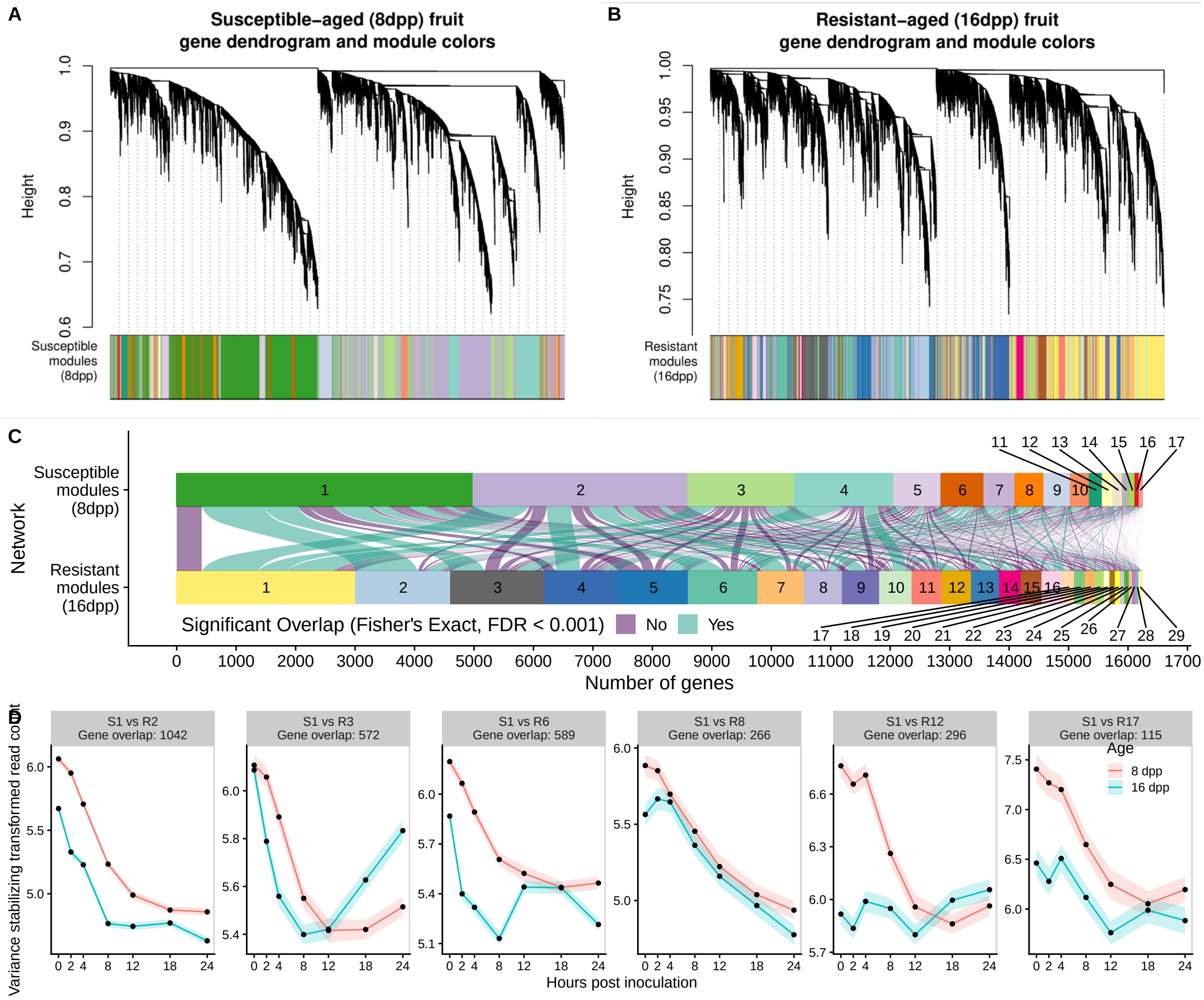


```r
# pdf(file = "fig6.pdf", width = 7, 9)
# cowplot::plot_grid(drPlot, poin, mic,
#                        ncol = 1,
#                        labels = "AUTO",
#                        rel_heights = c(3, 1.5, 1.8))
# dev.off()
```

Export all genes and GO term enrichment results for Resistant network modules

```r
outGenes <- list()
k <- 1
```

```
for (i in 1:max(modSubset)) {
    outGenes[[k]] <- modulecounts16 %>%
        filter(moduleLabel %in% i) %>%
        dplyr::select(-module,  -moduleLabel, -age, -timepoint, -treatment ,-n, -facetLabel, -logCounts
        spread(sample, counts)
    k <- k + 1
}

names(outGenes) <- paste0("Module R", 1:max(modSubset))

library(writexl)

write_xlsx(outGenes, "Supplementary File 3_Resistant_Modules_Genes.xlsx", col_names = T)
```

```
GOres <- list()
k <- 1
for (i in 1:max(modSubset)) {
GOres[[k]] <- moduleGOterms(
  network = "16dpp",
  mod = i,
  nodeSize = 100,
  n = 500
  ) %>% filter(Fisher.weight01 < 0.05)
k <- k + 1
}

names(GOres) <- paste0("Module R", 1:max(modSubset))
```

```
write_xlsx(GOres, "Supplementary File 5_Resistant_Modules_GO_Enrichment.xlsx", col_names = T)
```

Export gene module assignments for susceptible network:

```
outGenes <- list()
k <- 1
for (i in 1:max(modSubset)) {
    outGenes[[k]] <- modulecounts8 %>%
        filter(moduleLabel %in% i) %>%
        dplyr::select(-module,  -moduleLabel, -age, -timepoint, -treatment ,-n, -facetLabel, -logCounts
        spread(sample, counts)
    k <- k + 1
}

names(outGenes) <- paste0("Module R", 1:max(modSubset))

library(writexl)

write_xlsx(outGenes, "Supplementary File 4_Susceptible_Modules_Genes.xlsx", col_names = T)
```

Which have significant interaction effects and then in which spline portion do they have an interaction

```
library(splines)

modulecounts16 %>%
  filter(treatment == "Inoc") %>%
  group_by(moduleLabel) %>%
```

```r
  do(anova = anova(lm(logCounts ~ age*ns(timepoint, df = 4), data = .))) %>%
  broom::tidy(anova) %>%
  filter(str_detect(term, "age:ns"), p.value < 0.05) %>% kable()
```

| moduleLabel | term | df | sumsq | meansq | statistic | p.value |
|---:|---|---:|---:|---:|---:|---:|
| 1 | age:ns(timepoint, df = 4) | 4 | 329.99965 | 82.49991 | 15.934368 | 0.0000000 |
| 2 | age:ns(timepoint, df = 4) | 4 | 314.39321 | 78.59830 | 14.955229 | 0.0000000 |
| 3 | age:ns(timepoint, df = 4) | 4 | 556.14154 | 139.03539 | 27.257186 | 0.0000000 |
| 4 | age:ns(timepoint, df = 4) | 4 | 1372.47125 | 343.11781 | 64.269490 | 0.0000000 |
| 5 | age:ns(timepoint, df = 4) | 4 | 330.53709 | 82.63427 | 14.554743 | 0.0000000 |
| 6 | age:ns(timepoint, df = 4) | 4 | 405.35071 | 101.33768 | 23.749391 | 0.0000000 |
| 7 | age:ns(timepoint, df = 4) | 4 | 940.10468 | 235.02617 | 42.185627 | 0.0000000 |
| 8 | age:ns(timepoint, df = 4) | 4 | 126.22441 | 31.55610 | 5.047284 | 0.0004597 |
| 9 | age:ns(timepoint, df = 4) | 4 | 73.84148 | 18.46037 | 3.210159 | 0.0120955 |
| 10 | age:ns(timepoint, df = 4) | 4 | 172.79071 | 43.19768 | 8.528060 | 0.0000007 |
| 11 | age:ns(timepoint, df = 4) | 4 | 135.22117 | 33.80529 | 7.716836 | 0.0000033 |
| 12 | age:ns(timepoint, df = 4) | 4 | 503.94481 | 125.98620 | 25.395775 | 0.0000000 |
| 13 | age:ns(timepoint, df = 4) | 4 | 223.30571 | 55.82643 | 13.362317 | 0.0000000 |
| 14 | age:ns(timepoint, df = 4) | 4 | 106.54735 | 26.63684 | 6.508226 | 0.0000315 |
| 15 | age:ns(timepoint, df = 4) | 4 | 207.59172 | 51.89793 | 10.683718 | 0.0000000 |
| 16 | age:ns(timepoint, df = 4) | 4 | 72.22910 | 18.05727 | 4.950540 | 0.0005495 |
| 18 | age:ns(timepoint, df = 4) | 4 | 98.17228 | 24.54307 | 5.643348 | 0.0001563 |
| 19 | age:ns(timepoint, df = 4) | 4 | 68.86208 | 17.21552 | 3.174732 | 0.0129011 |
| 20 | age:ns(timepoint, df = 4) | 4 | 55.10566 | 13.77642 | 3.567641 | 0.0065185 |
| 21 | age:ns(timepoint, df = 4) | 4 | 41.93763 | 10.48441 | 2.463965 | 0.0431030 |
| 22 | age:ns(timepoint, df = 4) | 4 | 52.10105 | 13.02526 | 3.387247 | 0.0089643 |
| 23 | age:ns(timepoint, df = 4) | 4 | 41.79785 | 10.44946 | 3.288753 | 0.0106407 |
| 27 | age:ns(timepoint, df = 4) | 4 | 104.56769 | 26.14192 | 11.520982 | 0.0000000 |
| 29 | age:ns(timepoint, df = 4) | 4 | 410.30424 | 102.57606 | 19.212257 | 0.0000000 |

```r
modulecounts16 %>%
  filter(treatment == "Inoc") %>%
  group_by(moduleLabel) %>%
  do(fit = summary(lm(counts ~ age*ns(timepoint, df = 4), data = .))) %>%
  broom::tidy(fit) %>%
  filter(str_detect(term, "16dpp:ns"), p.value < 0.05) %>%
  kable()
```

| moduleLabel | term | estimate | std.error | statistic | p.value |
|---|---|---|---|---|---|
| 1 | age16dpp:ns(timepoint, df = 4)1 | 33.716112 | 12.587751 | 2.678486 | 0.0073966 |
| 1 | age16dpp:ns(timepoint, df = 4)4 | -17.708278 | 8.045915 | -2.200903 | 0.0277447 |
| 2 | age16dpp:ns(timepoint, df = 4)2 | 34.164062 | 12.604516 | 2.710462 | 0.0067207 |
| 2 | age16dpp:ns(timepoint, df = 4)4 | 27.622086 | 8.593784 | 3.214194 | 0.0013087 |
| 3 | age16dpp:ns(timepoint, df = 4)3 | -54.547120 | 24.827976 | -2.197002 | 0.0280238 |
| 3 | age16dpp:ns(timepoint, df = 4)4 | 42.540349 | 12.464383 | 3.412952 | 0.0006430 |
| 4 | age16dpp:ns(timepoint, df = 4)1 | -50.846834 | 25.008103 | -2.033214 | 0.0420360 |
| 4 | age16dpp:ns(timepoint, df = 4)2 | -47.739781 | 23.444978 | -2.036248 | 0.0417306 |
| 4 | age16dpp:ns(timepoint, df = 4)4 | -52.898562 | 15.984832 | -3.309297 | 0.0009360 |
| 6 | age16dpp:ns(timepoint, df = 4)2 | 25.219679 | 8.301942 | 3.037805 | 0.0023844 |
| 7 | age16dpp:ns(timepoint, df = 4)2 | -89.246529 | 39.228200 | -2.275060 | 0.0229089 |
| 7 | age16dpp:ns(timepoint, df = 4)4 | -66.615752 | 26.745863 | -2.490694 | 0.0127544 |
| 12 | age16dpp:ns(timepoint, df = 4)1 | 84.857949 | 42.235479 | 2.009163 | 0.0445331 |
| 12 | age16dpp:ns(timepoint, df = 4)2 | 79.431336 | 39.595562 | 2.006067 | 0.0448624 |
| 12 | age16dpp:ns(timepoint, df = 4)4 | 84.269099 | 26.996331 | 3.121502 | 0.0018018 |
| 13 | age16dpp:ns(timepoint, df = 4)2 | -72.656693 | 24.064806 | -3.019210 | 0.0025377 |
| 13 | age16dpp:ns(timepoint, df = 4)3 | -81.096923 | 32.682187 | -2.481380 | 0.0130959 |
| 13 | age16dpp:ns(timepoint, df = 4)4 | -85.829751 | 16.407431 | -5.231151 | 0.0000002 |
| 15 | age16dpp:ns(timepoint, df = 4)3 | 139.282482 | 68.398006 | 2.036353 | 0.0417333 |
| 16 | age16dpp:ns(timepoint, df = 4)3 | -93.437517 | 30.767404 | -3.036900 | 0.0023946 |
| 17 | age16dpp:ns(timepoint, df = 4)4 | 237.608300 | 115.102668 | 2.064316 | 0.0390184 |
| 18 | age16dpp:ns(timepoint, df = 4)3 | -46.351422 | 19.764345 | -2.345204 | 0.0190433 |
| 22 | age16dpp:ns(timepoint, df = 4)4 | -55.705134 | 28.027657 | -1.987506 | 0.0469395 |
| 27 | age16dpp:ns(timepoint, df = 4)1 | 13.974305 | 3.277604 | 4.263573 | 0.0000210 |
| 27 | age16dpp:ns(timepoint, df = 4)2 | 8.463938 | 3.072738 | 2.754526 | 0.0059260 |
| 27 | age16dpp:ns(timepoint, df = 4)4 | 14.381642 | 2.094999 | 6.864749 | 0.0000000 |
| 29 | age16dpp:ns(timepoint, df = 4)2 | -496.697392 | 99.904288 | -4.971732 | 0.0000007 |
| 29 | age16dpp:ns(timepoint, df = 4)3 | -413.950980 | 135.679075 | -3.050957 | 0.0023094 |
| 29 | age16dpp:ns(timepoint, df = 4)4 | -275.218091 | 68.114936 | -4.040495 | 0.0000552 |

normalized gene expression and fitted curves with splined time

```
pME <- modulecounts16 %>%
  filter(treatment == "Inoc") %>%
  group_by(moduleLabel) %>%
  ggplot(aes(x = timepoint, y = logCounts, color = age)) +
  stat_summary(fun.data = "mean_se", size = 0.25) +
  geom_smooth(method = lm,
              formula = y ~ splines::ns(x, df = 4), linetype = 2) +
  facet_wrap(~ moduleLabel, scales = "free", ncol = 5) +
  labs(x = "Hours post inoculation", y = "Log(Normalized counts)") +
  theme_gray() +
  theme(legend.position="bottom")

gME <- ggplot_gtable(ggplot_build(pME))
stripr <- which(grepl('strip-t', gME$layout$name) & grepl("gTree", gME$grobs))

fills <- names(sort(table(moduleColors16.custom), decreasing = T))
fills <- names(
  sort(
    table(
      moduleColors16.custom), decreasing = T)
  )[unlist(rev(split(1:length(fills), ceiling(seq_along(1:length(fills))/5)))),
```

```
           use.names = F)]


k <- 1

for (i in stripr) {
    j <- which(grepl('rect', gME$grobs[[i]]$grobs[[1]]$childrenOrder))
    gME$grobs[[i]]$grobs[[1]]$children[[j]]$gp$fill <- fills[k]
    k <- k+1
}
grid::grid.draw(gME)
```

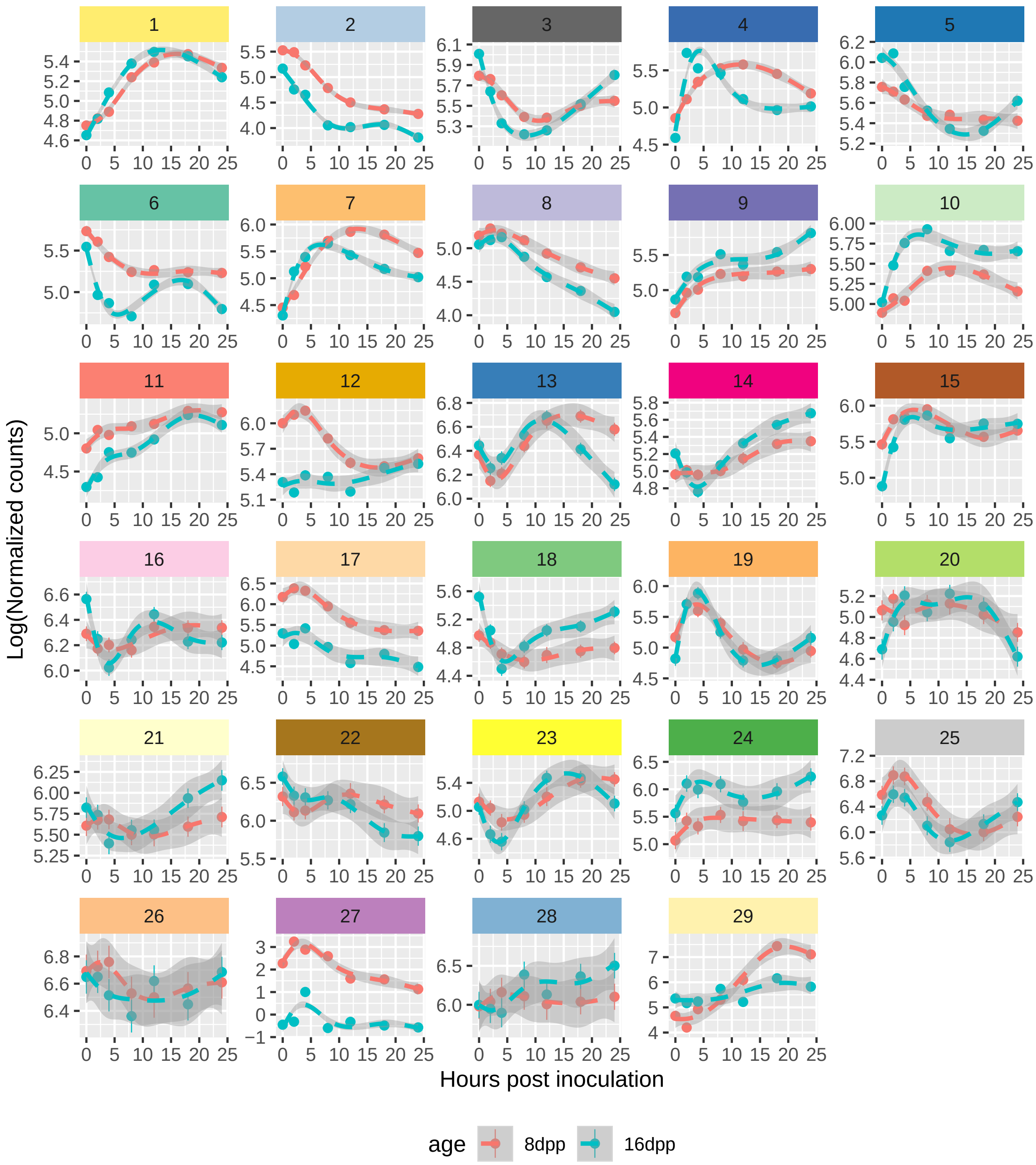


Which modules have an interaction in the first or second spline portions

```
library(splines)

anovaModExp <- modulecounts16 %>%
  filter(treatment == "Inoc") %>%
  group_by(moduleLabel) %>%
  do(fit = summary(lm(counts ~ age*ns(timepoint, df = 4), data = .))) %>%
  broom::tidy(fit) %>%
```

```r
  filter(str_detect(term, "age16dpp\\:ns.*[1-2]"), p.value < 0.05)

anovaModExp
```

```
## # A tibble: 12 x 6
## # Groups:   moduleLabel [9]
##    moduleLabel term                      estimate std.error statistic    p.value
##          <dbl> <chr>                        <dbl>     <dbl>     <dbl>      <dbl>
##  1           1 age16dpp:ns(timepoint, d...    33.7      12.6       2.68    7.40e-3
##  2           2 age16dpp:ns(timepoint, d...    34.2      12.6       2.71    6.72e-3
##  3           4 age16dpp:ns(timepoint, d...   -50.8      25.0      -2.03    4.20e-2
##  4           4 age16dpp:ns(timepoint, d...   -47.7      23.4      -2.04    4.17e-2
##  5           6 age16dpp:ns(timepoint, d...    25.2       8.30      3.04    2.38e-3
##  6           7 age16dpp:ns(timepoint, d...   -89.2      39.2      -2.28    2.29e-2
##  7          12 age16dpp:ns(timepoint, d...    84.9      42.2       2.01    4.45e-2
##  8          12 age16dpp:ns(timepoint, d...    79.4      39.6       2.01    4.49e-2
##  9          13 age16dpp:ns(timepoint, d...   -72.7      24.1      -3.02    2.54e-3
## 10          27 age16dpp:ns(timepoint, d...    14.0       3.28      4.26    2.10e-5
## 11          27 age16dpp:ns(timepoint, d...     8.46      3.07      2.75    5.93e-3
## 12          29 age16dpp:ns(timepoint, d...  -497.       99.9      -4.97    7.17e-7
```

```r
GOres <- list()

k <- 1
for (i in unique(anovaModExp$moduleLabel)) {
GOres[[k]] <- moduleGOterms(
  network = "16dpp",
  mod = i,
  nodeSize = 100,
  n = 5
  )
k <- k + 1
}

names(GOres) <- unique(anovaModExp$moduleLabel)
```

Plot log2Normalized read counts with a spline of time. Plot top 5 goterms for each module on the side. Figure 8

```r
fills <- rev(names(sort(table(moduleColors16.custom), decreasing = T))[unique(anovaModExp$moduleLabel)]]

tidyGOres <- bind_rows(GOres, .id = "moduleLabel") %>%
  mutate(moduleLabel = fct_relevel(moduleLabel, as.character(sort(unique(anovaModExp$moduleLabel))))) %
  mutate(color = rep(rev(fills), each = 5))


pME <- modulecounts16 %>%
  filter(moduleLabel %in% anovaModExp$moduleLabel) %>%
  ggplot(aes(x = timepoint, y = logCounts, color = age)) +
  stat_summary(fun.data = "mean_se", size = 0.25) +
  geom_smooth(method = lm,
              formula = y ~ splines::ns(x, df = 4), linetype = 2) +
  labs(x = "Hours post inoculation", y = expression(log[2]*"(Normalized read counts)")) +
  # theme_gray() +
  scale_x_continuous(breaks = c(0, 2, 4, 8, 12, 18, 24)) +
```

```r
 guides(color = guide_legend(title = "Age")) +
 scale_color_discrete(labels = c("8 dpp", "16 dpp")) +
    cowplot::theme_cowplot(font_size = 10) +
   theme(strip.text = element_text(
       colour = "grey10",
       size = rel(0.8),
       margin = margin(0.8 * 7, 0.8 * 7, 0.8 * 7, 0.8 * 7)
       ),
       plot.margin = unit(c(0, 0, 0, 0), "cm"),
       axis.line = element_line(),
       legend.position = "none") +
   lemon::facet_rep_wrap(~ facetLabel, scales = "free", ncol = 1) +
   cowplot::panel_border()



gME <- ggplot_gtable(ggplot_build(pME))
stripr <- which(grepl('strip-t', gME$layout$name) & grepl("gTree", gME$grobs))

k <- 1

for (i in stripr) {
    j <- which(grepl('rect', gME$grobs[[i]]$grobs[[1]]$childrenOrder))
    gME$grobs[[i]]$grobs[[1]]$children[[j]]$gp$fill <- fills[k]
    k <- k+1
}

reorder_within <- function(x, by, within, fun = mean, sep = "___", ...) {
  new_x <- paste(x, within, sep = sep)
  stats::reorder(new_x, by, FUN = fun)
}

scale_x_reordered <- function(..., sep = "___") {
  reg <- paste0(sep, ".+$")
  ggplot2::scale_x_discrete(labels = function(x) gsub(reg, "", x), ...)
}


GO_ME <- tidyGOres %>%
  ggplot() +
  geom_col(aes(y = -log10(Fisher.weight01),
               x = reorder_within(Term,
                                  by = -log10(Fisher.weight01),
                                  moduleLabel),
               fill = color)) +
  scale_x_reordered(position = "top") +
  scale_fill_identity() +
  facet_wrap(~ moduleLabel, scales = "free", ncol = 1) +
  labs(x = "", y = expression(log[10]*"(p-value)")) +
    cowplot::theme_cowplot(font_size = 10) +
  theme(strip.background = element_rect(fill = "transparent", color = "transparent"),
        axis.line = element_blank(),
        strip.text = element_text(color = "transparent",
```

```r
                                    size = rel(0.8),
                                    margin = margin(0.8 * 7, 0.8 * 7, 0.8 * 7, 0.8 * 7)),
          plot.margin = unit(c(0, 0, 0, 0), "cm"),
          axis.title.x = element_text(margin = margin(t = 5))) +
    coord_flip()

# cowplot::theme_cowplot(font_size = 10) +
#     theme(strip.text = element_text(
#         colour = "grey10",
#         size = rel(0.8),
#         margin = margin(0.8 * 7, 0.8 * 7, 0.8 * 7, 0.8 * 7)
#         ),
#         axis.line=element_line()


cowplot::plot_grid(gME, GO_ME, rel_widths = c(0.5, 1), align = "h", axis = "b")
```

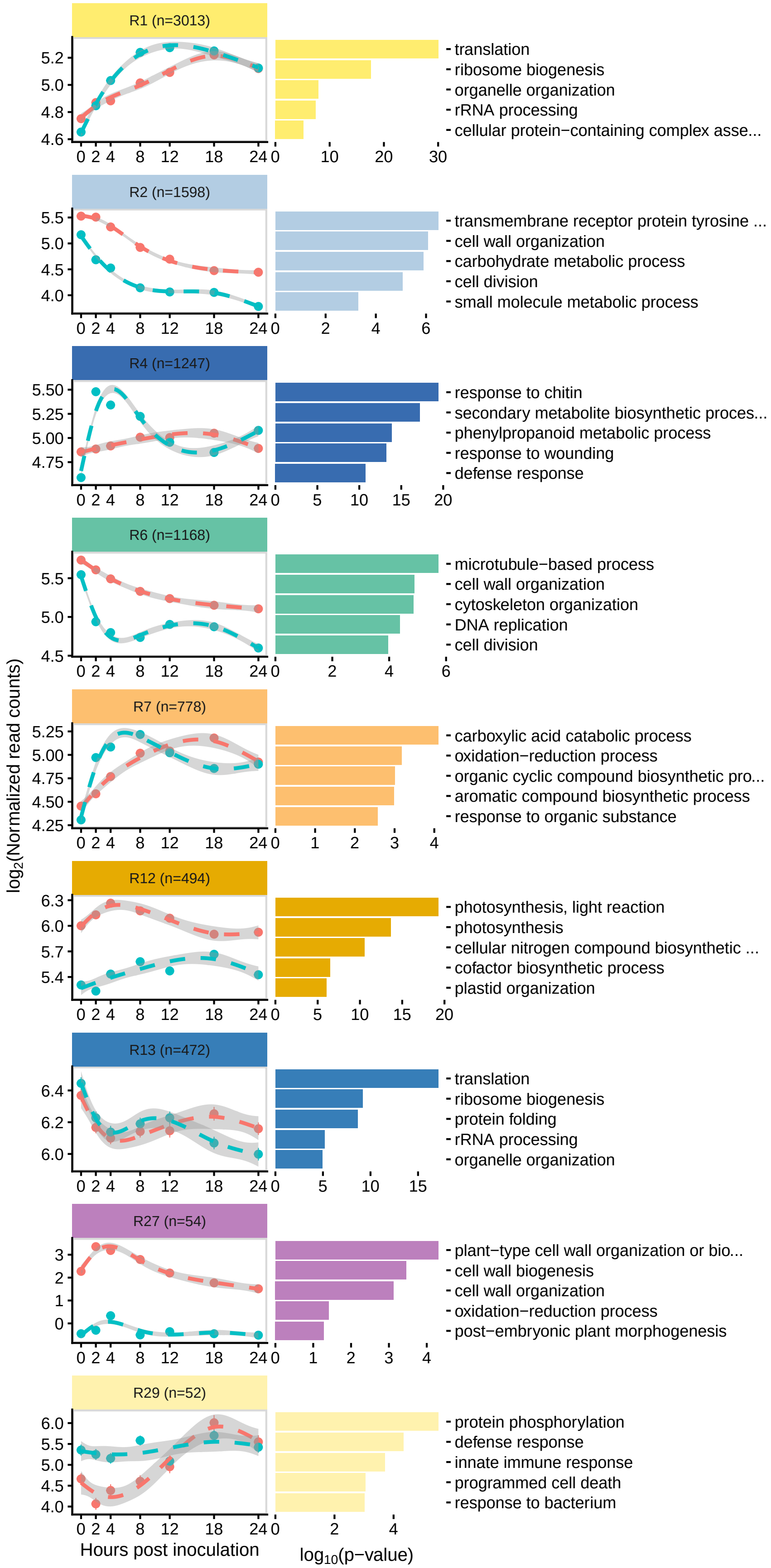
R1 (n=3013)
translation
ribosome biogenesis
organelle organization
rRNA processing
cellular protein−containing complex asse...
R2 (n=1598)
transmembrane receptor protein tyrosine ...
cell wall organization
carbohydrate metabolic process
cell division
small molecule metabolic process
R4 (n=1247)
response to chitin
secondary metabolite biosynthetic proces...
phenylpropanoid metabolic process
response to wounding
defense response
R6 (n=1168)
microtubule−based process
cell wall organization
cytoskeleton organization
DNA replication
cell division
R7 (n=778)
carboxylic acid catabolic process
oxidation−reduction process
organic cyclic compound biosynthetic pro...
aromatic compound biosynthetic process
response to organic substance
R12 (n=494)
photosynthesis, light reaction
photosynthesis
cellular nitrogen compound biosynthetic ...
cofactor biosynthetic process
plastid organization
R13 (n=472)
translation
ribosome biogenesis
protein folding
rRNA processing
organelle organization
R27 (n=54)
plant−type cell wall organization or bio...
cell wall biogenesis
cell wall organization
oxidation−reduction process
post−embryonic plant morphogenesis
R29 (n=52)
protein phosphorylation
defense response
innate immune response
programmed cell death
response to bacterium
log2(Normalized read counts)
Hours post inoculation
log10(p−value)

We can caluculate gene module membership ie the correlation between gene expression and the module eigengene

```
# module membership kME is the correlation to the module eigengene and represents the association to th
geneModuleMembership16 <- as.data.frame(cor(infection16dpp, MEs16Lables, use = "p"))
MMPvalue16 <- as.data.frame(corPvalueStudent(as.matrix(geneModuleMembership16), 21));
```

look for genes of interest in module 4 and 7:

Bring in contrast data from DESeq2 analysis comparing 16_Inoc_T2 vs 8_Inoc_T2

```
resT2 <- as.data.frame(lfcShrink(dds, contrast = c("condition", "16dpp_T2_Inoc", "8dpp_T2_Inoc"))) %>%
              mutate(gene = rownames(.))
resT2Cont <- as.data.frame(lfcShrink(dds, contrast = c("condition", "16dpp_T2_Inoc", "16dpp_T2_Cont")))
              mutate(gene = rownames(.))
```

```
resT4 <- as.data.frame(lfcShrink(dds, contrast = c("condition", "16dpp_T4_Inoc", "8dpp_T4_Inoc"))) %>%
              mutate(gene = rownames(.))
resT4Cont <- as.data.frame(lfcShrink(dds, contrast = c("condition", "16dpp_T4_Inoc", "16dpp_T4_Cont")))
              mutate(gene = rownames(.))
```

```
resT2 <- inner_join(resT2, resT2Cont, by = "gene", suffix = c("_T2vs8", "_T2vsC"))
resT4 <- inner_join(resT4, resT4Cont, by = "gene", suffix = c("_T4vs8", "_T4vsC"))

resultsEarly <- inner_join(resT2, resT4)
```

```
geneMod4 <-
  tibble(gene = colnames(infection16dpp),
         moduleLable = moduleLabels16) %>%
  mutate(
    modMembership = abs(geneModuleMembership16[gene, "ME4"]),
    modMemAdjPval = p.adjust(p = MMPvalue16$ME4, method = "BH") # [moduleLabels16 == 4]
  ) %>%
  left_join(resultsEarly)

geneMod7 <-
  tibble(gene = colnames(infection16dpp),
         moduleLable = moduleLabels16) %>%
  mutate(
    modMembership = abs(geneModuleMembership16[gene, "ME7"]),
    modMemAdjPval = p.adjust(p = MMPvalue16$ME7, method = "BH") # [moduleLabels16 == 7]
  ) %>%
    left_join(resultsEarly)
```

```
geneMod4_filt <- geneMod4 %>%
  mutate(passT2 = (padj_T2vs8 < 0.05 &
                     padj_T2vsC < 0.05 &
                     log2FoldChange_T2vs8 >= 1 &
                     log2FoldChange_T2vsC >= 1),
         passT4 = (padj_T4vs8 < 0.05 &
                     padj_T4vsC < 0.05 &
                     log2FoldChange_T4vs8 >= 1 &
                     log2FoldChange_T4vsC >= 1)
         ) %>%
  filter(passT2 == TRUE | passT4 == TRUE) %>%
  filter(modMembership >= 0.75) %>%
  #filter(modMembership >= quantile(geneMod4$modMembership, 0.75)) %>%
```

```r
  dplyr::select(-contains("lfcSE"),
                -contains("stat"),
                -contains("pvalue"),
                -contains("baseMean")) %>%
  arrange(desc(modMembership)) %>%
  rowwise() %>%
  mutate(Annotation = paste(html_text(html_nodes(read_html(paste0("http://cucurbitgenomics.org/feature/g
    ) %>%
  separate(Annotation, into = c("Match Name", "E-value", "Identity", "Description"), sep = "__")

geneMod7_filt <- geneMod7 %>%
  mutate(passT2 = (padj_T2vs8 < 0.05 &
                     padj_T2vsC < 0.05 &
                     log2FoldChange_T2vs8 >= 1 &
                     log2FoldChange_T2vsC >= 1),
         passT4 = (padj_T4vs8 < 0.05 &
                     padj_T4vsC < 0.05 &
                     log2FoldChange_T4vs8 >= 1 &
                     log2FoldChange_T4vsC >= 1)
         ) %>%
  filter(passT2 == TRUE | passT4 == TRUE) %>%
  filter(modMembership >= 0.75) %>%
  #filter(modMembership >= quantile(geneMod4$modMembership, 0.75)) %>%
  dplyr::select(-contains("lfcSE"),
                -contains("stat"),
                -contains("pvalue"),
                -contains("baseMean")) %>%
  arrange(desc(modMembership)) %>%
  rowwise() %>%
  mutate(Annotation = paste(html_text(html_nodes(read_html(paste0("http://cucurbitgenomics.org/feature/g
    ) %>%
  separate(Annotation, into = c("Match Name", "E-value", "Identity", "Description"), sep = "__")


mod4and7GOIs <- bind_rows("Mod. R4" = geneMod4_filt, "Mod. R7" = geneMod7_filt, .id = "Module") %>%
    arrange(desc(modMembership)) %>%
    distinct(gene, .keep_all = TRUE) %>%
    arrange(Module, desc(modMembership))


kableExtra::kable(mod4and7GOIs %>%
                    dplyr::select(-`Match Name`, -`E-value`, -Identity, -Description), format="latex", b
  kable_styling(latex_options="scale_down")
```

| Module | gene | moduleLable | modMembership | modMemAdjPval | log2FoldChange_T2vs8 | padj_T2vs8 | log2FoldChange_T2vsC | padj_T2vsC | log2FoldChange_T4vs8 | padj_T4vs8 | log2FoldChange_T4vsC | padj_T4vsC | passT2 | passT4 |
|---|---|---|---|---|---|---|---|---|---|---|---|---|---|---|
| Mod. R4 | Csa3G706170 | 4 | 0.9600105 | 0.0000000 | 1.3898709 | 0.0029402 | 1.0593253 | 0.3940515 | 1.1399296 | 0.0239798 | 1.2171884 | 0.0485826 | FALSE | TRUE |
| Mod. R4 | Csa2G351540 | 4 | 0.9406524 | 0.0000004 | 1.3616668 | 0.0327769 | 2.1497868 | 0.0131642 | 0.4948059 | 0.5235623 | 2.5436807 | 0.0003579 | TRUE | FALSE |
| Mod. R4 | Csa3G845500 | 4 | 0.9380197 | 0.0000005 | 1.1012177 | 0.0061480 | 1.2105717 | 0.0498648 | -0.0102767 | 0.9883439 | 0.5686060 | 0.3766573 | TRUE | FALSE |
| Mod. R4 | Csa3G782680 | 4 | 0.9246104 | 0.0000015 | 1.5068388 | 0.0000775 | 1.3838731 | 0.0116152 | 0.8281223 | 0.0582116 | 0.7758144 | 0.1816469 | TRUE | FALSE |
| Mod. R4 | Csa4G009900 | 4 | 0.9169922 | 0.0000026 | 1.8250168 | 0.0002261 | 1.9299967 | 0.0064796 | 0.6570751 | 0.2588914 | 2.2912030 | 0.0000406 | TRUE | FALSE |
| Mod. R4 | Csa5G524780 | 4 | 0.9140939 | 0.0000031 | 1.2371015 | 0.0433276 | 2.2021755 | 0.0066488 | 0.7709451 | 0.2597889 | 1.9153035 | 0.0078657 | TRUE | FALSE |
| Mod. R4 | Csa4G640960 | 4 | 0.9081028 | 0.0000051 | 1.5021135 | 0.0275712 | 2.5788916 | 0.0036529 | -0.3130345 | 0.7307392 | 2.3283935 | 0.0021297 | TRUE | FALSE |
| Mod. R4 | Csa3G271380 | 4 | 0.9040334 | 0.0000066 | 1.7537840 | 0.0072234 | 2.1971300 | 0.0187068 | -0.2687449 | 0.7661782 | 1.7768127 | 0.0347712 | TRUE | FALSE |
| Mod. R4 | Csa1G025960 | 4 | 0.9012187 | 0.0000077 | 1.1674401 | 0.0057501 | 1.5402297 | 0.0066488 | -0.0530878 | 0.9362960 | 1.1137028 | 0.0383710 | TRUE | FALSE |
| Mod. R4 | Csa2G069200 | 4 | 0.8947864 | 0.0000112 | 1.6331850 | 0.0321952 | 2.7294085 | 0.0066488 | -0.5669738 | 0.5465553 | 1.3857524 | 0.1924334 | TRUE | FALSE |
| Mod. R4 | Csa5G471600 | 4 | 0.8877080 | 0.0000173 | 1.4826746 | 0.0008038 | 1.9405516 | 0.0007089 | -0.0874055 | 0.9016484 | 0.9185410 | 0.1495727 | TRUE | FALSE |
| Mod. R4 | Csa1G439830 | 4 | 0.8867294 | 0.0000182 | 1.6141040 | 0.0005676 | 1.7716575 | 0.0107740 | 1.3765261 | 0.0074272 | 1.6918006 | 0.0059780 | TRUE | TRUE |
| Mod. R4 | Csa1G086390 | 4 | 0.8815762 | 0.0000259 | 1.3412863 | 0.0024172 | 1.4832317 | 0.0272324 | -0.5783553 | 0.2649027 | 1.4668518 | 0.0140072 | TRUE | FALSE |
| Mod. R4 | Csa4G411390 | 4 | 0.8773160 | 0.0000336 | 1.5769626 | 0.0088696 | 2.0508358 | 0.0207870 | -0.1900954 | 0.8252679 | 1.4721343 | 0.0701171 | TRUE | FALSE |
| Mod. R4 | Csa3G219720 | 4 | 0.8749464 | 0.0000385 | 1.0142010 | 0.0012354 | 1.2465548 | 0.0066488 | 0.3525669 | 0.3673588 | 0.6780432 | 0.1615285 | TRUE | FALSE |
| Mod. R4 | Csa4G049610 | 4 | 0.8359295 | 0.0002425 | 1.7271720 | 0.0196407 | 2.3454202 | 0.0242726 | 0.5878543 | 0.5239993 | 2.4457724 | 0.0034939 | TRUE | FALSE |
| Mod. R4 | Csa6G367150 | 4 | 0.8080432 | 0.0007112 | 1.4268949 | 0.0063074 | 0.7363852 | 0.9997456 | 1.3328035 | 0.0126231 | 1.7390608 | 0.0060632 | FALSE | TRUE |
| Mod. R7 | Csa6G495000 | 7 | 0.9501139 | 0.0000001 | 1.6807367 | 0.0038978 | 0.4131100 | 0.9997456 | 1.4258427 | 0.0225707 | 1.9807971 | 0.0040090 | FALSE | TRUE |
| Mod. R7 | Csa6G213910 | 7 | 0.9414933 | 0.0000003 | 2.1449441 | 0.0003620 | 0.3153180 | 0.9997456 | 1.8827572 | 0.0035827 | 1.9741861 | 0.0076968 | FALSE | TRUE |
| Mod. R7 | Csa3G567330 | 7 | 0.9293113 | 0.0000007 | 1.7431791 | 0.0043160 | 2.5289635 | 0.0057249 | -0.3364053 | 0.6690821 | 2.7319807 | 0.0002753 | TRUE | FALSE |
| Mod. R7 | Csa7G432140 | 7 | 0.9252129 | 0.0000010 | 1.0221420 | 0.0053574 | -0.0479017 | 0.9997456 | 1.3115666 | 0.0002649 | 1.0393706 | 0.0173064 | FALSE | TRUE |
| Mod. R7 | Csa6G403620 | 7 | 0.9088402 | 0.0000036 | 1.2508920 | 0.0284654 | 0.4634498 | 0.9997456 | 1.1533441 | 0.0479600 | 1.7205324 | 0.0106230 | FALSE | TRUE |
| Mod. R7 | Csa2G070840 | 4 | 0.9030608 | 0.0000050 | 1.1974971 | 0.0076648 | 1.4792913 | 0.0242726 | 0.1655289 | 0.7898207 | 1.1374185 | 0.0526066 | TRUE | FALSE |
| Mod. R7 | Csa2G349630 | 7 | 0.8989141 | 0.0000066 | 3.9272827 | 0.0000000 | 1.2940690 | 0.4154108 | 1.2826325 | 0.0315018 | 3.2447890 | 0.0000001 | FALSE | TRUE |
| Mod. R7 | Csa6G454420 | 7 | 0.8743069 | 0.0000283 | 1.2303361 | 0.0300144 | 0.2987049 | 0.9997456 | 1.3393730 | 0.0130673 | 1.3635669 | 0.0438137 | FALSE | TRUE |
| Mod. R7 | Csa4G285730 | 1 | 0.8734286 | 0.0000296 | 2.3203664 | 0.0003727 | 0.1597129 | 0.9997456 | 1.7036611 | 0.0213874 | 1.9710580 | 0.0213608 | FALSE | TRUE |
| Mod. R7 | Csa6G517010 | 1 | 0.8656750 | 0.0000429 | 1.6339378 | 0.0315379 | 3.0025607 | 0.0346821 | 0.1919984 | 0.8481977 | 2.8248933 | 0.0034480 | TRUE | FALSE |
| Mod. R7 | Csa5G152820 | 7 | 0.8620518 | 0.0000507 | 1.7054469 | 0.0214431 | 1.5574696 | 0.5199601 | 1.6705263 | 0.0322478 | 2.2387310 | 0.0172186 | FALSE | TRUE |
| Mod. R7 | Csa6G501260 | 1 | 0.8551843 | 0.0000686 | 1.3755091 | 0.0209634 | 0.1235727 | 0.9997456 | 1.1799402 | 0.0450604 | 1.4858990 | 0.0394962 | FALSE | TRUE |
| Mod. R7 | Csa6G160180 | 1 | 0.8463722 | 0.0001033 | 0.7401645 | 0.1071162 | 1.0438717 | 0.2656611 | 1.7760578 | 0.0000230 | 1.5466169 | 0.0013964 | FALSE | TRUE |
| Mod. R7 | Csa6G431740 | 1 | 0.8315800 | 0.0001785 | 1.4421363 | 0.0191510 | 0.1356076 | 0.9997456 | 2.0566746 | 0.0004271 | 1.8116614 | 0.0086191 | FALSE | TRUE |
| Mod. R7 | Csa4G646190 | 1 | 0.8231660 | 0.0002280 | 1.1297512 | 0.0060468 | 0.3227404 | 0.9997456 | 1.0901826 | 0.0087498 | 1.6538751 | 0.0002312 | FALSE | TRUE |
| Mod. R7 | Csa2G270790 | 7 | 0.7918045 | 0.0006021 | 2.3212630 | 0.0104803 | 1.5130716 | 0.8083711 | 1.6459654 | 0.0262906 | 4.4452942 | 0.0003231 | FALSE | TRUE |
| Mod. R7 | Csa7G390060 | 1 | 0.7678290 | 0.0011451 | 0.6244622 | 0.3177034 | 0.0474841 | 0.9997456 | 1.3076313 | 0.0113320 | 1.3026763 | 0.0425571 | FALSE | TRUE |

```r
write.csv(mod4and7GOIs, file = "Modules_4_7_GOIs.csv")

kableExtra::kable(mod4and7GOIs %>%
                    dplyr::select(Module, gene, `Match Name`, `E-value`, Identity, Description),
                  format="latex", booktabs=TRUE) %>%
  kable_styling(latex_options="scale_down")
```

| Module | gene | Match Name | E-value | Identity | Description |
|---|---|---|---|---|---|
| Mod. R4 | Csa3G706170 | AT2G31945.1 | 1.2e-06 | 60.00 | unknown protein |
| Mod. R4 | Csa2G351540 | AT5G43260.1 | 1.8e-34 | 65.63 | chaperone protein dnaJ-related |
| Mod. R4 | Csa3G845500 | AT5G47910.1 | 0.0e+00 | 70.86 | respiratory burst oxidase homologue D |
| Mod. R4 | Csa3G782680 | AT3G11820.1 | 4.5e-124 | 74.13 | syntaxin of plants 121 |
| Mod. R4 | Csa4G009900 | AT5G61760.1 | 9.7e-95 | 56.89 | inositol polyphosphate kinase 2 beta |
| Mod. R4 | Csa5G524780 | AT5G14700.1 | 2.1e-88 | 48.35 | NAD(P)-binding Rossmann-fold superfamily protein |
| Mod. R4 | Csa4G640960 | AT5G42830.1 | 2.2e-143 | 55.16 | HXXXD-type acyl-transferase family protein |
| Mod. R4 | Csa3G271380 | AT1G13340.1 | 2.5e-56 | 37.40 | Regulator of Vps4 activity in the MVB pathway protein |
| Mod. R4 | Csa1G025960 | AT2G38470.1 | 2.8e-119 | 47.19 | WRKY DNA-binding protein 33 |
| Mod. R4 | Csa2G069200 | AT2G30490.1 | 8.9e-30 | 63.83 | cinnamate-4-hydroxylase |
| Mod. R4 | Csa5G471600 | AT1G14040.1 | 4.5e-309 | 66.50 | EXS (ERD1/XPR1/SYG1) family protein |
| Mod. R4 | Csa1G439830 | AT1G30040.1 | 9.9e-117 | 63.55 | gibberellin 2-oxidase |
| Mod. R4 | Csa1G086390 | AT2G48010.1 | 5.7e-211 | 62.46 | receptor-like kinase in in flowers 3 |
| Mod. R4 | Csa4G411390 | AT2G41640.1 | 2.7e-161 | 58.30 | Glycosyltransferase family 61 protein |
| Mod. R4 | Csa3G219720 | AT1G11050.1 | 6.6e-234 | 65.27 | Protein kinase superfamily protein |
| Mod. R4 | Csa4G049610 | AT4G11280.1 | 3.8e-186 | 65.29 | 1-aminocyclopropane-1-carboxylic acid (acc) synthase 6 |
| Mod. R4 | Csa6G367150 | AT1G69040.2 | 1.3e-193 | 75.33 | ACT domain repeat 4 |
| Mod. R7 | Csa6G495000 | AT5G67400.1 | 1.0e-137 | 71.43 | root hair specific 19 |
| Mod. R7 | Csa6G213910 | AT5G05340.1 | 6.2e-116 | 65.62 | Peroxidase superfamily protein |
| Mod. R7 | Csa3G567330 | AT5G13080.1 | 7.4e-44 | 71.79 | WRKY DNA-binding protein 75 |
| Mod. R7 | Csa7G432140 | AT1G80600.1 | 6.3e-175 | 74.75 | HOPW1-1-interacting 1 |
| Mod. R7 | Csa6G403620 | AT1G07040.1 | 1.0e-130 | 71.43 | unknown protein |
| Mod. R7 | Csa2G070840 | AT1G08860.1 | 1.9e-227 | 68.62 | Calcium-dependent phospholipid-binding Copine family protein |
| Mod. R7 | Csa2G349630 | AT1G67810.1 | 5.4e-53 | 55.22 | sulfur E2 |
| Mod. R7 | Csa6G454420 | AT5G06280.1 | 9.7e-14 | 38.26 | unknown protein |
| Mod. R7 | Csa4G285730 | AT5G06720.1 | 6.0e-95 | 57.63 | peroxidase 2 |
| Mod. R7 | Csa6G517010 | AT2G29990.1 | 4.2e-64 | 80.99 | alternative NAD(P)H dehydrogenase 2 |
| Mod. R7 | Csa5G152820 | AT4G25700.1 | 2.0e-97 | 77.78 | beta-hydroxylase 1 |
| Mod. R7 | Csa6G501260 | AT4G39640.1 | 6.6e-209 | 64.92 | gamma-glutamyl transpeptidase 1 |
| Mod. R7 | Csa6G160180 | AT1G05010.1 | 3.2e-141 | 74.77 | ethylene-forming enzyme |
| Mod. R7 | Csa6G431740 | AT5G19410.1 | 6.4e-213 | 59.39 | ABC-2 type transporter family protein |
| Mod. R7 | Csa4G646190 | AT5G11950.1 | 1.1e-97 | 78.97 | Putative lysine decarboxylase family protein |
| Mod. R7 | Csa2G270790 | AT4G35070.1 | 1.2e-35 | 37.04 | SBP (S-ribonuclease binding protein) family protein |
| Mod. R7 | Csa7G390060 | AT4G02860.1 | 6.9e-93 | 59.15 | Phenazine biosynthesis PhzC/PhzF protein |

```
kableExtra::kable(mod4and7GOIs %>%
                          dplyr::select(Module, gene, modMembership, Description, -passT2, -passT4, -`Match
                      format="latex", booktabs=TRUE, digits = 20) %>%
  kable_styling(latex_options="scale_down")
```

| Module | gene | modMembership | moduleLable | modMemAdjPval | log2FoldChange_T2vs8 | padj_T2vs8 | log2FoldChange_T2vsC | padj_T2vsC | log2FoldChange_T4vs8 | padj_T4vs8 | log2FoldChange_T4vsC | padj_T4vsC | passT2 | passT4 | Match Name | E-value | Identity | Description |
|---|---|---|---|---|---|---|---|---|---|---|---|---|---|---|---|---|---|---|
| Mod. R4 | Csa3G706170 | 0.9600105 | 4 | 3.430158e-08 | 1.3898709 | 2.940210e-03 | 1.05932525 | 0.3940514824 | 1.13992964 | 0.0239797533 | 1.2171884 | 4.858260e-02 | FALSE | TRUE | AT2G31945.1 | 1.2e-06 | 60.00 | unknown protein |
| Mod. R4 | Csa2G351540 | 0.9406524 | 4 | 4.167916e-07 | 1.3616668 | 3.277691e-02 | 2.14978676 | 0.0131641630 | 0.49480588 | 0.5235623271 | 2.5436807 | 3.578704e-04 | TRUE | FALSE | AT5G43260.1 | 1.8e-34 | 65.63 | chaperone protein dnaJ-related |
| Mod. R4 | Csa3G845500 | 0.9380197 | 4 | 5.480415e-07 | 1.1012177 | 6.148018e-03 | 1.21057169 | 0.0498648135 | -0.01027673 | 0.9883439439 | 0.5686060 | 3.766573e-01 | TRUE | FALSE | AT5G47910.1 | 0.0e+00 | 70.86 | respiratory burst oxidase homologue D |
| Mod. R4 | Csa3G782680 | 0.9246104 | 4 | 1.485907e-06 | 1.5068388 | 7.754051e-05 | 1.38387309 | 0.0116152162 | 0.82812232 | 0.0582115578 | 0.7758144 | 1.816469e-01 | TRUE | FALSE | AT3G11820.1 | 4.5e-124 | 74.13 | syntaxin of plants 121 |
| Mod. R4 | Csa4G009900 | 0.9169922 | 4 | 2.585717e-06 | 1.8250168 | 2.261208e-04 | 1.92999668 | 0.0064795667 | 0.65707514 | 0.2588913696 | 2.2912030 | 4.061742e-05 | TRUE | FALSE | AT5G61760.1 | 9.7e-95 | 56.89 | inositol polyphosphate kinase 2 beta |
| Mod. R4 | Csa5G524780 | 0.9140939 | 4 | 3.062732e-06 | 1.2371015 | 4.332757e-02 | 2.20217552 | 0.0066487908 | 0.77094506 | 0.2597889098 | 1.9153035 | 7.865657e-03 | TRUE | FALSE | AT5G14700.1 | 2.1e-88 | 48.35 | NAD(P)-binding Rossmann-fold superfamily protein |
| Mod. R4 | Csa4G640960 | 0.9081028 | 4 | 5.120081e-06 | 1.5021135 | 2.757125e-02 | 2.57889158 | 0.0036529132 | -0.31303452 | 0.7307392113 | 2.3283935 | 2.129703e-03 | TRUE | FALSE | AT5G42830.1 | 2.2e-143 | 55.16 | HXXXD-type acyl-transferase family protein |
| Mod. R4 | Csa3G271380 | 0.9040334 | 4 | 6.632221e-06 | 1.7537840 | 7.223442e-03 | 2.19713004 | 0.0187067777 | -0.26874492 | 0.7661782475 | 1.7768127 | 3.477124e-02 | TRUE | FALSE | AT1G13340.1 | 2.5e-56 | 37.40 | Regulator of Vps4 activity in the MVB pathway protein |
| Mod. R4 | Csa1G025960 | 0.9012187 | 4 | 7.653347e-06 | 1.1674401 | 5.750120e-03 | 1.54022975 | 0.0066487908 | -0.05308777 | 0.9362960481 | 1.1137028 | 3.837098e-02 | TRUE | FALSE | AT2G38470.1 | 2.8e-119 | 47.19 | WRKY DNA-binding protein 33 |
| Mod. R4 | Csa2G069200 | 0.8947864 | 4 | 1.124597e-05 | 1.6331850 | 3.219522e-02 | 2.72940853 | 0.0066487908 | -0.56697381 | 0.5465552602 | 1.3857524 | 1.924334e-01 | TRUE | FALSE | AT2G30490.1 | 8.9e-30 | 63.83 | cinnamate-4-hydroxylase |
| Mod. R4 | Csa5G471600 | 0.8877080 | 4 | 1.731260e-05 | 1.4826746 | 8.037743e-04 | 1.94055156 | 0.0007089183 | -0.08740545 | 0.9016483555 | 0.9185410 | 1.495727e-01 | TRUE | FALSE | AT1G14040.1 | 4.5e-309 | 66.50 | EXS (ERD1/XPR1/SYG1) family protein |
| Mod. R4 | Csa1G439830 | 0.8867294 | 4 | 1.823974e-05 | 1.6141040 | 5.675615e-04 | 1.77165753 | 0.0107740244 | 1.37652615 | 0.0074271878 | 1.6918006 | 5.978015e-03 | TRUE | TRUE | AT1G30040.1 | 9.9e-117 | 63.55 | gibberellin 2-oxidase |
| Mod. R4 | Csa1G086390 | 0.8815762 | 4 | 2.591417e-05 | 1.3412863 | 2.417233e-03 | 1.48323175 | 0.0272323673 | -0.57835534 | 0.2649027018 | 1.4668518 | 1.400716e-02 | TRUE | FALSE | AT2G48010.1 | 5.7e-211 | 62.46 | receptor-like kinase in in flowers 3 |
| Mod. R4 | Csa4G411390 | 0.8773160 | 4 | 3.356186e-05 | 1.5769626 | 8.869603e-03 | 2.05083579 | 0.0207869976 | -0.19009539 | 0.8252678743 | 1.4721343 | 7.011708e-02 | TRUE | FALSE | AT2G41640.1 | 2.7e-161 | 58.30 | Glycosyltransferase family 61 protein |
| Mod. R4 | Csa3G219720 | 0.8749464 | 4 | 3.852191e-05 | 1.0142010 | 1.235416e-03 | 1.24655478 | 0.0066487908 | 0.35256687 | 0.3673588219 | 0.6780432 | 1.615285e-01 | TRUE | FALSE | AT1G11050.1 | 6.6e-234 | 65.27 | Protein kinase superfamily protein |
| Mod. R4 | Csa4G049610 | 0.8359295 | 4 | 2.425107e-04 | 1.7271720 | 1.964067e-02 | 2.34542015 | 0.0242726257 | 0.58785427 | 0.5239992763 | 2.4457724 | 3.493920e-03 | TRUE | FALSE | AT4G11280.1 | 3.8e-186 | 65.29 | 1-aminocyclopropane-1-carboxylic acid (acc) synthase 6 |
| Mod. R4 | Csa6G367150 | 0.8080432 | 4 | 7.111629e-04 | 1.4268949 | 6.307412e-03 | 0.73638522 | 0.9997455713 | 1.33280352 | 0.0126230863 | 1.7390608 | 6.063222e-03 | FALSE | TRUE | AT1G69040.2 | 1.3e-193 | 75.33 | ACT domain repeat 4 |
| Mod. R7 | Csa6G495000 | 0.9501139 | 7 | 1.068336e-07 | 1.6807367 | 3.897840e-03 | 0.41311001 | 0.9997455713 | 1.42584269 | 0.0225707044 | 1.9807971 | 4.009014e-03 | FALSE | TRUE | AT5G67400.1 | 1.0e-137 | 71.43 | root hair specific 19 |
| Mod. R7 | Csa6G213910 | 0.9414933 | 7 | 2.528006e-07 | 2.1449441 | 3.619690e-04 | 0.31531796 | 0.9997455713 | 1.88275725 | 0.0035826632 | 1.9741861 | 7.696816e-03 | FALSE | TRUE | AT5G05340.1 | 6.2e-116 | 65.62 | Peroxidase superfamily protein |
| Mod. R7 | Csa3G567330 | 0.9293113 | 7 | 7.264591e-07 | 1.7431791 | 4.316036e-03 | 2.52896350 | 0.0057249280 | -0.33640533 | 0.6690821039 | 2.7319807 | 2.752580e-04 | TRUE | FALSE | AT5G13080.1 | 7.4e-44 | 71.79 | WRKY DNA-binding protein 75 |
| Mod. R7 | Csa7G432140 | 0.9252129 | 7 | 1.023973e-06 | 1.0221420 | 5.357446e-03 | -0.04790169 | 0.9997455713 | 1.31156664 | 0.0002648513 | 1.0393706 | 1.730635e-02 | FALSE | TRUE | AT1G80600.1 | 6.3e-175 | 74.75 | HOPW1-1-interacting 1 |
| Mod. R7 | Csa6G403620 | 0.9088402 | 7 | 3.611683e-06 | 1.2508920 | 2.846545e-02 | 0.46344979 | 0.9997455713 | 1.15334406 | 0.0479600430 | 1.7205324 | 1.062303e-02 | FALSE | TRUE | AT1G07040.1 | 1.0e-130 | 71.43 | unknown protein |
| Mod. R7 | Csa2G070840 | 0.9030608 | 4 | 5.025251e-06 | 1.1974971 | 7.664827e-03 | 1.47929126 | 0.0242726257 | 0.16552892 | 0.7898207059 | 1.1374185 | 5.260656e-02 | TRUE | FALSE | AT1G08860.1 | 1.9e-227 | 68.62 | Calcium-dependent phospholipid-binding Copine family protein |
| Mod. R7 | Csa2G349630 | 0.8989141 | 7 | 6.615034e-06 | 3.9272827 | 4.186760e-10 | 1.29406905 | 0.4154108420 | 1.28263246 | 0.0315018244 | 3.2447890 | 8.437974e-08 | FALSE | TRUE | AT1G67810.1 | 5.4e-53 | 55.22 | sulfur E2 |
| Mod. R7 | Csa6G454420 | 0.8743069 | 7 | 2.826254e-05 | 1.2303361 | 3.001439e-02 | 0.29870486 | 0.9997455713 | 1.33937296 | 0.0130673427 | 1.3635669 | 4.381373e-02 | FALSE | TRUE | AT5G06280.1 | 9.7e-14 | 38.26 | unknown protein |
| Mod. R7 | Csa4G285730 | 0.8734286 | 1 | 2.962155e-05 | 2.3203664 | 3.726559e-04 | 0.15971293 | 0.9997455713 | 1.70366114 | 0.0213874463 | 1.9710580 | 2.136078e-02 | FALSE | TRUE | AT5G06720.1 | 6.0e-95 | 57.63 | peroxidase 2 |
| Mod. R7 | Csa6G517010 | 0.8656750 | 1 | 4.285083e-05 | 1.6339378 | 3.153789e-02 | 3.00256072 | 0.0346820738 | 0.19199843 | 0.8481977041 | 2.8248933 | 3.448031e-03 | TRUE | FALSE | AT2G29990.1 | 4.2e-64 | 80.99 | alternative NAD(P)H dehydrogenase 2 |
| Mod. R7 | Csa5G152820 | 0.8620518 | 7 | 5.070049e-05 | 1.7054469 | 2.144307e-02 | 1.55746963 | 0.5199600790 | 1.67052628 | 0.0322477640 | 2.2387310 | 1.721862e-02 | FALSE | TRUE | AT4G25700.1 | 2.0e-97 | 77.78 | beta-hydroxylase 1 |
| Mod. R7 | Csa6G501260 | 0.8551843 | 1 | 6.855180e-05 | 1.3755091 | 2.096341e-02 | 0.12357271 | 0.9997455713 | 1.17994024 | 0.0450604045 | 1.4858990 | 3.949623e-02 | FALSE | TRUE | AT4G39640.1 | 6.6e-209 | 64.92 | gamma-glutamyl transpeptidase 1 |
| Mod. R7 | Csa6G160180 | 0.8463722 | 1 | 1.033099e-04 | 0.7401645 | 1.071162e-01 | 1.04387167 | 0.2656611196 | 1.77605779 | 0.0000229799 | 1.5466169 | 1.396397e-03 | FALSE | TRUE | AT1G05010.1 | 3.2e-141 | 74.77 | ethylene-forming enzyme |
| Mod. R7 | Csa6G431740 | 0.8315800 | 1 | 1.785270e-04 | 1.4421363 | 1.915103e-02 | 0.13560759 | 0.9997455713 | 2.05667456 | 0.0004270713 | 1.8116614 | 8.619114e-03 | FALSE | TRUE | AT5G19410.1 | 6.4e-213 | 59.39 | ABC-2 type transporter family protein |
| Mod. R7 | Csa4G646190 | 0.8231660 | 1 | 2.280407e-04 | 1.1297512 | 6.046789e-03 | 0.32274039 | 0.9997455713 | 1.09018255 | 0.0087498062 | 1.6538751 | 2.311617e-04 | FALSE | TRUE | AT5G11950.1 | 1.1e-97 | 78.97 | Putative lysine decarboxylase family protein |
| Mod. R7 | Csa2G270790 | 0.7918045 | 7 | 6.020873e-04 | 2.3212630 | 1.048026e-02 | 1.51307158 | 0.8083711230 | 1.64596538 | 0.0262905635 | 4.4452942 | 3.231415e-04 | FALSE | TRUE | AT4G35070.1 | 1.2e-35 | 37.04 | SBP (S-ribonuclease binding protein) family protein |
| Mod. R7 | Csa7G390060 | 0.7678290 | 1 | 1.145150e-03 | 0.6244622 | 3.177034e-01 | 0.04748414 | 0.9997455713 | 1.30763132 | 0.0113319883 | 1.3026763 | 4.255711e-02 | FALSE | TRUE | AT4G02860.1 | 6.9e-93 | 59.15 | Phenazine biosynthesis PhzC/PhzF protein |

Plot all genes identified as early expressed. Supplemental figure 6.

```
moduleCount <-
  as.data.frame(counts(dds[colnames(infection16dpp), ], normalized = TRUE)) %>%
```

```r
  mutate(gene = rownames(.),
         module = moduleColors16.custom,
         moduleLabel = moduleLabels16) %>%
  gather("sample", "counts",-gene, -module, -moduleLabel) %>%
  mutate(age = rep(dds$age, each = ncol(infection16dpp)),
         timepoint = rep(str_remove(dds$timepoint, "T"), each =  ncol(infection16dpp)),
         treatment = rep(dds$treatment, each =  ncol(infection16dpp))
  ) %>%
  left_join(., as_tibble(table(moduleColors16.custom)), by = c("module" = "moduleColors16.custom")) %>%
  mutate(timepoint = as.numeric(timepoint),
         module = fct_infreq(module),
         facetLabel = fct_infreq(paste0(moduleLabel, " (n=", n,")"))
  )


geneplot <- moduleCount %>%
  # filter(gene %in% mod4and7GOIs$gene) %>%
  right_join(., mod4and7GOIs, by = "gene") %>%
  mutate(gene = fct_reorder(gene, .x = modMembership, .fun = "max", .desc = TRUE)) %>%
  ggplot() +
  stat_summary(
    fun.y = mean,
    geom = "line",
    aes(
      x = timepoint,
      y = counts,
      color = age,
      linetype = treatment
    )
  ) +
  stat_summary(fun.data = "mean_se", geom = "errorbar",
    aes(
      x = timepoint,
      y = counts,
      group = paste(age, treatment)
    ),
    color = rep(rep(c("#00BFC4", "#00BFC4", "#F8766D", "#F8766D"), each = 7),
                length(unique(mod4and7GOIs$gene)))
  ) +
  scale_x_continuous(breaks = c(0, 2, 4, 8, 12, 18, 24)) +
  labs(x = "Hours post inoculation", y = "Normalized read count") +
  cowplot::theme_cowplot(font_size = 10) +
    theme(strip.text =         element_text(
        colour = "grey10",
        size = rel(0.8) #,
        #margin = margin(0.8 * 7, 0.8 * 7, 0.8 * 7, 0.8 * 7)
        ),
        legend.position = "bottom") +
    facet_wrap(Module ~ gene, scales = "free", ncol = 5) +
    guides(linetype = guide_legend(title="Treatment"),
         color = guide_legend(title = "Age")) +
  scale_linetype_discrete(labels = c("Control", "Inoculated")) +
  scale_color_discrete(labels = c("8 dpp", "16 dpp")) +
```

```
  theme(legend.position=c(0.85, 0.05),
        legend.box = 'vertical',
        legend.margin = margin(t = -5)) +
    cowplot::panel_border()

g <- ggplot_gtable(ggplot_build(geneplot))
stripr <- which(grepl('strip-t', g$layout$name) & grepl("gTree", g$grobs))


# fills <- names(sort(table(moduleColors8.custom), decreasing = T))[unlist(rev(split(1:length(fills), c
#
fills <- names(sort(table(moduleColors16.custom), decreasing = T))[c(rep(7, 14), rep(4, 2), rep(7, 3), 

k <- 1

for (i in stripr) {
    j <- which(grepl('rect', g$grobs[[i]]$grobs[[1]]$childrenOrder))
    g$grobs[[i]]$grobs[[1]]$children[[j]]$gp$fill <- fills[k]
    k <- k+1
}
grid::grid.draw(g)
```

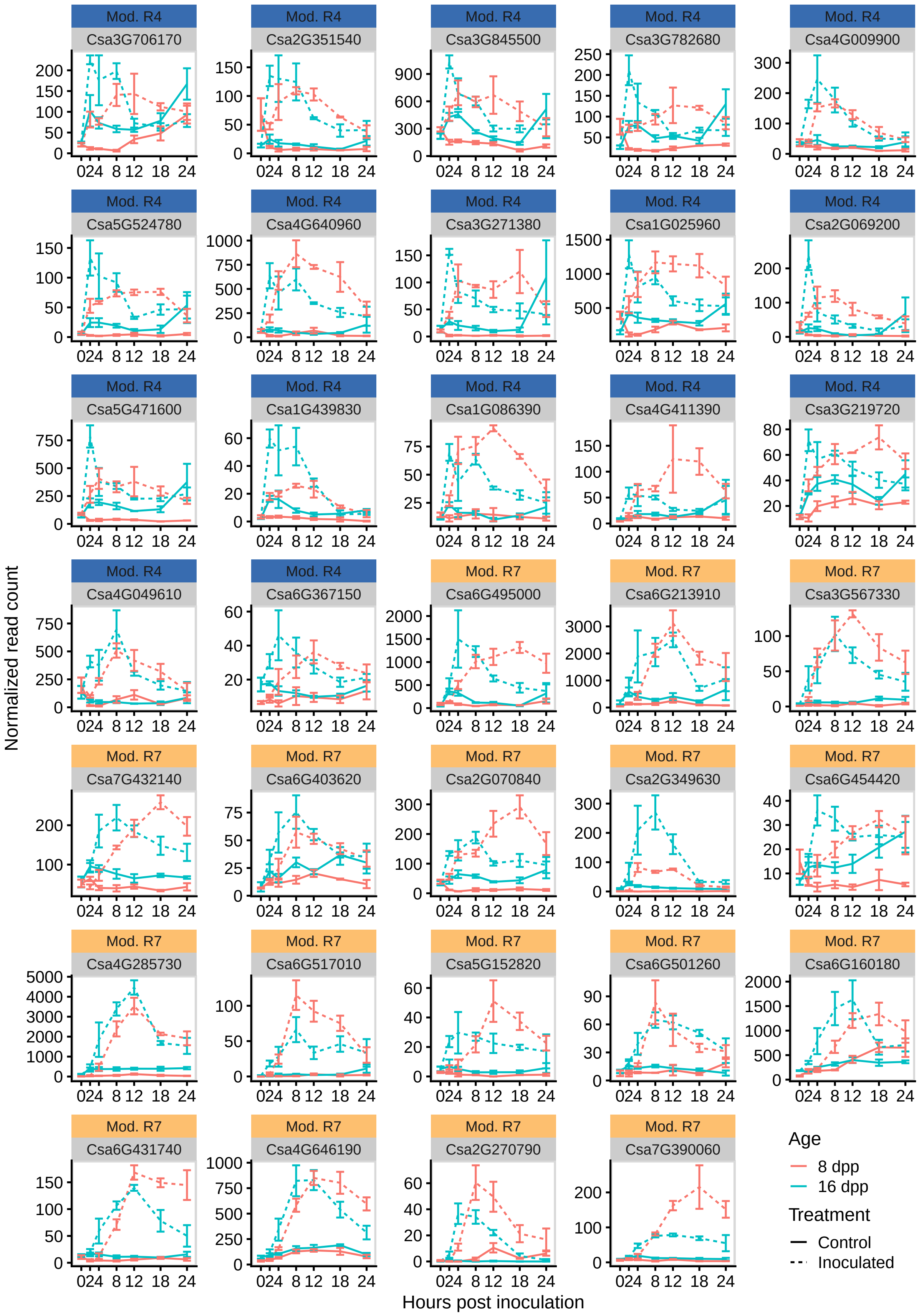
Mod. R4
Csa3G706170
Mod. R4
Csa2G351540
Mod. R4
Csa3G845500
Mod. R4
Csa3G782680
Mod. R4
Csa4G009900
Mod. R4
Csa5G524780
Mod. R4
Csa4G640960
Mod. R4
Csa3G271380
Mod. R4
Csa1G025960
Mod. R4
Csa2G069200
Mod. R4
Csa5G471600
Mod. R4
Csa1G439830
Mod. R4
Csa1G086390
Mod. R4
Csa4G411390
Mod. R4
Csa3G219720
Mod. R4
Csa4G049610
Mod. R4
Csa6G367150
Mod. R7
Csa6G495000
Mod. R7
Csa6G213910
Mod. R7
Csa3G567330
Mod. R7
Csa7G432140
Mod. R7
Csa6G403620
Mod. R7
Csa2G070840
Mod. R7
Csa2G349630
Mod. R7
Csa6G454420
Mod. R7
Csa4G285730
Mod. R7
Csa6G517010
Mod. R7
Csa5G152820
Mod. R7
Csa6G501260
Mod. R7
Csa6G160180
Mod. R7
Csa6G431740
Mod. R7
Csa4G646190
Mod. R7
Csa2G270790
Mod. R7
Csa7G390060
Normalized read count
Hours post inoculation
Age
8 dpp
16 dpp
Treatment
Control
Inoculated

Plot the subset of genes discussed in the discussion. Figure 9:

```
fd <- moduleCount %>%
    filter(gene %in% c("Csa1G025960", "Csa4G049610", "Csa6G160180",
                       "Csa4G285730", "Csa6G495000", "Csa3G845500",
                       "Csa3G271380", "Csa3G782680", "Csa6G431740",
                       "Csa1G086390", "Csa3G219720", "Csa4G009900",
                       "Csa2G069200", "Csa4G640960", "Csa4G411390")
    ) %>%
    mutate(
        gene = fct_relevel(
            gene,
            "Csa1G025960",
            "Csa4G049610",
            "Csa6G160180",
            "Csa4G285730",
            "Csa6G495000",
            "Csa3G845500",
            "Csa3G271380",
            "Csa3G782680",
            "Csa6G431740",
            "Csa1G086390",
            "Csa3G219720",
            "Csa4G009900",
            "Csa2G069200",
            "Csa4G640960",
            "Csa4G411390"
        ),
        `Annotation group` = fct_relevel(
            case_when(
                gene %in% c("Csa1G025960", "Csa4G049610", "Csa6G160180") ~ "Ethylene response",
                gene %in% c(
                    "Csa4G285730",
                    "Csa6G495000",
                    "Csa3G845500"
                ) ~ "Reactive oxygen metabolism",
                gene %in% c("Csa3G271380", "Csa3G782680", "Csa6G431740") ~ "Vesicle/Molecule transport"
                gene %in% c("Csa1G086390", "Csa3G219720", "Csa4G009900") ~ "Signal transduction",
                gene %in% c("Csa2G069200", "Csa4G640960", "Csa4G411390") ~ "Specialized metabolism"
            ),
            "Ethylene response",
            "Reactive oxygen metabolism",
            "Vesicle/Molecule transport",
            "Signal transduction",
            "Specialized metabolism"
        ),
        facet_order = case_when(
            gene == "Csa1G025960" ~ "Csa1G025960\nWRKY33",
            gene == "Csa4G049610" ~  "Csa4G049610\nACC synthase",
            gene == "Csa6G160180" ~ "Csa6G160180\nACC oxidase",
            gene == "Csa4G285730" ~ "Csa4G285730\nPeroxidase 2",
            gene == "Csa6G495000" ~  "Csa6G495000\nPeroxidase superfamily",
            gene == "Csa3G845500" ~  "Csa3G845500\nRespiratory burst oxidase homologue D",
            gene == "Csa3G271380" ~ "Csa3G271380\nIST1-LIKE 6",
```

```r
            gene == "Csa3G782680" ~  "Csa3G782680\nSyntaxin 121",
            gene == "Csa6G431740" ~ "Csa6G431740\nABC-2 type transporter",
            gene == "Csa1G086390" ~  "Csa1G086390\nRLK IN FLOWERS 3",
            gene == "Csa3G219720" ~ "Csa3G219720\nProtein kinase superfamily",
            gene == "Csa4G009900" ~  "Csa4G009900\n inositol polyphosphate kinase",
            gene == "Csa2G069200" ~ "Csa2G069200\nCinnamate-4-hydroxylase",
            gene == "Csa4G640960" ~  "Csa4G640960\nHXXXD-type acyl-transferase",
            gene == "Csa4G411390" ~ "Csa4G411390\nGlycosyltransferase family 61"
        ),
        facet_order = factor(
            facet_order,
            levels = c(
                "Csa1G025960\nWRKY33",
                "Csa4G049610\nACC synthase",
                "Csa6G160180\nACC oxidase",
                "Csa4G285730\nPeroxidase 2",
                "Csa6G495000\nPeroxidase superfamily",
                "Csa3G845500\nRespiratory burst oxidase homologue D",
                "Csa3G271380\nIST1-LIKE 6",
                "Csa3G782680\nSyntaxin 121",
                "Csa6G431740\nABC-2 type transporter",
                "Csa1G086390\nRLK IN FLOWERS 3",
                "Csa3G219720\nProtein kinase superfamily",
                "Csa4G009900\n inositol polyphosphate kinase",
                "Csa2G069200\nCinnamate-4-hydroxylase",
                "Csa4G640960\nHXXXD-type acyl-transferase",
                "Csa4G411390\nGlycosyltransferase family 61"
            )
        )
    )

pGeneExp <- fd %>%
  ggplot() +
  stat_summary(
    fun.y = mean,
    geom = "line",
    aes(
      x = timepoint,
      y = counts,
      color = age,
      linetype = treatment
    )
  ) +
  stat_summary(fun.data = "mean_se", geom = "errorbar",
    aes(
      x = timepoint,
      y = counts,
      group = paste(age, treatment)
    ),
    color = rep(rep(c("#00BFC4", "#00BFC4", "#F8766D", "#F8766D"), each = 7), 15)
  ) +
  scale_x_continuous(breaks = c(0, 2, 4, 8, 12, 18, 24)) +
  labs(x = "Hours post inoculation", y = "Normalized read count") +
```

```r
  cowplot::theme_cowplot(font_size = 10) +
    theme(strip.text =         element_text(
        colour = "grey10",
        size = rel(0.8) #,
        #margin = margin(0.8 * 7, 0.8 * 7, 0.8 * 7, 0.8 * 7)
        ),
        legend.position = "bottom") +
    facet_wrap(~ facet_order, ncol = 3,
               scales = "free", drop = FALSE) + #paste(gene, Description, sep = "\n")
    guides(linetype = guide_legend(title="Treatment"),
         color = guide_legend(title = "Age")) +
  scale_linetype_discrete(labels = c("Control", "Inoculated")) +
  scale_color_discrete(labels = c("8 dpp", "16 dpp")) +
  # theme(legend.position = c(0.75, 0.15),
  #       legend.box = 'vertical',
  #       legend.margin = margin(t = -5)) +
    cowplot::panel_border()

pGeneExp
```

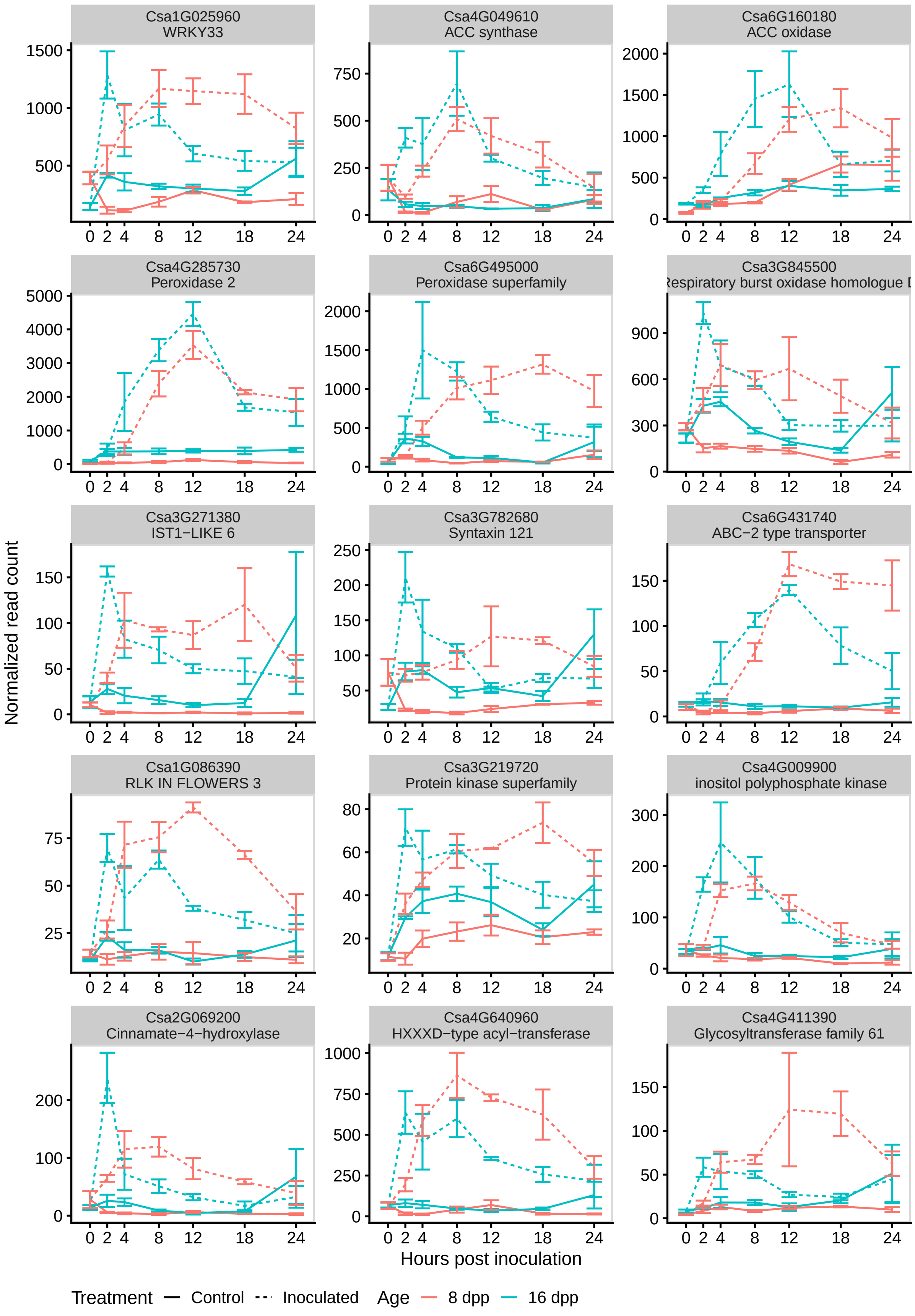
Csa1G025960
WRKY33
Csa4G049610
ACC synthase
Csa6G160180
ACC oxidase
Csa4G285730
Peroxidase 2
Csa6G495000
Peroxidase superfamily
Csa3G845500
Respiratory burst oxidase homologue D
Csa3G271380
IST1−LIKE 6
Csa3G782680
Syntaxin 121
Csa6G431740
ABC−2 type transporter
Csa1G086390
RLK IN FLOWERS 3
Csa3G219720
Protein kinase superfamily
Csa4G009900
inositol polyphosphate kinase
Csa2G069200
Cinnamate−4−hydroxylase
Csa4G640960
HXXXD−type acyl−transferase
Csa4G411390
Glycosyltransferase family 61
Normalized read count
Hours post inoculation
Treatment — Control - - Inoculated
Age — 8 dpp — 16 dpp

```
library(grid)
library(gtable)
z <- ggplotGrob(pGeneExp)
z <- gtable_add_rows(z, unit(13/14, "line"), 6)
z <- gtable_add_grob(z,
                     list(rectGrob(gp = gpar(col=NA, fill = gray(0.80), size = 0.5)),
                          textGrob("Ethylene response",
                                   rot = 0, gp = gpar(col = gray(0),fontsize=12))),
                     7, 5, 7,14, name = paste(runif(2)))
z <- gtable_add_rows(z, unit(13/14, "line"), 12)
z <- gtable_add_grob(z,
                     list(rectGrob(gp = gpar(col=NA, fill = gray(0.80), size = 0.5,face="bold")),
                          textGrob("Reactive oxygen metabolism",
                                   rot = 0, gp = gpar(col = gray(0),fontsize=12))),
                     13, 5, 13, 14, name = paste(runif(2)))

z <- gtable_add_rows(z, unit(13/14, "line"), 18)
z <- gtable_add_grob(z,
                     list(rectGrob(gp = gpar(col=NA, fill = gray(0.80), size = 0.5)),
                          textGrob("Vesicle/Molecule transport",
                                   rot = 0, gp = gpar(col = gray(0),fontsize=12))),
                     19, 5, 19, 14, name = paste(runif(2)))
z <- gtable_add_rows(z, unit(13/14, "line"), 24)
z <- gtable_add_grob(z,
                     list(rectGrob(gp = gpar(col=NA, fill = gray(0.80), size = 0.5)),
                          textGrob("Signal transduction",
                                   rot = 0, gp = gpar(col = gray(0),fontsize=12))),
                     25, 5, 25, 14, name = paste(runif(2)))
z <- gtable_add_rows(z, unit(13/14, "line"), 30)
z <- gtable_add_grob(z,
                     list(rectGrob(gp = gpar(col=NA, fill = gray(0.80), size = 0.5)),
                          textGrob("Specialized metabolism",
                                   rot = 0, gp = gpar(col = gray(0),fontsize=12))),
                     31, 5, 31, 14, name = paste(runif(2)))

grid.newpage()
grid.draw(z)
```

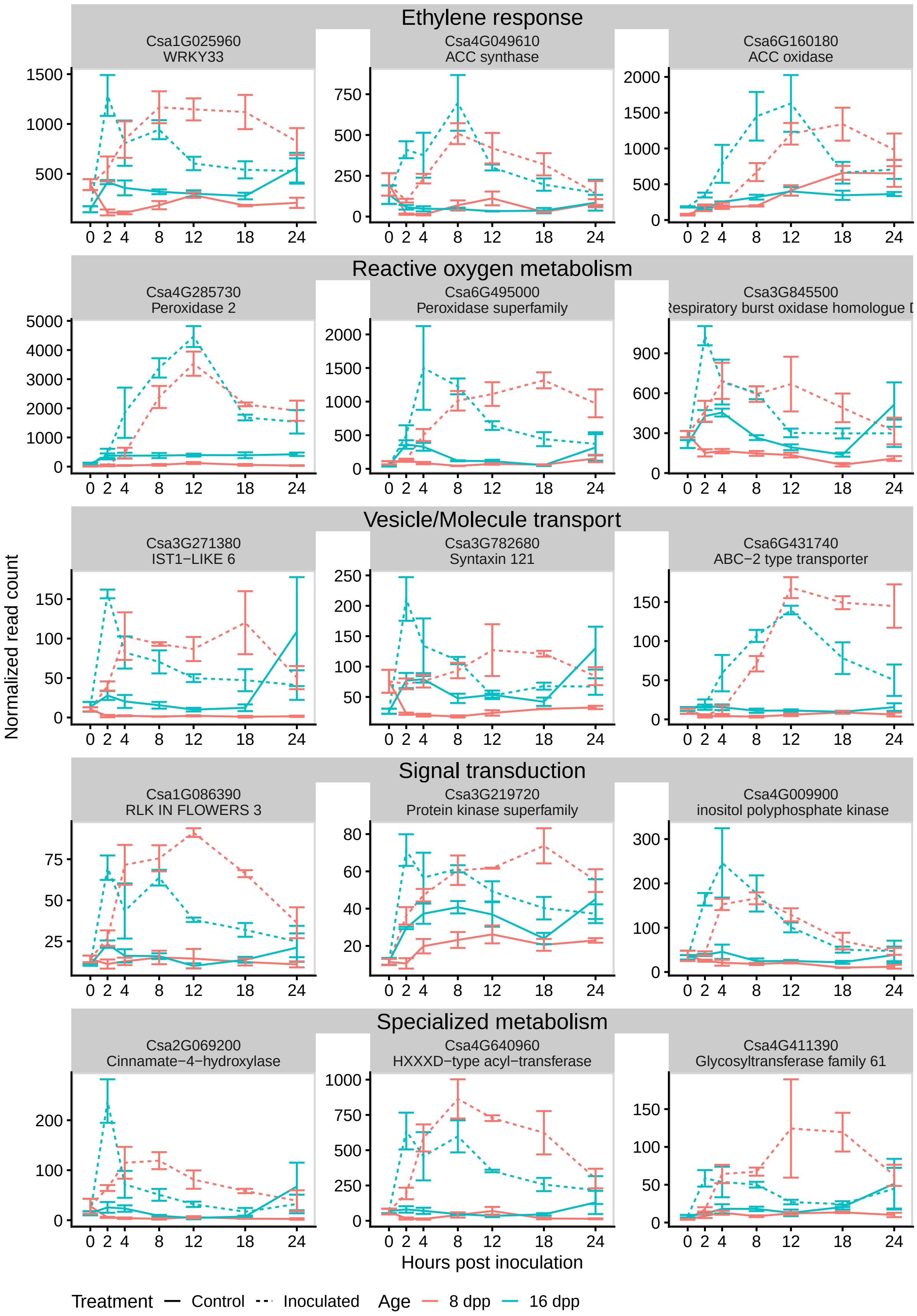
Ethylene response
Csa1G025960
WRKY33
Csa4G049610
ACC synthase
Csa6G160180
ACC oxidase
Reactive oxygen metabolism
Csa4G285730
Peroxidase 2
Csa6G495000
Peroxidase superfamily
Csa3G845500
Respiratory burst oxidase homologue D
Vesicle/Molecule transport
Csa3G271380
IST1−LIKE 6
Csa3G782680
Syntaxin 121
Csa6G431740
ABC−2 type transporter
Signal transduction
Csa1G086390
RLK IN FLOWERS 3
Csa3G219720
Protein kinase superfamily
Csa4G009900
inositol polyphosphate kinase
Specialized metabolism
Csa2G069200
Cinnamate−4−hydroxylase
Csa4G640960
HXXXD−type acyl−transferase
Csa4G411390
Glycosyltransferase family 61
Normalized read count
Hours post inoculation
Treatment
Control
Inoculated
Age
8 dpp
16 dpp

```r
pdf(file = "fig9.pdf", width = 7.5, height = 10)
    grid.draw(z)
dev.off()
```

```
## pdf
##   2
```

Plot the subset of genes discussed in the discussion from data in RNAseq exp1. Supp. Fig. 7:

```r
exp1Count <-
  as.data.frame(counts(dds_exp1, normalized = TRUE)) %>%
  mutate(gene = rownames(.),
         # module = moduleColors16.custom,
         # moduleLabel = moduleLabels16
         ) %>%
  gather("sample", "counts",-gene
         # , -module, -moduleLabel
          ) %>%
  mutate(age = rep(dds_exp1$age, each = nrow(dds_exp1)),
         timepoint = rep(str_remove(dds_exp1$timepoint, "T"), each =  nrow(dds_exp1)),
         timepoint = fct_relevel(timepoint, "0", "4", "24", "48")
  )

GOIs <- exp1Count %>%
    filter(gene %in% c("Csa1G025960", "Csa4G049610", "Csa6G160180",
                       "Csa4G285730", "Csa6G495000", "Csa3G845500",
                       "Csa3G271380", "Csa3G782680", "Csa6G431740",
                       "Csa1G086390", "Csa3G219720", "Csa4G009900",
                       "Csa2G069200", "Csa4G640960", "Csa4G411390")
    ) %>%
    mutate(
        gene = fct_relevel(
            gene,
            "Csa1G025960",
            "Csa4G049610",
            "Csa6G160180",
            "Csa4G285730",
            "Csa6G495000",
            "Csa3G845500",
            "Csa3G271380",
            "Csa3G782680",
            "Csa6G431740",
            "Csa1G086390",
            "Csa3G219720",
            "Csa4G009900",
            "Csa2G069200",
            "Csa4G640960",
            "Csa4G411390"
        ),
        `Annotation group` = fct_relevel(
            case_when(
                gene %in% c("Csa1G025960", "Csa4G049610", "Csa6G160180") ~ "Ethylene response",
                gene %in% c(
                    "Csa4G285730",
                    "Csa6G495000",
```

```
                    "Csa3G845500"
                ) ~ "Reactive oxygen metabolism",
                gene %in% c("Csa3G271380", "Csa3G782680", "Csa6G431740") ~ "Vesicle/Molecule transport"
                gene %in% c("Csa1G086390", "Csa3G219720", "Csa4G009900") ~ "Signal transduction",
                gene %in% c("Csa2G069200", "Csa4G640960", "Csa4G411390") ~ "Specialized metabolism"
            ),
            "Ethylene response",
            "Reactive oxygen metabolism",
            "Vesicle/Molecule transport",
            "Signal transduction",
            "Specialized metabolism"
        ),
        facet_order = case_when(
            gene == "Csa1G025960" ~ "Csa1G025960\nWRKY33",
            gene == "Csa4G049610" ~  "Csa4G049610\nACC synthase",
            gene == "Csa6G160180" ~ "Csa6G160180\nACC oxidase",
            gene == "Csa4G285730" ~ "Csa4G285730\nPeroxidase 2",
            gene == "Csa6G495000" ~  "Csa6G495000\nPeroxidase superfamily",
            gene == "Csa3G845500" ~  "Csa3G845500\nRespiratory burst oxidase homologue D",
            gene == "Csa3G271380" ~ "Csa3G271380\nIST1-LIKE 6",
            gene == "Csa3G782680" ~  "Csa3G782680\nSyntaxin 121",
            gene == "Csa6G431740" ~ "Csa6G431740\nABC-2 type transporter",
            gene == "Csa1G086390" ~  "Csa1G086390\nRLK IN FLOWERS 3",
            gene == "Csa3G219720" ~ "Csa3G219720\nProtein kinase superfamily",
            gene == "Csa4G009900" ~  "Csa4G009900\n inositol polyphosphate kinase",
            gene == "Csa2G069200" ~ "Csa2G069200\nCinnamate-4-hydroxylase",
            gene == "Csa4G640960" ~  "Csa4G640960\nHXXXD-type acyl-transferase",
            gene == "Csa4G411390" ~ "Csa4G411390\nGlycosyltransferase family 61"
        ),
        facet_order = factor(
            facet_order,
            levels = c(
                "Csa1G025960\nWRKY33",
                "Csa4G049610\nACC synthase",
                "Csa6G160180\nACC oxidase",
                "Csa4G285730\nPeroxidase 2",
                "Csa6G495000\nPeroxidase superfamily",
                "Csa3G845500\nRespiratory burst oxidase homologue D",
                "Csa3G271380\nIST1-LIKE 6",
                "Csa3G782680\nSyntaxin 121",
                "Csa6G431740\nABC-2 type transporter",
                "Csa1G086390\nRLK IN FLOWERS 3",
                "Csa3G219720\nProtein kinase superfamily",
                "Csa4G009900\n inositol polyphosphate kinase",
                "Csa2G069200\nCinnamate-4-hydroxylase",
                "Csa4G640960\nHXXXD-type acyl-transferase",
                "Csa4G411390\nGlycosyltransferase family 61"
            )
        )
    )


GOIs %>%
```

```
ggplot() +
stat_summary(
  fun.y = mean,
  geom = "line",
  aes(
    x = timepoint,
    y = counts,
    color = age,
    group = age
  )
) +
stat_summary(fun.data = "mean_se", geom = "errorbar",
  aes(
    x = timepoint,
    y = counts,
    color = age,
    group = age
  ), width = 0.2
) +
# scale_x_continuous(breaks = c(0, 2, 4, 8, 12, 18, 24)) +
labs(x = "Hours post inoculation", y = "Normalized read count") +
cowplot::theme_cowplot(font_size = 10) +
  theme(strip.text =         element_text(
      colour = "grey10",
      size = rel(0.8) #,
      #margin = margin(0.8 * 7, 0.8 * 7, 0.8 * 7, 0.8 * 7)
      ),
      legend.position = "bottom") +
  facet_wrap(~ facet_order, ncol = 3,
             scales = "free", drop = FALSE) + #paste(gene, Description, sep = "\n")
  guides(color = guide_legend(title = "Age")) +
scale_linetype_discrete(labels = c("Control", "Inoculated")) +
scale_color_discrete(labels = c("8 dpp", "16 dpp")) +
# theme(legend.position = c(0.75, 0.15),
#       legend.box = 'vertical',
#       legend.margin = margin(t = -5)) +
  cowplot::panel_border()
```

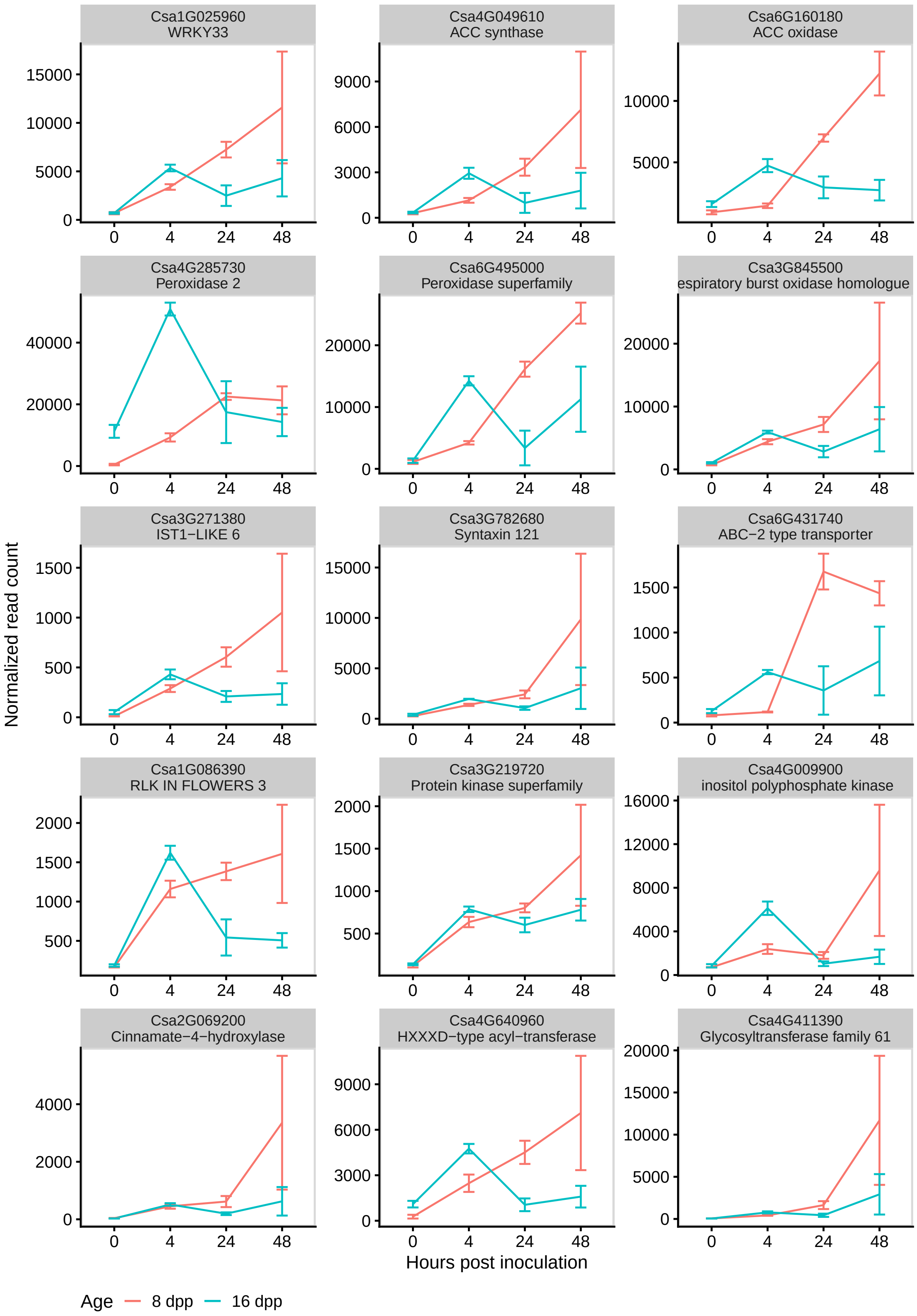

Csa1G025960
WRKY33
Csa4G049610
ACC synthase
Csa6G160180
ACC oxidase
Csa4G285730
Peroxidase 2
Csa6G495000
Peroxidase superfamily
Csa3G845500
espiratory burst oxidase homologue
Csa3G271380
IST1−LIKE 6
Csa3G782680
Syntaxin 121
Csa6G431740
ABC−2 type transporter
Csa1G086390
RLK IN FLOWERS 3
Csa3G219720
Protein kinase superfamily
Csa4G009900
inositol polyphosphate kinase
Csa2G069200
Cinnamate−4−hydroxylase
Csa4G640960
HXXXD−type acyl−transferase
Csa4G411390
Glycosyltransferase family 61
Normalized read count
Hours post inoculation
Age
8 dpp
16 dpp

Correlation of the subset of genes discussed in the discussion between Exp1 and 2. Supp. Fig. 8:

```
moduleCount %>%
    filter(timepoint %in% c(0, 4, 24),
           treatment == "Inoc" | timepoint == 0
           ) %>%
    filter(gene %in% c(
            "Csa1G025960",
            "Csa4G049610",
            "Csa6G160180",
            "Csa4G285730",
            "Csa6G495000",
            "Csa3G845500",
            "Csa3G271380",
            "Csa3G782680",
            "Csa6G431740",
            "Csa1G086390",
            "Csa3G219720",
            "Csa4G009900",
            "Csa2G069200",
            "Csa4G640960",
            "Csa4G411390"
            )
    ) %>%
    mutate(timepoint = as.factor(timepoint)) %>%
    group_by(gene, timepoint, age) %>%
    summarize(avgCount = mean(counts)) %>%
    left_join(GOIs %>%
    group_by(gene, timepoint, age) %>%
    summarize(avgCount = mean(counts)), by = c("gene", "timepoint", "age")) %>%
    ggplot(aes(x = log10(avgCount.x), y = log10(avgCount.y))) +
        geom_point() +
        geom_smooth(method = "lm") +
        ggpubr::stat_cor(method = "pearson") +
        labs(x = "log10(Avg. Counts) - Exp2", y = "log10(Avg. Counts) - Exp1") +
    cowplot::theme_cowplot(font_size = 11) +
    theme(strip.text = element_text(
        colour = "grey10",
        size = rel(0.8),
        margin = margin(0.5 * 7, 0.5 * 7, 0.5 * 7, 0.5 * 7)
        )
    ) +
    theme(legend.position="bottom",
        legend.box = 'vertical',
        legend.margin = margin(t = -5)) +
    cowplot::panel_border()
```

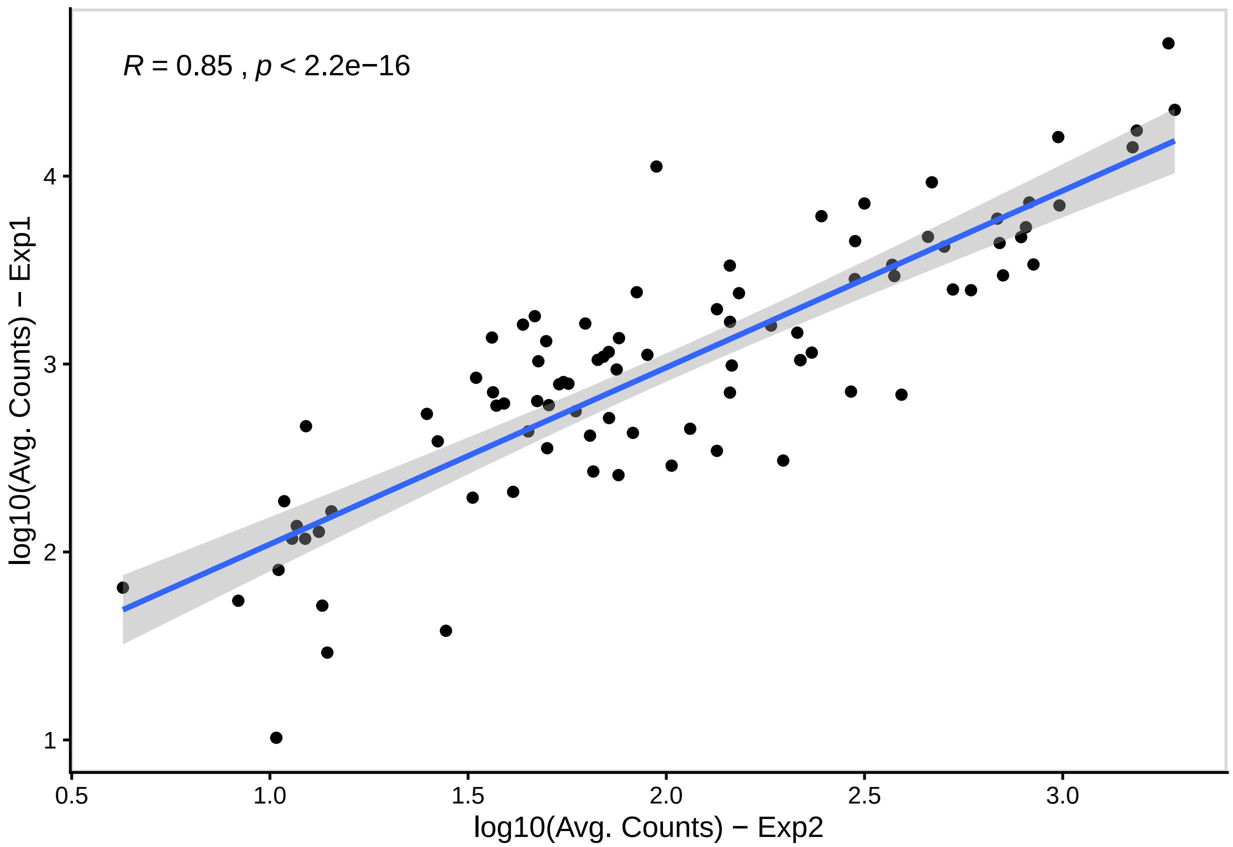


```
sessionInfo()
```

```
## R version 3.6.1 (2019-07-05)
## Platform: x86_64-w64-mingw32/x64 (64-bit)
## Running under: Windows 10 x64 (build 18362)
##
## Matrix products: default
##
## locale:
## [1] LC_COLLATE=English_United States.1252
## [2] LC_CTYPE=English_United States.1252
## [3] LC_MONETARY=English_United States.1252
## [4] LC_NUMERIC=C
## [5] LC_TIME=English_United States.1252
##
## attached base packages:
##  [1] grid      splines   stats4    parallel  stats     graphics  grDevices
##  [8] utils     datasets  methods   base
##
## other attached packages:
##  [1] gtable_0.3.0            writexl_1.2
##  [3] rgeos_0.5-2             sp_1.3-2
##  [5] ggdendro_0.1-20         magick_2.3
##  [7] readxl_1.3.1            rvest_0.3.5
##  [9] xml2_1.2.2              RColorBrewer_1.1-2
## [11] topGO_2.38.1            SparseM_1.78
```

```
## [13] GO.db_3.10.0               AnnotationDbi_1.48.0
## [15] graph_1.64.0                pheatmap_1.0.12
## [17] WGCNA_1.68                  fastcluster_1.1.25
## [19] dynamicTreeCut_1.63-1       ggalluvial_0.11.1
## [21] rjson_0.2.20                kableExtra_1.1.0
## [23] DESeq2_1.26.0               SummarizedExperiment_1.16.1
## [25] DelayedArray_0.12.1         BiocParallel_1.20.1
## [27] matrixStats_0.55.0          Biobase_2.46.0
## [29] GenomicRanges_1.38.0        GenomeInfoDb_1.22.0
## [31] IRanges_2.20.1              S4Vectors_0.24.1
## [33] BiocGenerics_0.32.0         tximport_1.14.0
## [35] UpSetR_1.4.0                forcats_0.4.0
## [37] stringr_1.4.0               dplyr_0.8.3
## [39] purrr_0.3.3                 readr_1.3.1
## [41] tidyr_1.0.0                 tibble_2.1.3
## [43] ggplot2_3.2.1               tidyverse_1.3.0
##
## loaded via a namespace (and not attached):
##   [1] backports_1.1.5        Hmisc_4.2-0            selectr_0.4-2
##   [4] plyr_1.8.5             lazyeval_0.2.2         robust_0.4-18.2
##   [7] digest_0.6.22          foreach_1.4.7          htmltools_0.4.0
##  [10] fansi_0.4.0            magrittr_1.5           checkmate_1.9.4
##  [13] memoise_1.1.0          fit.models_0.5-14      cluster_2.1.0
##  [16] doParallel_1.0.15      limma_3.42.2           annotate_1.64.0
##  [19] modelr_0.1.5           jpeg_0.1-8.1           colorspace_1.4-1
##  [22] ggrepel_0.8.1          blob_1.2.0             rrcov_1.5-2
##  [25] haven_2.2.0            xfun_0.11              crayon_1.3.4
##  [28] RCurl_1.95-4.12        jsonlite_1.6.1         genefilter_1.68.0
##  [31] zeallot_0.1.0          impute_1.60.0          survival_2.44-1.1
##  [34] iterators_1.0.12       glue_1.3.1             zlibbioc_1.32.0
##  [37] XVector_0.26.0         webshot_0.5.2          DEoptimR_1.0-8
##  [40] scales_1.0.0           mvtnorm_1.0-12         DBI_1.1.0
##  [43] Rcpp_1.0.2             viridisLite_0.3.0      xtable_1.8-4
##  [46] htmlTable_1.13.3       foreign_0.8-71         bit_1.1-14
##  [49] preprocessCore_1.48.0  Formula_1.2-3          htmlwidgets_1.5.1
##  [52] httr_1.4.1             ellipsis_0.3.0         acepack_1.4.1
##  [55] pkgconfig_2.0.3        XML_3.98-1.20          nnet_7.3-12
##  [58] dbplyr_1.4.2           utf8_1.1.4             locfit_1.5-9.1
##  [61] reshape2_1.4.3         labeling_0.3           tidyselect_0.2.5
##  [64] rlang_0.4.1            munsell_0.5.0          cellranger_1.1.0
##  [67] tools_3.6.1            cli_2.0.0              generics_0.0.2
##  [70] RSQLite_2.1.5          broom_0.5.3            evaluate_0.14
##  [73] yaml_2.2.0             knitr_1.26             bit64_0.9-7
##  [76] fs_1.3.1               robustbase_0.93-5      lemon_0.4.3
##  [79] nlme_3.1-140           compiler_3.6.1         rstudioapi_0.10
##  [82] curl_4.3               png_0.1-7              ggsignif_0.6.0
##  [85] reprex_0.3.0           geneplotter_1.64.0     pcaPP_1.9-73
##  [88] stringi_1.4.3          lattice_0.20-38        Matrix_1.2-17
##  [91] vctrs_0.2.0            pillar_1.4.3           lifecycle_0.1.0
##  [94] cowplot_1.0.0          data.table_1.12.6      bitops_1.0-6
##  [97] R6_2.4.1               latticeExtra_0.6-29    gridExtra_2.3
## [100] codetools_0.2-16       MASS_7.3-51.4          assertthat_0.2.1
## [103] withr_2.1.2            GenomeInfoDbData_1.2.2 hms_0.5.2
## [106] rpart_4.1-15           rmarkdown_2.0          ggpubr_0.2.4
```

```
## [109] ggnewscale_0.4.1       lubridate_1.7.4        base64enc_0.1-3
```
